# A 2D Fragment-Assisted Protein Mimetic Approach to Rescue α-Synuclein Aggregation Mediated Early and Post-Disease Parkinson’s Phenotypes

**DOI:** 10.1101/2022.07.11.499659

**Authors:** Nicholas H. Stillman, Johnson A. Joseph, Jemil Ahmed, Ryan A. Dohoney, Tyler D. Ball, Alexandra G. Thomas, Tessa C. Fitch, Courtney M. Donnelly, Sunil Kumar

**Affiliations:** Department of Chemistry and Biochemistry, University of Denver, Denver, CO 80210; The Knoebel Institute for Healthy Aging, University of Denver, Denver, CO 80210; Molecular and Cellular Biophysics Program, University of Denver, Denver, CO 80210

## Abstract

We have developed a Oligopyridylamide (OP) based 2-Dimensional Fragment-Assisted Structure-based Technique (2D-FAST) to identify potent antagonists of α-Synuclein (αS) aggregation, a process central to Parkinson’s disease (PD). The 2D-FAST utilizes a fragment-based screening of large chemical space in OPs, which led to the identification of NS132 as an antagonist of the multiple facets of αS aggregation. We also identified a better cell permeability analog (NS163) without sacrificing activity. OPs rescue αS aggregation mediated PD phenotypes in muscle cells and dopaminergic (DA) neurons in *C. elegans* models. OPs prevent the progression of PD phenotypes in a novel post-disease onset PD model.

This is one of the first examples of a synthetic mimetic-based 2D-FAST to identify antagonists of toxic αS self-assembly. We envision that 2D-FAST will have tremendous potential as it is expandable for other oligoamide scaffolds and for a much larger chemical space to identify lead therapeutics for various diseases.

## INTRODUCTION

Abberent protein-protein interactions (aPPIs) are associated with a plethora of pathological conditions, including infectious diseases, cancer, neurodegenerative diseases, and amyloid diseases^1–11^. Consequently, modulation of aPPIs is considered to be a promising therapeutic intervention toward various pathologies. The pathological aPPIs are mediated via specific chemical interactions that often sample dynamic and transient conformations, which spread over a large and hydrophobic surface^1–7^. One such example is the aggregation of αS, which is a neuronal protein expressed at high levels in DA neurons in the brain and implicated in the regulation of synaptic vesicle trafficking and recycling and neurotransmitter release^12–18^. The aggregation of αS is associated with the impaired DA neurons, which is a pathological hallmark of PD^12–18^. Therefore, one of the potential disease-modifying therapeutic strategies for PD is the modulation of αS aggregation^19–29^. A few small molecules have been shown to inhibit αS aggregation^19–29^ (ref. within 19); however, some of them have complex chemical structures, which might limit their ability for synthetic tuning and further optimization of the antagonist activity against αS aggregation. Also, protein mimetics have been identified to inhibit αS aggregation; however, the chemical space on them was limited and there was no systematic optimization carried out against αS aggregation^23–29^ Moreover, most of these ligands were not tested against PD phenotypes in DA neurons in *in vivo models* to further assess their therapeutic potential^23–25,28,29^. Therefore, ligands with the ability to manipulate aggregation with a large chemical space and having the tendency for systematic optimization of the antagonist activity could lead to potent antagonism of αS aggregation.

Oligopyridylmides (OPs) are a class of synthetic protein mimetics that have been shown to manipulate the aggregation of multiple proteins, including islet amyloid polypeptide^30–32^, Aβ peptide^33,34^, and mutant p53^35^, which are associated with type 2 diabetes (T2D), Alzheimer’s disease (AD), and cancer, respectively. OPs have a large surface area and synthetically tunable side-chain functionalities that can complement the topography and side-chain residues of proteins such as those present at the interfaces of aPPIs during protein aggregation^30–38^. However, the OP library used in the screening to identify antagonists of protein aggregation was moderate in size (~30 OPs) with limited chemical diversity (~10 side chains), which may have precluded the opportunity for the optimization of the antagonist activity of OPs against the aggregation of various amyloid proteins^30–37^. There were several limitations with the previous method to generate OP libraries with larger chemical space to identify antagonists for the aggregation of proteins^30–37^, including tedious synthesis with several chromatography steps (Fig. 1A) and pre-synthesized and screened OPs, which lacked a systematic optimization against the dynamic and transient nature of protein aggregation surfaces.^27–38^

**Fig. 1.**
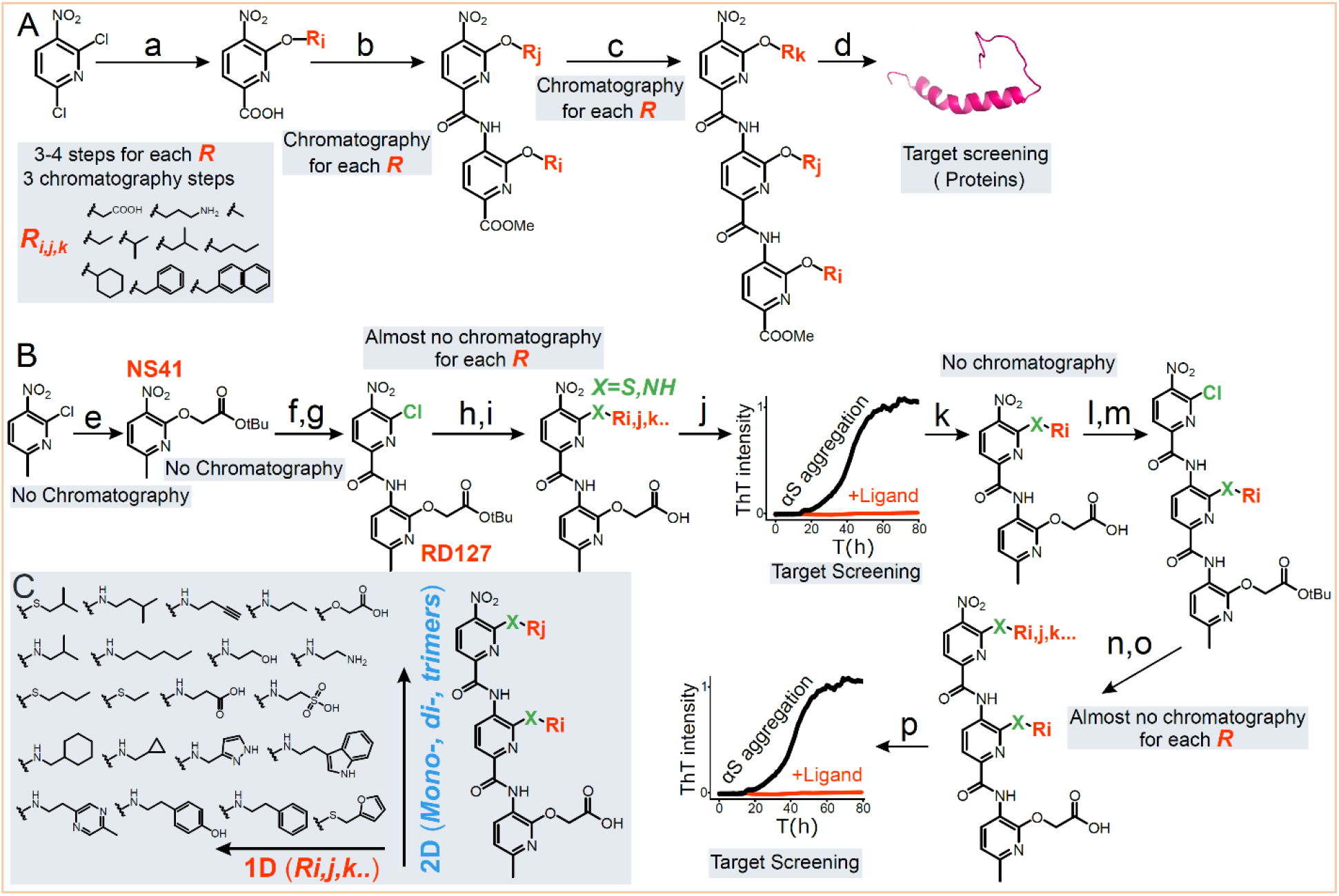
A schematic for the comparison of the old method (A) and the novel 2D-FAST (B). **a**, Synthesis of the monopyridyls with various side chains (R_i,j,k_). (**Inset**) The chemical structures of the side chains on OPs. **b-d**, A flowchart for the synthesis of dipyridyls and tripyridyls and their testing against various biological targets. **e**, *Tert*-Butyl 2-hydroxyacetate, NaH (60% dispersion in mineral oil), toluene, 50 min. at 0 °C, then 5 h at r.t. **f**, **l**, Pd/C, H_2_ (g), EtOAc, 3 h at r.t. **g**, 6-chloro-5-nitropicolinic acid, DCM (anhydrous), triethylamine, thionyl chloride, 0 °C to r.t., 45 min. **h**, Primary amine/thiol, DIPEA, DCM, 3 h at r.t. **i**,**o**, TFA, DCM, TES, 3h, r.t. **j**, Screening of the dipyridyls against αS aggregation using high throughput ThT kinetic aggregation assay. **k**, The identification of the most potent dipyridyl antagonist of αS aggregation (X-Ri on the second side chain of the dipyridyl). **m**, triethylamine, 6-chloro-5-nitropicolinoyl chloride, THF (anhydrous) 0 °C to r.t., 40 min. **n**, Primary amine, DIPEA, DCM, 3 h at r.t. **p**, Screening of the tripyridyls against αS aggregation using ThT kinetic aggregation assay. **C**, The representation of the 2D-FAST approach.

We have developed a novel 2D-FAST by combining fragment and structure-based techniques into the OP scaffold in order to systematically optimize the antagonist activity against αS aggregation (Fig. 1B,C). The fragment-based approach has emerged as a promising method for drug discovery to identify high-affinity ligands against various pathological targets, including aPPIs^39–46^. The OP is an ideal scaffold for the fragment-based approach because the antagonist activity of OPs against their biological targets have been shown to increase with increasing side chains (monopyridyl<dipyridyl<tripyridyl)^32–35^. In 2D-FAST, the 2D consist of the side chains and the number of pyridyls groups in OPs (Fig. 1C). There are several novel features of our 2D-FAST for OPs, including (1) Use of common precursors for the elongation of OP from mono-to di-to tripyridyl synthesis; (2) Use of a chromatography-free amide coupling method for the elongation of the backbone chain; (3) Introduction of a large chemically diverse library of side chains on OPs with no column chromatography for most of the products; (4) Use of a fragment-based approach for systematic optimization of the antagonist activity of OPs against αS aggregation.

Using the synthetic 2D-FAST and an array of biophysical and cellular assays, we have identified NS132 as the most potent antagonist of *de novo* and fibers catalyzed aggregation of αS. NS132 was able to wholly inhibit αS aggregation, even at a substoichiometric ratio (αS:NS132, 1:0.2). In contrast, the peptidomimetic approaches without the novel features of our 2D-FAST, have identified ligands that require 5-100 fold molar excess to inhibit the aggregation of αS^28,29^. This observation highlights the novel aspects of our 2D-FAST approach, which entails a systematic fragment-based screening of a very large chemical space against αS aggregation that allows the identification of a very potent antagonist. A structure-activity relationship demonstrated that the side chains of NS132 are essential for its antagonist activity. The HEK cells-based assays demonstrated that NS132 potently rescues cytotoxicity and inhibits the formation of intracellular inclusions. The 2D HSQC NMR study demonstrates that NS132 interacts with specific sequences of αS, which have been previously suggested to be the key aggregation-prone sequences^24,47^. The study also led to the synthesis of an analog of NS132 (NS163, Supplementary Fig. 2) with improved cell permeability without sacrificing the antagonist activity against αS aggregation. The antagonist activity of NS163 and NS132 was tested against αS aggregation-mediated PD phenotypes in two *C. elegans* PD models. Both ligands (NS163 and NS132) were very effective in rescuing various PD phenotypes in two *C. elegans* PD models, including neuroprotective effect against degeneration of DA neurons, motility recovery, improved food sensing behavioral deficits, increased dopamine synthesis, and reduced ROS level. Moreover, the OPs were very effective in rescuing further progression of PD phenotypes in DA neurons when administered in a post-disease-onset PD model, a model that mimics the current therapeutic intervention strategies, where the treatment starts during post-diagnosis of PD.

Overall, we have developed a novel 2D-FAST and demonstrated its utility in the identification of potent antagonists of αS aggregation, a process that is associated with PD. We have used a comprehensive study to establish the synthetic protein mimetic-based 2D-FAST approach and identified potent ligands, which were very effective in rescuing αS aggregation mediated PD phenotypes in physiologically relevant PD models.

## RESULTS

### Design and Synthesis of the 2D-FAST for OPs

Foldamers with carboxylic acid functional groups as side chains have been shown to effectively modulate αS aggregation and they partly mimic the topography of OPs^24,48^. Also, OPs with a minimum of two side chains (in dipyridyls) have been shown to achieve moderate antagonist activity against protein aggregation^31,33–35^. Therefore, for the 2D-FAST, we synthesized a library of dipyridyls with the carboxylic acid functional group as the first side chain and appended diverse chemical side chains on the second pyridyl position (Fig. 1B). The 2-chloro-6-methyl-3-nitropyridine was used as a precursor to synthesize the nitro *tert*-butyl protected carboxylic acid monopyridyl (NS41, Fig. 1e) using our novel chromatography free method. Subsequently, we synthesized the chloro-dipyridyl using a newly developed chromatography-free amide coupling in our lab (RD127, Fig. 1f,g, and Supplementary Fig. 1), which was used as a common precursor to synthesize a library of dipyridyls with diverse side chains using primary amines/thiols via a one-pot reaction (Fig. 1h). All reactions went to completion and a large number of dipyridyl products did not require column chromatography as the excess primary amines/thiols were evaporated on rotovap or lyophilizer. However, a few dipyridyls required column chromatography to separate them from the starting material side chains because of their very high boiling point (7 out of 21 dipyridyls required column, Supplementary Fig. 1).

### Biophysical characterization of OPs against the aggregation of αS

The dipyridyl library was screened against the aggregation of 100 μM αS (in 1 × PBS buffer) at an equimolar ratio using Thioflavin T (ThT) dye-based aggregation assay (Fig. 1j)^49^. The screening led to the identification of NS55 as the most potent antagonist as it was able to completely suppress the aggregation of αS (Figure 2a-d), reflected by a low ThT fluorescence signal. The inhibition of αS aggregation by NS55 was also confirmed by transmission electron microscopy (TEM) images, which show an abundance (Fig. 2e) and no (Fig. 2f) αS fibers in the absence and presence of NS55, respectively. Next, we used NS55 (dipyridyl) as a precursor to synthesize and generate a tripyridyl library because we have shown that tripyridyls are better antagonists than dipyridyls for various amyloid proteins^31,33–35^. Surprisingly, all OP dimers synthesized using primary thiols were agonists of αS aggregation; therefore, we did not pursue primary thiols for the synthesis of tripyridyl. We used similar synthetic steps to generate tripyridyls as we used to generate dipyridyls (Fig. 1l-o, and Supplementary Fig. 1). All reactions went to completion and most of the tripyridyls products did not require column chromatography (6 out of 15 tripyridyls required column, Supplementary Fig. 1).

**Fig.2.**
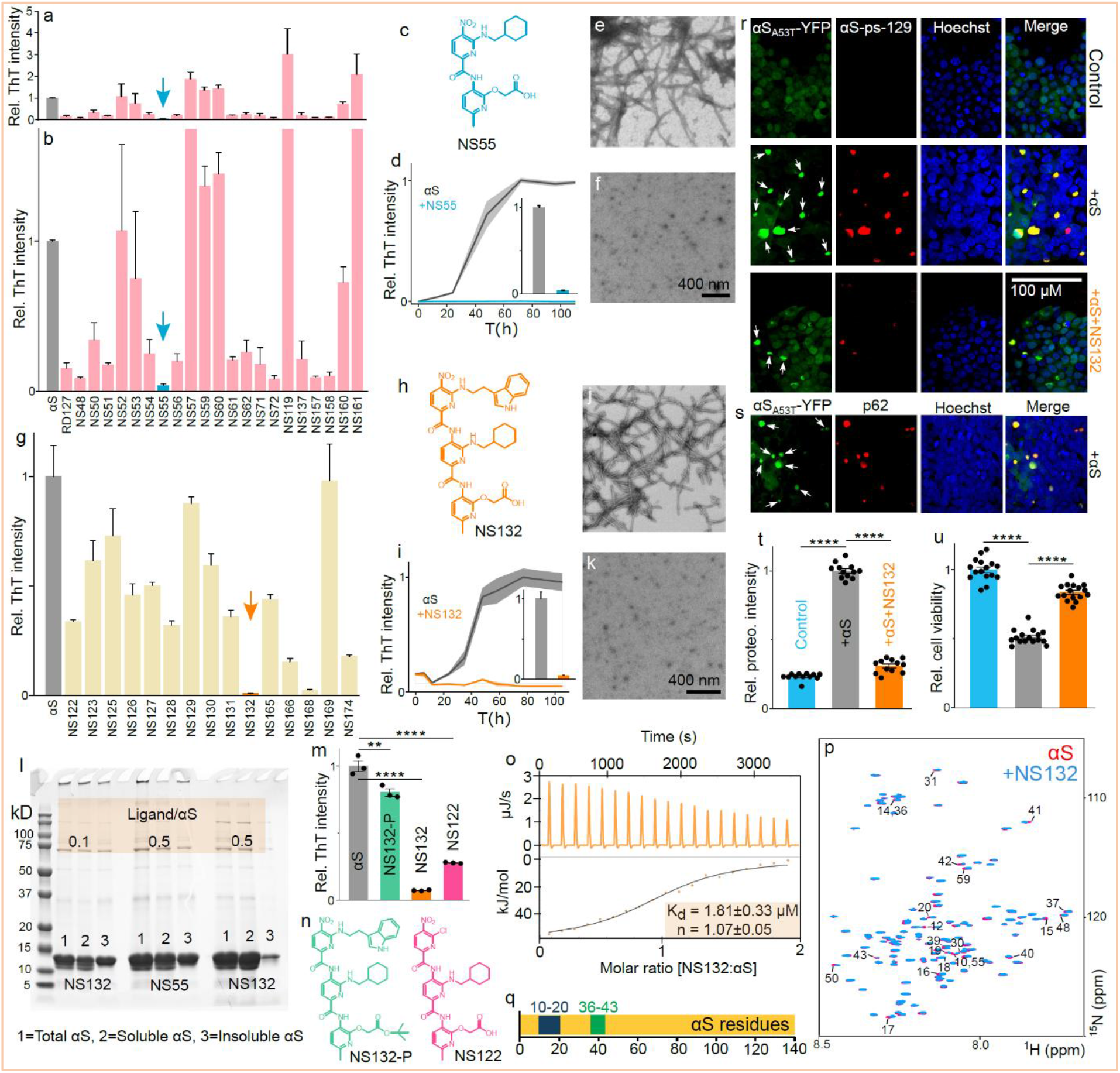
Identification and characterization of the most potent antagonist against αS aggregation using the novel 2D-FAST. **a**, The graphical representation and zoom in version (**b**) of the ThT intensity of 100 μM αS aggregation for four days in the absence and presence of dipyridyls and tripyridyls (**g**) at an equimolar ratio. The arrow indicates the most potent antagonist of αS aggregation. **c**, The chemical structure of NS55 and NS163 (**h**). The aggregation profile (**d**) and TEM image of 100 μM αS in the absence (**e**) and presence (**f**) of NS55 at an equimolar ratio. (Inset) The ThT intensity of 100 μM αS aggregation after four days in the absence and presence of NS55 at an equimolar ratio. **i**, The aggregation profile and TEM image of 100 μM αS in the absence (**j**) and presence (**k**) of NS132 at an equimolar ratio. (Inset) The ThT intensity of 100 μM αS aggregation after four days in the absence and presence of NS132 at an equimolar ratio. **l**, SDS-PAGE gel analysis of 100 μM αS aggregation after four days in the absence and presence of NS55 and NS132 at the indicated molar ratios. **m**, The ThT intensity of 100 μM αS aggregation after four days in the absence and presence of the indicated ligands (chemical structures, **n**) at an equimolar ratio.**o**, The ITC thermogram for the titration of a solution of NS132 into αS where heat burst curves (upper panel) and the corrected injection heats (lower panel). **p**, Overlay of 2D HSQC (^1^H, ^15^N) NMR spectra of 70 μM uniformly ^15^N-labelled αS in the absence (red) and presence (blue) of NS132 at an equimolar ratio. **q**, The potential binding site of NS132 on αS, represented by blue and green boxes. **r,s**, Confocal images of HEK cells treated with the aggregated solution of 5 μM αS in the absence and presence of NS132 at an equimolar ratio. Inclusions of αS_A53T_-YFP = white arrows, Hoechst = blue, αS-ps-129 = red, p62 = red (**s**), merge = Hoechst, αS-ps-129, and αS_A53T_-YFP. **j**, Confocal images of HEK cells treated with the aggregated solution of 5 μM αS. Inclusions of αS_A53T_-YFP = white arrows, Hoechst = blue, p62 = red, merge = Hoechst, p62, and αS_A53T_-YFP. The relative intensity of Proteostat dye-stained aggregates of αS_A53T_-YFP inclusions (**k**) and relative viability (**l**) of HEK cells treated with the aggregated solution of 5 μM αS in the absence and presence of NS132 at an equimolar ratio. The ThT experiments were conducted three times and the reported change in the ThT intensity was an average of three separate experiments. The cell viability or Proteostat assays were conducted with at least four biological replicates and four technical replicates for each biological replicate. The data were expressed as mean and the error bars report the s.e.m. (n = 3 to 4 independent experiments and each n consisted of three technical replicates). The statistical analysis was performed using ANOVA with Tukey’s multiple comparison test. *p < 0.05, **p < 0.01, ***p < 0.001.

The screening of tripyridyls against αS aggregation at an equimolar ratio using ThT assay led to the identification of NS132 as the most potent antagonist (Fig. 1p, 2g-i). The absence of αS fibers in the presence of NS132 was also confirmed by TEM images (Figure 2j,k). Both dipyridyl (NS55) and tripyridyl (NS132) were very effective inhibitors of αS aggregation at an equimolar ratio; however, NS132 was a far more effective antagonist than NS55 at a substoichiometric ratio of 1:0.5 (αS:ligand) and NS132 was almost equally effective at 1/5 of the conc. of NS55 in inhibiting the aggregation of αS (αS:ligand, Figure 2l), reflected by SDS-PAGE (Sodium dodecyl sulphate–polyacrylamide gel electrophoresis) and ThT assay (Supplementary Fig. 3a,b). For SDS-PAGE analysis, a solution of 100 μM αS was aggregated for four days in the absence and presence of NS55 and NS132 at various substoichiometric ratios (αS:ligand, 1:0.5, 1:0.1). Subsequently, the αS solutions were centrifuged to separate soluble and insoluble fractions, which were analyzed by SDS-PAGE (Fig. 2l). In the case of NS132, αS was predominantly detected in the soluble fraction at a substoichiometric ratio (αS:ligand, 1:0.5). In marked contrast, in the case of NS55, a significant amount of αS protein was found in the insoluble form (αS:ligand, 1:0.5). The soluble and insoluble amounts of αS were comparable when the concentration of NS132 was 5-fold less than NS55 (Fig. 2l). Collectively, both the ThT assay and SDS-PAGE analysis demonstrate that NS132 is a far better antagonist than NS55 of αS aggregation. These results highlight the validity of our 2D-FAST, where we were able to identify NS132 (tripyridyl) as a better antagonist than NS55 (dipyridyl) of αS aggregation.

To confirm that the side chains of NS132 are essential for its antagonist activity, we used various analogs of NS132 and compared their antagonist activity for αS aggregation. The ThT signal of αS aggregation was decreased from 1.0 to 0.80, 0.30, and 0.07 in the presence of NS132-P (Protected COOH group), NS122 (Chloro side chain), and NS132 at an equimolar ratio, respectively (Fig. 2m-n). The SDS-PAGE analysis also validated the ThT results, suggesting that NS132-P was a poor antagonist of αS aggregation (Supplementary Fig. 3c-f). Collectively, both ThT assay and SDS-PAGE analysis demonstrate that NS132 is a far better antagonist than NS132-P and NS122 and the side chains are important for the antagonist activity of NS132 against αS aggregation. Under matching conditions of the ThT aggregation assay, we did not observe any significant quenching of the ThT fluorescence signal by NS132 (Supplementary Fig. 4). We also characterized the binding interaction between αS and NS132 using the isothermal calorimetry titration (ITC) (Fig. 2o). The ITC titration yielded the dissociation constant (Kd) of 1.81±0.33 μM with a binding stoichiometry of 1:1 (αS:NS132) (Fig. 2o). We utilized two-dimensional heteronuclear single quantum coherence NMR spectroscopy (2D NMR HSQC) to gain insights into the binding site of NS132 on αS. We collected the HSQC NMR of 70 μM ^15^N-^1^H-uniformly labeled αS in the absence (Fig. 2p, red) and presence of NS132 (Fig. 2p, blue) and compared the signal intensity of the amide peaks. In the presence of NS132, we observed noticeable changes in the amide peaks for specific residues toward the N-terminus, indicative of the interaction and binding site of NS132 on αS, especially residues 10-20, 36-43, and 50,55,59 (Fig. 2q). The binding sites of NS132 have been suggested to be the essential sequences for αS aggregation and these sequences have been considered to be the potential therapeutic targets for the effective inhibition of αS aggregation and associated PD phenotypes^24,47^. Our study supports the hypothesis that the targeting of these sequences will effectively inhibit αS aggregation.

To confirm that NS132 did not generate fiber-competent cytotoxic structures during αS aggregation inhibition, we utilized a well-established model of HEK293 cells, which stably express YFP-labeled αS-A53T mutant (αS-A53T-YFP)^23,24,50^. The endogenous monomeric αS-A53T-YFP have been shown to template into fibers when transfected with exogenously added αS fibers in the presence of Lipofectamine 3000 (Fig. 2r,s)^23,24,50^. The aggregation of endogenous monomeric αS-_A53T_-YFP into fibers can be detected by the intracellular fluorescent puncta (Fig. 2r,s). A solution of 100 μM αS was aggregated in the absence and presence of NS132 at an equimolar ratio for four days. The resulting solutions of αS fibers (5 μM in monomeric αS, ±NS132) were introduced to HEK cells in the presence of Lipofectamine 3000 for 24 h. In contrast to the control (no fibers), we observed a significant number of fluorescent inclusions in the presence of αS fibers (Fig. 2r,s, white arrows, αS-_A53T_-YFP), which was a consequence of the templating of endogenous monomeric αS-_A53T_-YFP by the exogenously added αS fibers. The αS inclusions were colocalized in the cytoplasm of HEK cells as suggested by others as well^24,51–54^. In addition, we also observed that the αS-pS-129 protein (Phosphorylated αS at residue 129) colocalized in the aggresome of αS inclusions in HEK cells (Fig. 2r, red). Moreover, we observed the colocalization of an adaptor protein, p62, in the aggresome of αS inclusions in HEK cells (Fig. 2s, red) ^24,51–54^. It has been suggested that during αS aggregation, p62 recruits the autophagy machinery to the αS inclusions^51–54^. The autophagy machinery regulates many vital cellular processes and its impairment due to αS aggregation can lead to PD and other neurodegenerative disorders^51–54^. In marked contrast, in HEK293 cells transfected with αS aggregated in the presence of NS132, there was a significant decrease in the intracellular αS-_A53T_-YFP inclusions (Fig. 2r, + αS+NS132). We also quantified the inclusions (αS-_A53T_-YFP) using a novel ProteoStat dye-based high throughput 96-well plate reader-based assay recently developed in our lab^24^. The ProteoStat dye-based intensity of HEK cells treated with αS fibers was ~4-5 fold higher than the control (no fibers) (Fig. 2t). In marked contrast, we did not observe a significant difference in the ProteoStat intensity of HEK cells treated with αS fibers+NS132 and the control conditions (Fig. 2t). Both proteins, including αS-pS-129 and p62 have been shown to be key pathological biomarkers for the formation of Lewy body like aggregates^24,51–54^. The presence of both proteins in the inclusions suggests that these inclusions mimic some features of Lewy-body like structures that are important events in inducing cytotoxicity in HEK cells^24,51–54^. Therefore, we used HEK cells to test the cytotoxicity of the aggregated solution of αS in the absence and presence of NS132 (Fig. 2u). The viability of HEK cells was measured using the (3-(4,5-dimethylthiazol-2-yl)-2,5-diphenyltetrazolium bromide) (MTT) reduction based assay. The viability of HEK cells decreased to 52% in the presence of the aggregated solution of αS; however, the viability of HEK cells was improved to 85% in the presence of the αS aggregated solution with an equimolar ratio of NS132 (Fig. 2u). This data suggest that NS132 doesn’t promote the formation of seed competent αS assemblies and the higer order aggresome.

### Antagonist effect of OPs against fibers catalyzed aggregation of αS

In addition to the spontaneous accumulation of αS via *de novo* aggregation, another crucial mechanism for inducing pathology in PD is αS fibers catalyzed aggregation of αS^20,24,55–61^. Therefore, we monitored the effect of NS132 on the αS fibers catalyzed aggregation of αS. The aggregation of 100 μM αS monomer in the presence of preformed αS fibers (20%, monomer eq.) resulted in the acceleration of monomeric αS aggregation, reflected by a significant increase in the ThT signal after 20 h (Fig. 3a). The αS fibers catalyzed aggregation of αS was wholly suppressed by NS132 at an equimolar ratio as evidenced by significantly lower ThT signal (Fig. 3a). The antagonist activity of NS132 was also assessed on more robust αS fibers generated using the protein misfolding cyclic amplification (PMCA) technique^20,24,57–60^. The PMCA technique is used to amplify the aggregation of proteins from a small number of fibers from the previous cycle, which generates robust fibers via a nucleation-dependent polymerization model^20,24,57–60^. In the PMCA assay, the fibers of αS are amplified for five cycles using αS monomer and αS fibers from the previous cycle. The ThT intensity for αS aggregation via PMCA assay increases significantly after cycle two and the intensity stays consistent up to cycle five (Fig. 3b, grey bar). Additionally, PMCA samples (from αS aggregation) from cycle three to cycle five were PK (proteinase K) resistant (Fig. 3c, white arrow), which indicates these αS fibers are very robust and non-degradable after cycle three (Fig. 3c). In marked contrast, in the presence of NS132 at an equimolar ratio, we did not observe any significant change in the ThT intensity up to cycle five (Fig. 3b, orange bar). More importantly, the PMCA assay samples of αS aggregation in the presence of NS132 from cycle three to cycle five were completely degraded by PK treatment (Fig. 3d, orange arrow). The data suggest that NS132 interacts with αS and generates off-pathway fiber-incompetent structures, which are easily degradable with PK treatment. To further validate that NS132 generates fiber incompetent structures from the fiber catalyzed aggregation, we utilized HEK cells that stably express αS-_A53T_-YFP^24,51–54^. The solutions of αS aggregation in the absence and presence of NS132 from the PMCA cycle five (2 μM in monomeric αS, ±NS132) were introduced to the HEK cells in the presence of Lipofectamine 3000 for 24 h. In contrast to the control (no fibers) (Fig. 3e, control), we observed a significant number of inclusions in the presence of αS fibers (Fig. 3e, white arrows, αS-_A53T_-YFP). In addition, we also observed the colocalization of αS-pS-129 in the aggresome of αS inclusions in HEK cells (Fig. 3e, αS-pS-129, red). In contrast, in the presence of the PMCA assay sample from cycle five (+NS132), there was a significant reduction in the number of αS inclusions (Fig. 3e, +NS132). The ProteoStat intensity of inclusions in HEK cells treated with the PMCA sample from cycle five was ~2-3 fold higher than the control condition (no fibers) (Fig. 3f). In marked contrast, we observed a significantly lower ProteoStat intensity in the presence of the PMCA sample from cycle five in the presence of NS132 (Fig. 3f). Clearly, NS132 was a potent antagonist of the fibers catalyzed aggregation of αS and it generates fiber incompetent off-pathway structures.

**Fig.3.**
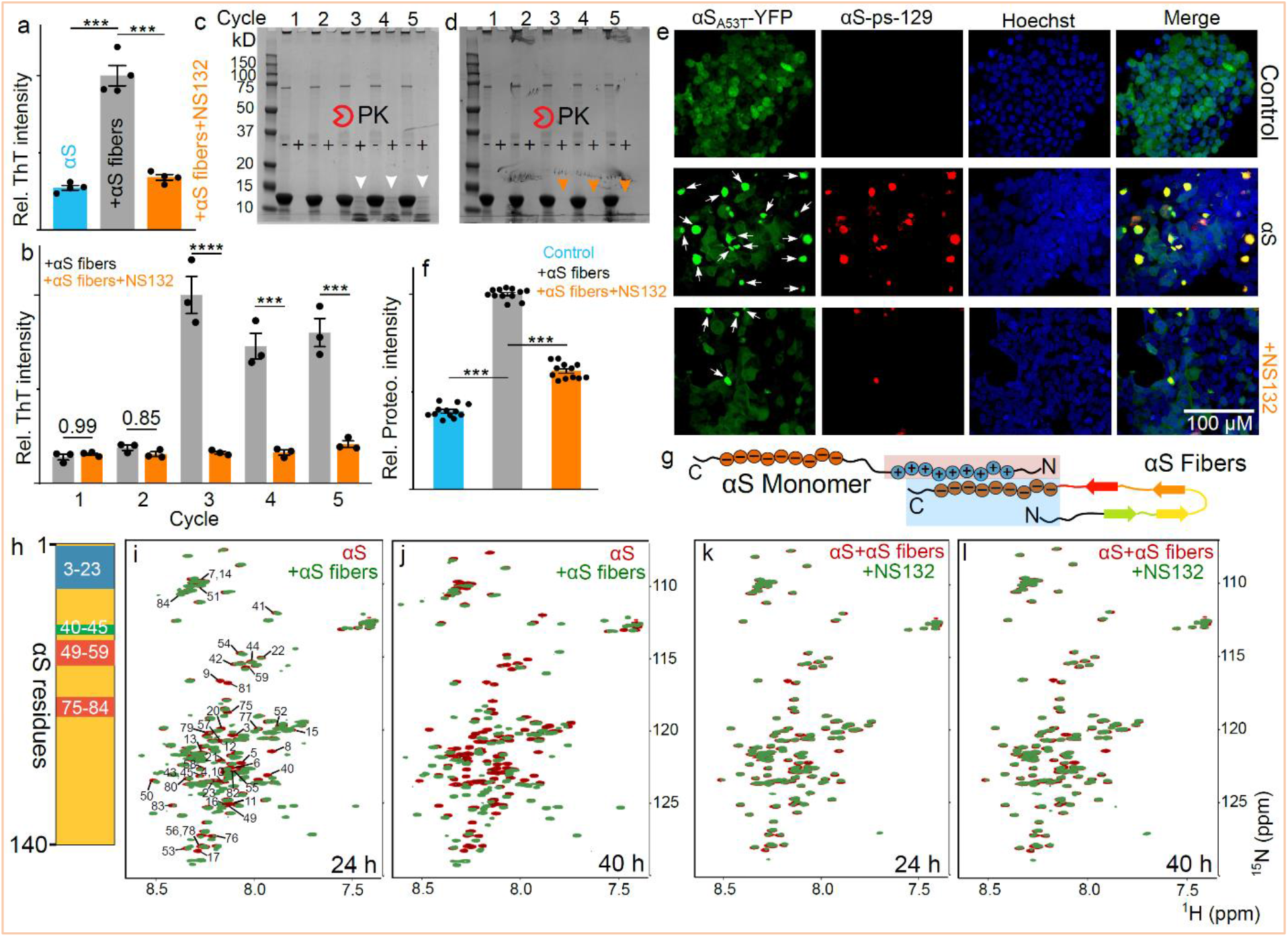
The effect of NS132 on the fibers catalyzed aggregation of αS. **a**, The relative ThT intensity of αS fibers (20% monomer) catalyzed aggregation of 100 μM αS in the absence and presence of NS132 at an equimolar ratio after 24 h in the aggregation conditions. The Bis-tris gels of PMCA samples from the firstto the fifth cycle in the absence (**b**) and presence (**c**) of NS132. The (-) and (+) signs indicate the amplified samples without and with the treatment of PK, respectively. The arrows indicate the effect of PK on PMCA samples from the indicated cycles. **d**, The statistical analysis of the relative ThT intensity of various PMCA samples in the absence (grey bar) and presence (orange bar) of NS132, before treating these samples with PK. **e**, The representative confocal images of HEK cells after treatment with PMCA samples from the fifth cycle under the indicated conditions. The αS_A53T_-YFP (green) inclusions are indicated by white arrows. Hoechst = blue, αS-ps-129 = red, merge = Hoechst, αS-ps-129, and αS_A53T_-YFP. **f**, The relative intensity of Proteostat dye-stained aggregates of αS_A53T_-YFP inclusions in HEK cells treated with PMCA samples from the fifth cycle (2 μM αS in monomer) in the absence and presence of NS132. **g**, A model for the proposed interaction of αS monomer with αS fibers. **h**, The proposed binding interaction sites of αS fibers on the αS monomer residues. The comparison of the HSQC NMR of the 70 μM ^15^N-labeled αS in the absence (red) and presence (green) of αS fibers after 24 h (**i**) and 40 h (**j**). The comparison of the HSQC NMR of the premixed solution of ^15^N-labeled αS+αS fibers in the absence (red) and presence (green) of 70 μM NS132 after 24 h (**k**) and 40 h (**l**). The ThT experiments were conducted three times and the reported change in the ThT intensity was an average of three independent experiments. The Proteostat assay was conducted with at least four biological replicates and three technical replicates for each biological replicate. The data were expressed as mean and the error bars report the s.e.m. (n = 3 independent experiments and each n consisted of three technical replicates). The statistical analysis was performed using ANOVA with Tukey’s multiple comparison test. *p < 0.05, **p < 0.01, ***p < 0.001.

We also employed 2D NMR HSQC for atomic-level insights into fibers catalyzed aggregation of αS and its inhibition by NS132. It has been suggested that the negatively charged flexible C-terminal tail of αS (in fibers) interacts and recruits the positively charged N-terminal segment of αS (in monomer) during the fibers catalyzed aggregation of αS (Fig. 3g)^62^. We have also shown that NS132 specifically interacts with the N-terminal residues of αS. Therefore, we hypothesize that NS132 will be able to inhibit the interaction of αS (monomer) with αS fibers, which is suggested to be the prerequisite interaction to initiate the seed-catalyzed aggregation of αS. We incubated preformed fibers of αS with 70 μM ^15^N-^1^H-uniformly labeled αS monomer and used HSQC NMR to characterize the kinetics of fibers catalyzed aggregation of αS on a molecular level. The total changes in the intensity of the amide peaks of ^15^N αS monomer in the presence of αS fibers suggest that the binding interaction was predominantly toward the N-terminus of αS (Fig. 3h,i); more specifically, αS fibers interact specifically with four αS sequences, including 3-23, 40-45, 49-59, and 75-84 (Fig. 3h,i). Also, there was an induction of a secondary structure in the αS monomer, indicated by the spreading of the amide peaks toward the ^1^H resonances (Fig. 3i, green). At 40 h, more pronounced changes were observed in the location and intensity of the amide peaks of αS monomer, suggesting a much stronger interaction with αS fibers and further induction of a secondary structure in ^15^N-^1^H-αS monomer (Fig. 3j). In marked contrast, no significant change in the amide peaks of ^15^N-^1^H-uniformly labeled αS monomer was observed in the presence of NS132 at an equimolar ratio (Fig. 3k). Even after 40 h, there were fewer and smaller changes in the amide peaks of ^15^N-^1^H-uniformly labeled αS monomer in the presence of NS132 (Fig. 3l). The NMR study suggests that NS132 inhibits the αS monomer-αS fibers interaction by potentially interacting at the N-terminal of αS. Clearly, NS132 is a potent inhibitor of fibers catalyzed aggregation of αS.

### Effect of OPs on intracellular αS aggregation in a C. *elegans* PD model

Next, we investigated the antagonist activity of NS132 against αS aggregation in a well-established *C. elegans* PD model. The *C. elegans* models have been extensively used to study the underlying mechanisms and therapeutic interventions for neurodegenerative diseases associated with protein aggregation because of the short lifespan (2-3 weeks), tractability to genetic manipulation, distinctive behavioral and neuropathological defects, and high degree of genetic relevance compared to humans.^20,24,63^ We utilized NL5901 worms, which express αS-fused yellow fluorescent protein (αS-YFP) in the body wall muscle cells^20,24,63^. The PD phenotypic readouts in the NL5901 worms include a gradual increase in inclusions (αS-YFP) in body wall muscle cells and a decline in motility during the aging of the worms^20,24,63^. To act as a potent antagonist of αS aggregation in *C. elegans* model, NS132 should permeate the cell membrane of the body wall muscle cells of *C. elegans*. We used the parallel artificial membrane permeation assay (PAMPA) to test the cell permeability of NS132 and compare it with various PAMPA standards of cell permeabilities (Fig. 4a). The cell permeability of NS132 was lower in comparison to the PAMPA medium standard (Fig. 4a), which was most likely a consequence of the COOH functional group. We have also shown that COOH is a very important side chain for the antagonist activity of NS132 against αS aggregation (NS132 vs NS132-P). We surmise that we may underachieve the overall antagonist effect of NS132 against αS aggregation in *C. elegans* due to its less than moderate cell permeability. Therefore, to enhance the cell permeability without sacrificing the antagonist activity of NS132, we synthetically replaced the carboxylic acid with hydroxamic acid (NS163, Fig. 4a,b), which is considered to be one of the most common and successful carboxylic acid isosteres in the pharmaceutical industry and has shown higher cell permeability than the former^64^. The cell permeability of the hydroxamic acid analog (NS163) was higher than NS132 (Fig. 4a,b). NS163, similar to NS132, was a potent antagonist of αS aggregation, confirmed by ThT assay (Fig. 4c) and TEM images (Supplementary Fig. 5). Clearly, the PAMPA assay, ThT assay, and TEM demonstrate that we have improved the cell permeability of NS132 without sacrificing the antagonist activity against αS aggregation. Subsequently, we tested the antagonist activity of NS163 against αS aggregation-mediated PD phenotypes in NL5901 worms. The NL5901 worms were treated with 50 μM NS163 on day two, followed by incubation at r.t. for one day before assessing the effect of NS163 on PD phenotypes. We observed a gradual increase in the number of inclusions from day five (~15 inclusions/worms) to day eight (~34 inclusions/worms) and a slight decrease on day nine (~32 inclusions/worms), suggesting a saturation in the number of inclusions after day eight in the animals (Fig. 4d,e, white arrow). In marked contrast, there was a substantial decline in the number of inclusions in the presence of NS163 from day five (~4 inclusions/worms) to day nine (~2 inclusions/worms) (Fig. 4d,g). We also observed a decrease in the number of inclusions in the presence of NS132 (lower effect than NS163), despite its lower cell permeability (Fig. 4d,f). As we predicted earlier, the cell permeability of NS132 was lower than NS163, which is the likely reason for the lower effect of NS132 (Fig. 4d,f). The motility rate of the NL5901 worms decreases during the aging process due to an increase in αS aggregation, which impairs the muscle cells. We utilized the WMicroTracker ARENA plate reader^24,65^ to measure the motility rate of NL5901 worms in the absence and presence of NS163. We observed a significant decline in the activity of NL5901 worms in comparison to the control worms (N2, healthy *C. elegans* strain) (Fig. 4h,i). In marked contrast, the NL5901 worms treated with NS163 on day two resulted in a significant improvement in the motility rate (Fig. 4h,i). The *C. elegans-* based study suggests that NS163 permeates the body wall muscle cell membrane of the worms and inhibits αS aggregation. NS132 also demonstrated an increase in the motility rate of NL5901 worms, albeit with a less pronounced effect than NS163 (Supplementary Fig. 6).

**Fig. 4.**
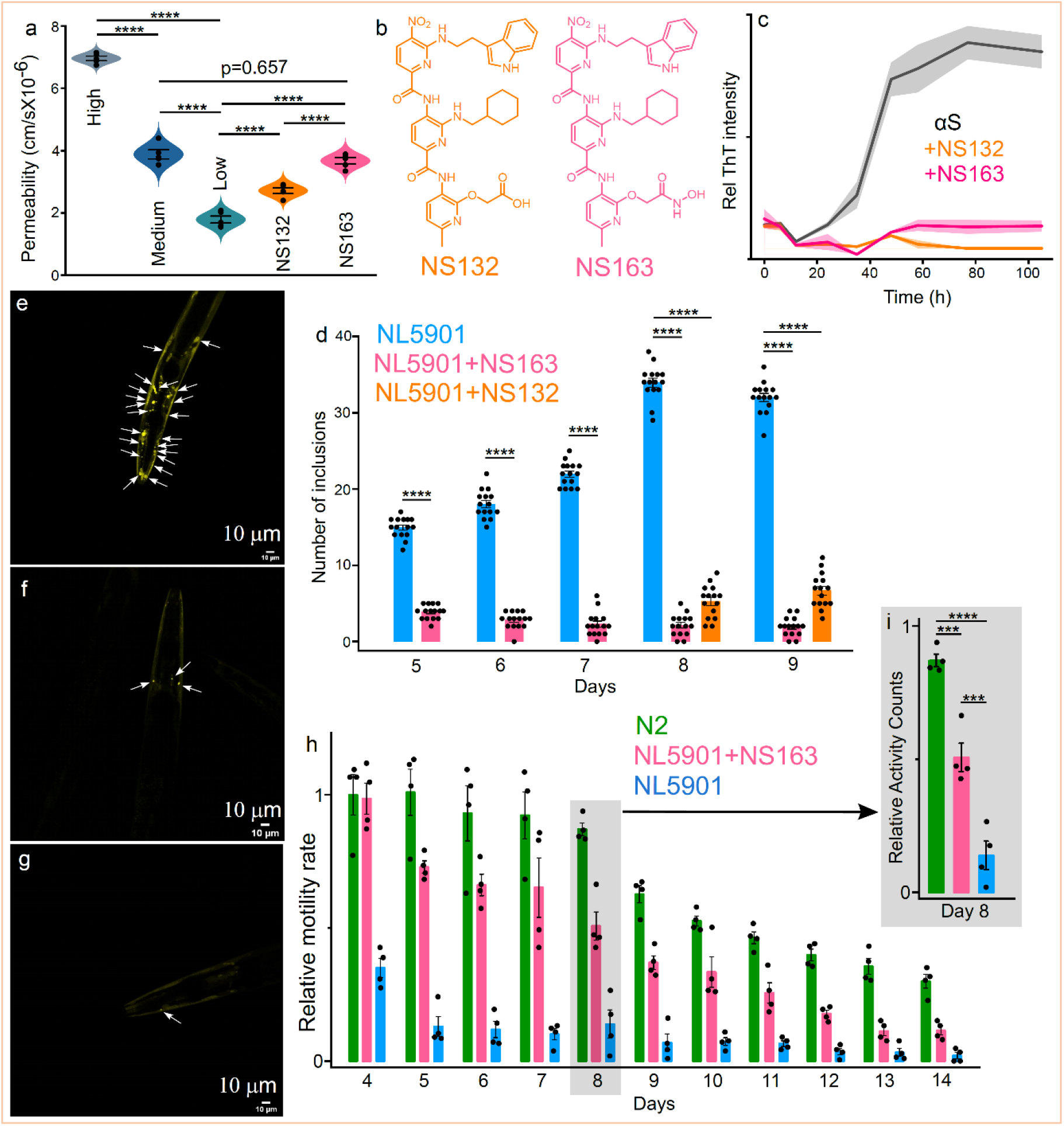
The intracellular inhibition of αS aggregation by OPs in an *in vivo* PD model. **a**, Assessment of cell permeability of the indicated ligands using the PAMPA. **b**, The chemical structures of NS132 and NS163. **c**, The ThT-based aggregation profile of 100 μM αS in the absence and presence of the indicated ligands at an equimolar ratio. **d**, The number of inclusions for experiment ‘**e-g**’ in NL5901 in the absence and presence of NS132 and NS163 from day four to day nine of the adulthood. The representative confocal images of αS-YFP inclusions (white arrows) in the body wall muscle cells of NL5901 (Days = 8) in the absence (**e**) and presence of 50 μM NS163 (**f**) and 50 μM NS132 (**g**). **h**, The motility rate of N2 and NL5901 and statistics (Day 8, **i**) in the absence and presence of 50 μM NS163 during the aging process. For each confocal imaging experiment (e-g), at least 10 worms were used, and the inclusions were counted manually, and each condition (Each day) consisted of at least four independent experiments. For motility experiments, a total of 50 worms were used in duplicate for each experiment and each condition consisted of at least four independent experiments. The data were expressed as mean and the error bars report the s.e.m. (n = 3 or 4 independent experiments and each n consisted of a minimum of three technical replicates). The statistical analysis was performed using ANOVA with Tukey’s multiple comparison test. *p < 0.05, **p < 0.01, ***p < 0.001.

### Effect of OPs on the degeneration of DA neurons in a *C. elegans* PD model

The aggregation of αS is associated with the neurodegeneration of DA neurons, which is a pathological hallmark of PD. We have also shown that NS163 rescues αS aggregation-mediated PD phenotypes in the body wall muscle cells of *C. elegans* worms. Next, we investigated the neuroprotective effect of NS163 on αS aggregation-mediated degeneration of DA neurons in a well-established *C. elegans* PD model (UA196)^20,66–69^. The UA196 worms express both human αS and GFP in DA neurons under the control of the dopamine promoter genotype (Pdat-1::GFP; Pdat-1::α-SYN). The expression and aggregation of αS in six DA neurons that are located within the anterior region of worms lead to progressive neurodegeneration characteristics during the aging of UA196 worms. This strain has been used to gain insights into the PD-associated mechanisms and to assess the neuroprotective effect of ligands against αS aggregation^20,66–69^. The DA neurons in UA196 worms degenerate from day three to day 15, represented by a gradual decline in the GFP fluorescence in DA neurons shown by others as well (Fig. 5a-g)^66–69^. The number of DA neurons decreased from 6 (day three) to 1 (day 15) in UA196 worms (Fig. 5b-g). In marked contrast, the % loss of six intact DA neurons in UA196 worms was 95%, 87%, and 78% on day five, 10, and 15, respectively, after treatment with 50 μM NS163 on day two (Fig. 5h-k). The degeneration and subsequent loss of DA neurons is a consequence of αS aggregation during the aging of worms (Fig. 5d-f). The % decline in the total DA neurons in UA196 worms was 72%, 36%, and 23% after five, 10, and 15 days, respectively (Fig. 5k, blue). In contrast, in NS163 treated UA196 worms, the % loss of the total number of DA neurons was <5% even up to 15 days (Fig. 5k, red). The data suggest a remarkable neuroprotective effect of NS163 as a significant number of DA neurons were intact up to day 15 (Fig. 5h-k). Other reported ligands were not able to achieve such a remarkable neuroprotective effect against αS aggregation mediated degeneration of DA neurons in *C. elegans* PD model even at 1 mM concentration^70^. Under matched conditions, NS132 also displayed a very good neuroprotective effect on the degeneration of DA neurons in UA196 worms, confirmed by confocal imaging (Supplementary Fig. 7a-f). The neuroprotective effect of NS163 was better than NS132, indicated by a higher number of healthy neurons (Fig. 5k, Supplementary Fig. 7f), most likely due to the former’s better ability to permeate the cell membranes.

**Fig.5.**
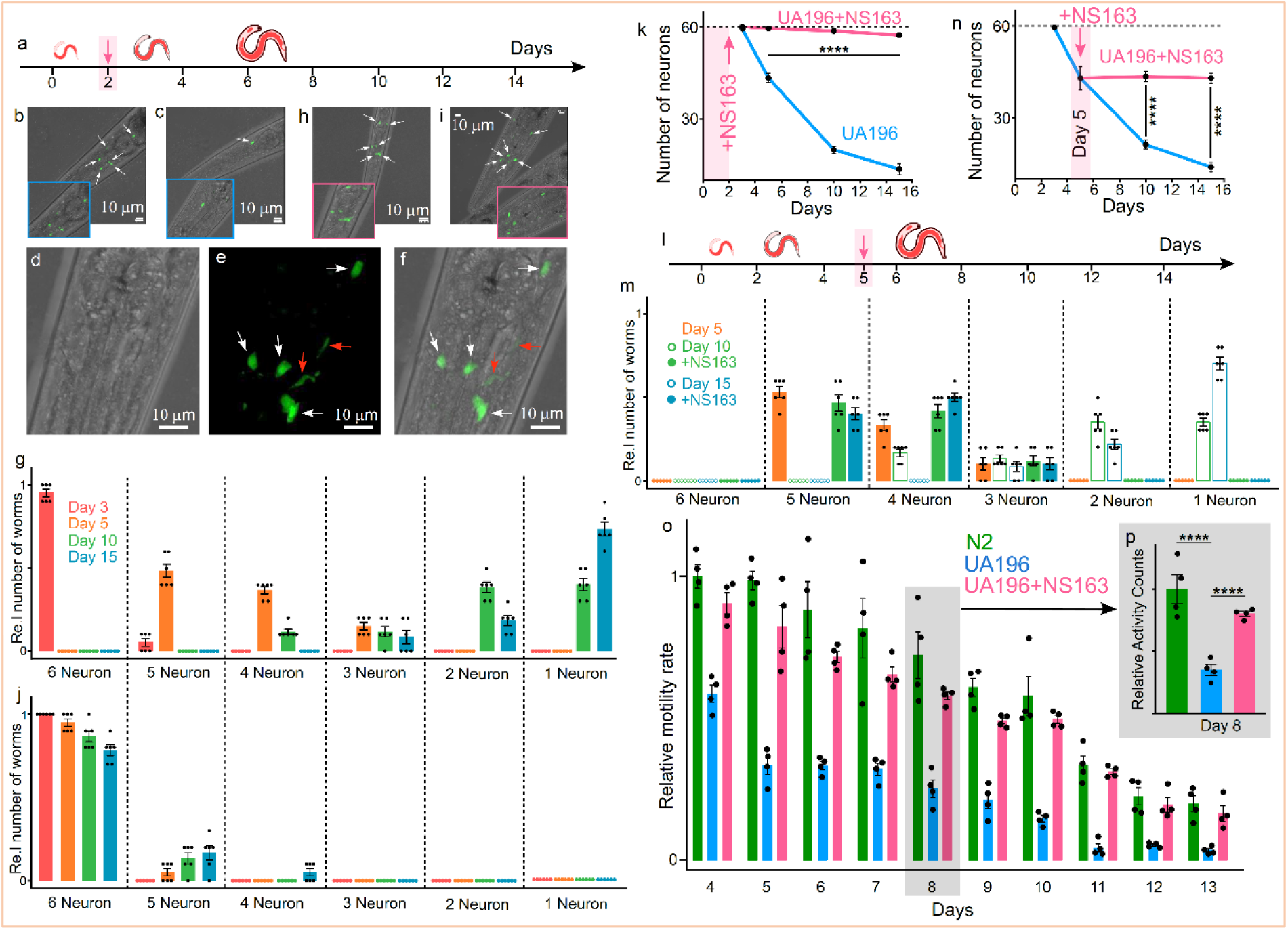
Neuroprotective effect of OPs on the degeneration of DA neurons. **a,** Schematic of the aging process of UA196 worms and their treatment with the ligands. Representative confocal images of UA196 worms in the absence (**b,c**) and presence (**h,i**) of 50 μM NS163 on day three and day 15. **d,e**,**f**, The healthy (white arrow) and degenerated (red arrow) DA neurons in UA196 worms on day five. **g,** The relative number of neurons in UA196 worms during the aging process in the absence (**g**) and presence of 50 μM NS163 (**j**). **k**, Statistics for the total number of neurons in UA196 worms during the aging process in the absence and presence of 50 μM NS163. **l**, Schematic of the aging process of UA196 worms and their treatment with the ligands in a late-onset disease model (day five). **m**, The relative number of neurons in UA196 worms during the aging process when treated on day five with 50 μM NS163. **n**, Statistics for the total number of neurons in UA196 worms when treated on day five with 50 μM NS163. **o**, The comparison of the motility rate for 13 days of N2 and UA196 and statistics (for day eight, **p**) in the absence and presence of 50 μM NS163. For each confocal imaging experiment (**b-f, h-i**), at least 10 worms were used, and the healthy neurons (GFP signal) were counted manually, and each condition (day) consisted of six independent experiments. For motility experiments, a total of 50 worms were used in duplicate for each experiment and each condition consisted of at least four independent experiments. The data were expressed as mean and the error bars report the s.e.m. (n = 4 independent experiments and each n consisted of at least four technical replicates). The statistical analysis was performed using ANOVA with Tukey’s multiple comparison test. *p < 0.05, **p < 0.01, ***p < 0.001, ****p < 0.0001.

### Effect of OPs on the motility rate of UA196 worms

The degeneration of DA neurons has been directly linked with the loss of motor functions resulting in slow motility rate^13–18,67^. Therefore, we assessed the motility rate of UA196 in the absence and presence of NS163 using WMicroTracker ARENA plate reader (Fig. 5o,p). There was a significant decline in the motility rate of the UA196 worms (Fig. 5o,p, blue) during the aging process in comparison to the control worms (Fig. 5o,p, N2, green). In marked contrast, the motility rate of UA196 worms treated with 50 μM NS163 (day two) was significantly improved during the aging process. The improvement in the motility rate is likely due to the rescue of the degeneration of DA neurons by NS163. NS132 also displayed a neuroprotective effect; therefore, we also assessed its effect on the motility rate of UA196 worms. Under matched conditions to NS163, we observed a noticeable rescue of the motility rate of UA196 worms in the presence of NS132 during the aging process (Supplementary Fig. 8). The effect of NS163 was better than NS132 in rescuing the motility rate of UA196 worms.

### Effect of OPs on the ROS level in UA196 worms

One of the causal agents associated with the etiology of PD is the generation of ROS, which oxidizes lipids, proteins, and DNA^66,68,70^. The neurodegeneration in UA196 worms due to αS aggregation leads to the production of intraworm ROS. The ROS level was determined using a fluorescent probe (CM-H2DCFDA), which reacts with intraworm ROS level in UA196 worms (day eight) and produces a green fluorescent signal, whose intensity increases up to 2 h (Fig. 6a,c)^70^. In marked contrast, UA196 worms treated with 50 μM NS163 (on day two) displayed a significant decrease in the intracellular ROS level on day eight (Fig. 6b,c). The signal intensity was very similar for the dye sample and UA196 worms treated with NS163 (Fig. 6c). The data suggest that the decrease in the ROS level in UA196 worms in the presence of NS163 is a consequence of the rescue of degeneration of DA neurons. Similarly, in the presence of NS132, the ROS level was low in UA196 worms (Supplementary Fig. 9a). It has been shown earlier that the GFP signal in DA neurons does not interfere significantly with the detection of the ROS level by the green fluorescent dye because of the weak signal of the former in comparison to the later.^70^

**Fig.6.**
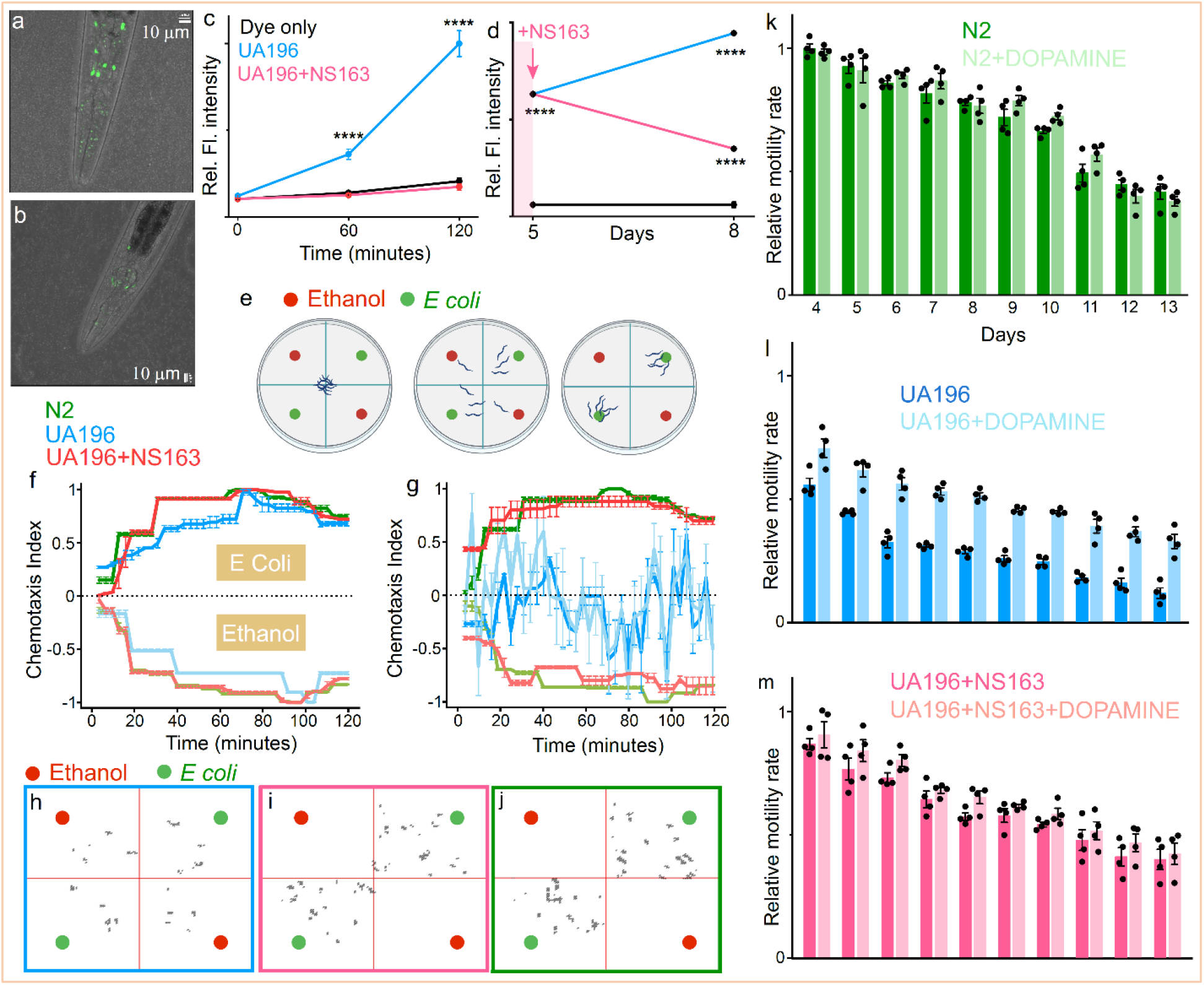
Effect of OPs on the PD phenotypes in UA196 worms. Representative confocal images of UA196 worms (day eight) treated with CM-H2DCFDA dye to quantify the ROS level (green) in the abence (**a**) and presence of 50 μM NS163 (**b**, day two treatment with NS163). **c**, Statistical analysis of the quantification of the ROS level for experiment **a-b**. **d**, Statistical analysis of the ROS level in UA196 worms when 50 μM NS163 was added on day five (arrow). **e**, Schematic to assess the behavioral deficits in UA196 worms in a petri dish in the presence of ethanol and *E. coli* as a function of time. The CI graph for N2, UA196 worms, and UA196 worms treated with 50 μM NS163 under the indicated conditions on day three (**f**) and day 10 (**g**). Snapshots at 60 min. of the animated videos collected for the CI for UA196 (**h**), UA196+50 μM NS163 (**i**), and N2 (**j**) under the indicated conditions on day 10. The motility rate for N2 (**k**), UA196 worms (**l**) and UA196 worms treated with 50 μM NS163 (**m**) in the absence and presence of 2 mM dopamine. For the ROS level quantification, at least 50 worms were used and each condition consisted of three independent experiments. For motility assay experiment, a total of 50 worms were used in duplicate for each experiment and each condition consisted of four independent experiments. For chemotaxis assays, a total of 50 worms were used for each experiment and each condition consisted of three independent experiments. The data were expressed as mean and the error bars report the s.e.m. (n = 3 or 4 independent experiments and each n consisted of a minimum of two technical replicates). For ROS assay, the data were expressed as mean and the error bars report the s.d. (n = 3 independent experiments and each n consisted of five technical replicates). The statistical analysis was performed using ANOVA with Tukey’s multiple comparison test. *p < 0.05, **p < 0.01, ***p < 0.001, ****p < 0.0001.

### Effect of OPs on Behavioral deficits in UA196 worms

It has been shown that the lack of dopamine synthesis in DA neurons in *C. elegans* leads to behavioral deficits, including food sensing behavior^66,67,72^. The DA neurons in UA196 worms degenerate over time, which leads to a decrease in the amount of dopamine and NS163 rescues the degeneration of DA neurons. Therefore, we hypothesized that NS163 would be able to rescue behavioral deficits of UA196 worms. We used a chemotaxis assay to assess the effect of NS163 on the behavioral deficits of UA196 worms. In this assay, a petri dish is divided into four quadrants where two opposite quadrants were treated with a toxic chemical (ethanol, repellent) or food *(E. coli,* attractant) for worms ^71–74^. For each experiment, 50 worms were placed at the center of the dish and ethanol (red dots) and *E. coli* (green dots) were placed at the polar ends of the petri dish (Fig. 6e). We used the ARENA plate reader to measure the chemotaxis index (CI) over time with values from −1.0 to +1.0. The kinetics of the CI index over time was generated based on the time spent by worms in ethanol or *E. coli* quadrants. The kinetics of the CI over time was monitored on day three for 2 h for various worms, including N2 (Fig. 6f, Movie S1), UA196 (Fig. 6f, Movie S2), and UA196+50 μM NS163 (Fig. 6f, Movie S3). The kinetic data for the CI suggest that all the worms spent most of their time in the *E. coli* quadrants reflected by a value of ~1 during the course of 2h (Fig. 6f). None of the worms displayed behavioral deficit on day three (Fig. 6f, Movie S1, S2, S3). All the DA neurons in UA196 worms on day three were intact (Fig. 5b,g,k); therefore, we anticipate a similar behavioral response of UA196 (Fig. 6f, Movie S2) and N2 (Fig. 6f, Movie S1) worms. In marked contrast, on day 10, the kinetics of the CI of UA196 did not display any preference for ethanol (repellent) or *E. coli* (attractant) for the whole time course of the experiment (Fig. 6g,h, Movie S6). The lack of preference of UA196 worms is due to the behavioral deficits caused by the decrease in dopamine as a result of the degeneration of DA neurons. However, the UA196 worms treated with NS163 strongly favored *E. coli* (Fig. 6g,i, Movie S7) than ethanol, similar to the control worms (Fig. 6g,j, Movie S5), indicated by the chemotaxis indices. Clearly, NS163 was able to rescue the behavioral deficits of UA196. Similarly, NS132 was also able to rescue behavioral deficits in UA196 worms under the matched conditions to NS163 (Supplementary Fig. 10, Movie S4,S8).

### Effect of OPs on the dopamine level in UA196 worms

We have shown that a decrease in the motility rate in UA196 worms is a consequence of the loss of DA neurons, which is potentially associated with a substantial decrease in dopamine synthesis^66,67^. Therefore, we hypothesized that the motility rate could be enhanced by administrating dopamine in UA196 worms. The 2 mM dopamine treated UA196 worms displayed a much higher motility rate during the aging process (Fig. 6l). In marked contrast, we did not observe any significant difference in the motility rate for 50 μM NS163 treated UA196 worms in the absence and presence of 2 mM dopamine (Fig. 6m). We observed similar behavior for the N2 worms (to UA196+NS163) in the presence of 2 mM dopamine (Fig. 6k). The data suggest that the improvement in the motility rate of the dopamine treated UA196 worms is the compensation for the decrease in the dopamine synthesis due to the loss of DA neurons. However, we did not observe any change in the motility rate for N2 and NS163 treated UA196 worms because of the intact DA neurons and dopamine synthesis.

### Effect of OPs on the neurodegeneration in a post-disease onset PD model

NS163 had shown a remarkable neuroprotective effect against the degeneration of DA neurons in UA196 worms, when it was added on day two to UA196 worms. However, the effect of NS163 has not been tested in a post-disease onset PD model. For PD, most of the current therapeutic intervention strategies predominantly rely on the post-disease-onset model and the treatment occurs after the diagnosis of PD^69,75,76^. Also, during the post-disease onset of PD, the aggregation of αS is facilitated by multiple mechanisms, including the *de novo* αS aggregation, fibers catalyzed αS aggregation, and prion-like spread of αS fibers^15–18,24^. Moreover, we have identified that NS163 is a potent inhibitor of both the *de novo* (Fig. 2g-m) and the fibers catalyzed αS aggregation (Supplementary Fig. 11). Therefore, we envision that NS163 will be effective in rescuing the degeneration of DA neurons in the post-disease onset model of PD in UA196 worms. We have already shown that the DA neuron loss in UA196 worms was 28%, 67%, and 78% after five, 10, and 15 days, respectively (Fig. 5m,n). It has been suggested that ~30% neuronal loss or day four timepoint of adulthood (total five days) in *C. elegans* is considered to be a post-disease-onset PD model^69,75,76^. Therefore, we chose day five of the UA196 worms as the post-disease onset PD model, where 28% of the total DA neurons were degenerated (Fig. 5m,n). The UA196 worms were treated with 50 μM NS163 on day five and the number of healthy DA neurons was counted on day 10 and day 15 (Fig. 5l-n). NS163 was very effective in rescuing the degeneration of DA neurons as we did not observe any further loss of DA neurons in the presence of NS163 (Fig. 5l-n). The effect of NS163 was also tested on the intraworm ROS level in the post-disease model of *C. elegans*. The ROS level of UA196 worms was measured on day five, followed by the addition of 50 μM NS163 and the ROS level was assessed on day eight in the absence and presence of NS163 (Fig. 6d). In UA196 worms, the ROS level increased from day five to day eight due to the increase in the degeneration of DA neurons mediated by αS aggregation (Fig. 6d). However, the ROS level was significantly decreased in UA196 worms treated with NS163 (Fig. 6d). Clearly NS163 is a potent ligand in rescuing degeneration of DA neurons (and ROS level) when added to a post-disease onset PD model. Similarly, NS132 was also effective (less than NS163) in rescuing degeneration of DA neurons in a post-disease onset PD model, characterized by confocal imaging and the number of healthy neurons (Supplementary Fig. 12a-h). Additionally, NS132 was also very effective in decreasing the ROS level when added on day five to UA196 worms (Supplementary Fig. 9b). It is a remarkable finding that NS163 and NS132 are able to rescue PD phenotypes in a post-disease onset PD model. Most of the ligands reported in the literature rescue PD phenotypes when added in the early stages of PD models, which does not mimic the clinical landscape for the current therapeutic interventions that rely on the post-diagnosis of PD^69,75,76^.

Our data suggest that NS163 and NS132 are potent inhibitors of αS aggregation both *in vitro* and *in vivo* models. NS163 and NS132 displayed potent efficacy to rescue PD phenotypes in a DA neuron *C. elegans* PD model, including degeneration of DA neurons, impaired motility rate, decreased dopamine synthesis, behavioral deficits, and increased ROS level. More importantly, OPs were very effective in rescuing PD phenotypes in a post-disease onset model. Overall, we have developed a novel technique to identify potent ligands with tremendous therapeutic potential for the treatment of PD.

## DISCUSSION

The aPPIs are elusive targets as they are of pathological significance and their modulation approaches are directly linked to the discovery of lead therapeutics^1–11^. One of the most effective approaches to modulate aPPIs is the design of synthetic protein mimetics with diverse chemical space, which can complement the sequence and structural topography of aPPI’s interfaces^1–11^. OPs are a class of synthetic protein mimetics that imitate the secondary structure of proteins and have been shown to modulate the aggregation of multiple proteins^30–38^. Despite the overall success of OPs as antagonists, no attention has been directed towards enhancing their antagonist activity, which is predominantly dependent on the extension of the chemical diversity of the side chains on OPs^30–38^ and the onus is on the tedious synthetic route for the generation of OP libraries as the synthesis requires multiple chromatography steps to add individual side chains on OPs.

We have developed a novel fragment-based approach (2D-FAST), in tandem with novel chemistry, to append a large number of side chains with a very diverse chemical space on OPs using a highly efficient synthetic method. We used this approach to modulate a dynamic and transient target, which is the self-assembly of αS, a process linked to the onset of PD^12–18^. PD is the second most common neurodegenerative disorder affecting more than 10 million people worldwide and there is no cure for the disease^20,24^. Therefore, there is a pressing need to identify therapeutic strategies that can prevent or slow down PD. We have used 2D-FAST approach to identify ligands that can modulate the aggregation of αS, which led to the identification of a potent antagonist of *de novo* and fibers catalyzed aggregation of αS, and *in vitro* model of the prion-like spread (PMCA) of αS fibers.

For the first time, we also demonstrated that OPs are synthetically tunable ligands with the ability to improve their cell permeability (from NS132 to NS163) without sacrificing their antagonist activity. Under matched conditions, NS163 was a better antagonist than NS132 in rescuing αS aggregation-mediated PD phenotypes in muscle cells and DA neurons in two *C. elegans* PD models, most likely due to the better cell permeability of the former. Moreover, we have developed a novel post-disease onset PD model to study the effect of ligands on the PD phenotypes mediated by αS aggregation. Both OPs have shown remarkable rescue of the degeneration of DA neurons in this post-disease onset PD model. This study also shows a good correlation between the inhibition of *de novo* and fibers catalyzed aggregation of αS and the rescue of PD phenotypes in DA neurons in UA196 worms in both early-stage and post-disease onset models. Moreover, the inhibition of αS aggregation and prevention of PD phenotypes in *vitro and in vivo* models is a consequence of the interaction of OPs toward the N-terminal of αS and, more specifically, to the suggested aggregation-prone αS sequences^24,47^. The study further highlights the targeting of these sequences as a potential therapeutic approach for the treatment of PD.

We have also shown in the past and the current study that OPs are enzymatically stable in the biological milieu^34,35,37^. Therefore, in the near future, the most potent OPs will be tested for their ability to cross the blood-brain barrier and their pharmacokinetics and pharmacodynamics properties. Subsequently, we will use the most potent OPs in PD mouse models to further assess their pharmaceutical properties and the antagonist activity against PD phenotypes mediated by αS aggregation. The overall neuroprotective effect of OPs on the degeneration of DA neurons (both early and post-disease onset PD models) promises the identification of lead therapeutics for the treatment of PD.

We envision that the OP scaffold-based 2D-FAST could be used to modulate numerous pathological targets, including aPPIs and RNA-protein interactions. This approach is expandable as a greater number of side chains can be appended on OPs because of a very convenient synthetic pathway. We envisage that the novel 2D-FAST will have a broader impact as both the chemistry and the fragment-based approach could be used for various foldamers (aromatic oligoamides, synthetic protein mimetics, and hybrid macrocycles of peptide and synthetic ligands) for the development of potent antagonists for various pathological protein and nucleic acid targets. The combination of the novel and efficient synthetic methodology and the fragment-based approach allows a systematic screening of large chemical space in a succinct time, which will aid in the identification of high affinity and specificity ligands for various pathological targets. To the best of our knowledge, this is the first example of a fragment-based approach for synthetic protein mimetics to successfully identify potent antagonists of a pathological protein target. We envision that the 2D-FAST will have a profound impact on the development of lead therapeutics for various diseases.

## DATA AVAILABILITY

All the datasets generated and analyzed during the current study are also available from the corresponding author.

## ADDITIONAL INFORMATION

Correspondence and requests for materials should be addressed to S.K.

## ACKNOLEDGMENTS

The authors would like to thank the department of chemistry and biochemistry, The Knoebel Institute for Healthy Aging, and the University of Denver for the startup funds. The authors also thank the PinS program (University of Denver) for awarding summer undergraduate fellowship to C.M.D. and T.C.F. We would also like to thank Prof. Marc Diamond’s lab for the wonderful gift of the HEK cells that stably express YFP-labeled A53T αS mutant (αS-_A53T_-YFP). We would also like to thank the American Parkinson Disease Association Research Grant (S.K.) to support the research conducted in this manuscript. We would also like to thank the Parkinson’s Foundation for the summer student fellowship to C.M.D.

## AUTHOR CONTRIBUTIONS

S.K. designed and conceived the project with assistance from N.H.S., J.A.J., and J.A. The synthesis of the OP libraries was carried out by N.H.S. and R.A.D. The biophysical study was carried out by J.A., N.H.S., and T.C.F. The NMR study was carried out by J.A. and N.H.S. The HEK cell-based cytotoxicity and confocal microscopy imaging were carried out by T.D.B. and C.M.D. The *C. elegans-*based *in vivo* experiments (for both PD models), including the confocal imaging, behavioral experiments, dopamine study, and the motility study were carried out by J.A.J. The paper was written by S.K. with assistance from N.H.S., J.A., J.A.J., T.D.B., and R.A.D.

## REFERENCES

1. Lage K. Protein-protein interactions and genetic diseases: the interactome. Biochim. Biophys. Acta. 1842, 1971–1980 (2014).

2. Fry D. C. Targeting protein-protein interactions for drug discovery. Methods Mol. Biol. 1278, 93–106 (2015).

3. Modell A. E., Blosser S. L. & Arora P. S. Systematic targeting of protein-protein interactions. Trends Pharmacol. Sci. 37, 702–713 (2016).

4. Arkin M. R., Tang Y. & Wells J. A. Small-molecule inhibitors of protein-protein interactions: progressing toward the reality. Chem. Biol. 21, 1102–1114 (2014).

5. Gonzalez M. W. & Kann M. G. Chapter 4: protein interactions and disease. PLoS Comput. Biol. 8, e1002819 (2012).

6. Cummings C. G. & Hamilton A. D. Disrupting protein-protein interactions with non-peptidic, small molecule α-helix mimetics. Curr. Opin. Chem. Biol. 14, 341–346 (2010).

7. Cheng, F. et al. Comprehensive characterization of protein-protein interactions perturbed by disease mutations. Nat. Genet. 53, 342–353 (2021).

8. Biza K. V. et al. The amyloid interactome: exploring protein aggregation. PLoS One 12, e0173163 (2017).

9. Brown J. & Horrocks M. H. A sticky situation: aberrant protein-protein interactions in Parkinson’s disease. Semin. Cell Dev. Biol. 99, 65–77 (2020).

10. Vabulas R. M. & Hartl F. U. Aberrant protein interactions in amyloid disease. Cell Cycle 10, 1512–1513 (2011).

11. Richards A. L., Eckhardt M. & Krogan N. J. Mass spectrometry-based protein-protein networks for the study of human diseases. Mol. Syst. Biol. 17, e8792 (2021).

12. Spillantini, M. G. et al. α-Synuclein in lewy bodies. Nature 388, 839–840 (1997).

13. Dettmer, U., Selkoe, D. & Bartels, T. New insights into cellular α-synuclein homeostasis in health and disease. Curr. Opin. Neurobiol. 36, 15–22 (2016).

14. Chiti, F. & Dobson, C. M. Protein misfolding, functional amyloid, and human disease. Annu. Rev. Biochem. 75, 333–366 (2006).

15. Dawson, T. M. & Dawson V. L. Molecular pathways of neurodegeneration in Parkinson’s disease. Science 302, 819–822 (2003).

16. Goedert, M. Alpha-synuclein and neurodegenerative diseases. Nat. Rev. Neurosci. 2, 492–501 (2001).

17. Burré, J., Sharma, M. & Südhof, T. C. Cell biology and pathophysiology of α-Synuclein. Cold Spring Harb. Perspect. Med. 8, a024091 (2018).

18. Kingwell, K. Zeroing in on neurodegenerative α-Synuclein. Nat. Rev. Drug Discov. 16, 371–373 (2017).

19. Pujols, J., Peña-Díaz, S., Pallarès, I. & Ventura S. Chemical chaperones as novel drugs for Parkinson’s disease. Trends Mol. Med. 26, 408–421 (2020).

20. Pujols, J. et al. Small molecule inhibits α-Synuclein aggregation, disrupts amyloid fibrils, and prevents degeneration of dopaminergic neurons. Proc. Natl. Acad. Sci. USA. 115, 10481–10486 (2018).

21. Pineda, A. & Burré, J. Modulating membrane binding of α-Synuclein as a therapeutic strategy. Proc. Natl. Acad. Sci. USA. 114, 1223–1225 (2017).

22. Perni, M. et al. A natural product inhibits the initiation of α-Synuclein aggregation and suppresses its toxicity. Proc. Natl. Acad. Sci. USA. 114, E1009–E1017 (2017).

23. Sangwan, S. et al. Inhibition of synucleinopathic seeding by rationally designed inhibitors. Elife 9, e46775 (2020).

24. Ahmed, J. et al. Foldamers reveal and validate novel therapeutic targets associated with toxic α-Synuclein self-assembly. Nat. Commun. 13, 2273 (2022).

25. Marafon, G. et al. Photoresponsive prion-mimic foldamer to induce controlled protein aggregation. Angew. Chem. Intl. Ed. 60, 5173–5178 (2020).

26. Agerschou, E. D. et al. An engineered monomer binding-protein for α-Synuclein efficiently inhibits the proliferation of amyloid fibrils. ELife 9, e46112 (2019).

27. Mirecka, E. A. et al. Sequestration of a β-hairpin for control of α-Synuclein aggregation. Angew. Chem. Int. Ed. Engl. 53, 4227–4230 (2014).

28. Bavinton, C. E. et al. Rationally designed helical peptidomimetics disrupt α-Synuclein fibrillation. Chem. Commun. 58, 5132–5135 (2022).

29. Oh, M., et al. Potential pharmacological chaperones targeting cancer-associated MCL-1 and Parkinson disease-associated α-Synuclein. Proc. Natl. Acad. Sci. USA. 111, 11007–11012 (2014).

30. Kumar, S., et al. Islet amyloid-induced cell death and bilayer integrity loss share a molecular origin targetable with oligopyridylamide-based α-helical mimetics. Chem. Biol. 22, 369–378 (2015).

31. Saraogi, I. et al. Synthetic alpha-helix mimetics as agonists and antagonists of islet amyloid polypeptide aggregation. Angew. Chem. Int. Ed. Engl. 49, 736–739 (2010).

32. Hebda, J. A., Saraogi, I., Magzoub, M., Hamilton, A. D. & Miranker, A. D. A peptidomimetic approach to targeting pre-amyloidogenic states in type II diabetes. Chem. Biol. 16, 943–950 (2009).

33. Kumar, S. & Hamilton, A. D. α-helix mimetics as modulators of Aβ self-assembly. J. Am. Chem. Soc. 139, 5744–5755 (2017).

34. Kumar, S., Henning-Knechtel, A., Magzoub, M. & Hamilton, A. D. Peptidomimetic-based multidomain targeting offers critical evaluation of Aβ structure and toxic function. J. Am. Chem. Soc. 140, 6562–6574 (2018).

35. Palanikumar, L. et al. Protein mimetic amyloid inhibitor potently abrogates cancer-associated mutant p53 aggregation and restores tumor suppressor function. Nat. Commun. 12, 3962 (2021).

36. Ernst, J. T., Becerril, J., Park, H. S., Yin, H. & Hamilton, A. D. Design and application of an alpha-helix-mimetic scaffold based on an oligoamide-foldamer strategy: antagonism of the Bak BH3/Bcl-xL complex. Angew. Chem. Int. Ed. Engl. 42, 535–539 (2003).

37. Maity, D., Kumar, S., Curreli, F., Debnath, A. K. & Hamilton, A. D. α-helix-mimetic foldamers for targeting HIV-1 TAR RNA. Chemistry 25, 7265–7269 (2019).

38. Yin, H. & Hamilton, A. D. Strategies for targeting protein-protein interactions with synthetic agents. Angew. Chem. Int. Ed. Engl. 44, 4130–4163 (2005).

39. Murray, C. W. & Rees, D. C. The rise of fragment-based drug discovery. Nat. Chem. 1, 187–192 (2009).

40. Scott, D. E., Coyne, A. G., Hudson, S. A. & Abell, C. Fragment-based approaches in drug discovery and chemical biology. Biochemistry 51, 4990–5003 (2012).

41. Li, Q. Application of fragment-based drug discovery to versatile targets. Front. Mol. Biosci. 7, 180 (2020).

42. Rees, D. C., Congreve, M., Murray, C. W. & Carr, R. Fragment-based lead discovery. Nat. Rev. Drug Discov. 3, 660–672 (2004).

43. Magee, T. V. Progress in discovery of small-molecule modulators of protein–protein interactions via fragment screening. Bioorg. Med. Chem. Lett. 25, 2461–2468 (2015).

44. Lu, H. et al. Recent advances in the development of protein–protein interactions modulators: mechanisms and clinical trials. Sig. Transduct. Target Ther. 5, 1–23 (2020).

45. Scott, D. E. et al. Using a fragment-based approach to target protein–protein interactions. ChemBioChem 14, 332–342 (2013).

46. Turnbull, A. P., Boyd, S. M. & Walse, B. Fragment-based drug discovery and protein-protein interactions. RRBC 4, 13–26 (2014).

47. Doherty, C. P. A. et al. A short motif in the N-terminal region of α-synuclein is critical for both aggregation and function. Nat. Struct. Mol. Biol. 27, 249–259 (2020).

48. Kumar, S., Birol, M. & Miranker, A. D. Foldamer scaffolds suggest distinct structures are associated with alternative gains-of-function in a preamyloid toxin. Chem. Commun. 52, 6391–6394 (2016).

49. Levine III, H. Thioflavine T interaction with synthetic Alzheimer’s disease β-amyloid peptides: detection of amyloid aggregation in solution. Protein Science 2, 404–410 (1993).

50. Sanders, D. W. et al. Distinct tau prion strains propagate in cells and mice and define different tauopathies. Neuron 82, 1271–1288 (2014).

51. Tanik, S. A., Schultheiss, C. E., Volpicelli-Daley, L. A., Brunden, K. R. & Lee, V. M. Y. Lewy body-like α-Synuclein aggregates resist degradation and impair macroautophagy. JBC 288, 15194–15210 (2013).

52. Luk, K. C. et al. Exogenous α-synuclein fibrils seed the formation of lewy body-like intracellular inclusions in cultured cells. Proc. Natl. Acad. Sci. USA. 106, 20051–20056 (2009).

53. Volpicelli-Daley, L. A. et al. Exogenous α-Synuclein fibrils induce lewy body pathology leading to synaptic dysfunction and neuron death. Neuron 72, 57–71 (2011).

54. Mahul-Mellier, A. et al. The process of lewy body formation, rather than simply α-Synuclein fibrillization, is one of the major drivers of neurodegeneration. Proc. Natl. Acad. Sci. USA. 117, 4971–4982 (2020).

55. Masuda-Suzukake M. et al. Prion-like spreading of pathological α-Synuclein in brain. Brain 136, 1128–1138 (2013).

56. Desplats P. et al. Inclusion formation and neuronal cell death through neuron-to-neuron transmission of α-synuclein. Proc. Natl. Acad. Sci. USA. 106, 13010–13015 (2009).

57. Barria, M. A., Gonzalez-Romero, D. & Soto, C. Cyclic amplification of prion protein misfolding. Methods Mol. Biol. 849, 199–212 (2012).

58. Shahnawaz, M., et al. Discriminating α-Synuclein strains in Parkinson’s disease and multiple system atrophy. Nature 578, 273–277 (2020).

59. Guerrero-Ferreira, R. et al. Two new polymorphic structures of human full-length α-synuclein fibrils solved by cryo-electron microscopy. ELife 8, e48907 (2019).

60. Strohäker, T. et al. Structural heterogeneity of α-synuclein fibrils amplified from patient brain extracts. Nat. Commun. 10, 5535 (2019).

61. Bousset, L. et al. Structural and functional characterization of two α-synuclein strains. Nat. Commun. 4, 2575 (2013).

62. Kumari, P. et al. Structural insights into α-synuclein monomer-fibril interactions. Proc. Natl. Acad. Sci. USA. 118, e2012171118 (2021).

63. Van Ham, T. J. et al. C. elegans model identifies genetic modifiers of α-synuclein inclusion formation during aging. PLoS Genet. 4, 1000027 (2008).

64. Lassalas, P. et al. Structure property relationships of Carboxylic acid isosteres. J. Med. Chem. 59, 3183–3203 (2016).

65. Currey, H. N., Malinkevich, A., Melquist, P., & Liachko, N. F. ARENA-based activity profiling of tau and TDP-43 transgenic *C. elegans*. MicroPubl. Biol. 2020, 000278 (2020).

66. Ray, A., Martinez, B. A., Berkowitz, L. A., Caldwell, G. A. & Caldwell, K. A. Mitochondrial dysfunction, oxidative stress, and neurodegeneration elicited by a bacterial metabolite in a *C. elegans* Parkinson’s model. Cell Death Dis. 5, e984 (2014).

67. Kuwahara, T. et al. Familial Parkinson mutant α-synuclein causes dopamine neuron dysfunction in transgenic *Caenorhabditis elegans*. J. Biol. Chem. 281, 334–340 (2006).

68. Wang, S. et al. Phosphatidylethanolamine deficiency disrupts α-Synuclein homeostasis in yeast and worm models of Parkinson disease. Proc. Natl. Acad. Sci. USA. 111, E3976–E3985 (2014).

69. Mor, D. E. et al. Metformin rescues Parkinson’s disease phenotypes caused by hyperactive mitochondria. Proc. Natl. Acad. Sci. USA. 117, 26438–26447 (2020).

70. Garcia-Moreno, J. C., Porta de la Riva, M., Martínez-Lara, E., Siles, E. & Cañuelo, A. Tyrosol, a simple phenol from EVOO, targets multiple pathogenic mechanisms of neurodegeneration in a C. elegans model of Parkinson’s disease. Neurobiol. Aging 82, 60–68 (2019).

71. Neto, M. F., Nguyen, Q. H., Marsili, J., McFall, S. M. & Voisine, C. The nematode *Caenorhabditis elegans* displays a chemotaxis behavior to tuberculosis-specific odorants. J. Clin. Tuberc. Other Mycobact. Dis. 4, 44–49 (2016).

72. Park, C. et al. Roles of the ClC chloride channel CLH-1 in food-associated salt chemotaxis behavior of *C. elegans*. Elife 10, 55701 (2021).

73. Margie, O., Palmer, C. & Chin-Sang, I. C. elegans chemotaxis assay. J. Vis. Exp. 74, e50069 (2013).

74. Zhang, X. et al. Scorpion venom heat-resistant peptide protects transgenic *Caenorhabditis elegans* from β-amyloid toxicity. Front. Pharmacol. 7, 227 (2016).

75. Cheng, H-C., Ulane, C. M. & Burke, R. E. Clinical progression in Parkinson disease and the neurobiology of axons. Ann. Neurol. 67, 715–725 (2010).

76. Levin, J. et al. The oligomer modulator anle138b inhibits disease progression in a Parkinson mouse model even with treatment started after disease onset. Acta Neuropathol. 127, 779–780 (2014).

