## Supplementary information file for "A 2D Fragment-Assisted Protein Mimetic Approach to Rescue α-Synuclein Aggregation Mediated Early and Post-Disease Parkinson’s Phenotypes"

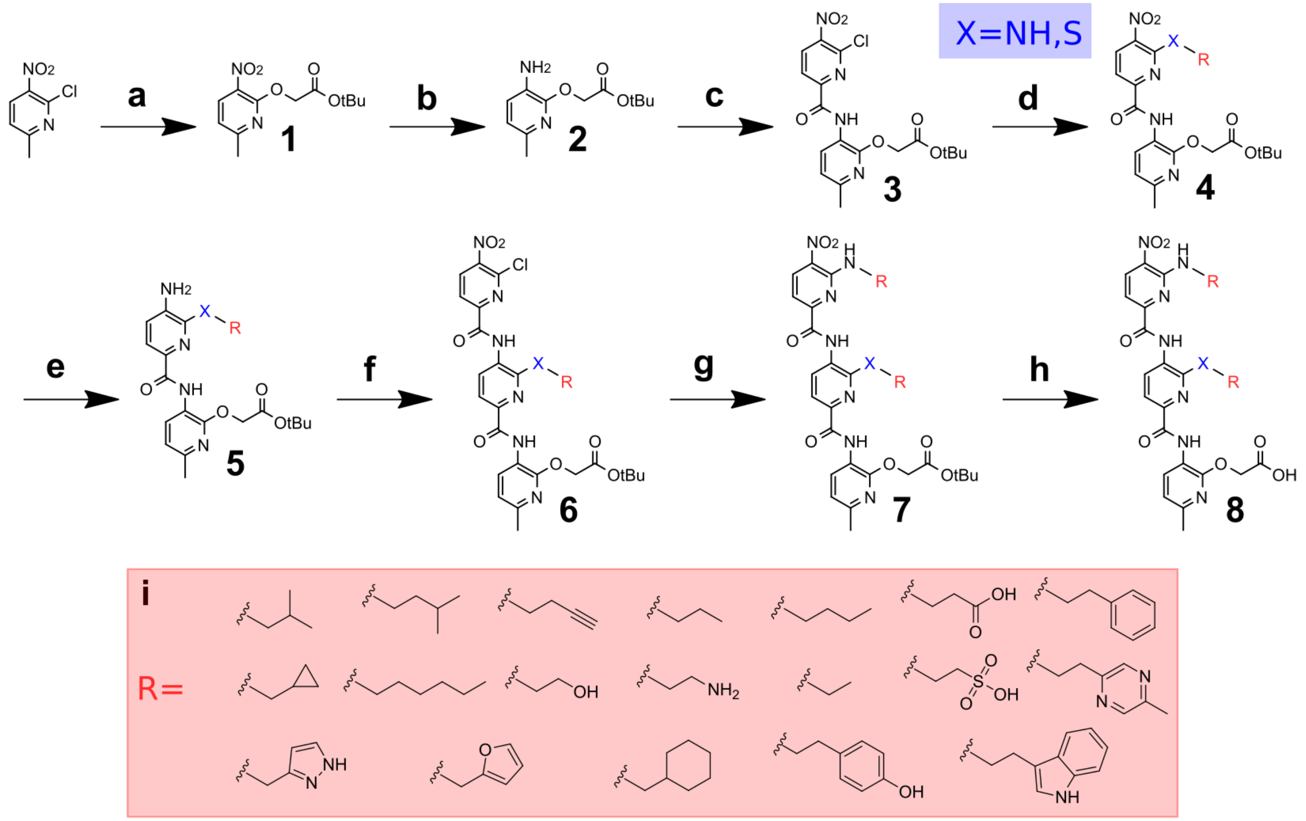

**Supplementary Fig. 1.** General Synthesis of Ops using 2D-FAST. **a**, *tert*-Butyl 2-hydroxyacetate, NaH (60% dispersion in mineral oil), toluene, 50 min at 0 °C, then 5 h at room temperature (r.t.) **b,e**, Palladium on activated carbon (Pd/C), $H_2$ (g), ethyl acetate (EtOAc), 3 h at r.t. **c**, 6-chloro-5-nitropicolinic acid, dichloromethane (DCM, anhydrous), triethylamine (TEA), thionyl chloride 0 °C to r.t., 45 min. **d,g**, Primary amine/thiol, N,N-Diisopropylethylamine (DIPEA), DCM 3 h at r.t. **f**, TEA, 6-chloro-5-nitropicolinoyl chloride, THF (anhydrous) 0 °C to r.t., 40 min **h**, Triethylsilane (TES):trifluoroacetic acid (TFA) (1:1), DCM, 3-6 h at r.t. **i**, Chemical structures of various side chains appended on the dipyridyls and tripyridyls.

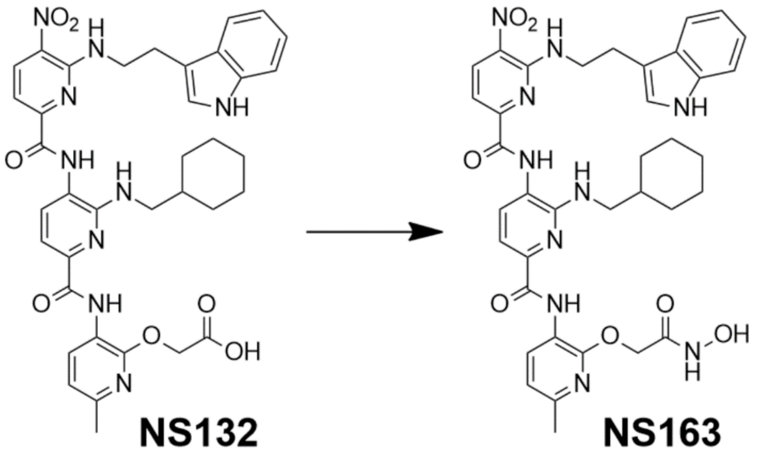


**Supplementary Fig. 2.** Synthesis of NS163. Reagents and conditions: Carbonyldiimidazole (CDI), tetrahydrofuran (THF, anhydrous), stir 1 h at r.t., hydroxylamine hydrochloride, 20 h, r.t.

**Synthesis of Starting Monomer (1)**

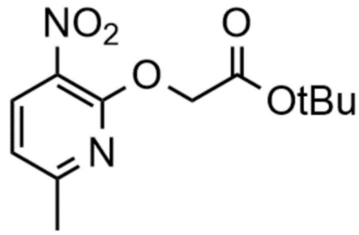

A solution of 6-chloro-5-nitro-2-picoline (2.00 g, 11.6 mmol) in toluene (30 mL) under argon (g) was stirred at 0 °C for 15 min before adding tert-Butyl 2-hydroxyacetate (2.45 g, 18.5 mmol, 1.6 eq.). After stirring the solution for an additional 15 min at 0 °C, NaH (60% dispersion in mineral oil, 0.445 g, 18.5 mmol, 1.6 eq.) was added incrementally over 20 min. The reaction was then stirred at 0 °C for 30 min and then at r.t. for 5 h. After the disappearance of the starting material was observed via TLC, the reaction mixture was partitioned between EtOAc and brine. The organic layers were combined, dried over anhydrous $Na_2SO_4$, and concentrated under vacuum (yellow solid, 3.02 g, 97.2%).

**General method to reduce arylamides (2,5)**

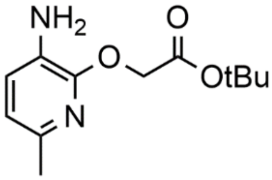

Pd/C (20% by wt., 0.298 g) was added to a solution of nitro arylamide (1.49 g, 5.55 mmol) in EtOAc (15 mL) and continuously stirred while bubbling $H_2$ (g) for 3 h at r.t. After confirming the disappearance of the starting material via TLC, the reaction was filtered and concentrated. The use of a rotary evaporator in this reaction is ill-advised due to the risk of product degradation. The product of this reaction was used in the next step without being further characterized (red/brown oil, 95.6%).

**Synthesis of 6-chloro-5-nitropicolinic acid**

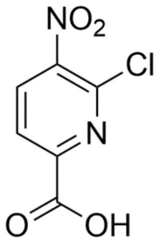


To a stirring solution of 6-chloro-5-nitro-2-picoline (5 g, 29.0 mmol) in conc. $H_2SO_4$ (45 mL), potassium dichromate (11.4 g, 38.6 mmol, 1.3 eq.) was added portionwise over 20 min and refluxed at 60 °C overnight. After confirming the absence of starting material via TLC, the reaction was placed on ice and quenched with water (25 mL). The reaction was then extracted with EtOAc (6×125 mL), dried over $Na_2SO_4$, and concentrated under vacuum to yield the pure product (pale yellow solid, 5.63 g, 96.0%).

**One-pot Amide Coupling (3)**

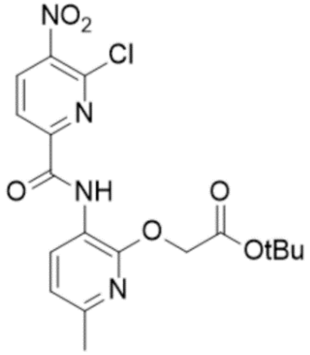


A solution of 6-chloro-5-nitropicolinic acid (0.335 g, 1.65 mmol, 1 eq.) and compound **(2)** (0.394 g, 1.7 mmol) in DCM (anhydrous, 10mL) was equilibrated for 15 min at 0 °C under argon (g). To the stirring solution was added TEA (0.690 mL, 4.96 mmol, 3 eq.) and equilibrated for an additional 60 sec at 0 °C, followed by the addition of thionyl chloride (0.360 mL, 4.96 mmol, 3 eq.). The reaction mixture was stirred for 45 min at r.t. and the disappearance of starting material was confirmed by TLC. The volatiles were removed on a rotovap, and the resulting product was partitioned between EtOAc and 1M HCl (1×15 mL). The organic layer was then washed with 1M NaOH (3×15 mL), followed by brine (1×15 mL), dried over anhydrous $Na_2SO_4$, and concentrated under vacuum (yellow solid, 0.699 g, 94.3%).

**General Method for Aromatic Substitution (4,7)**

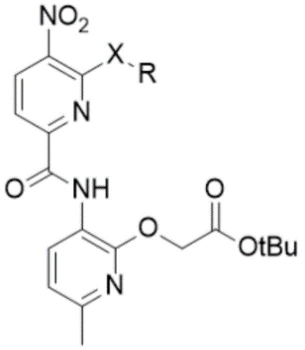

To a solution of compound **(3)** or **(6)** (0.015 g, 0.036 mmol) in DCM (3 mL), a primary amine or thiol (0.355 mmol, 10 eq.) and N,N-Diisopropylethylamine (DIPEA) (0.071 mmol, 20 eq.) were added and stirred at r.t. for 3 h. For reactions involving reagents that exhibit low solubility in DCM, e.g., tryptamine and C-(1-H_Pyrazol-3-yl)-methylamine, DMF was substituted as the solvent. The reaction mixture was dried following confirmation that the starting material had disappeared via TLC. For the removal of non-volatile reagents, flash chromatography is required (10 to 80% EtOAc in hexanes, v/v, over 8 min). The products appear as yellow solids with a yield ranging from 48.0% to 94.3%.

**Synthesis of 6-chloro-5-nitropicolinoyl chloride**

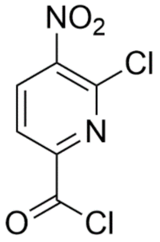


A solution of 6-chloro-5-nitropicolinic acid (0.500 g, 2.47 mmol) in $SOCl_2$ (10 mL) was refluxed overnight at 70 °C. After confirming the reaction was complete via $^1$H NMR, the volatiles were removed under vacuum and the final product was preserved under argon (g) at -20 °C (pale brown solid, 0.516 g, 94.8%).

**General method for amide coupling (6)**

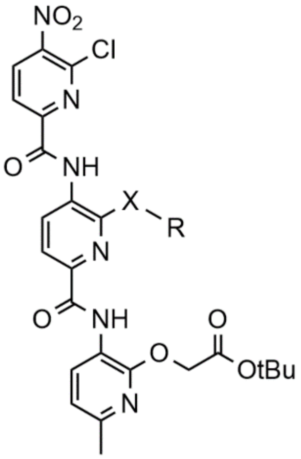


A solution of compound **(5)** (0.033 g, 0.070 mmol) in THF (anhydrous, 4 mL) with TEA (0.028 g, 0.280 mmol, 4 eq.) was stirred at 0 °C for 5 min under argon (g). After equilibration, 6-chloro-5-nitropicolinoyl chloride (0.062 g, 0.280 mmol, 4 eq) in THF (anhydrous, 2 mL) was added dropwise over 60 sec. The solution was stirred at 0 °C for 10 min, then allowed to come to room temperature. TLC confirmed the disappearance of the starting material after 30 min and the resulting solution was partitioned between EtOAc and saturated NaOH. The combined organic layers were washed with brine, dried over $Na_2SO_4$, and concentrated under vacuum (orange solid, 0.042 g, 90.8 %).

**General Method for Deprotection (8)**

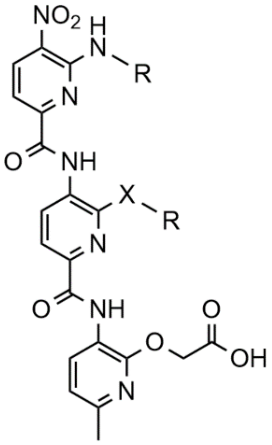


Compound (7) (0.004 - 0.010 g) was dissolved in 3 mL of DCM followed by the addition of 1:1 TES:TFA (0.5 mL each), stirring 60 sec prior to the addition of TFA. The solution was allowed to stir at r.t. for 3 h, confirming the disappearance of starting material by TLC. Upon completion of the reaction, volatiles were removed under vacuum and the resulting product was washed with diethyl ether (3×2 mL, 0 °C) to yield the pure product; Yellow-Orange solids with a yield ranging from 50.8 to 97.1%.

Due to the sensitivity of indoles to acidic conditions, alternative methods of deprotection were required for molecules containing this functional group[1,2]. To a stirring solution of NS72 (0.012 g, 0.022 mmol) in acetonitrile (6 mL), water (40 µL) and elemental iodine (0.002g, 30% mol) were added, and the solution refluxed at 75 °C for 5.5 h. After equilibrium was reached, the solution was extracted with $Na_2S_2O_3$ (aq., 4 mL) and DCM (2×10 mL), and the combined organic layers were dried over $Na_2SO_4$. Flash chromatography was used to recover the starting material (0 to 40% EtOAc in hexanes, v/v, over 6 min, 0.002 g, 16.4%) and to isolate the pure product (0 to 20% methanol in DCM, v/v over 6 min, 0.005g 42.9%).

To a stirring solution of NS132 (0.023g, 0.030 mmol) in DCM (6 mL), zinc bromide ($ZnBr_2$, 0.033 g, 0.147 mmol, 5 eq) was added and stirred 72 h at r.t. Additional solvent was added as needed to replace evaporated solvent. Subsequently, deionized water (30 mL) was added, and the solution stirred vigorously for an additional 2 h. The solution was then extracted with DCM (3×10 mL) and the combined organic layers were washed with brine, dried over $Na_2SO_4$, and concentrated under vacuum (orange solid, 0.014 g, 65.2%).

**Synthesis of NS163**

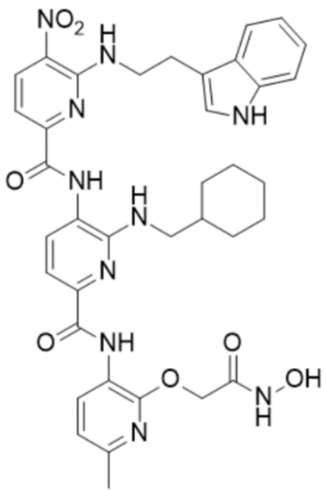


A solution of NS132 (0.009g, 0.011 mmol) and CDI (0.003g, 0.016 mmol, 1.5 eq.) in 2 mL of THF (anhydrous) was stirred at r.t. for 1 h. After adding hydroxylamine hydrochloride (0.002g, 0.022 mmol, 2 eq.), the reaction was stirred at r.t. for 20 hours. The volatiles were removed on a rotovap, and the resulting product was redissolved in EtOAc and washed with brine (3×5 mL). The organic layer was dried over $Na_2SO_4$ and concentrated under vacuum. Flash chromatography (0 to 15 % EtOAc in Hexanes, v/v, over 15 min) yielded the final product (yellow solid, 0.002g, 23.7%).

### Protein Expression and Purification

The plasmid construct pET11-αS (Addgene, Watertown, MA) was chemically transformed into competent BL21(DE3) cells and plated on ampicillin selection plate. A single colony was used to inoculate a 5 mL starter culture which was kept overnight at 37 °C while shaking at 200 rpm. The following day, the starter culture was used to inoculate 4 L of autoclaved Luria Broth (LB). The culture was incubated at 37 °C and shaken at 200 rpm until the optical density ($OD_{600nm}$) reached 0.8. Then, protein expression was induced for 5 h by adding isopropyl β-D-thiogalactoside (IPTG) at a final 1 mM concentration. Cells were then harvested by centrifugation at 7,000 rpm and 4 °C for 15 min. αS purification was carried out using a previously described Osmotic Shock protocol[3]. Briefly, harvested cells were resuspended in Osmotic Shock Buffer (30 mM Tris pH 7.2, 30% sucrose, 2 mM EDTA) and stirred for 15 min at 4 °C. The cells were centrifuged again at 12,000 rpm for 10 min at 4 °C and resuspended in cold Milli-Q $H_2O$. The cell suspension was spiked with $MgCl_2$ at a final 5 mM concentration. The suspension was stirred for an additional 5 min and centrifuged at 6,000 rpm for 10 min to remove cells. The supernatant was boiled at 95 °C for 15 min to precipitate proteins other than αS. The resulting precipitate was removed by centrifugation at 6000xg for 20 min and the solution was loaded on Bio-Scale Macro-Prep High Q ion-exchange column (Bio-Rad, Hercules, CA) equilibrated with 20 mM Tris pH 8.0, 25 mM NaCl, 1 mM EDTA, and αS was eluted using 20 mM Tris pH 8.0, 1 M NaCl, 1 mM EDTA. Pure fractions were collected, buffer exchanged to Milli-Q water using Amicon ultra 3K filters (MilliPoreSigma, Burlington, MA), before lyophilizing and storing the dried protein at -80 °C.

### Aggregation Kinetics Measurement

Aggregation Kinetics were measured according to a previously described protocol[4]. Briefly, 100 μM αS solutions (250 μL, with or without ligands) were prepared in phosphate buffer saline (PBS, 137 mM NaCl, 2.7 mM KCl, 8 mM $Na_2HPO_4$, 2 mM $KH_2PO_4$, pH 7.3). The solutions were placed in a ThermoMixer (Eppendorf, Hamberg, Germany) at 37 °C with constant shaking at 1300 rpm (rpm). At different time points, 5 μL aliquots of each αS solution were added to 95 μL PBS containing 50 μM Thioflavin T (ThT) dye in a Costar black 96 well plate (Corning Inc., Kennebunk, ME). ThT fluorescence intensity was measured ($\lambda_{ex}$ = 450 nm and $\lambda_{em}$ = 490 nm) on an Infinite M200PRO plate reader (Tecan, Männedorf, Switzerland), and plotted against time to give a sigmoidal curve typical of amyloid aggregation. For the single point ThT aggregation, 100 μM αS, in the absence and presence of ligands, was aggerated under the described conditions and the ThT fluorescence was reported at 96 h. The reported ThT intensity values are the average of each experiment conducted in triplicate, with the error bars representing the standard deviations or standard error of the mean (sem) as described in the main manuscript.

### Spin-down assay

An αS solution (100 μM) was prepared in PBS with or without ligands and shaken (13,000 rpm, 37 °C) until the fluorescence plateaued, as observed using the ThT aggregation assay, indicative of the aggregation of αS as a control. The solutions were separated into soluble and insoluble fractions by centrifugation at 22,000×g for 20 min. The fractions were diluted in 2×Laemmli protein sample loading buffer (Bio-rad, Hercules, CA), boiled at 95 °C for 5 min, and ran on a 12% Mini-PROTEAN precast protein gel (Bio-rad, Hercules, CA). The gel was visualized using ChemiDoc MP Imaging System (Bio-rad, Hercules, CA) and band intensity was quantified using Image Lab image acquisition and analysis software (Bio-rad, Hercules, CA). Densitometry values were averaged from three technical replicates and reported with the error bars representing their standard deviations.

### Transmission Electron Microscopy (TEM)

Transmission Electron Microscopy was used to visualize protein assemblies after aggregation experiments as previously described[4]. Briefly, aliquots (5 μL) of αS solutions (with or without ligands) were applied on glow-discharged carbon-coated 300-mesh copper grids for 2 min and dried using Kimwipes (Kimberly-Clark, Irving, TX). The copper grids were negatively stained for 1 min with uranyl acetate (0.75%, w/v). The micrographs were taken on an FEI Tecnai G2 Biotwin TEM at 80 kV accelerating voltages.

**Isothermal Titration Calorimetry (ITC)**

The ITC experiments were performed in a NANO-ITC (TA Instruments, New Castle, DE). For each ITC experiment, a solution of 250 μM NS132 in PBS was serially titrated (2 μL injections every 10 sec via rotary syringe, stirring speed at 300 rpm) into a sample cell containing 350 μL of 15 μM αS in the same buffer conditions at 300 sec intervals. The heat associated with each injection was calculated by integrating each heat burst curve using NanoAnalyze software (TA Instruments, New Castle, DE). The associated heat for each injection of NS132 in αS solution was corrected by subtracting heat resulted from the titration of NS132 into buffer (in 1 × PBS) under identical conditions. The corrected heat values were plotted as a function of the molar ratio of NS132 to αS and the plot was fitted using a one binding site independent model available in NanoAnalyze software. The fitting was carried out using 10,000 iterations in the software without any data constraints during the fitting. The ITC titrations were conducted in triplicate and the reported thermodynamic parameters were calculated as an average of three independent experiments.

**Protein Misfolding Cyclic Amplification Assay (PMCA)**

The PMCA assay was performed according to a previously described protocol[4]. Briefly, an αS solution (100 μM) was prepared in PBS and 60 μL of the solution was placed in a 200 μL Polymerase Chain Reaction (PCR) tube and the mixture was subjected to five 24 h cycles of 1 min shaking (1200 rpm) and 29 min incubation at 37 °C. Every 24 h, 1 μL of αS solution was passaged to seed 60 μL of fresh soluble monomeric αS solution. The seeding cycle was repeated for five days, after which the ThT signal was measured for all five passages. For the preparation of PMCA samples of αS in the presence of NS132, a molar ratio of 1:1 (αS:NS132) was maintained. All experiments were performed in triplicate. The reported ThT intensity values are the average of three separate experiments with the error bars representing the standard deviations.

**Proteinase K Digestion of PMCA Samples**

A Protease K (PK) solution at 50 μg/mL (IBI Scientific, Dubuque, IA) in digestion buffer (10 mM Tris, pH 8.0, 2 mM $CaCl_2$) was diluted (0.1 eq.) into 30 μL of PMCA solution (with or without NS132) and incubated for 30 min at 37 °C. Subsequently, the samples were diluted (1:1, v/v) in SDS Protein Gel Loading Dye 2× (Quality Biological, Gaithersburg, MD) and loaded on Mini-PROTEAN TGX Stain-Free Protein Gel (BioRad, Hercules, CA). The gel was stained using the Fairbanks staining method and imaged using ChemiDoc MP (BioRad, Hercules, CA).

**Parallel Artificial Membrane Permeability Assay (PAMPA)**

The permeability of various ligands was assessed with a PAMPA kit (BioAssay Systems, Hayward, CA) according to a previously described protocol[2]. Briefly, 4% lecithin solution (LS) was prepared in dodecane and 5 μL was applied to the donor plate membrane. Then, 300 μL PBS was added to the acceptor plate. Subsequently, 200 μL of ligand solutions (500 μM) and various permeability standards (high, medium, and low, 500 μM) were added to the donor plate membrane. The donor plate was placed in the acceptor plate and incubated at r.t. for 18 h. The solutions were then moved from the acceptor plate into a clear-bottom 96-well plate (Corning Inc., Corning, NY) and absorbance was recorded at 360 nm for various ligands and 275 nm for permeability standards. The permeability was calculated using Eq. 1 below, where the permeability rate, $C$, equals $7.72 \times 10^{-6}$, $OD_A$ is the absorbance of the acceptor solution, and $OD_E$ is the absorbance of the equilibrium standard.

$$P_e = C \times -\ln\left(1 - \frac{OD_A}{OD_E}\right) cm/s$$

**Heteronuclear Single Quantum Coherence (HSQC) NMR Spectroscopy**

A solution of $^{15}$N αS was prepared by resuspending 1 mg of uniformly labelled $^{15}$N αS (rpeptide, Bogart, GA) in 1 mL Milli-Q water to yield a 70 µM αS solution in 20 mM Tris-HCl pH 7.4, 100 mM NaCl. The protein was buffer exchanged to Milli-Q water, lyophilized, and stored at -80 °C. For the preparation of NMR experiments, the lyophilized $^{15}$N αS powder was dissolved in the NMR buffer (300µL, 20 mM NaPO pH 6.4, 5% $D_2O$, v/v) to make a final concentration of 70 µM. The protein concentration was confirmed with a NanoDrop One (Thermo Scientific, Waltham, MA) at 280 nm using an extinction coefficient of 5960 $M^{-1}cm^{-1}$. The solutions were transferred into Shigemi BMS-005 NMR tube (Shigemi, Tokyo, Japan) and the two-dimensional $^{1}$H-$^{15}$N HSQC NMR experiments were performed on a 600 MHz Bruker Avance Neo (CU Anschutz, NMR core facility, CO) instrument equipped with a triple resonance HCN cryoprobe. Data was collected at 12 °C on Topspin 4.9.0 software (Bruker, Billerica, MA). The HSQC spectra were recorded using a data matrix consisting of 1024 ($t_2$, $^{1}$H) × 160 ($t_1$, $^{15}$N) complex points. Resonance assignments were determined utilizing a previous publication[5] and the data analysis was performed using MestReNova software.

For the HSQC experiments titrating NS132 into $^{15}$N αS, the same experimental conditions were used as above. Incrementally, NS132 was added to a single $^{15}$N αS solution, mixed, kept at r.t. for 10 min and HSQC was recorded under the conditions stated above. For HSQC experiments containing pre-formed fibrils (PFF) of αS, 5.4 eq. of PFF solution (500 µM) was added to monomeric $^{15}$N αS solution, and the spectra was recorded as previously stated.

**Preparation of Lipofectamine solution**

The Lipofectamine solution was made fresh for each transfection to avoid possible denaturation of reagents in a stored stock solution. For each transfection protocol, the total volume of lipofectamine solution required is equivalent to 30% of the total volume of αS/lipofectamine solution needed. The solution was made by mixing OptiMEM media (Thermo Fisher Scientific, Waltham, MA), Lipofectamine, and P3000 reagents (Invitrogen, Carlsbad, CA) at a ratio of 50:1:1 (v/v/v), homogenized, and incubated at r.t. for 20 min. To prepare a 5 µM αS/Lipofectamine solution, 250 µL of an aggregated solution of αS (100 µM) was diluted with OptiMEM to a final volume of 3.5 mL and sonicated for 10 min. Subsequently, the Lipofectamine solution (1.5 mL) was combined with the αS protein solution for a total volume of 5 mL. The resulting solution (5 µM) was homogenized and added into their respective wells for transfection of cells.

**MTT (3-(4,5-dimethylthiazol-2-yl)-2,5-diphenyltetrazolium bromide) reduction Assay**

This assay was performed based on a previous protocol with slight modifications[4]. Unless stated otherwise, the density of Human Embryonic Kidney (HEK) cells (Generous gift by Prof. Mark Diamond' lab) expressing αS$_{A53T}$-YFP was 150,000 cells/mL for each experiment. The HEK cells were plated (100 µL/ well) and incubated (24 h, 37 °C, 5% $CO_2$) in a Costar 96-well transparent plate (Corning, Kennebunk, ME) with complete DMEM (Thermo Fisher Scientific, Waltham, MA) supplemented with 10% FBS (Hyclone, Logan, UT) media containing 1% penicillin/streptomycin (Life Technologies Corporation, Grand Island, NY). The cell solution replaced with OptiMEM (Life Technologies Co., Grand Island, NY) containing αS aggregated solution in the absence and presence of NS132 at an equimolar ratio, containing the lipofectamine solution. A solution of MTT dye (10 µL, 5 mg/mL in PBS) was added to each well and the plate was covered with aluminum foil and incubated for 3 h. Afterward, the solution was carefully aspirated out without disturbing the formazan crystals, and replaced with DMSO (100 µL/well). The crystals were dissolved with gentle shaking of the plate, and subsequently, the absorbance was measured at 570 nm using an Infinite M200 Pro Plate Reader (Tecan, Grŏdig, Austria). For all MTT assays, four biological replicates were performed and each biological assay consisted of four technical replicates.

**ProteoStat-Dye Assay**

This assay was performed based on a previous protocol with slight modifications[4]. The HEK cells (αS$_{A53T}$-YFP) were plated in 35 mm dishes (Celltreat, Pepperell, MA) with a density of 300,000 cells/dish. The cells were then transfected with αS fibrils (aggregated in the absence and presence of NS132 at an equal molar ratio) using lipofectamine and incubated for 24 h. The Petri dishes were checked for puncta using an Axio Observer Microscope (Carl Zeiss Microscopy, Göttingen, Germany) before measuring intracellular aggregation using the PROTEOSTAT Protein Aggregation Assay Kit (Enzolifesciences, Farmingdale, NY). The media containing dead cells was then transferred into 15 mL Falcon tubes (Corning Science Mexico, Tamaulipas, Mexico). Subsequently, a solution of Detachin (Genlantis, San Diego, CA) was added to detach the live cells, which were then transferred into their respective 15 mL Falcon tubes. After the cells were transferred, the solution was centrifuged for 10 min, and the detachin was aspirated out and replaced with 500 µL PBS. The resulting cell pellets were then homogenized, transferred into an autoclaved 1.7 mL microcentrifuge tube, and counted to normalize the data. The microcentrifuge tubes were then centrifuged for 5 min at 4000 rpm and 4 °C (unless stated otherwise, centrifugation parameters remained the same throughout the assay). The process was repeated two more times with 500 µL PBS to properly wash the cell pellets and remove any excess of detachin or FBS media. The cells were treated with 4% paraformaldehyde (Electron Microscopy Sciences, Hatfield, PA), homogenized, and incubated on ice for 30 min. The tubes were then centrifuged for 5 min, the paraformaldehyde solution was replaced with 100 µL PBS, and centrifuged for another 5 min. The 100 µL PBS was replaced with 500 µL PBS containing 0.15% (v/v) Triton X-100 (Oakwood Chemical, Estill, SC), homogenized, and incubated on ice for 20 min. The tubes were then centrifuged again, and the supernatant was replaced with 100 µL PBS. Following another round of centrifugation, the supernatant was replaced with 375 µL PBS containing the ProteoStat buffer (10%, v/v) and ProteoStat Dye (1%, v/v), according to the manufacturer's guidelines. The cell solutions were homogenized, protected from ambient light, and incubated at r.t. for 20 min. The tubes were centrifuged, and the dye solution was replaced with 400 µL PBS. The cell solution was homogenized and equally aliquoted into the desired number of wells in a Costar 96-well flat black plate and fluorescence was measured ($\lambda_{ex}$ = 550 nm, $\lambda_{em}$ = 600 nm) on an Infinite M200 Pro Plate Reader. For each condition, four biological replicates were performed, each consisting of three technical replicates.

### Immunofluorescence staining and confocal imaging

The staining and confocal imaging of HEK cells ($\alpha S_{A53T}$-YFP) was carried out using a previously reported protocol with slight modifications[4]. The HEK cells were plated (300 µL/well) in a µ-slide 8-well plate (Ibidi, Gräfelfing, Germany) and incubated (24 h at 37 °C and 5% $CO_2$). The cells were then transfected with αS fibrils aggregated in the absence and presence of NS132, using lipofectamine and incubated for 24 h as previously described. The transfection was confirmed by the appearance of puncta within the plated cells using an Axio Observer microscope. The cells were then fixed with 4% paraformaldehyde for 10 min and subsequently washed with PBS (3×). The paraformaldehyde was then replaced with PBS containing 0.15% Triton X-100 for 10 min and washed with PBS (3×). A PBS solution containing 1% (w/v) BSA (Thermo-Fischer Scientific, Rockford, IL) and 0.1% (v/v) Tween-20 (Sigma-Aldrich, St. Louis, MO) was added and incubated at r.t. for an additional 30 min then washed with PBS (3×). Next, the cells were stained with anti-α-Synuclein Phospho Ser129 (phosphorylated αS at serine residue 129) mouse antibody (BioLegend, San Diego, CA) for 1 h and washed with PBS (3×). The cells were then treated with Donkey anti-mouse tagged with Alexa Fluor Plus 647 secondary antibody (Invitrogen, Rockford, IL) for 1 h and washed with PBS (3×). Lastly, the cell nuclei were stained with DAPI (Cayman Chemical Company, Ann Arbor, MI) for 10 min and washed with PBS (2×). Confocal imaging was conducted on an Olympus Fluoview (FV3000 confocal/2-photon microscope). The images produced by the confocal microscope were analyzed on OlympusViewer in ImageJ processing software.

### Culture methods for *C. elegans* strains

The N2 (wild-type *C. elegans* Bristol strain), NL5901 (*C. elegans* model of PD), and *Escherichia coli* OP50 (*E. coli*, a uracil requiring mutant) strains were obtained from Caenorhabditis Genomics Center (Minneapolis, MN). Dopaminergic neuron-specific RNAi UA196 strain was generously donated by the laboratory of Dr. Guy Caldwell (Department of Biological Science, The University of Alabama, Tuscaloosa, AL, United States)[6]. The worms were maintained at standard conditions on nematode growth media (NGM) agar in 60 mm plates (CytoOne, Ocala, FL) using *E. coli* OP50 as the food source following the previous protocols[4,7,8]. NGM agar plates, M9 buffer (3 g $KH_2PO_4$, 6 g $Na_2HPO_4$, 5 g NaCl, 1 mL 1 M $MgSO_4$, milli-Q $H_2O$ to 1 L), Chemotaxis (CTX) media plates (2% Agar, 5 mM $KH_2PO_4$, 1 mM $CaCl_2$, and 1 mM $MgSO_4$), CTX buffer (5 mM $KH_2PO_4$, 1 mM $CaCl_2$, and 1 mM $MgSO_4$) and OP50 solution at 0.5 $OD_{600nm}$ were prepared using previous protocols[4,8–10].

### Motility assay for *C. elegans* (N2, NL5901 and UA196)

The motility assay was conducted based on previously described protocols[4,8,11,12] with slight modifications. On day one, N2 and NL5901 strains were synchronized using the bleaching process, involving egglay[7] and incubation of the eggs (23 °C, 30 h) on a solution of NGM in 60 mm culture plates (CytoOne, Ocala, FL) with OP50 (350 µL, 0.5 $OD_{600nm}$) as a food source. On day two, the worms were transferred (using M9 buffer) to 35 mm NGM plates (CellTreat Scientific, Pepperell, MA) containing 75 µM Fluorodeoxyuridine (FUDR; to prevent worm reproduction and ensure that equal ages of worms were used for the experiment)[4,8] and ligand (NS132 and NS163, 50 µM) and incubated at r.t. for 24 h (up to day four). On day three, fresh stock of OP50 was prepared by diluting 1 µL of OP50 in 5 mL of LB Miller media (Neogen, Lansing, MI) and incubated in a shaking incubator (Eppendorf, Hamburg, Germany) at 37 °C and 200 rpm for ~24 h. On day four, liquid media was prepared with 67.28% (v/v) of M9 buffer, 0.018% (75 µM) of FUDR solution (v/v), 0.1% of 1 M magnesium sulfate (v/v), 0.1% of 1 M calcium chloride (v/v), 2.5% of 1 M potassium phosphate solution (pH 6, v/v)[8], and 30% of 0.5 $OD_{600nm}$ OP50 (v/v). The experiment was carried out using a sterile 24 well plate (CellTreat Scientific, Pepperell, MA) containing liquid media (500 µL/well). The three conditions included: (1) N2 worms (positive control), (2) NL5901 worms (untreated, negative control) and (3) NL5901 worms treated with the ligands (50 µM NS132 and NS163). A total of 50 worms per well were manually transferred into a 24 well plate and incubated at r.t. for 24 h. On day four, before conducting each motility assay, the 24 well plate was mechanically tapped for about 30 sec to make the worms more active in the liquid media. The assay was started on day four where 20 activity scores per well were collected using the WMicroTracker ARENA plate reader (Phylumtech, Santa Fe, Argentina) at 23 °C for 1 h per day over a 14-day period. For each condition, four biological replicates were performed and each biological replicate consisted of two technical replicates. For each condition, the data were expressed as mean and the error bars report the s.e.m. (n = 4 independent experiments and each n consisted of a minimum of two technical replicates). The motility rate experiment was repeated for the UA196 strain (in the absence and presence of NS132 and NS163 at 50 µM) following the same conditions and protocol as used for the NL5901 strain.

**Motility assay for *C. elegans* (N2 and UA196) in the presence of dopamine and ligands**

The motility assay was conducted for UA196 and N2 worms in the absence and presence of Dopamine (2 mM, treated on day two) and ligand (50 µM NS163, treated on day two) using the previous protocols[4,8,11,12]. The six worm conditions for this experiment included: (1,2) N2 in the absence and presence of Dopamine (2 mM), (3) UA196 (negative untreated control), (4) UA196 treated with 2 mM Dopamine, (5) UA196 treated with ligand (50 µM NS163), and (6) UA196 treated with both Dopamine (2 mM) and ligand (50 µM NS163). The worms were treated with NS163 and/or dopamine on day two and the motility assay was conducted as described in the previous section.

**Confocal microscopy of early stage treated *C. elegans* (NL5901 and UA196) with ligands**

This experiment was performed based on previously described protocols[4,11,13] with slight modifications. The NL5901 worms in the absence and presence of ligands (50 µM NS132 or NS163 treated on day two) were used for this experiment under conditions identical to the motility assay. At least 10 worms per condition were transferred to a cover slide containing an anesthetic (40 mM sodium azide), and mounted on glass microscope slide containing 2% agarose pads for imaging[13]. The images of the worms were collected using an Olympus Fluoview FV3000 confocal/2-photon microscope (40 x Plan-Apo/1.3 NA objective with DIC capability) from day five through day nine (for NL5901 strain) and processed using the OlympusViewer in ImageJ software[4]. The inclusions of aggregated αS (in the muscle cells of NL5901 strain) were manually counted (10 worms per condition) for the five-day imaging period. For UA196 strain, the confocal imaging of the worms was conducted on day 3, day 5, day 10, and day 15. For each condition, three biological replicates were performed and at least 10 technical replicates were used for each biological replicate. The data were expressed as mean and the error bars report the s.e.m. (n = 3 or 4 independent experiments and each n consisted of a minimum of ten technical replicates).

**Confocal imaging of a post-disease onset PD model of UA196 worms in the presence of ligands**

The UA196 worms were synchronized by bleaching and transferred into FUDR plates as described in the previous section. For this experiment, the UA196 worms were treated with 50 µM ligands (NS132 or NS163) on day five and then incubated at r.t. Confocal imaging was performed with worms immediately prior to treatment with the ligand on day five. The confocal imaging of UA196 worms was then performed on day 10 and day 15 in the absence and presence of ligands. For each condition, three biological replicates were performed and at least 10 technical replicates were used for each biological replicate. The data were expressed as mean and the error bars report the s.e.m. (n = 3 or 4 independent experiments and each n consisted of a minimum of ten technical replicates).

**Chemotaxis Assay for *C. elegans* (N2 and UA196) in the presence of ligands**

The previous protocols[9,10] with slight modifications were used to perform this assay in CTX solid media 35 mm plates (CellTreat Scientific, Pepperell, MA). The UA196 (treated and untreated with 50 µM ligands on day two) and N2 worms were prepared similar to the motility assay conditions as described above. Three CTX 35 mm plates (for the N2, treated and untreated UA196 strains) were divided into four equal quadrants designated A and C (the left diagonal quadrant), B and D (the right diagonal quadrant). A solution of *E coli* (attractant) was placed at ~ 0.4 cm from the edge of quadrants B and D and ethanol (10 µL, repellant) was placed at ~ 0.4 cm from the edge of quadrants A and C. The solutions in the quadrants were allowed to dry for about 1 h at r.t. to develop the gradient. CTX buffer (1 mL) was used to transfer adult worms from the 35 mm plates (in the absence and presence of ligands) for each condition in sterile 1.7 mL microcentrifuge tubes (Eppendorf, Hamburg, Germany). The worm solution was centrifuged for 10 sec using a mini-Vortex mixer (VWR, China) to allow the worm pellets to settle at the bottom of the microcentrifuge tubes. The supernatant solution was aspirated out and replaced with CTX buffer. This buffer exchange step was conducted three times to ensure that the worms are re-suspended in only 100 µL of CTX buffer. After homogenizing the worm solution, 10 µL of the worm solution was transferred into a blank solid media 35 mm plate and the number of worms were counted under an Olympus microscope (SZ-6145, Waltham, MA) in triplicate. A worm suspension was then prepared with the CTX buffer containing 50 worms/10 µL. Approximately 50 worms were transferred to the center of the CTX plate, and the lids were covered with parafilm (Bemis Company, Inc., Neenah, WI). Subsequently, the worm activity was monitored using a WMicroTracker ARENA plate reader (Phylumtech, Santa Fe, Argentina) at ~23 °C for 2 h. The report was generated using MapPlot option on the plate reader. Three biological replicates of this experiment were performed. The experiment was conducted for the UA196 worms (in the absence and presence of ligands) and N2 worms on day three and day 10 of the aging process. For each condition, three biological replicates were performed and at least two technical replicates were used for each biological replicate. The data were expressed as mean and the error bars report the s.e.m. (n = 3 independent experiments and each n consisted of two technical replicates).

**Measurement of the ROS level in UA196 worms in the presence of ligands (Quantification of ROS and confocal imaging)**

A fluorescent probe 2',7'-dichlorofluorescein diacetate ($H_2$DCFDA) was used to measure the intracellular ROS based on previously established protocols[11,14,15] with slight modifications. The UA196 worms were synchronized and treated with ligands (NS163 and NS132, 50 µM) on day two as described in the previous sections. On day eight, the worms were transferred into 1.7 mL microcentrifuge tubes using M9 buffer (1 mL). The samples were centrifuged for 2 min at 2,500 rpm and 20 °C. Subsequently, 0.8 mL of the supernatant was discarded. After, homogenizing the worm pellet in the remaining solution, 10 µL of the suspension was placed onto a glass slide for counting using an Olympus microscope (SZ-6145, Waltham, MA) in triplicate. The worm solution was then diluted with M9 buffer to approximately 50 worms/10 µL. Each well of a Costar 96-well black plate (Corning, Kennebunk, ME) contained M9 buffer (40 µL), worm solution (10 µL), and $H_2$DCFDA (50 µL, 50 µM in DMSO). For the vehicle, 50 µL of M9 buffer and 50 µL of $H_2$DCFDA reaction solution were placed in the wells (as a control). Subsequently, the Costar 96-well plate was gently shaken for 30 sec at r.t. and the fluorescence intensity was quantified ($\lambda_{ex}$ = 485 nm and $\lambda_{em}$ = 530 nm) at multiple time points (0 to 120 min) using the Infinite M200 Pro Plate Reader.

For confocal imaging, the same UA196 worms used for running the ROS quantification, were transferred into different 35 mm NGM plates, air-dried using the fume hood, and transferred to a cover slide containing an anesthetic (40 mM sodium azide), and mounted on a glass microscope slide containing 2% agarose pads for imaging as described previously.[13] This experiment consisted of three biological replicates and each biological replicate consisted of three technical replicates. The data were expressed as mean and the error bars report the sd's (n = 3 independent experiments and each n consisted of three technical replicates).

**Measurement of intracellular ROS level in a post-disease onset PD model of UA196 worms in the presence of ligands**

The UA196 worms were synchronized by bleaching and transferred into FUDR treated plates as described in the previous sections. On day five, the quantification of ROS level was conducted as described in the previous section. Subsequently, the UA196 worms were treated with ligands (NS132 and NS163, 50 µM) on day five and then incubated at r.t. for three days. On day eight, the quantification of ROS level was carried out for UA196 worms in the absence and presence of ligands. This experiment consisted of one biological replicate and four technical replicates. This experiment consisted of three biological replicates and each biological replicate consisted of three technical replicates. The data were expressed as mean and the error bars report the sd's (n = 3 independent experiments and each n consisted of three technical replicates).

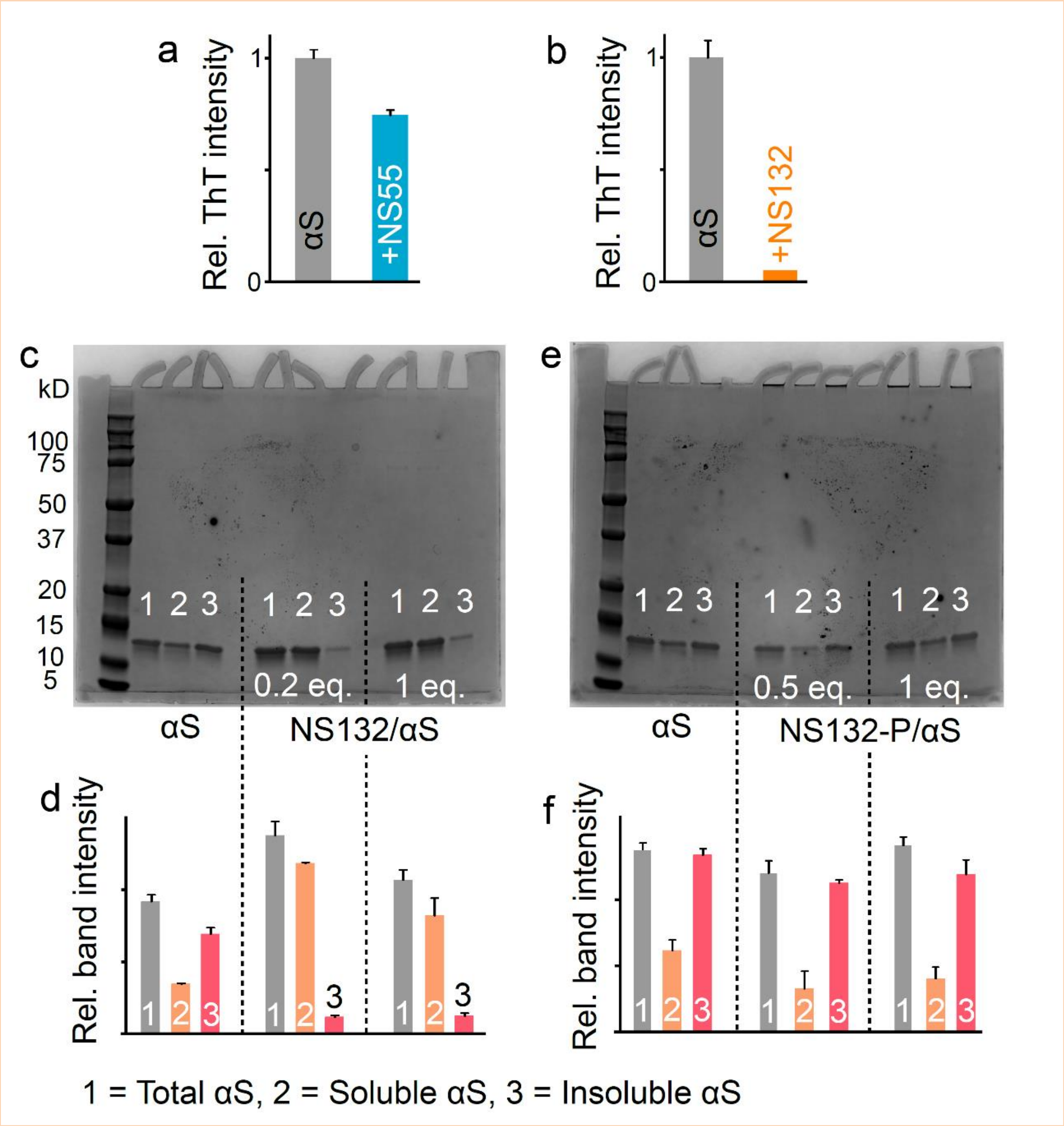


**Supplementary Fig. 3. a**, The graphical representation of the ThT intensity of 100 μM αS aggregation for four days in the absence and presence of NS55 (**a**) and NS132 (**b**) in 1 × PBS at substoichiometric ratio (for NS55, αS:ligand, 1:0.5, and for NS132, αS:ligand, 1:0.1, ). **c**, Representative SDS-PAGE gel images and band intensities of 100 μM αS aggregation for four days in the absence and presence of NS132 (**c,d**) and NS132-P (**e,f**) at the indicated molar ratios. The ThT experiments were conducted three times and the reported change in the ThT intensity was an average of three separate experiments. The gel shift assay experiments were conducted three times and the reported intensity changes were an average of three separate experiments. The data were expressed as mean and the error bars report the standard deviation (s.d.) (n = 3 independent experiments and each n consisted of three technical replicates).

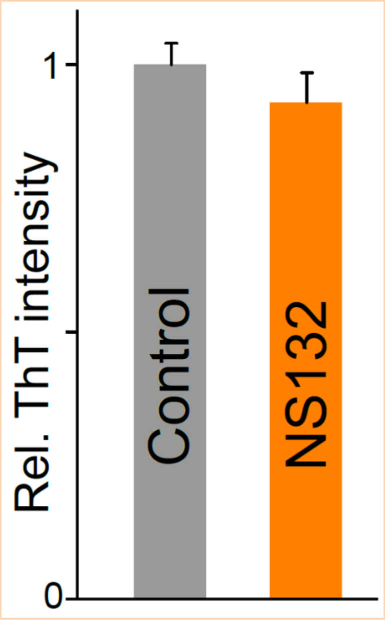


**Supplementary Fig. 4.** The comparison of the fluorescence intensity of the ThT dye (50 μM) in the absence and presence of NS132 (100 μM) in 1 × PBS. The ThT experiments were conducted three times and the reported change in the ThT intensity was an average of three separate experiments. The data were expressed as mean and the error bars report the s.d. (n = 3 independent experiments and each n consisted of three technical replicates).

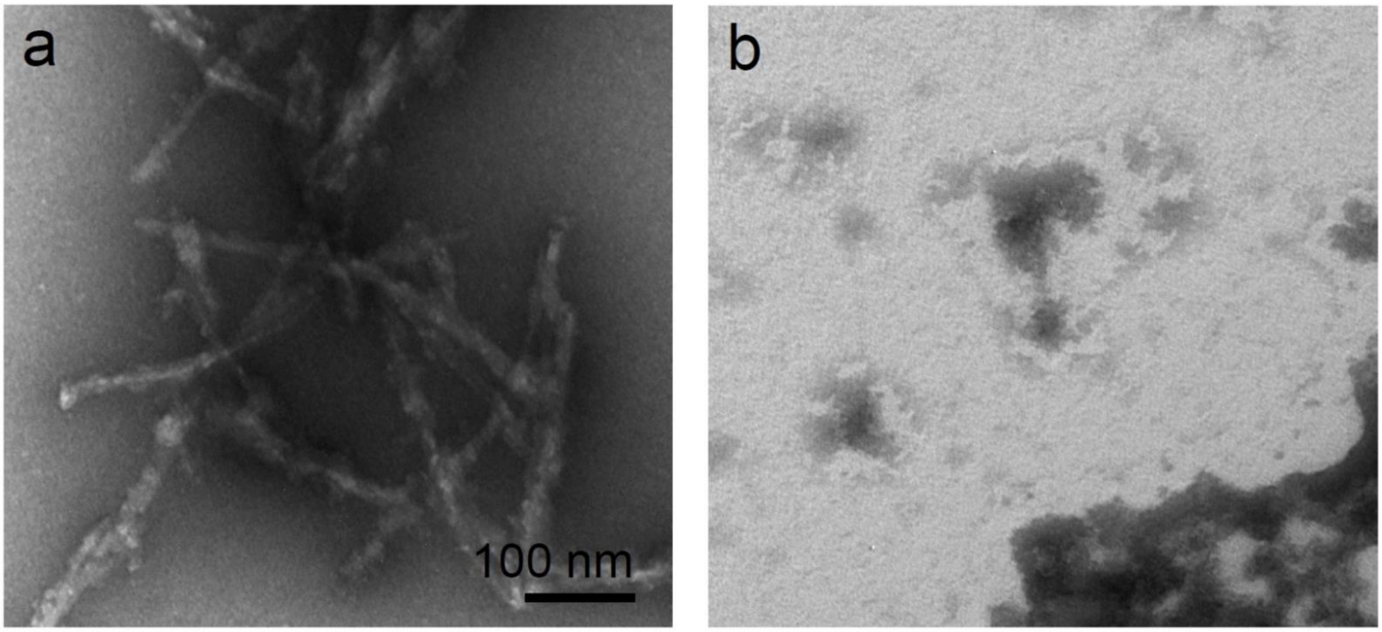


**Supplementary Fig. 5.** The TEM images of 100 μM αS solution aggregated for four days under aggregation conditions (1 × PBS buffer) in the absence (a) and presence (b) of NS163 at an equimolar ratio.

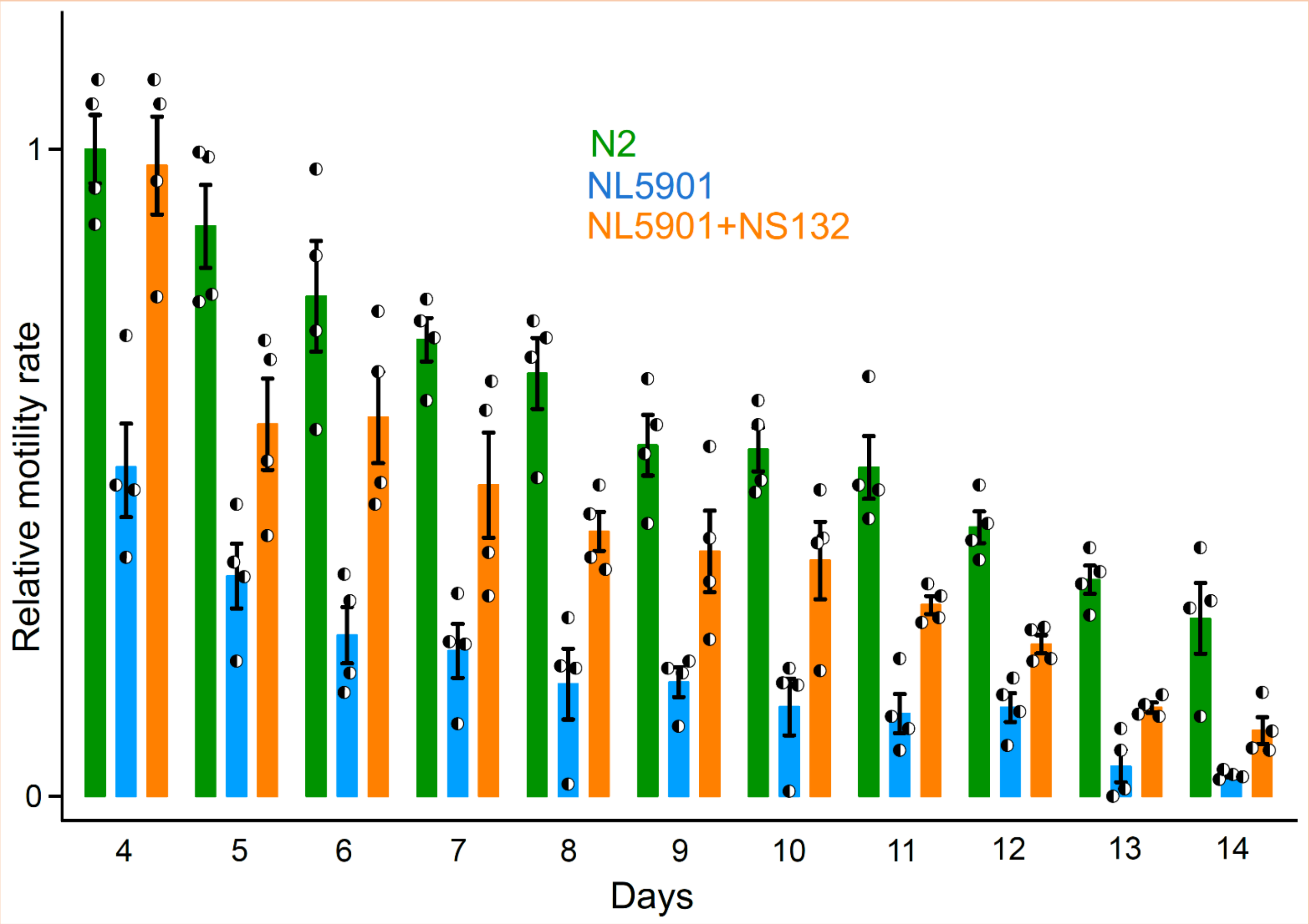


**Supplementary Fig. 6. The intracellular inhibition of αS aggregation by NS132 in an *in vivo* PD model.** The comparison of the motility rate of N2 and NL5901 and statistics in the absence and presence of 50 µM NS132 (treatment on day two). For motility rate experiment, a total of 50 worms were used in duplicate for each experiment and each condition consisted of at least four independent experiments. The data were expressed as mean and the error bars report the s.e.m. (n = 4 independent experiments and each n consisted of two technical replicates). The statistical analysis was performed using ANOVA with Tukey's multiple comparison test. *$p < 0.05$, **$p < 0.01$, ***$p < 0.001$.

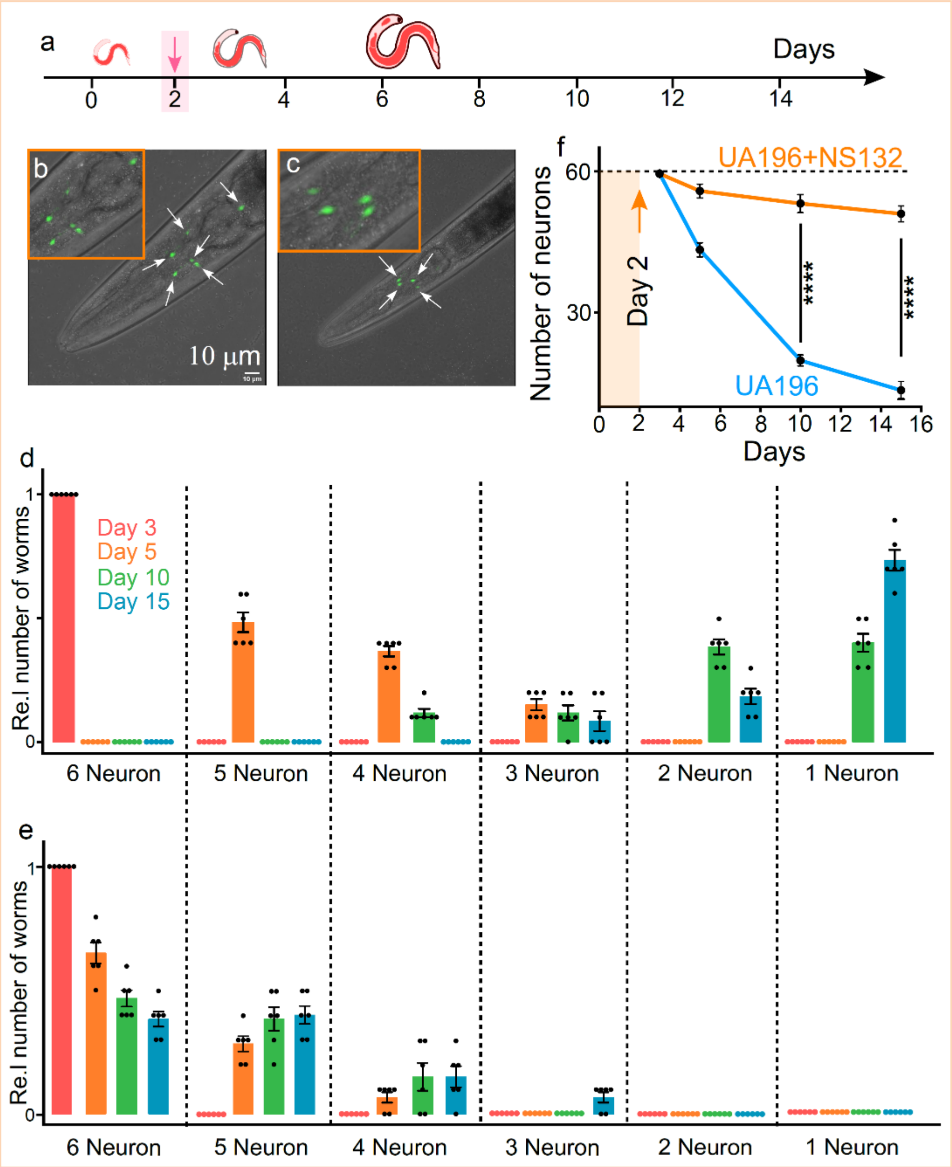


**Supplementary Fig. 7. Neuroprotective effect of NS132 on the degeneration of DA neurons. a,** Schematic of the aging process of UA196 worms and their treatment with the ligands. Representative confocal images of UA196 worms in the presence of 50 µM NS132 on day 5 (**b**) and day 15 (**c**). The healthy DA neurons (white arrow) in UA196 worms on day 5. The relative number of neurons in UA196 worms during the aging process in the absence (**d**) and presence of 50 µM NS132 (**e**). **f**, Statistics for the total number of neurons in UA196 worms during the aging process in the absence (blue) and presence (orange) of 50 µM NS132. For each confocal imaging experiment (**b,c**), at least 10 worms were used, and the healthy neurons were counted manually, and each condition (day) consisted of six independent experiments with freshly bleached worms. For motility rate experiment, a total of 50 worms were used in duplicate for each experiment and each condition consisted of four independent experiments with freshly bleached worms. The data were expressed as mean and the error bars report the s.e.m. (n = at least 4 independent experiments and each n consisted of at least two technical replicates). The statistical analysis was performed using ANOVA with Tukey's multiple comparison test. $*p < 0.05$, $**p < 0.01$, $***p < 0.001$, $****p < 0.0001$.

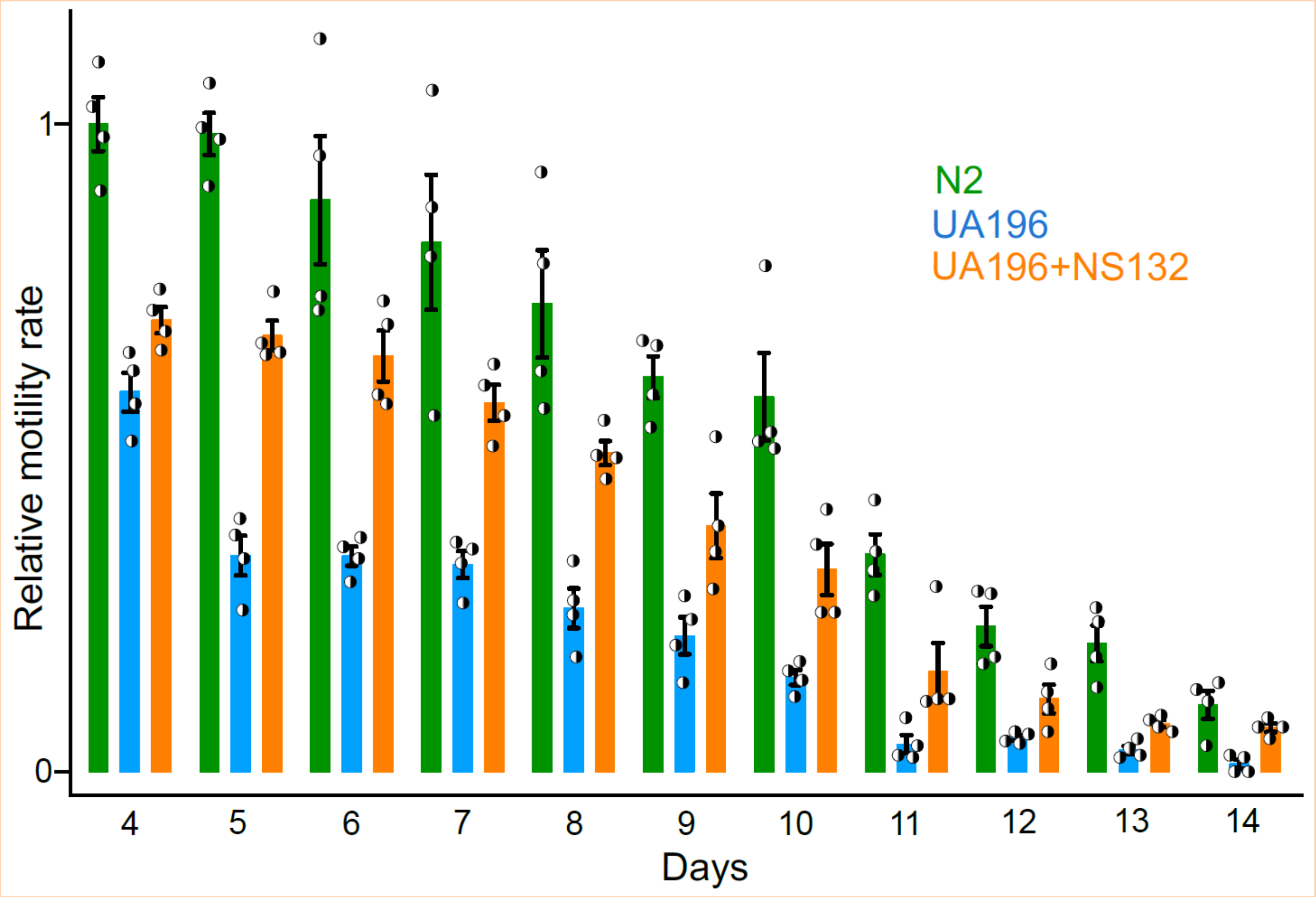


**Supplementary Fig. 8. The rescue of motility rate in UA196 worms by NS132.** The comparison of the motility rate of N2 and UA196 in the absence and presence of 50 µM NS132 (treatment on day two). For motility rate experiment, a total of 50 worms were used in duplicate for each experiment and each condition consisted of four independent experiments. The data were expressed as mean and the error bars report the s.e.m. (n = 4 independent experiments and each n consisted of two technical replicates). The statistical analysis was performed using ANOVA with Tukey's multiple comparison test. *$p < 0.05$, **$p < 0.01$, ***$p < 0.001$.

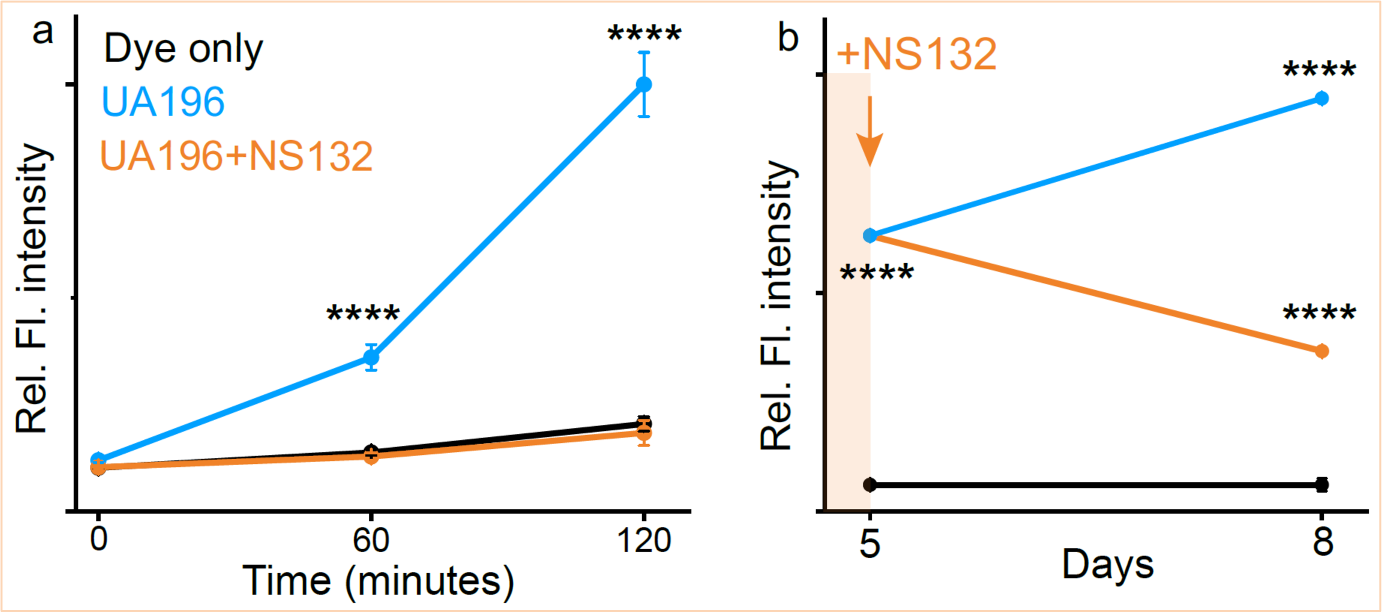


**Supplementary Fig. 9. The effect of NS132 on the ROS level in UA196 worms. a**, The comparison of the ROS level in UA196 worms in the absence and presence of 50 µM NS132 at the indicated time points. The UA196 worms were treated with NS132 on day 2 and the ROS level was measured on day 10. **b**, The comparison of the ROS level in UA196 worms (blue) on day 5 and day 8 when treated (orange) with 50 µM NS132 on day 5. For ROS level quantification, at least 50 worms were used and each condition consisted of three independent experiments with freshly bleached worms. The data were expressed as mean and the error bars report the s.d. ($n = 3$ independent experiments and each n consisted of three technical replicates). The statistical analysis was performed using ANOVA with Tukey's multiple comparison test. *$p < 0.05$, **$p < 0.01$, ***$p < 0.001$, ****$p < 0.0001$.

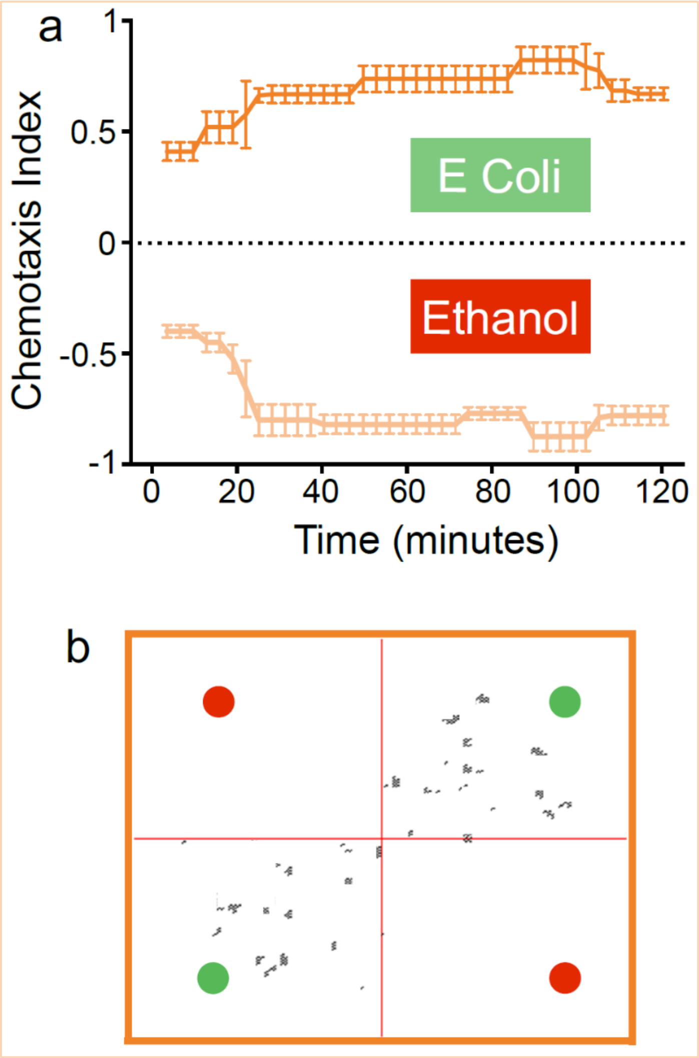


**Supplementary Fig. 10. The effect of NS132 on the behavioral deficits mediated by αS aggregation in DA neurons in UA196 worms. a,** The CI graph for UA196 worms treated with 50 μM NS132 (treatment on day two) under the indicated conditions on day 10. **b**, The snapshots at 60 min. of the animated videos (Movie S8) collected for the CI for UA196+NS132 under the indicated conditions. For chemotaxis assays, a total of 50 worms were used in duplicate for each experiment and each condition consisted of three independent experiments with freshly bleached worms. The data were expressed as mean and the error bars report the s.e.m. ($n = 3$ independent experiments and each n consisted of two technical replicates). The statistical analysis was performed using ANOVA with Tukey's multiple comparison test. *$p < 0.05$, **$p < 0.01$, ***$p < 0.001$, ****$p < 0.0001$.

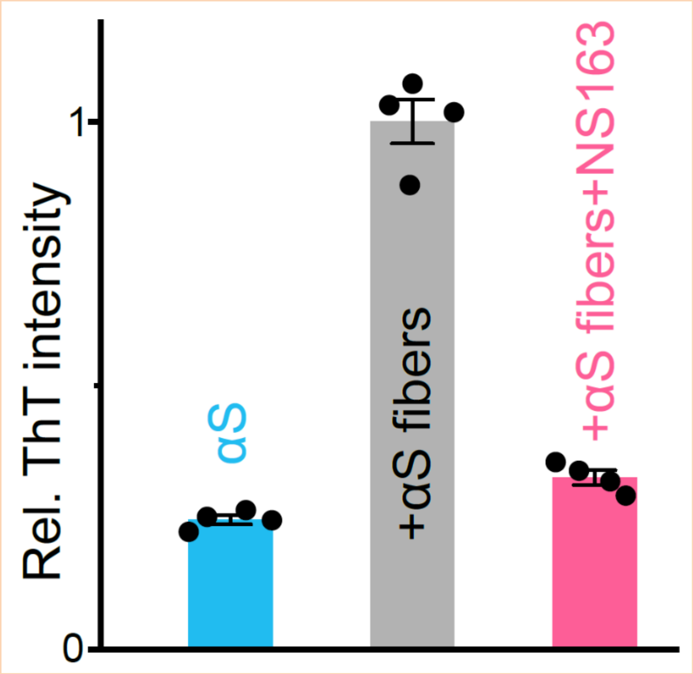


**Supplementary Fig. 11. Effect of NS132 on the seed catalyzed aggregation of αS.** The relative ThT intensity of the aggregation of αS monomer (100 µM), αS monomer (100 µM) + αS fibers (20% monomer), and αS monomer (100 µM) + αS fibers (20% monomer) + NS163 (100 µM) after 20 h in the aggregation buffer conditions.

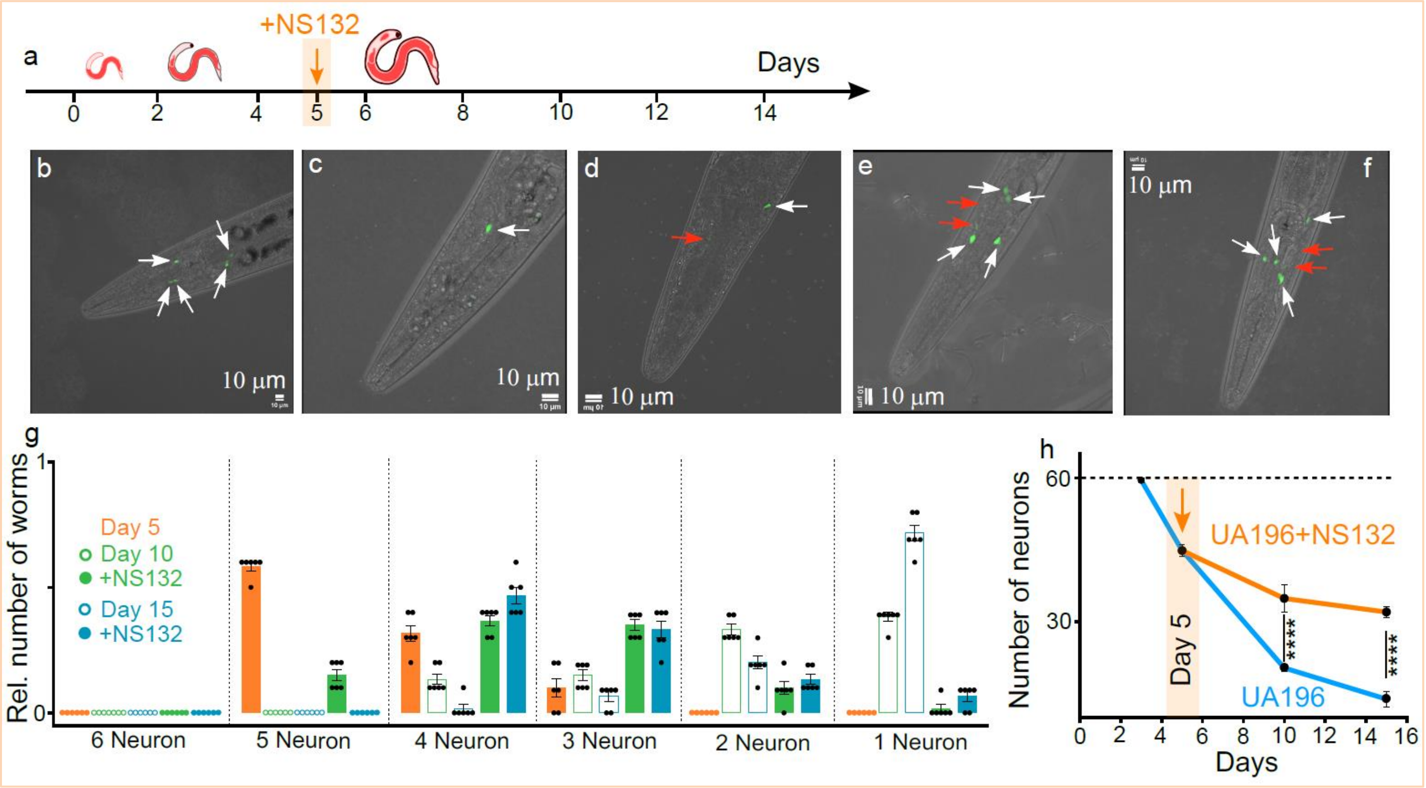


**Supplementary Fig. 12. Neuroprotective effect of NS132 on the preexiting PD *C. elegans* model. a,** Schematic of the aging process of UA196 worms and their treatment with NS-132 at the mid-stage pathology (day 5). Representative confocal images of UA196 worms on day 5 (**b**), day 10 (**c**), and day 15 (**d**). Representative confocal images of UA196 worms on day 10 (**e**) and day 15 (**f**) when treated with 50 μM NS132 on day 5. The healthy (white) and degenerated (red) DA neurons in UA196 worms. **g**, The relative number of healthy DA neurons in UA196 worms during the aging process when treated with 50 μM NS132 on day 5. **h**, Statistics for the total number of neurons in UA196 worms during the aging process in the absence and presence of 50 μM NS132. For each confocal imaging experiment at least 10 worms were used, and the healthy neurons were counted manually, and each condition (day) consisted of six independent experiments. The data were expressed as mean and the error bars report the s.e.m. (n = 6 independent experiments and each n consisted of 10 technical replicates). The statistical analysis was performed using ANOVA with Tukey's multiple comparison test. $^*p < 0.05$, $^{**}p < 0.01$, $^{***}p < 0.001$, $^{****}p < 0.0001$.

**Supplementary Fig. 13. $^1$H-NMR of 6-chloro-5-nitro-2-picoline**

$^1$H NMR (500 MHz, $CDCl_3$) δ 2.63 – 2.66 (s, 3H), 7.26 – 7.29 (d, *J* = 8.0 Hz, 1H), 8.13 – 8.18 (d, *J* = 8.1 Hz, 1H). HRMS (m/z): [M]+ calcd. for $C_6H_5ClN_2O_2$, 173.0112; found, 173.0114.

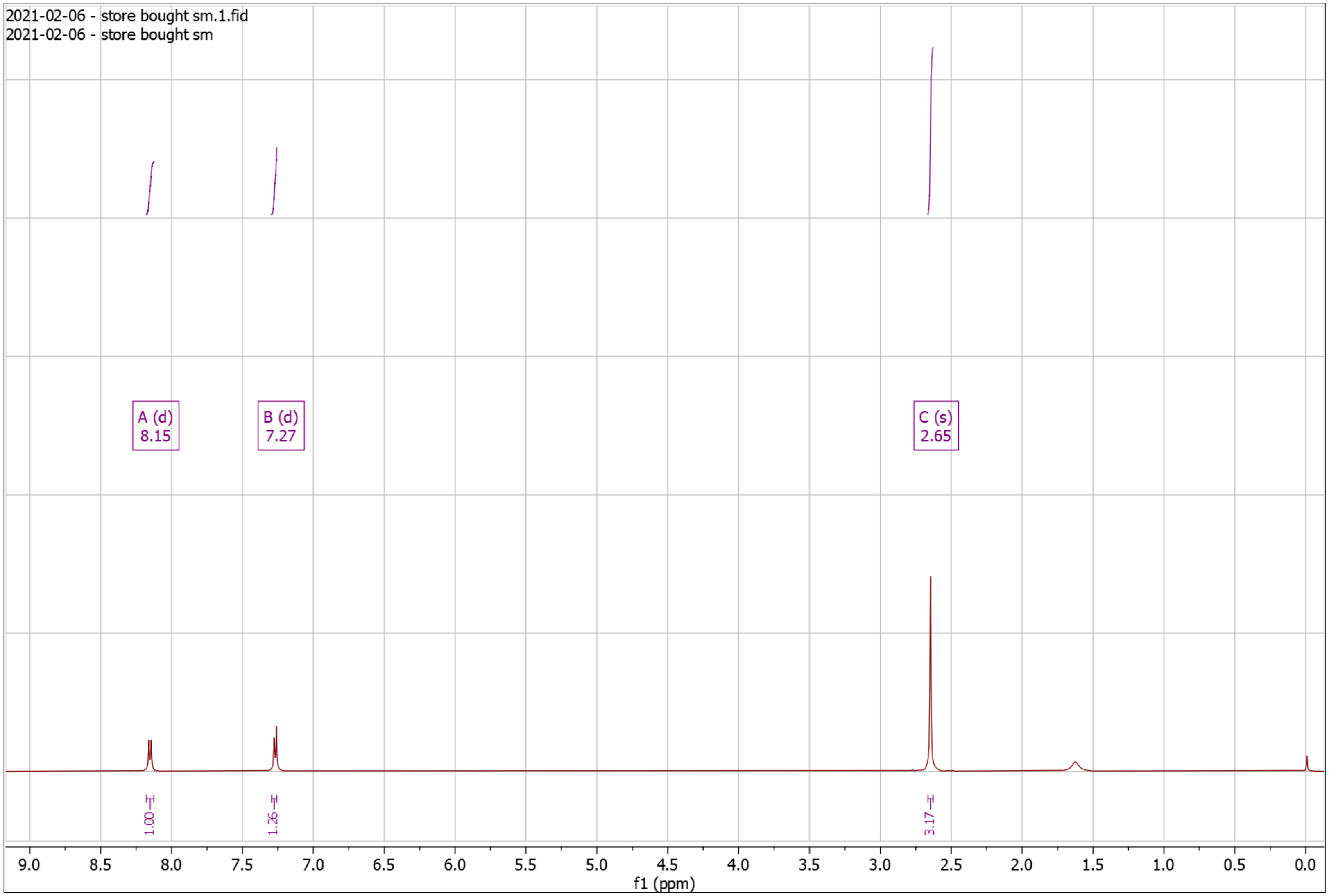


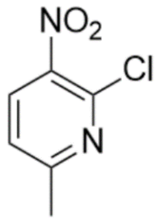

**Supplementary Fig. 14. $^{1}$H-NMR of NS41 Pro**

$^{1}$H NMR (500 MHz, $CDCl_3$) δ 1.42 – 1.45 (s, 9H), 2.46 – 2.48 (s, 3H), 4.88 – 4.92 (s, 2H), 6.87 – 6.91 (d, *J* = 8.1 Hz, 1H), 8.22 – 8.25 (d, *J* = 8.1 Hz, 1H).

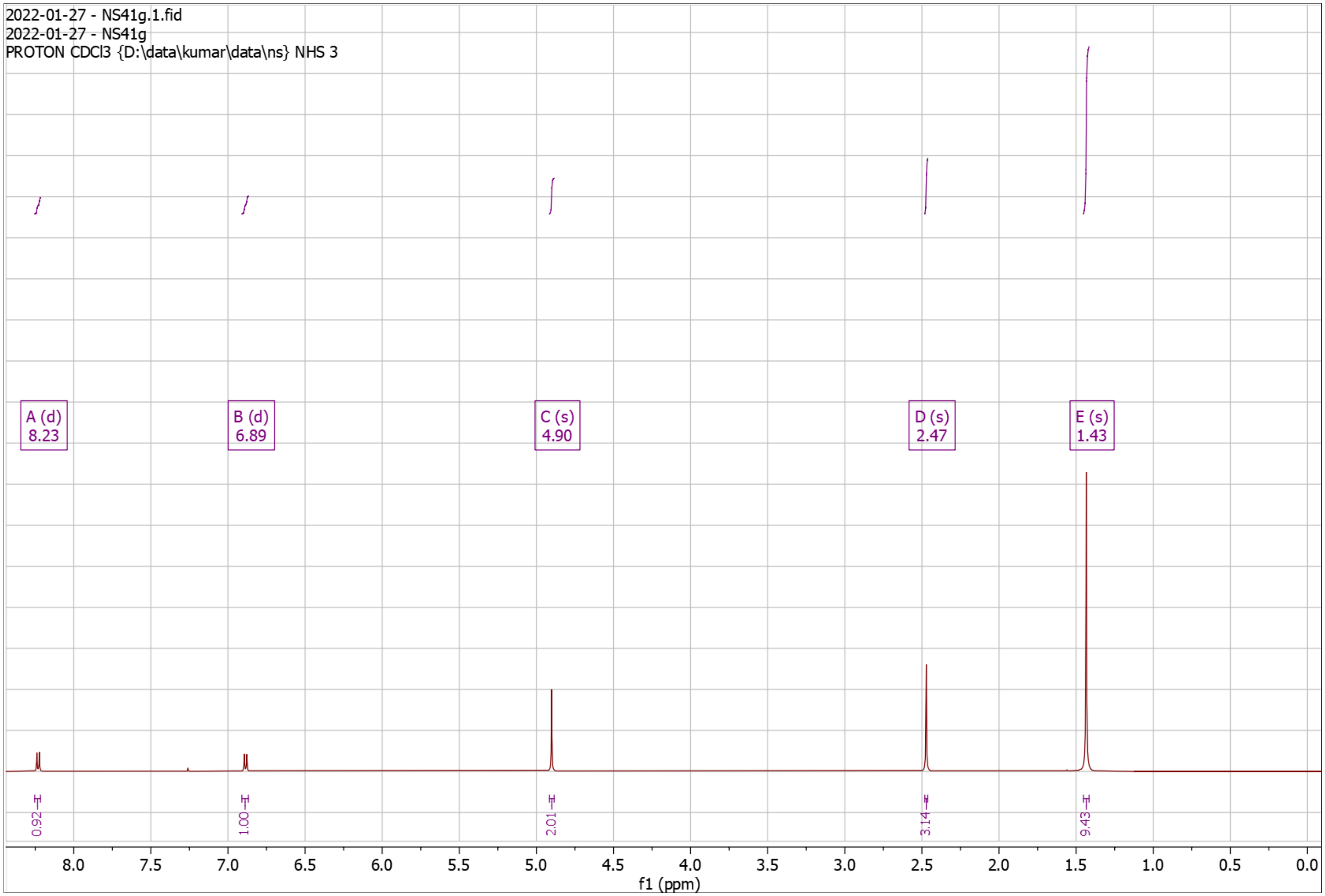


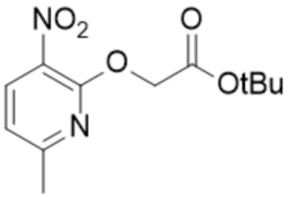

**Supplementary Fig. 15. $^{1}$H-NMR of NS101**

$^{1}$H NMR (500 MHz, DMSO) δ 8.19 – 8.29 (d, *J* = 8.2 Hz, 1H), 8.65 – 8.73 (d, *J* = 8.2 Hz, 1H). HRMS (m/z): [M]+ calcd. for $C_6H_3ClN_2O_4$, 202.9854; found, 202.9859.

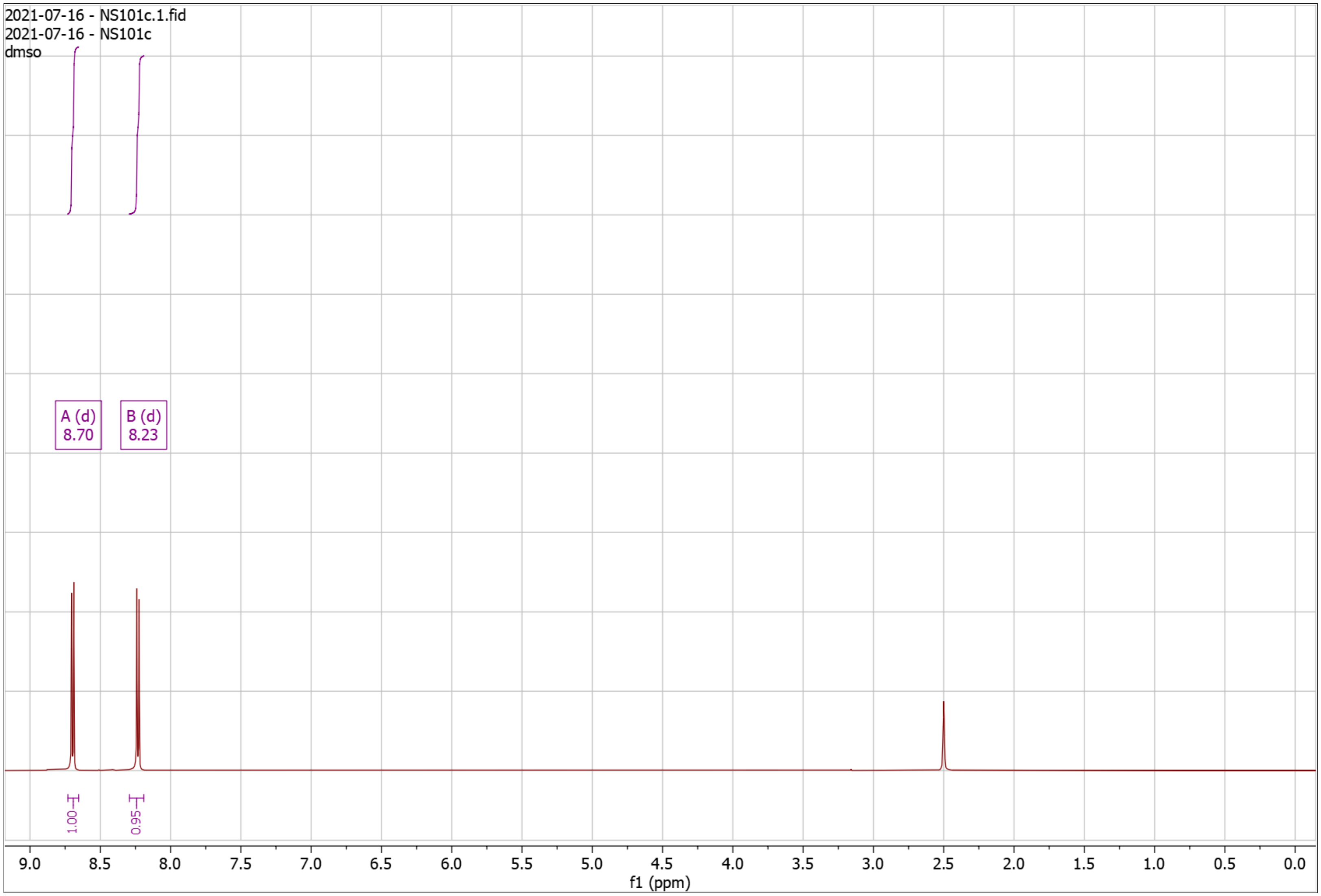


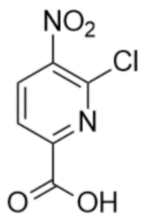

**Supplementary Fig. 16. $^{1}$H-NMR of RD121**

$^{1}$H NMR (500 MHz, $CDCl_3$) δ 8.15 – 8.25 (d, *J* = 8.1 Hz, 1H), 8.33 – 8.38 (d, *J* = 8.2 Hz, 1H). HRMS (m/z): [M]+ calcd. for $C_6H_2Cl_2N_2O_3$, 220.9515; found, 220.9516.

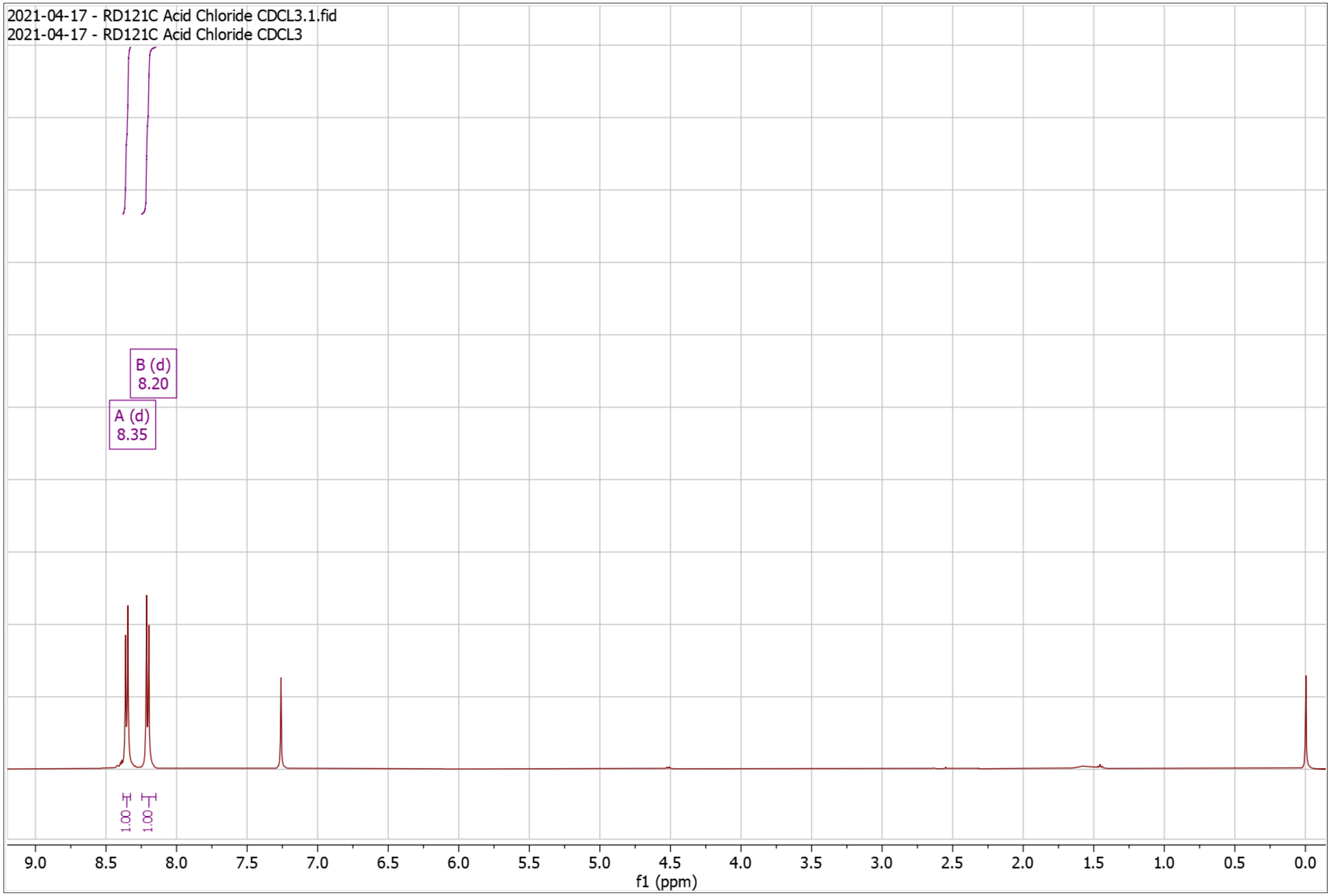


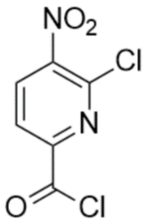

**Supplementary Fig. 17. $^{1}$H-NMR of RD127 Pro**

$^{1}$H NMR (500 MHz, $CDCl_3$) δ 1.47 – 1.50 (s, 9H), 2.38 – 2.42 (s, 3H), 4.87 – 4.91 (s, 2H), 6.80 – 6.86 (d, *J* = 7.9 Hz, 1H), 8.34 – 8.42 (m, 2H), 8.60 – 8.66 (d, *J* = 7.9 Hz, 1H), 10.02 – 10.06 (s, 1H); HRMS (m/z): [M]+ calcd. for $C_{18}H_{19}ClN_4O_6$, 423.1066; found, 423.1055.

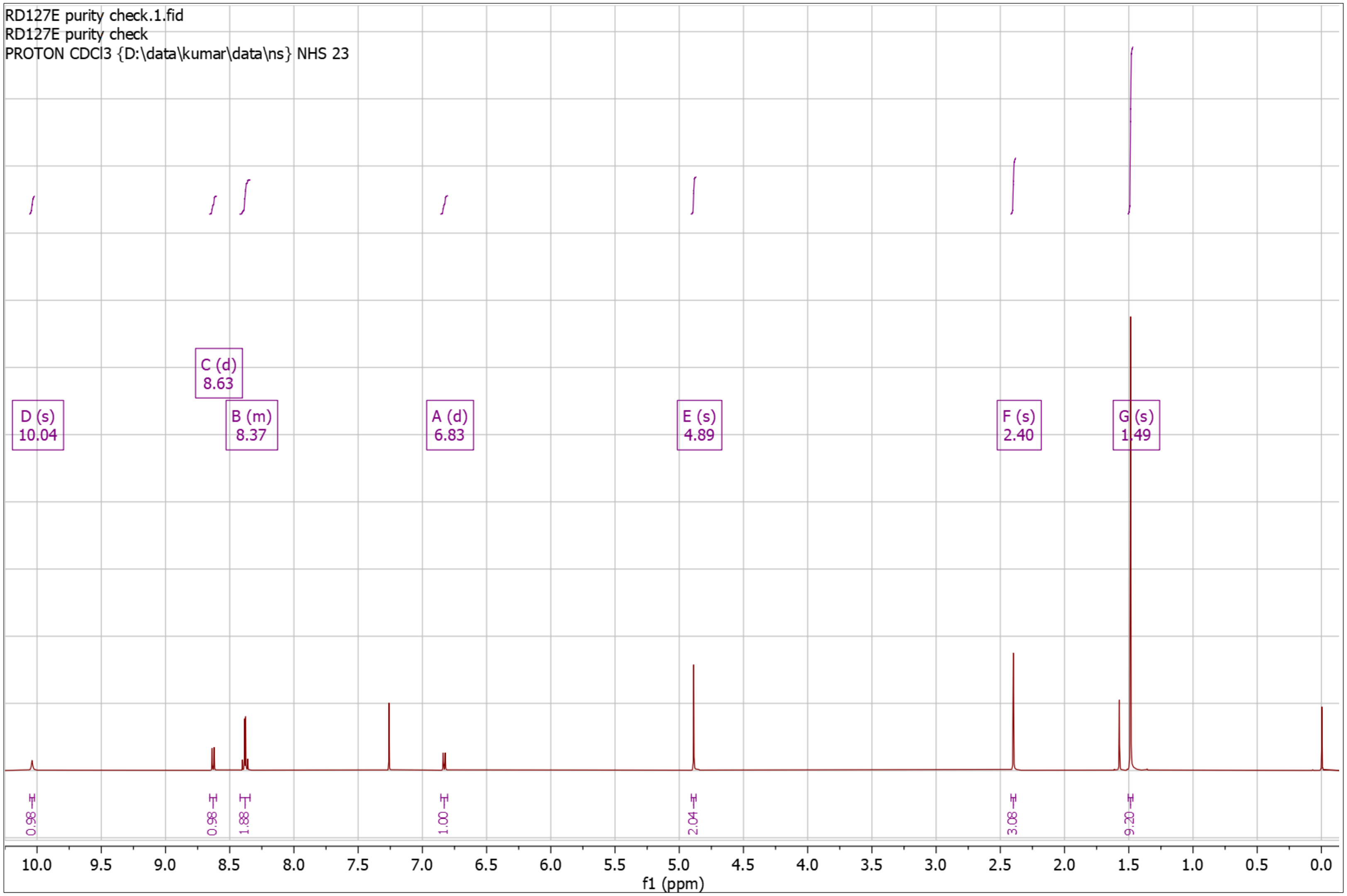


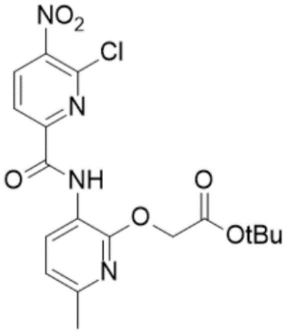

**Supplementary Fig. 18. [1]H-NMR of RD127 Dep**

$^1$H NMR (500 MHz, DMSO) δ 2.31 – 2.40 (s, 3H), 4.92 – 5.00 (s, 2H), 6.93 – 7.00 (d, *J* = 7.9 Hz, 1H), 8.33 – 8.42 (d, *J* = 8.3 Hz, 1H), 8.42 – 8.53 (d, *J* = 7.9 Hz, 1H), 8.74 – 8.86 (d, *J* = 8.3 Hz, 1H), 9.96 – 10.01 (s, 1H), 12.84 – 13.06 (s, 1H); HRMS (m/z): [M]+ calcd. for $C_{14}H_{11}ClN_4O_6$, 367.0440; found, 367.0423.

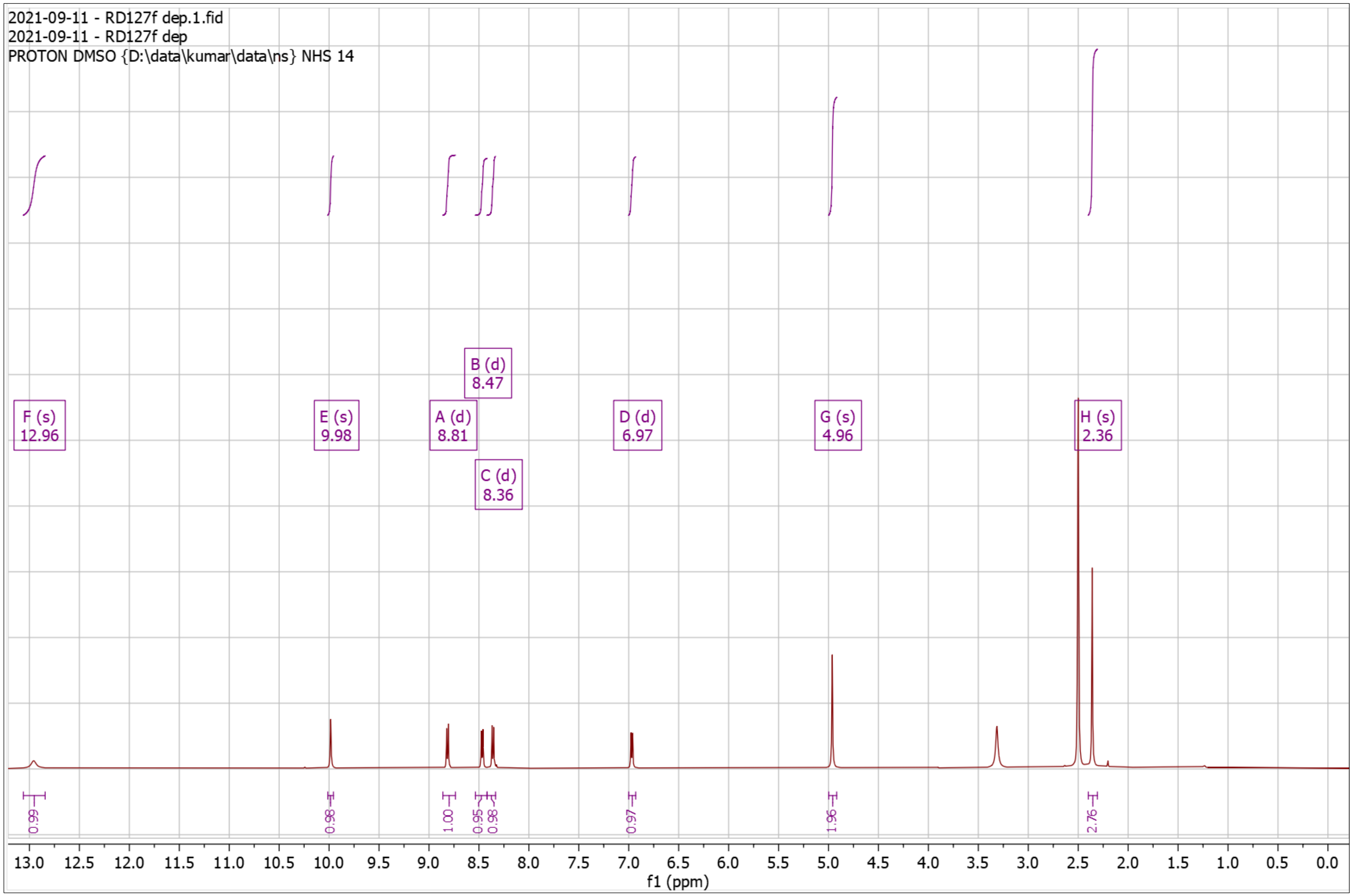


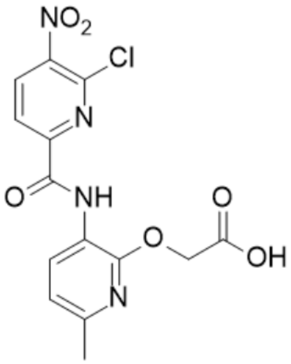

**Supplementary Fig. 19. $^{1}$H-NMR of NS48 Pro**

$^{1}$H NMR (500 MHz, $CDCl_3$) δ 0.96 – 1.03 (d, $J$ = 6.6 Hz, 6H), 1.42 – 1.48 (s, 9H), 1.63 – 1.69 (q, $J$ = 7.0 Hz, 2H), 1.71 – 1.83 (dh, $J$ = 13.3, 6.6 Hz, 1H), 2.36 – 2.44 (s, 3H), 3.72 – 3.83 (td, $J$ = 7.3, 5.4 Hz, 2H), 4.80 – 4.88 (s, 2H), 6.77 – 6.88 (d, $J$ = 8.0 Hz, 1H), 7.51 – 7.59 (d, $J$ = 8.5 Hz, 1H), 8.15 – 8.27 (t, $J$ = 5.4 Hz, 1H), 8.56 – 8.63 (d, $J$ = 8.4 Hz, 1H), 8.65 – 8.71 (d, $J$ = 7.9 Hz, 1H), 10.16 – 10.28 (s, 1H); HRMS (m/z): [M]+ calcd. for $C_{23}H_{31}N_5O_6$, 474.2347; found, 474.2349.

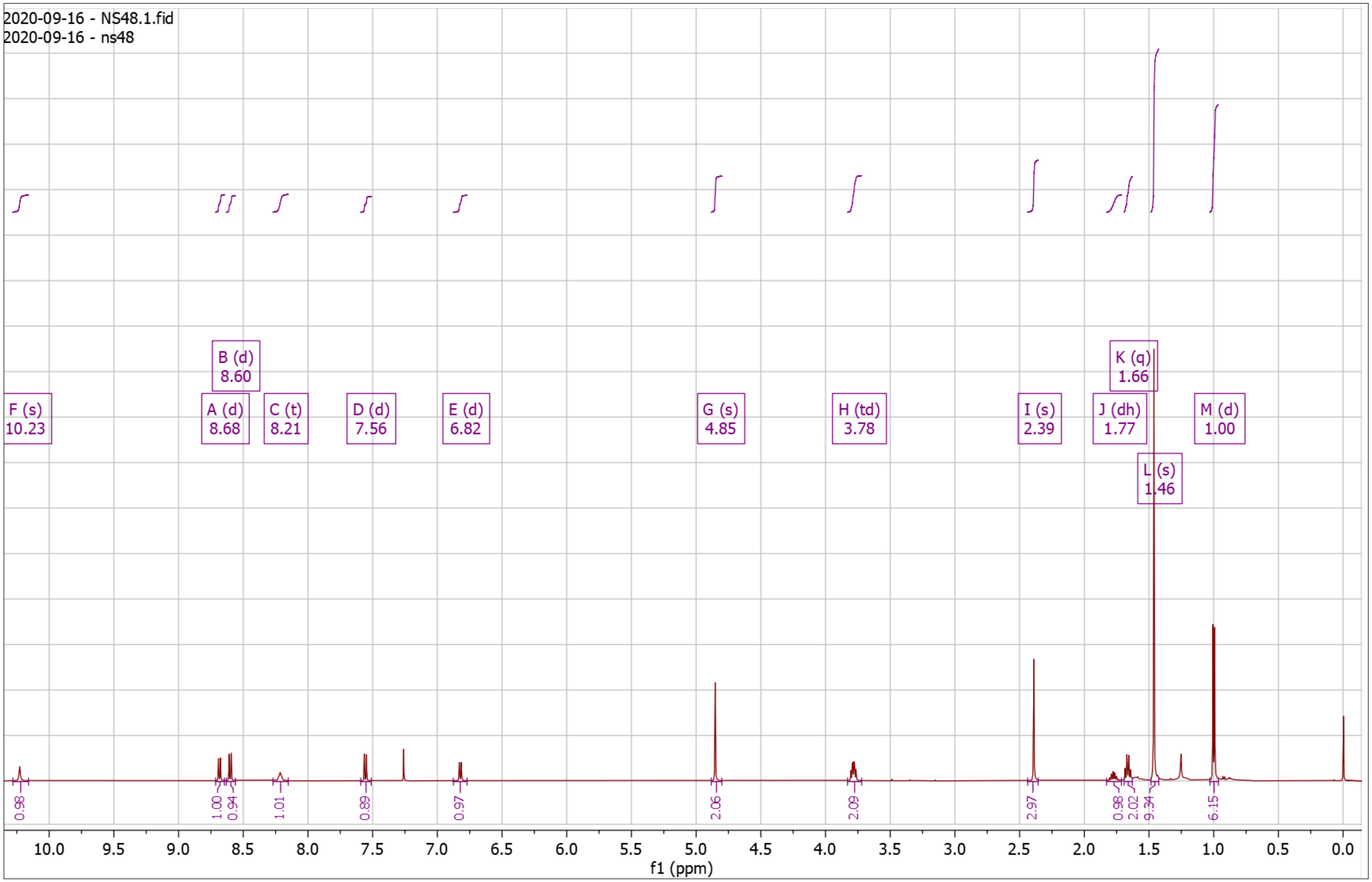


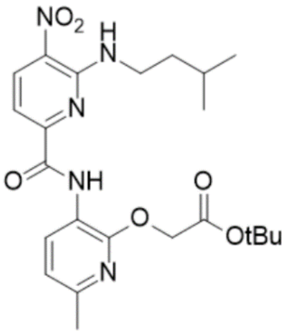

**Supplementary Fig. 20. $^{1}$H-NMR of NS48 Dep**

$^{1}$H NMR (500 MHz, DMSO) δ 0.90 – 0.93 (d, *J* = 6.5 Hz, 6H), 1.54 – 1.60 (q, *J* = 7.1 Hz, 2H), 1.62 – 1.74 (hept, *J* = 6.6 Hz, 1H), 2.31 – 2.35 (s, 3H), 3.56 – 3.73 (q, *J* = 6.6 Hz, 2H), 4.91 – 4.95 (s, 2H), 6.87 – 6.97 (d, *J* = 8.0 Hz, 1H), 7.33 – 7.44 (d, *J* = 8.4 Hz, 1H), 8.47 – 8.52 (t, *J* = 5.7 Hz, 1H), 8.52 – 8.56 (d, *J* = 7.9 Hz, 1H), 8.56 – 8.64 (d, *J* = 8.4 Hz, 1H), 10.09 – 10.13 (s, 1H), 12.66 – 13.23 (s, 1H); HRMS (m/z): [M]+ calcd. for $C_{19}H_{23}N_5O_6$, 418.1721; found, 418.1704.

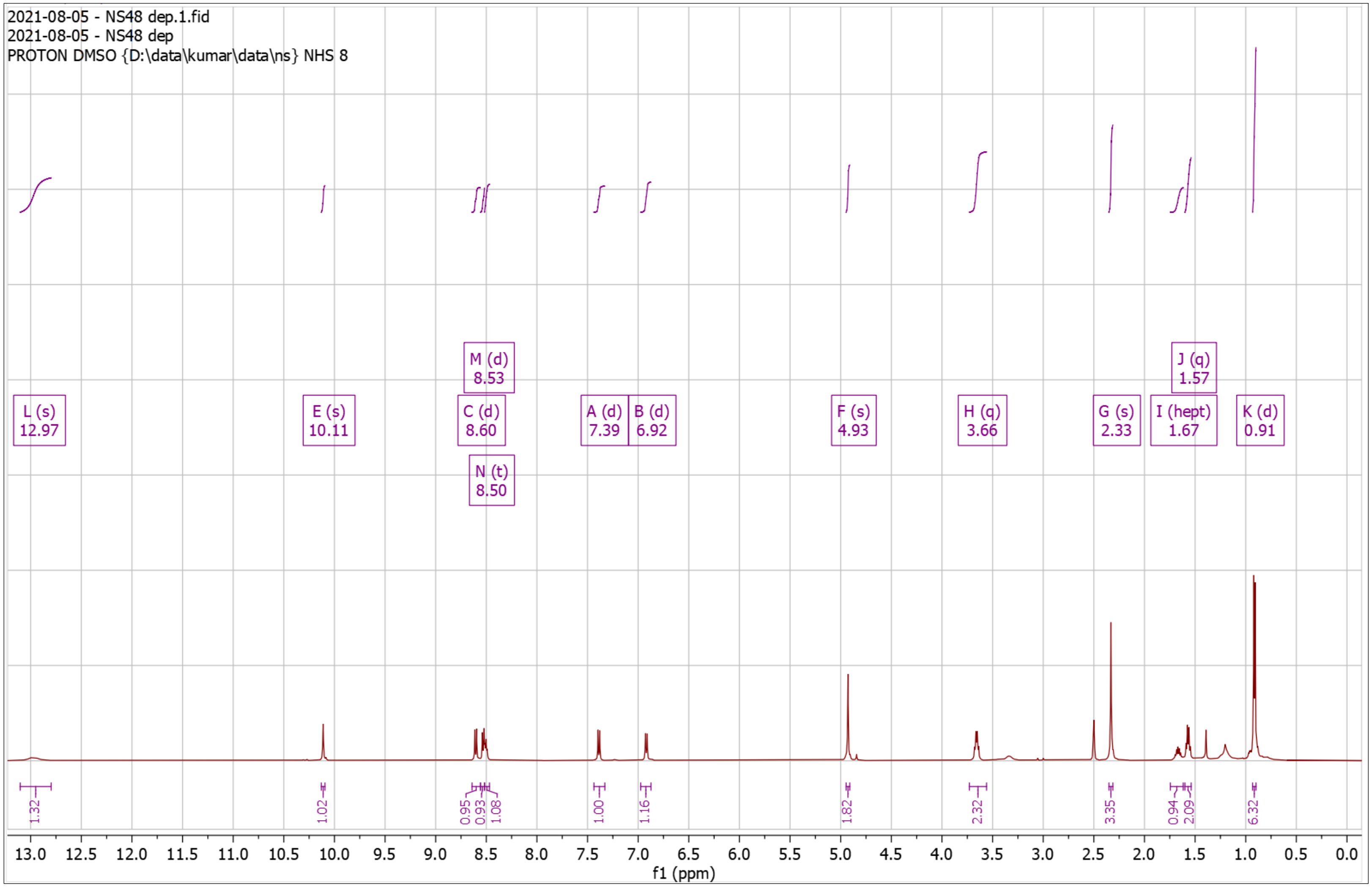


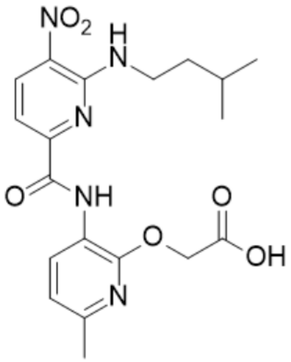

**Supplementary Fig. 21. $^1$H-NMR of NS50 Pro**

$^1$H NMR (500 MHz, $CDCl_3$) δ 1.47 – 1.48 (s, 9H), 2.06 – 2.11 (t, *J* = 2.7 Hz, 1H), 2.38 – 2.41 (s, 3H), 2.65 – 2.72 (td, *J* = 6.8, 2.7 Hz, 2H), 3.93 – 4.00 (q, *J* = 6.6 Hz, 2H), 4.90 – 4.94 (s, 2H), 6.80 – 6.85 (d, *J* = 7.9 Hz, 1H), 7.58 – 7.63 (d, *J* = 8.4 Hz, 1H), 8.45 – 8.51 (t, *J* = 6.0 Hz, 1H), 8.60 – 8.65 (d, *J* = 8.4 Hz, 1H), 8.65 – 8.70 (d, *J* = 7.9 Hz, 1H), 10.22 – 10.26 (s, 1H); HRMS (m/z): [M]+ calcd. for $C_{22}H_{25}N_5O_6$, 456.1878; found, 456.1867.

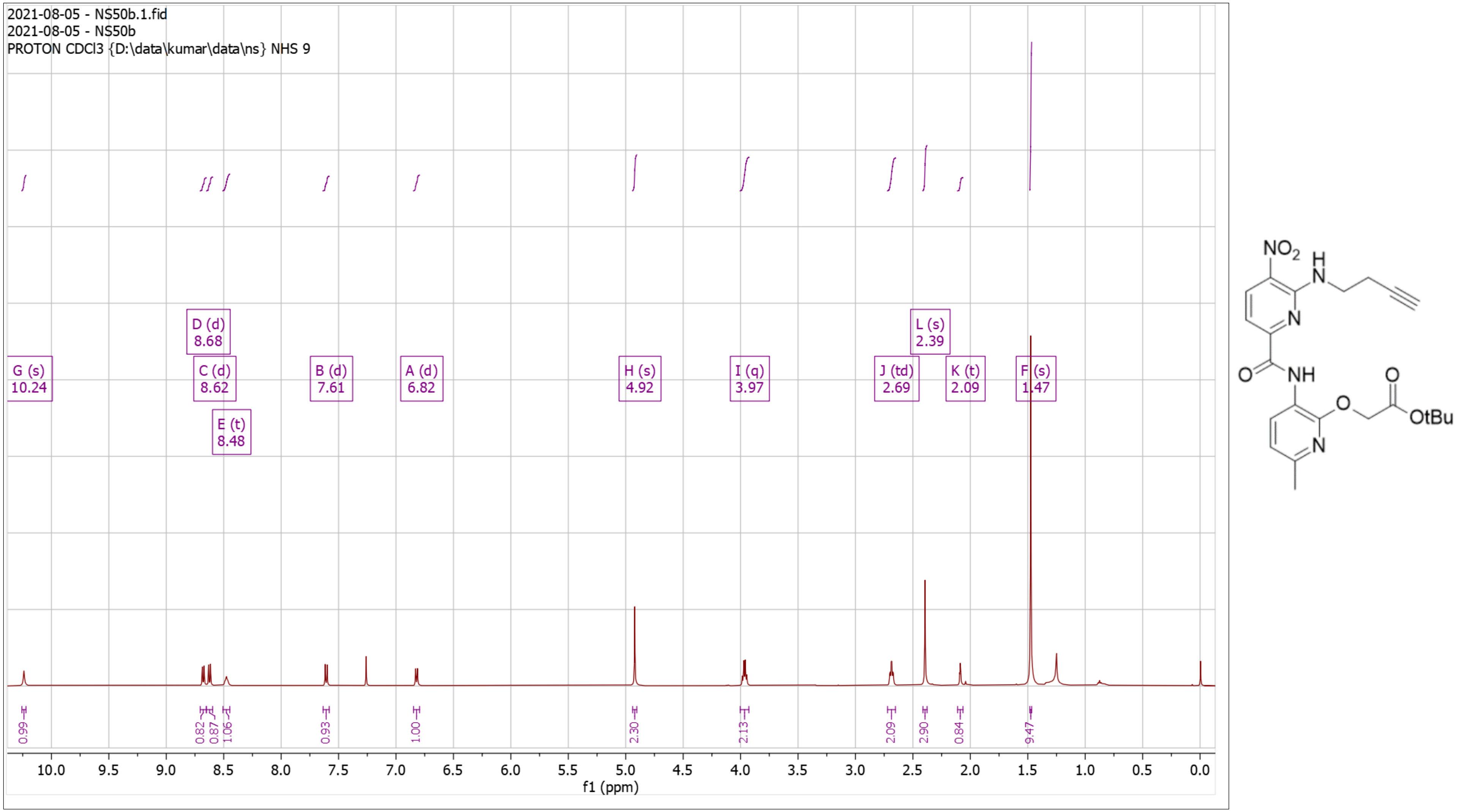

**Supplementary Fig. 22. [1]H-NMR of NS50 Dep**

[1]H NMR (500 MHz, DMSO) δ 2.32 – 2.36 (s, 3H), 2.55 – 2.67 (td, *J* = 7.2, 2.7 Hz, 2H), 2.85 – 2.90 (d, *J* = 2.7 Hz, 1H), 3.75 – 3.87 (q, *J* = 6.8 Hz, 2H), 4.95 – 5.03 (s, 2H), 6.89 – 6.98 (d, *J* = 7.8 Hz, 1H), 7.40 – 7.50 (d, *J* = 8.1 Hz, 1H), 8.49 – 8.57 (d, *J* = 7.8 Hz, 1H), 8.59 – 8.68 (d, *J* = 8.1 Hz, 1H), 8.68 – 8.78 (t, *J* = 6.0 Hz, 1H), 10.16 – 10.23 (s, 1H); HRMS (m/z): [M]+ calcd. for $C_{18}H_{17}N_5O_6$, 400.1252; found, 400.1251.

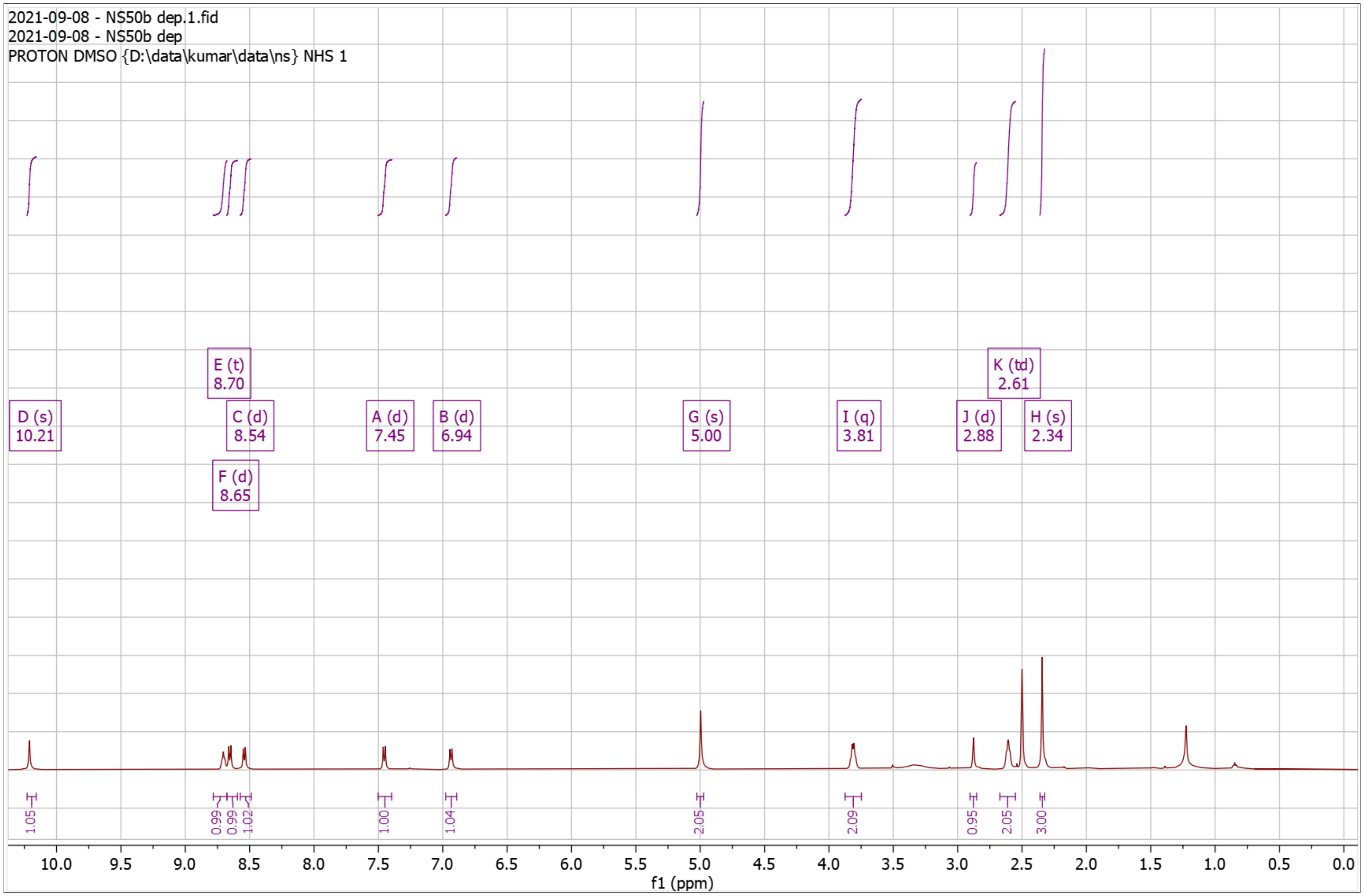


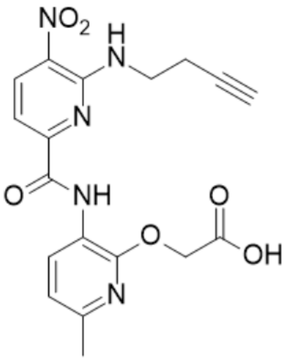

**Supplementary Fig. 23. $^{1}$H-NMR of NS52 Pro**

$^{1}$H NMR (500 MHz, $CDCl_3$) δ 1.03 – 1.12 (t, *J* = 7.4 Hz, 3H), 1.44 – 1.49 (s, 9H), 1.73 – 1.87 (h, *J* = 7.3 Hz, 2H), 2.37 – 2.41 (s, 3H), 3.68 – 3.79 (td, *J* = 7.0, 5.6 Hz, 2H), 4.84 – 4.93 (s, 2H), 6.78 – 6.89 (d, *J* = 7.9 Hz, 1H), 7.50 – 7.59 (d, *J* = 8.4 Hz, 1H), 8.23 – 8.32 (t, 1H), 8.57 – 8.64 (d, *J* = 8.4 Hz, 1H), 8.65 – 8.72 (d, *J* = 7.9 Hz, 1H), 10.18 – 10.36 (s, 1H); HRMS (m/z): [M]+ calcd. for $C_{21}H_{27}N_5O_6$, 446.2034; found, 446.2037.

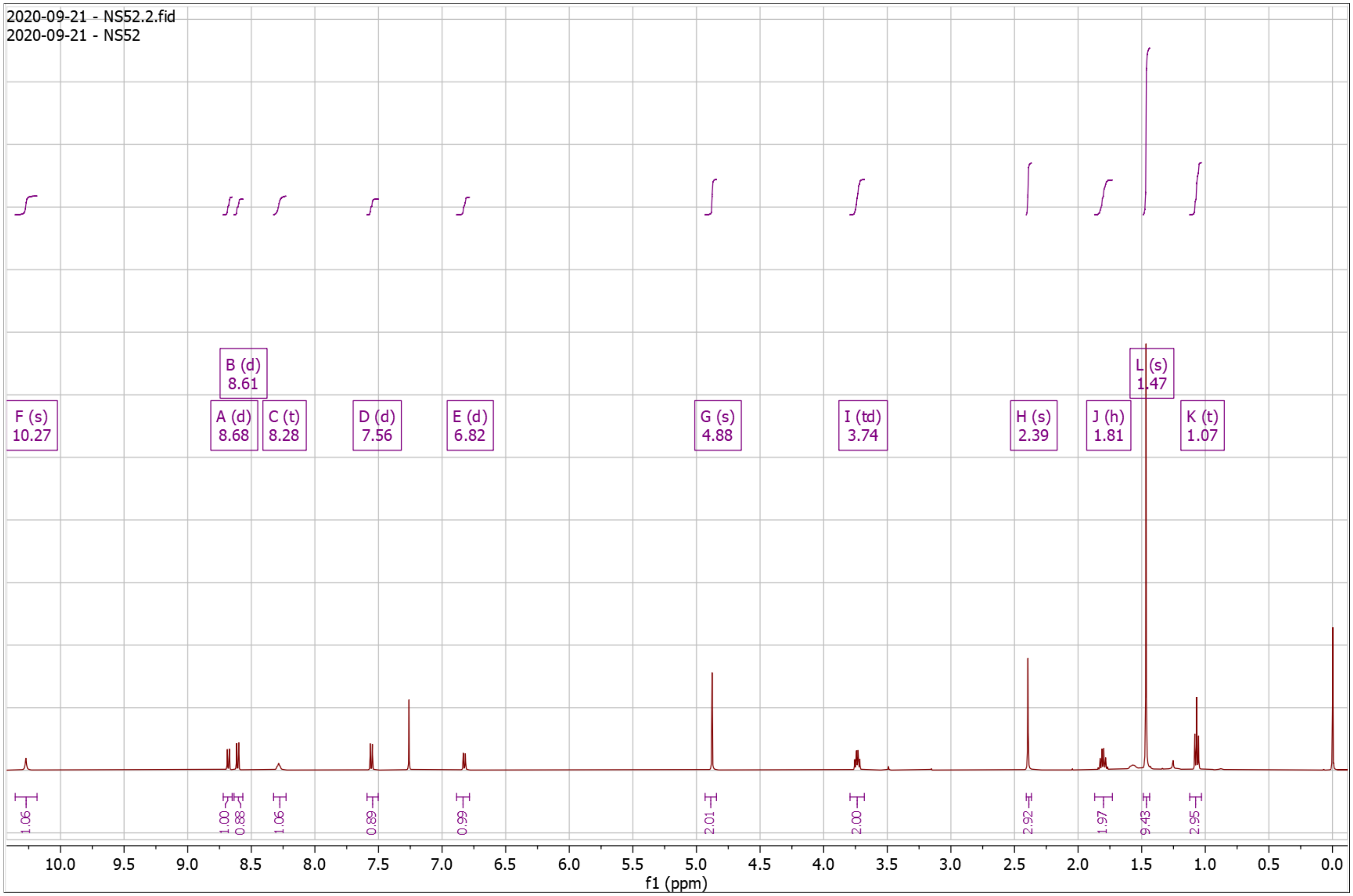


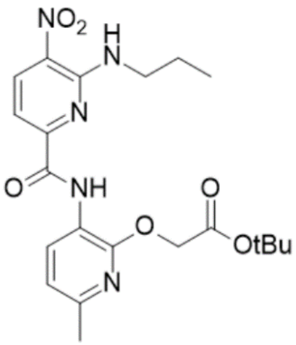

**Supplementary Fig. 24. $^1$H-NMR of NS52 Dep**

$^1$H NMR (500 MHz, DMSO) δ 0.89 – 0.98 (t, *J* = 7.4 Hz, 3H), 1.63 – 1.77 (h, *J* = 7.4 Hz, 2H), 2.28 – 2.40 (s, 3H), 3.55 – 3.68 (q, *J* = 6.6 Hz, 2H), 4.89 – 5.05 (s, 2H), 6.90 – 7.04 (d, *J* = 7.9 Hz, 1H), 7.35 – 7.50 (d, *J* = 8.2 Hz, 1H), 8.52 – 8.59 (d, *J* = 7.9 Hz, 1H), 8.59 – 8.64 (t, 1H), 8.64 – 8.69 (d, *J* = 8.3 Hz, 1H), 10.12 – 10.29 (s, 1H), 12.74 – 13.08 (s, 1H); HRMS (m/z): [M]+ calcd. for $C_{17}H_{19}N_5O_6$, 390.1408; found, 390.1403.

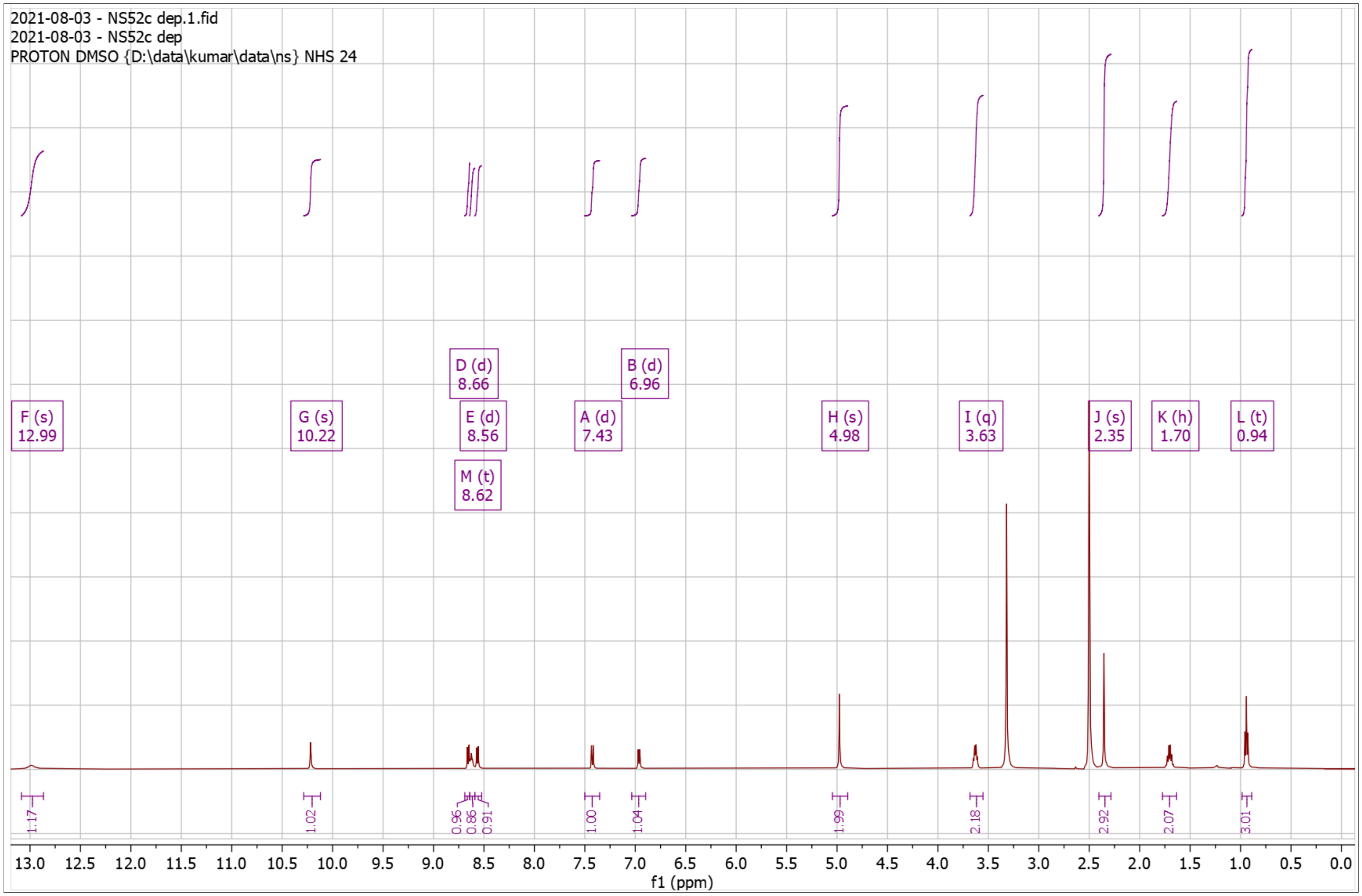


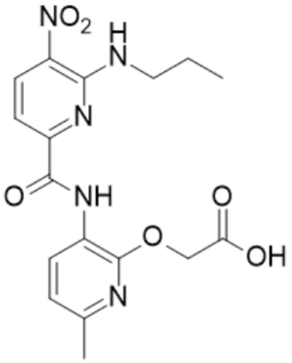

**Supplementary Fig. 25. [1]H-NMR of NS53 Pro**

$^{1}$H NMR (500 MHz, $CDCl_3$) δ 1.46 – 1.50 (s, 9H), 2.38 – 2.42 (s, 3H), 4.81 – 4.85 (s, 2H), 4.93 – 4.98 (d, *J* = 5.5 Hz, 2H), 6.30 – 6.34 (d, *J* = 2.2 Hz, 1H), 6.80 – 6.85 (d, *J* = 7.9 Hz, 1H), 7.47 – 7.51 (d, *J* = 2.2 Hz, 1H), 7.56 – 7.61 (d, *J* = 8.4 Hz, 1H), 8.47 – 8.52 (d, *J* = 7.9 Hz, 1H), 8.57 – 8.62 (d, *J* = 8.5 Hz, 1H), 8.62 – 8.67 (t, *J* = 5.4 Hz, 1H), 10.31 – 10.35 (s, 1H); HRMS (m/z): [M]+ calcd. for $C_{22}H_{25}N_7O_6$, 484.1939; found, 484.1934.

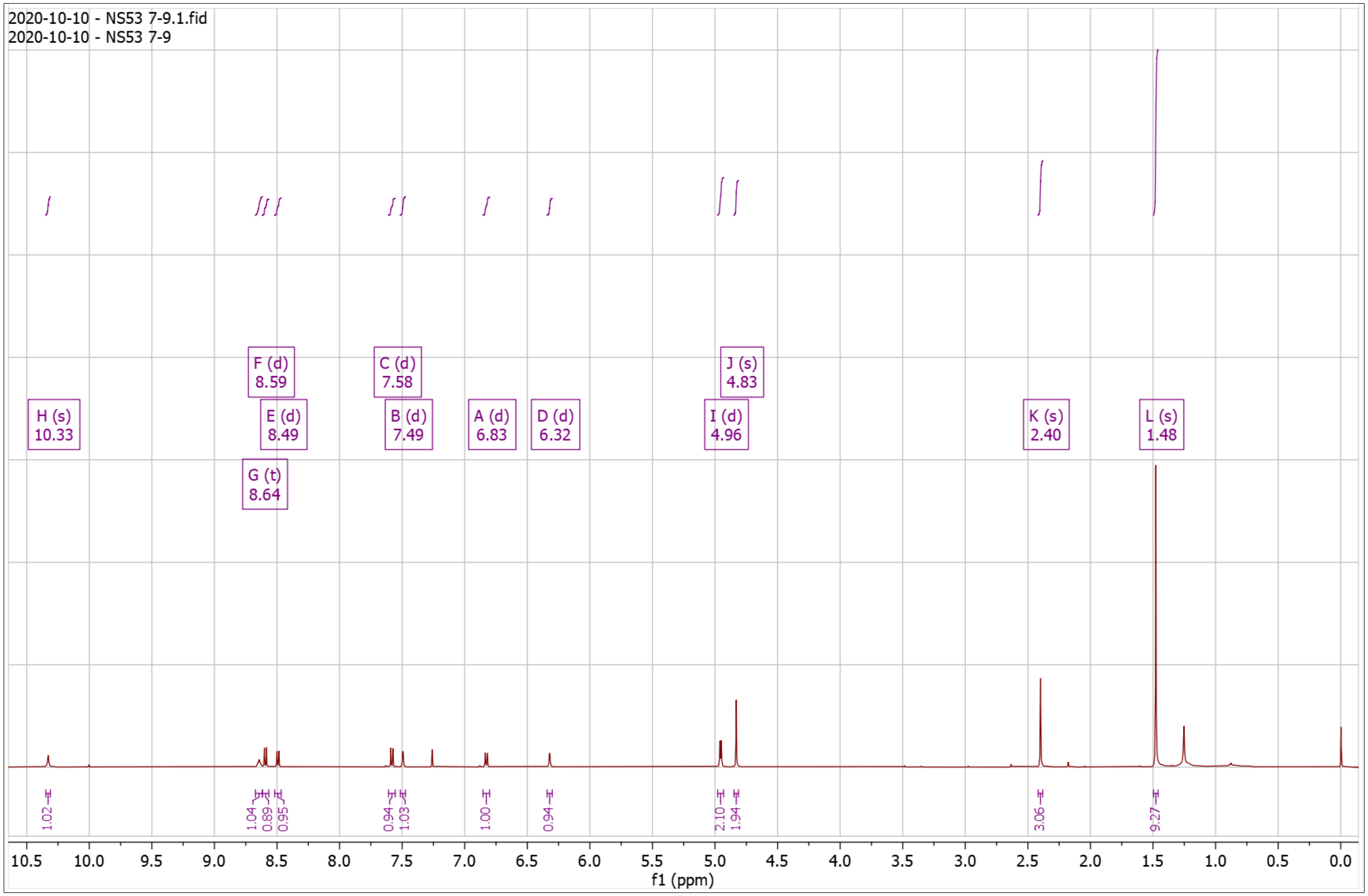


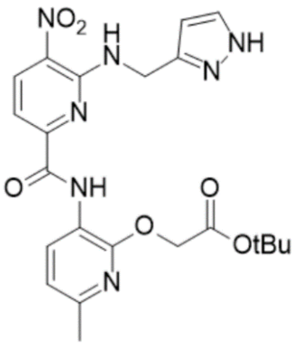

**Supplementary Fig. 26. $^{1}$H-NMR of NS53 Dep**

$^{1}$H NMR (500 MHz, DMSO) δ 2.34 – 2.37 (s, 3H), 4.87 – 4.90 (s, 3H), 6.22 – 6.31 (s, 1H), 6.89 – 7.00 (d, $J$ = 7.9 Hz, 1H), 7.42 – 7.51 (d, $J$ = 8.4 Hz, 1H), 7.54 – 7.63 (s, 1H), 8.41 – 8.54 (d, $J$ = 7.9 Hz, 1H), 8.64 – 8.73 (d, $J$ = 8.4 Hz, 1H), 8.88 – 9.06 (t, $J$ = 5.8 Hz, 1H), 10.15 – 10.39 (s, 1H); HRMS (m/z): [M]+ calcd. for $C_{18}H_{17}N_7O_6$, 428.1313; found, 428.1313.

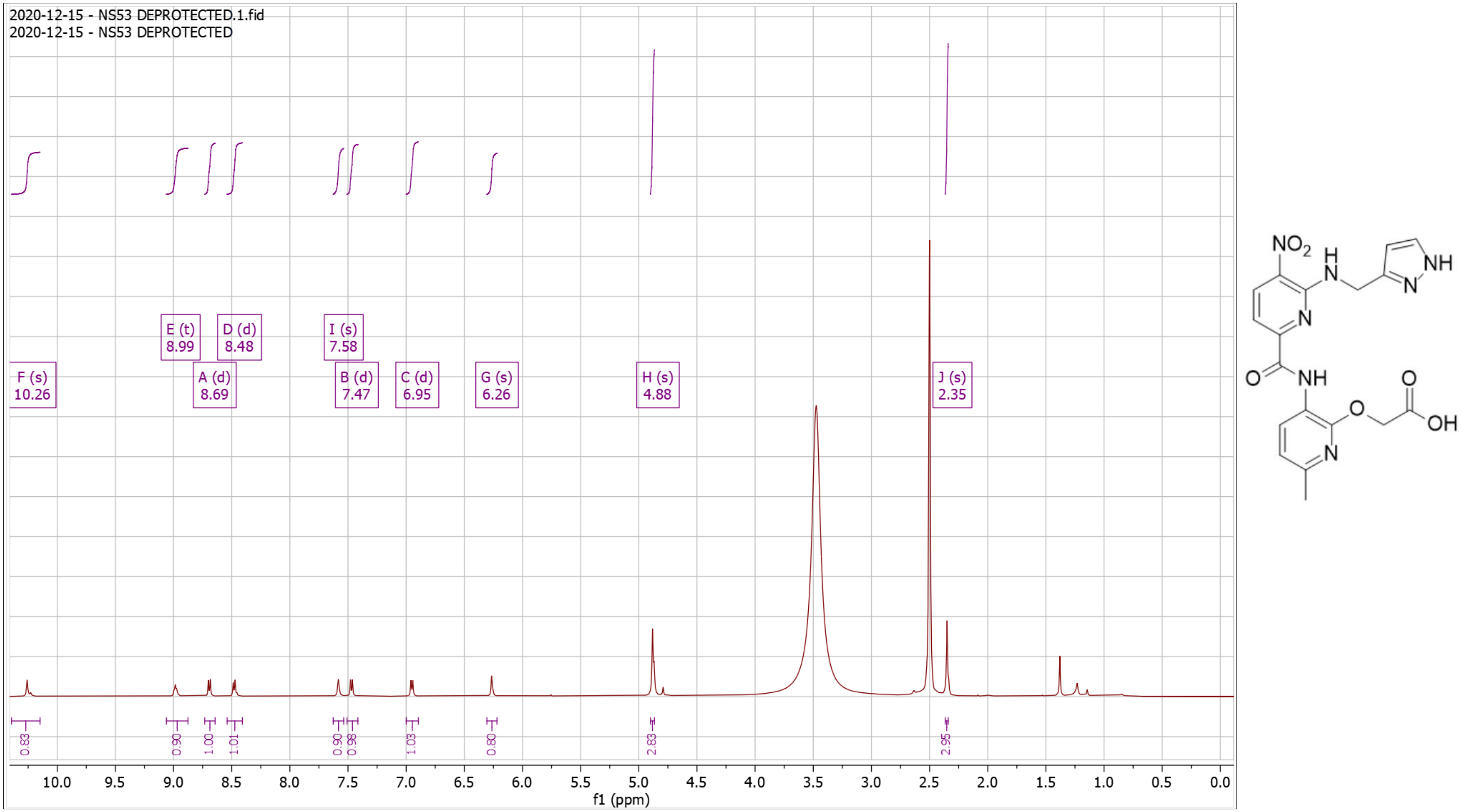

**Supplementary Fig. 27. $^{1}$H-NMR of NS54 Pro**

$^{1}$H NMR (500 MHz, $CDCl_3$) δ 0.35 – 0.40 (m, 2H), 0.60 – 0.66 (m, 2H), 1.26 – 1.32 (tdd, *J* = 8.0, 5.2, 1.9 Hz, 1H), 1.45 – 1.49 (s, 9H), 2.37 – 2.42 (s, 3H), 3.58 – 3.65 (dd, *J* = 7.2, 5.2 Hz, 2H), 4.87 – 4.91 (s, 2H), 6.80 – 6.85 (d, *J* = 7.9 Hz, 1H), 7.54 – 7.60 (d, *J* = 8.4 Hz, 1H), 8.29 – 8.38 (t, 1H), 8.58 – 8.64 (d, *J* = 8.4 Hz, 1H), 8.64 – 8.70 (d, *J* = 7.9 Hz, 1H), 10.25 – 10.34 (s, 1H); HRMS (m/z): [M]+ calcd. for $C_{22}H_{27}N_5O_6$, 458.2034; found, 458.2034.

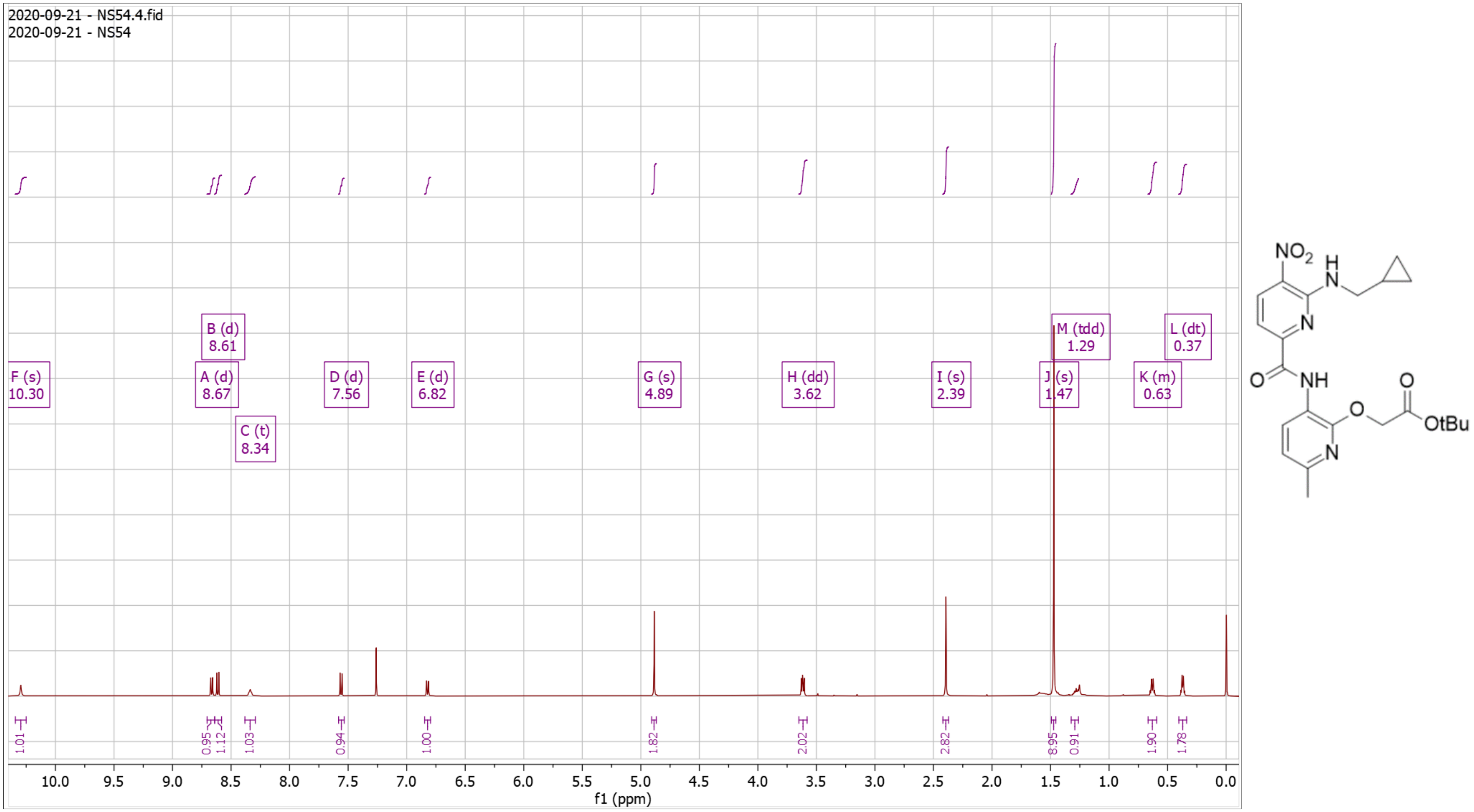

**Supplementary Fig. 28. $^{1}$H-NMR of NS54 Dep**

$^{1}$H NMR (500 MHz, DMSO) δ 0.32 – 0.38 (q, *J* = 4.8 Hz, 2H), 0.42 – 0.50 (d, *J* = 7.6 Hz, 2H), 1.26 – 1.34 (m, 1H), 2.33 – 2.38 (s, 3H), 3.48 – 3.56 (t, *J* = 6.1 Hz, 2H), 4.96 – 5.01 (s, 2H), 6.91 – 7.01 (d, *J* = 7.9 Hz, 1H), 7.38 – 7.49 (d, *J* = 8.3 Hz, 1H), 8.50 – 8.59 (d, *J* = 7.8 Hz, 1H), 8.63 – 8.69 (d, *J* = 8.4 Hz, 1H), 8.69 – 8.77 (t, *J* = 6.0 Hz, 1H), 10.29 – 10.34 (s, 1H); HRMS (m/z): [M]+ calcd. for $C_{18}H_{19}N_5O_6$, 402.1408; found, 402.1402.

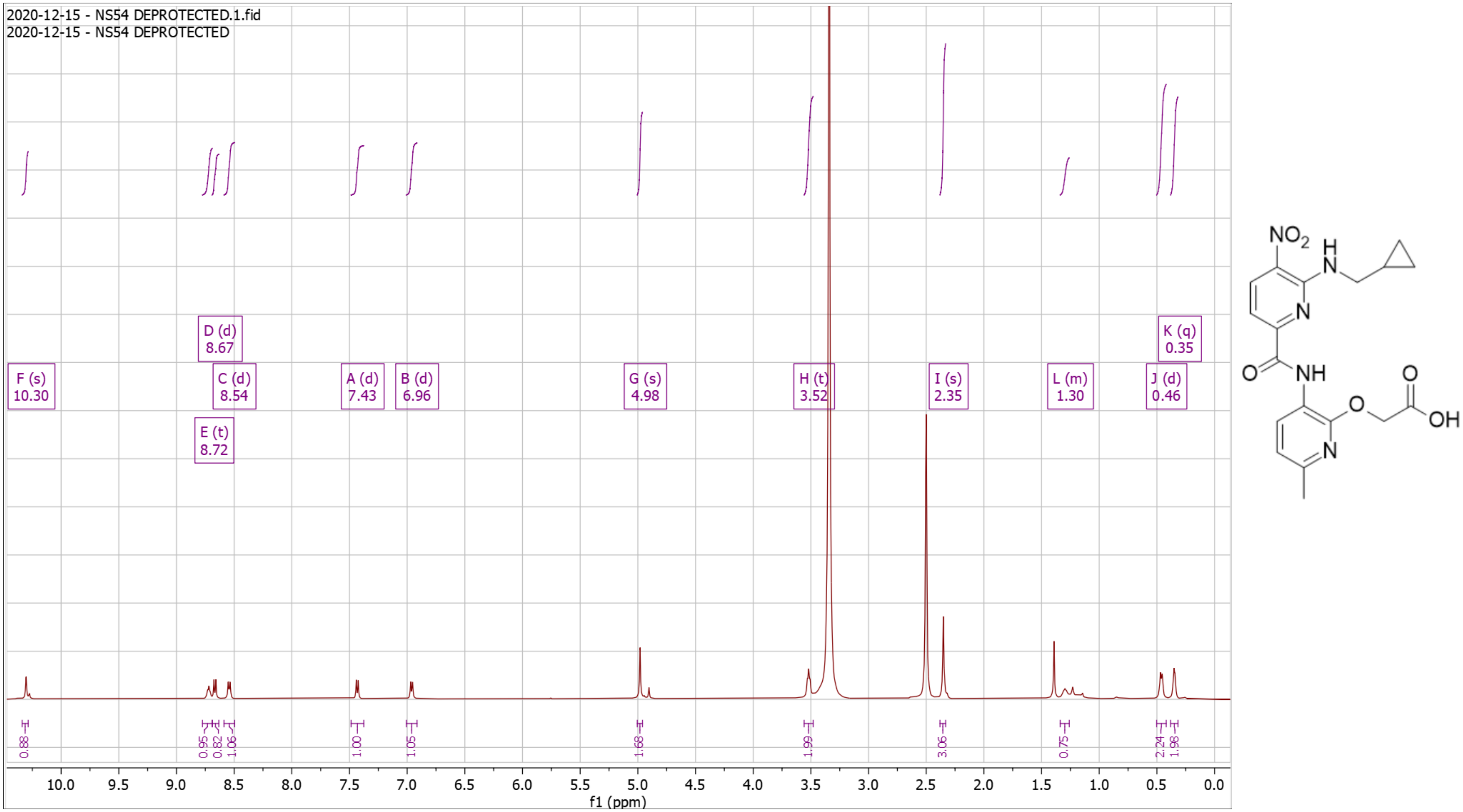

**Supplementary Fig. 29. $^{1}$H-NMR of NS55 Pro**

$^{1}$H NMR (500 MHz, $CDCl_3$) δ 1.01 – 1.34 (m, 5H), 1.46 – 1.46 (s, 9H), 1.66 – 1.91 (m, 6H), 2.37 – 2.41 (s, 3H), 3.59 – 3.65 (t, *J* = 6.3 Hz, 2H), 4.84 – 4.89 (s, 2H), 6.79 – 6.85 (d, *J* = 7.9 Hz, 1H), 7.50 – 7.57 (d, *J* = 8.4 Hz, 1H), 8.31 – 8.37 (t, *J* = 5.8 Hz, 1H), 8.57 – 8.62 (d, *J* = 8.4 Hz, 1H), 8.66 – 8.71 (d, *J* = 8.0 Hz, 1H), 10.21 – 10.24 (s, 1H); HRMS (m/z): [M]+ calcd. for $C_{25}H_{33}N_5O_6$, 500.2504; found, 500.2500.

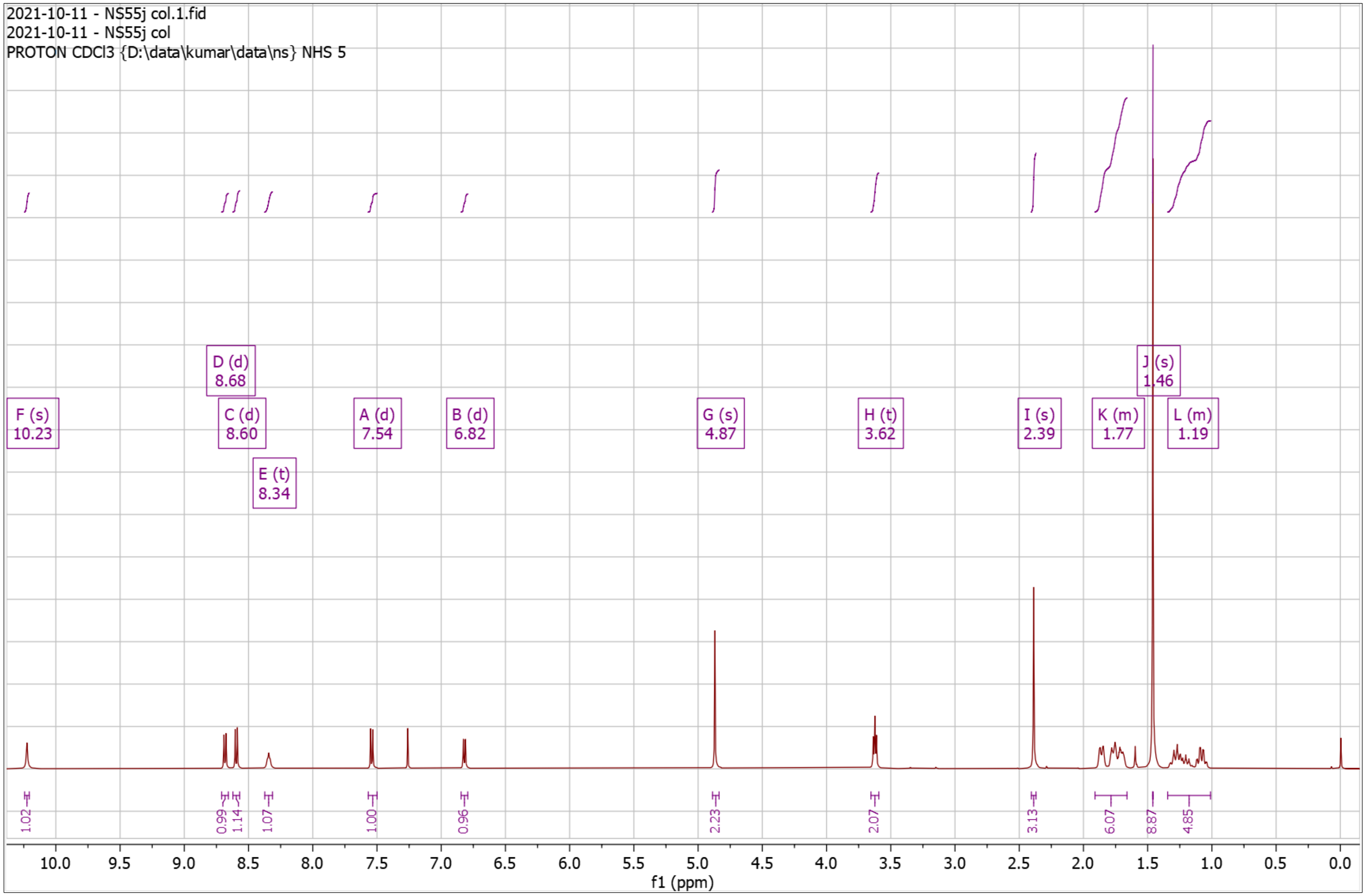


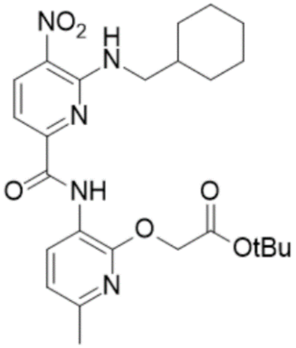

**Supplementary Fig. 30. $^{1}$H-NMR of NS55 Dep**

$^{1}$H NMR (500 MHz, DMSO) δ 10.19 – 10.10 (s, 1H), 8.67 – 8.62 (d, *J* = 8.4 Hz, 1H), 8.60 – 8.50 (m, 2H), 7.48 – 7.39 (d, *J* = 8.4 Hz, 1H), 6.99 – 6.93 (d, *J* = 8.0 Hz, 1H), 4.99 – 4.95 (s, 2H), 3.59 – 3.53 (t, *J* = 6.4 Hz, 2H), 2.37 – 2.33 (s, 3H), 1.81 – 1.63 (m, 6H), 1.24 – 0.94 (m, 5H); HRMS (m/z): [M]+ calcd. for $C_{21}H_{25}N_5O_6$ , 444.1878; found, 444.1878.

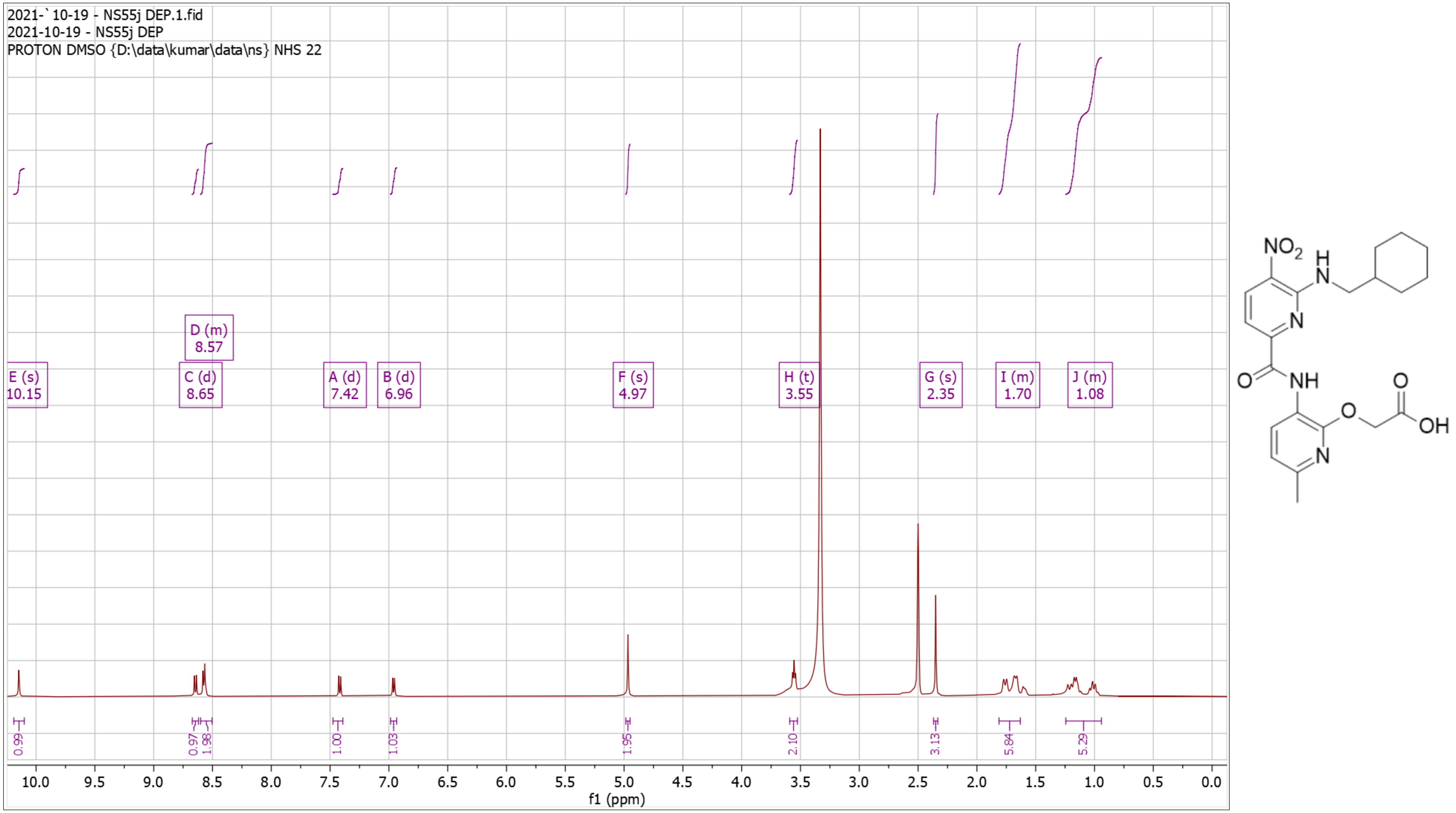

**Supplementary Fig. 31. $^{13}$C-NMR of NS55 Dep**

$^{13}$C NMR (126 MHz, DMSO) δ 23.27, 25.47, 25.94, 30.45, 37.46, 46.79, 62.14, 109.03, 116.85, 118.94, 127.13, 129.82, 138.05, 149.60, 150.92, 151.09, 152.25, 160.39, 169.73.

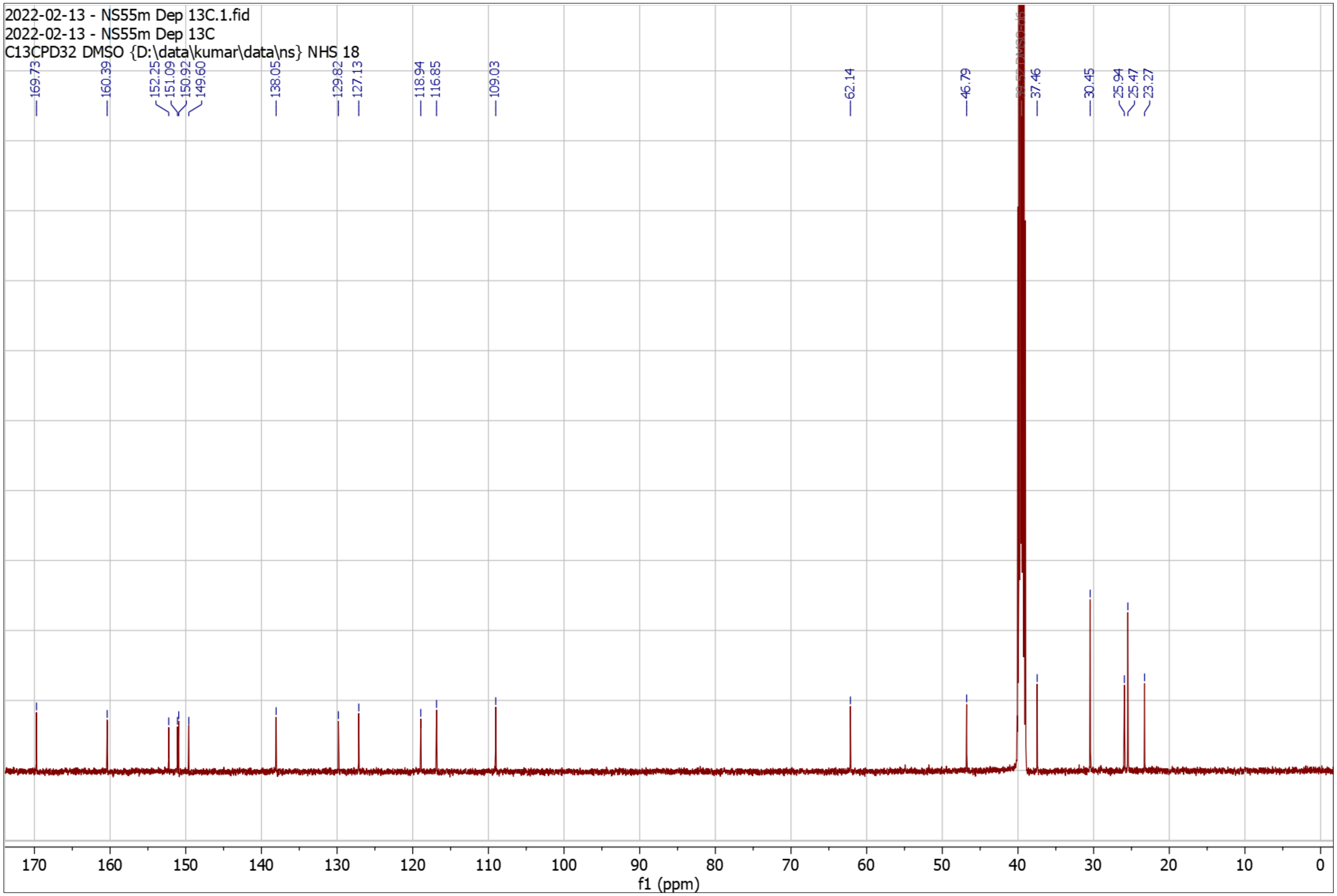


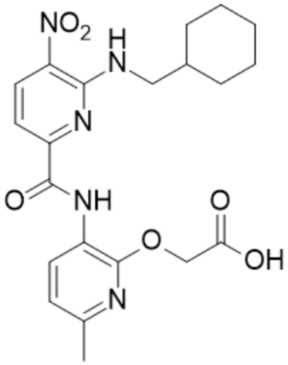

**Supplementary Fig. 32. $^{1}$H-NMR of NS56 Pro**

$^{1}$H NMR (500 MHz, $CDCl_3$) δ 1.46 – 1.46 (s, 9H), 2.34 – 2.44 (s, 3H), 3.02 – 3.12 (t, $J$ = 7.1 Hz, 2H), 3.97 – 4.12 (td, $J$ = 7.1, 5.4 Hz, 2H), 4.62 – 4.69 (s, 2H), 6.77 – 6.86 (d, $J$ = 7.9 Hz, 1H), 7.20 – 7.36 (m, 5H), 7.53 – 7.59 (d, $J$ = 8.5 Hz, 1H), 8.21 – 8.30 (t, $J$ = 5.3 Hz, 1H), 8.54 – 8.63 (d, $J$ = 8.3 Hz, 1H), 8.64 – 8.73 (d, $J$ = 7.9 Hz, 1H), 10.20 – 10.32 (s, 1H); HRMS (m/z): [M]+ calcd. for $C_{26}H_{29}N_5O_6$, 508.2191 ; found, 508.2190.

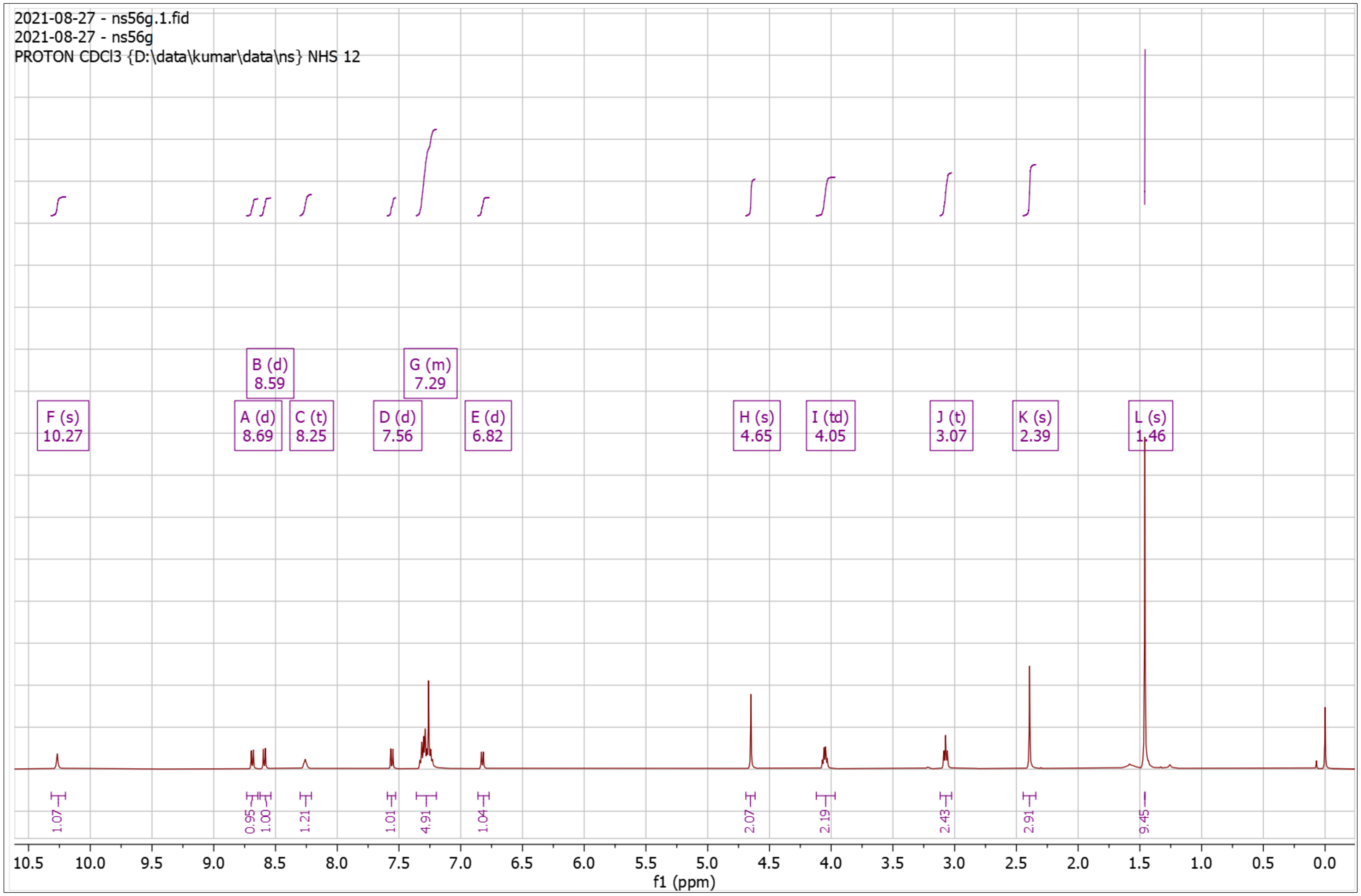


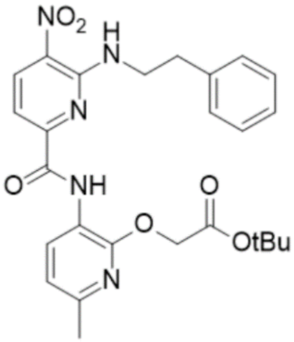

**Supplementary Fig. 33. ¹H-NMR of NS56 Dep**

¹H NMR (500 MHz, DMSO) δ 2.31 – 2.38 (s, 3H), 2.95 – 3.07 (t, *J* = 7.1 Hz, 2H), 3.88 – 4.00 (q, *J* = 6.7 Hz, 2H), 4.69 – 4.77 (s, 2H), 6.90 – 6.99 (d, *J* = 8.0 Hz, 1H), 7.12 – 7.29 (m, 5H), 7.36 – 7.44 (d, *J* = 8.4 Hz, 1H), 8.48 – 8.57 (d, *J* = 7.9 Hz, 1H), 8.57 – 8.61 (t, *J* = 5.8 Hz, 1H), 8.61 – 8.67 (d, *J* = 8.4 Hz, 1H), 10.10 – 10.17 (s, 1H); HRMS (m/z): [M]+ calcd. for $C_{22}H_{21}N_5O_6$, 452.1565; found, 452.1565.

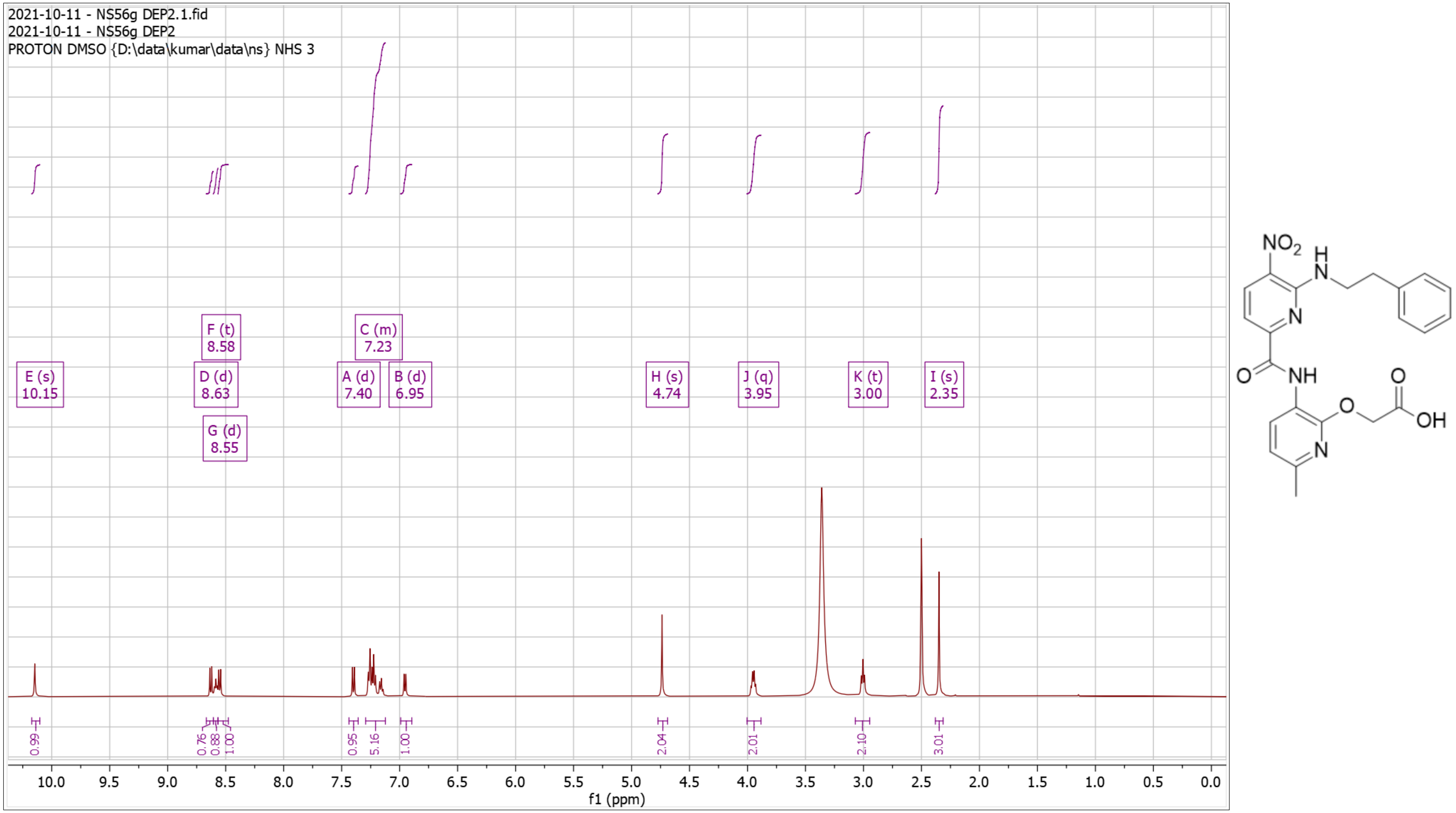

**Supplementary Fig. 34. $^1$H-NMR of NS57 Pro**

$^1$H NMR (500 MHz, $CDCl_3$) δ 1.44 – 1.47 (s, 9H), 1.47 – 1.51 (t, *J* = 7.4 Hz, 3H), 2.26 – 2.50 (s, 3H), 3.27 – 3.47 (q, *J* = 7.4 Hz, 2H), 4.80 – 4.97 (s, 2H), 6.75 – 6.94 (d, *J* = 7.9 Hz, 1H), 8.05 – 8.13 (d, 1H), 8.56 – 8.79 (dd, *J* = 8.2, 5.9 Hz, 2H), 10.21 – 10.35 (s, 1H); HRMS (m/z): [M]+ calcd. for $C_{20}H_{24}N_4O_6S$, 449.1489; found, 449.1488.

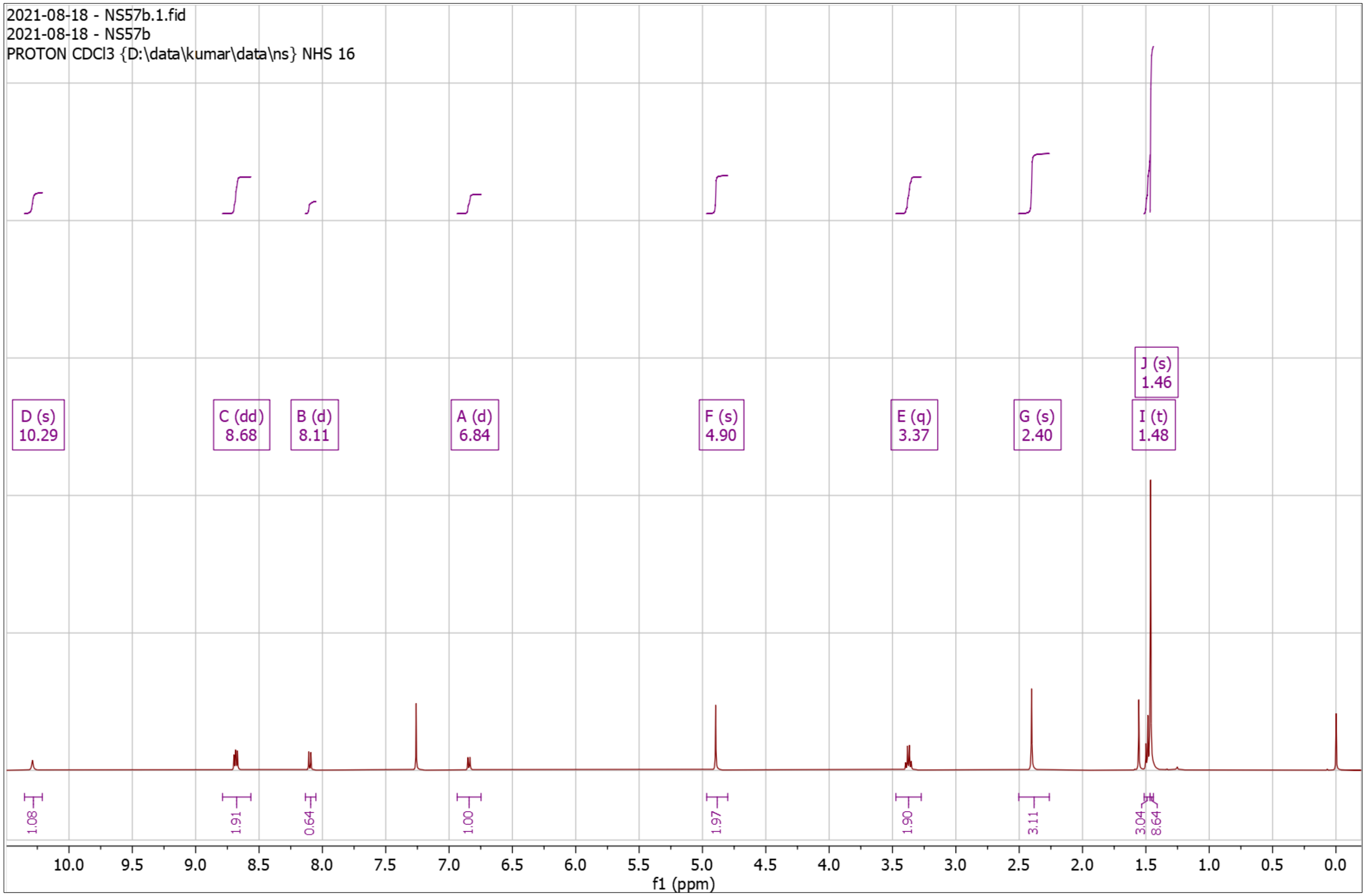


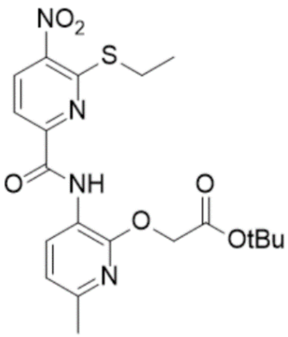

**Supplementary Fig. 35. $^{1}$H-NMR of NS57 Dep**

$^{1}$H NMR (500 MHz, DMSO) δ 1.32 – 1.36 (t, *J* = 7.3 Hz, 3H), 2.31 – 2.37 (s, 3H), 3.09 – 3.20 (q, *J* = 7.3 Hz, 2H), 4.92 – 4.97 (s, 2H), 6.88 – 6.99 (d, *J* = 7.9 Hz, 1H), 7.99 – 8.06 (d, *J* = 8.4 Hz, 1H), 8.06 – 8.14 (d, *J* = 8.1 Hz, 1H), 8.46 – 8.54 (d, *J* = 7.9 Hz, 1H), 9.81 – 9.88 (s, 1H); HRMS (m/z): [M]+ calcd. for $C_{16}H_{16}N_4O_6S$, 393.0863; found, 393.0862.

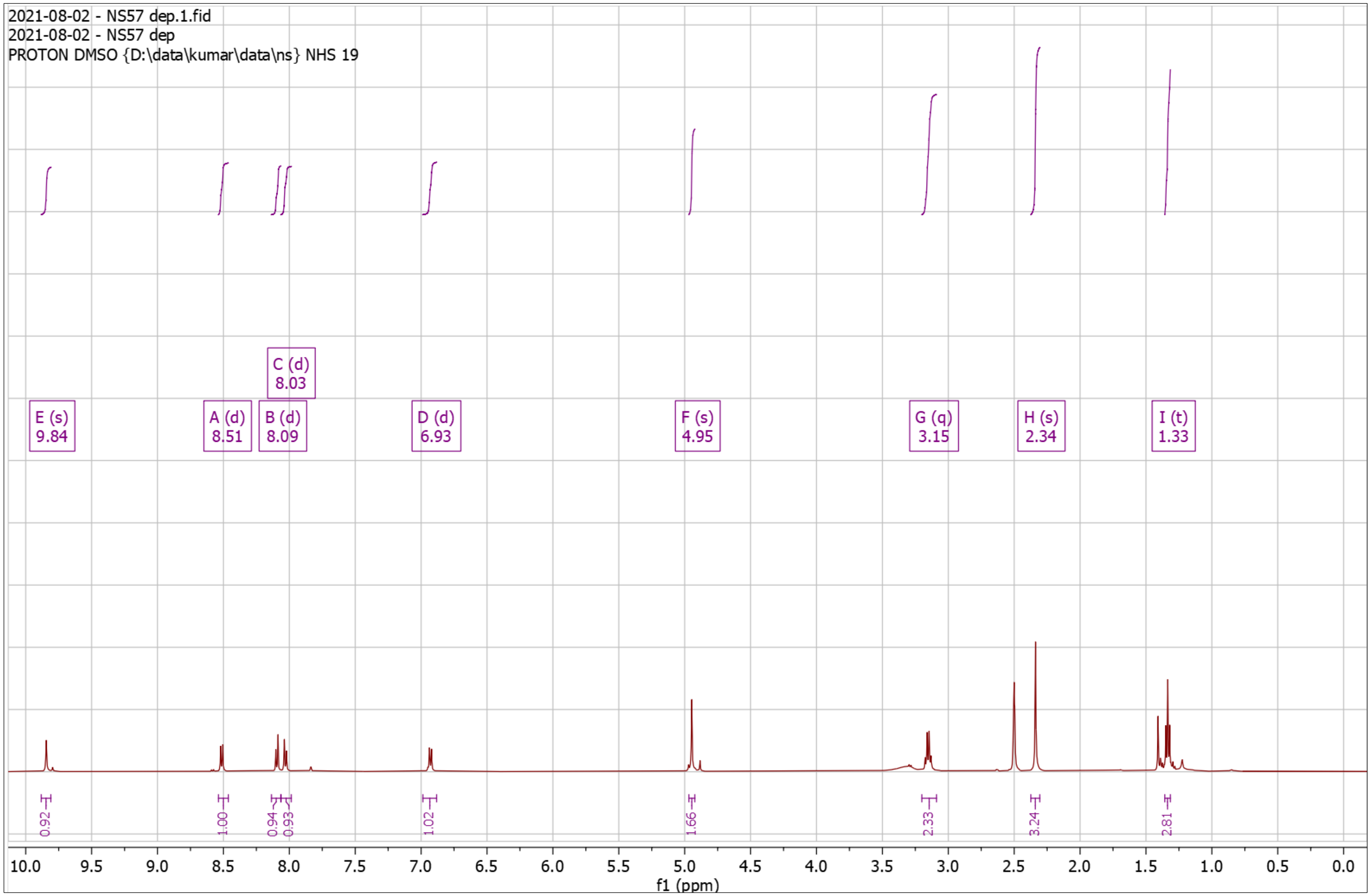


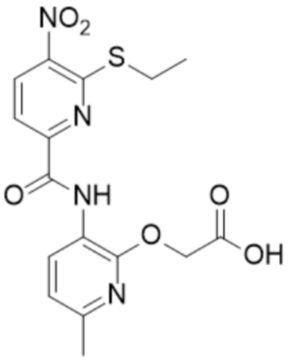

**Supplementary Fig. 36. $^{1}$H-NMR of NS59 Pro**

$^{1}$H NMR (500 MHz, $CDCl_3$) δ 1.10 – 1.15 (d, $J$ = 6.6 Hz, 6H), 1.44 – 1.47 (s, 9H), 1.99 – 2.11 (hept, $J$ = 6.7 Hz, 1H), 2.38 – 2.42 (s, 3H), 3.25 – 3.30 (d, $J$ = 6.7 Hz, 2H), 4.86 – 4.90 (s, 2H), 6.78 – 6.89 (d, $J$ = 8.0 Hz, 1H), 8.03 – 8.13 (d, $J$ = 8.4 Hz, 1H), 8.62 – 8.67 (d, $J$ = 8.3 Hz, 1H), 8.67 – 8.72 (d, $J$ = 7.9 Hz, 1H), 10.12 – 10.15 (s, 1H); HRMS (m/z): [M]+ calcd. for $C_{22}H_{28}N_4O_6S$, 477.1802; found, 477.1803.

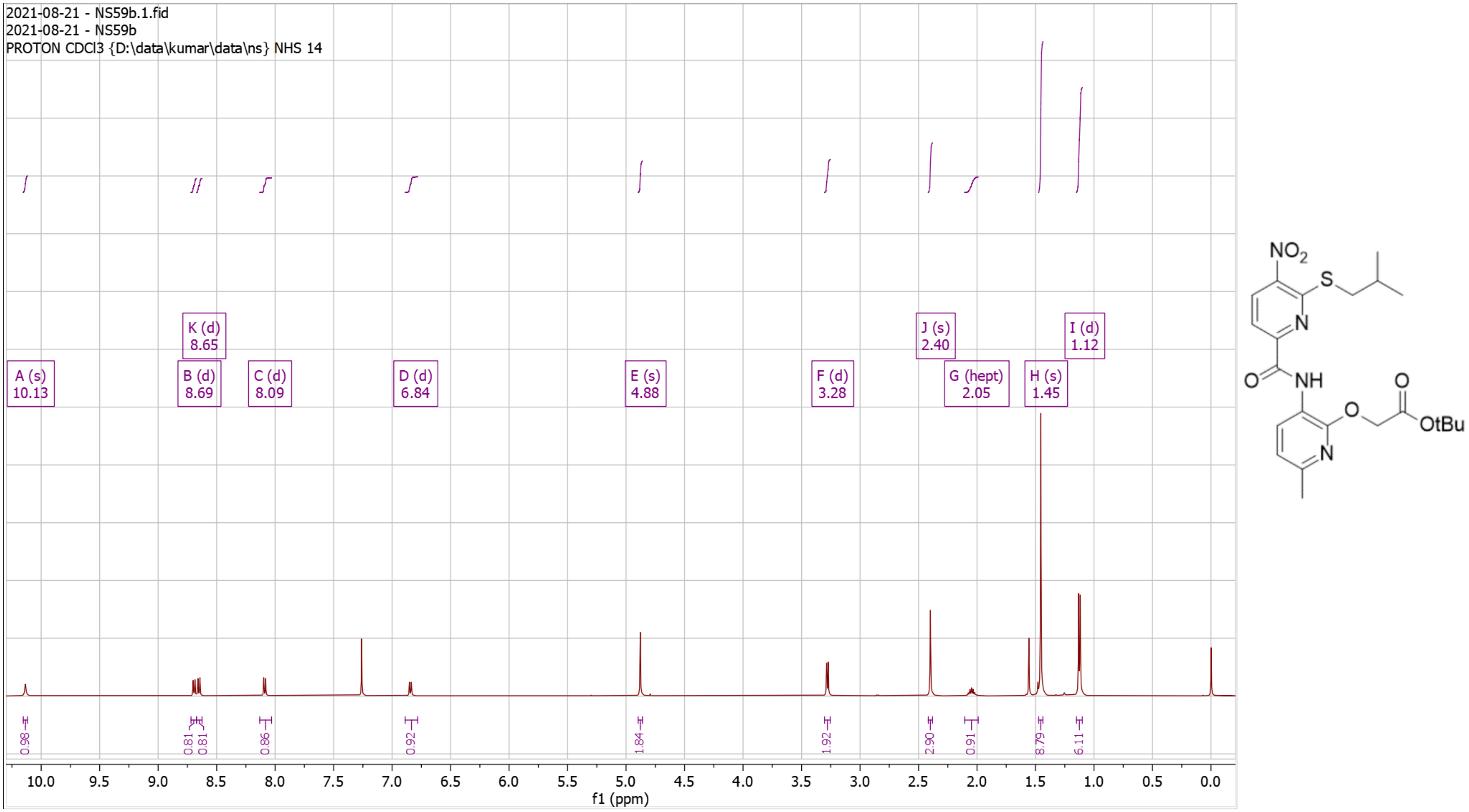

**Supplementary Fig. 37. $^1$H-NMR of NS59 Dep**

$^1$H NMR (500 MHz, DMSO) δ 1.03 – 1.09 (d, *J* = 6.6 Hz, 6H), 1.89 – 1.99 (hept, *J* = 6.7 Hz, 1H), 2.31 – 2.36 (s, 3H), 3.01 – 3.05 (d, *J* = 6.7 Hz, 2H), 4.88 – 4.93 (s, 2H), 6.88 – 6.93 (d, *J* = 8.0 Hz, 1H), 8.01 – 8.06 (d, *J* = 8.2 Hz, 1H), 8.06 – 8.12 (d, *J* = 8.1 Hz, 1H), 8.47 – 8.55 (d, *J* = 7.9 Hz, 1H), 9.84 – 9.93 (s, 1H); HRMS (m/z): [M]+ calcd. for $C_{18}H_{20}N_4O_6S$, 421.1176; found, 421.1171.

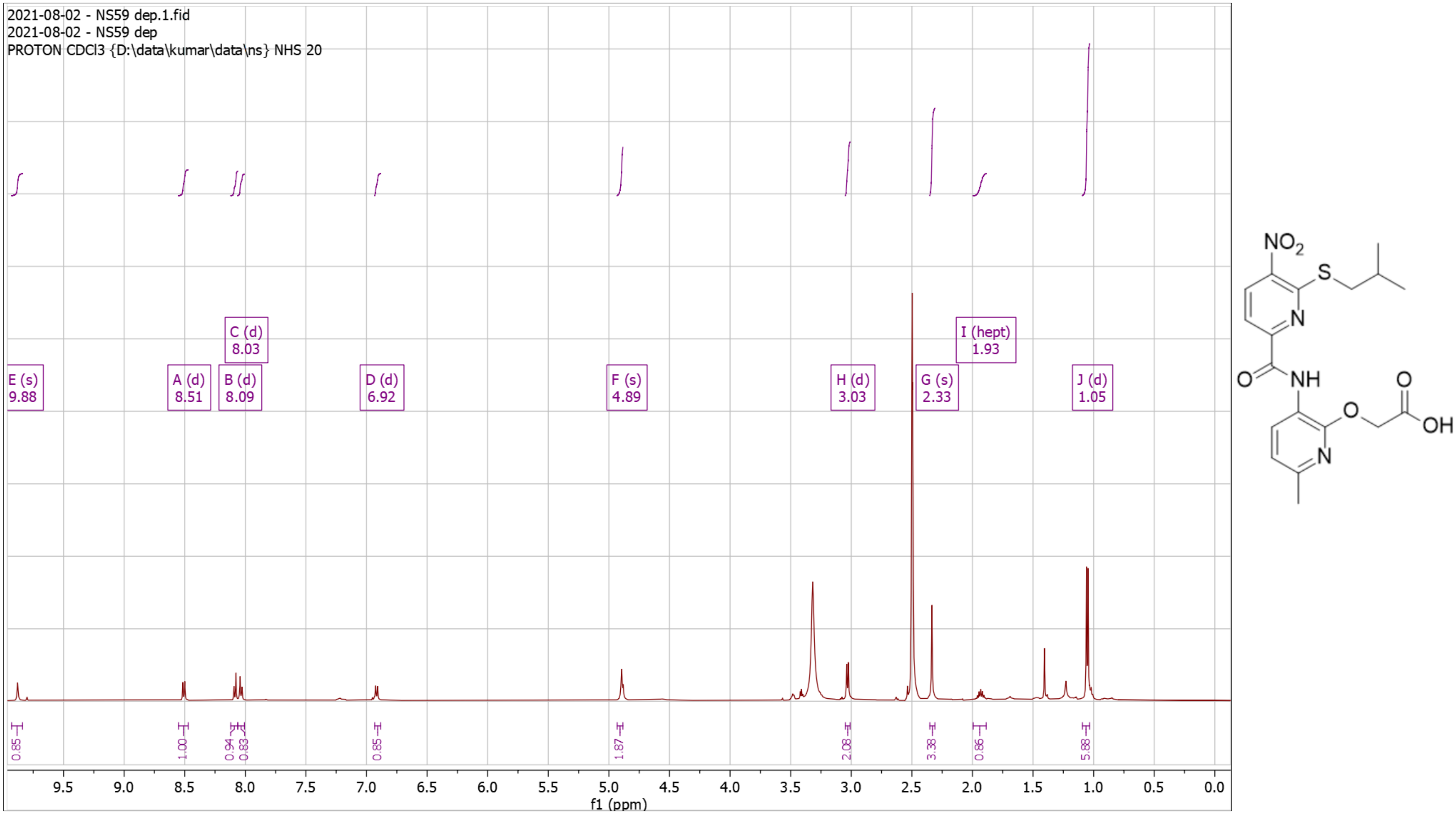

**Supplementary Fig. 38. $^{1}$H-NMR of NS60 Pro**

$^{1}$H NMR (500 MHz, $CDCl_3$) δ 0.95 – 1.03 (t, *J* = 7.4 Hz, 3H), 1.46 – 1.49 (s, 9H), 1.49 – 1.61 (h, *J* = 7.3 Hz, 2H), 1.70 – 1.80 (ddd, *J* = 15.1, 8.6, 6.3 Hz, 2H), 2.35 – 2.39 (s, 3H), 2.95 – 3.01 (t, *J* = 7.4 Hz, 2H), 4.86 – 4.89 (s, 2H), 6.76 – 6.85 (d, *J* = 7.9 Hz, 1H), 7.58 – 7.66 (d, *J* = 8.1 Hz, 1H), 8.07 – 8.14 (d, *J* = 8.1 Hz, 1H), 8.61 – 8.67 (d, *J* = 7.8 Hz, 1H), 9.96 – 10.05 (s, 1H); HRMS (m/z): [M]+ calcd. for $C_{22}H_{28}N_4O_6S$, 477.1802; found, 477.1805.

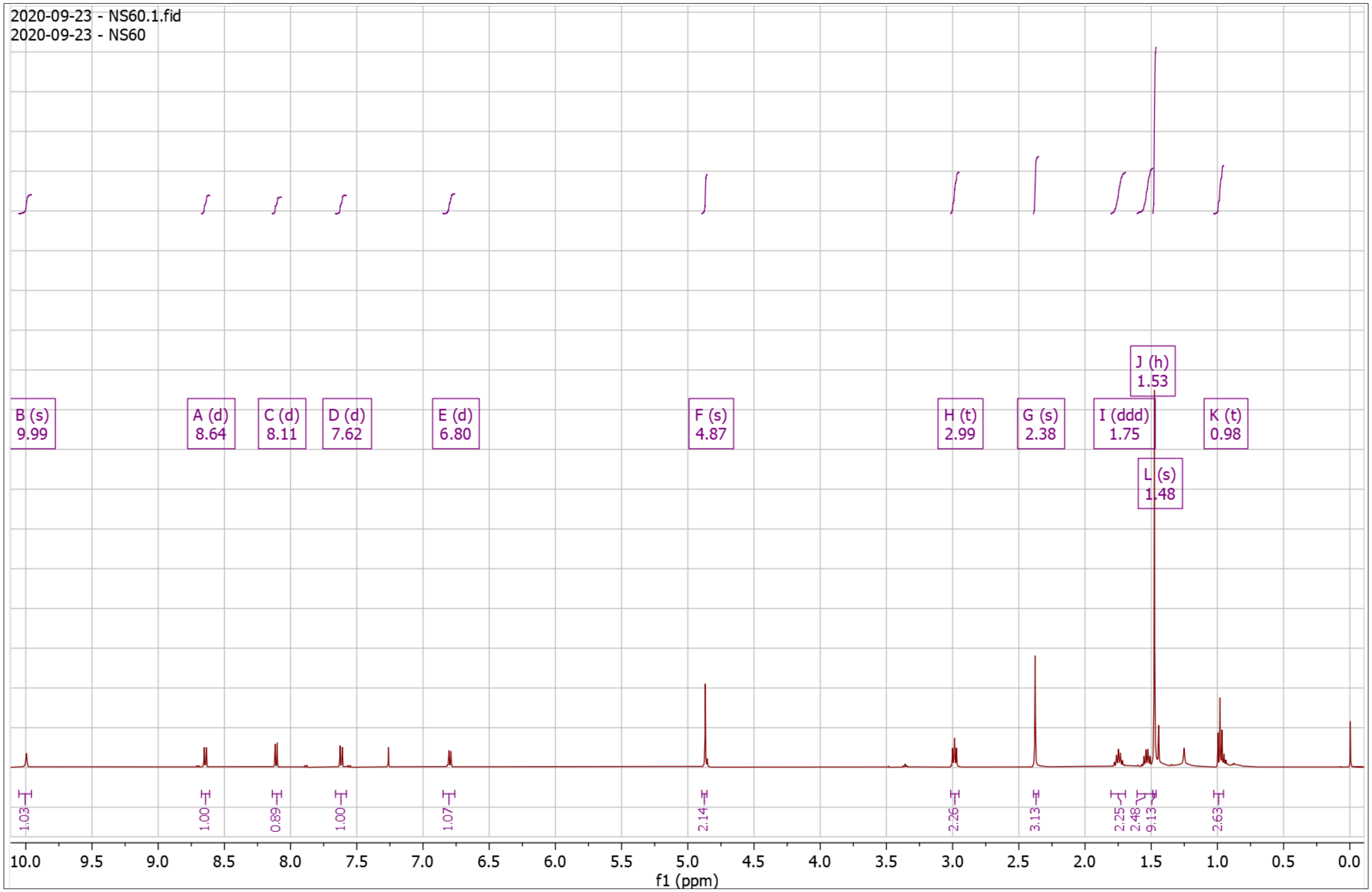


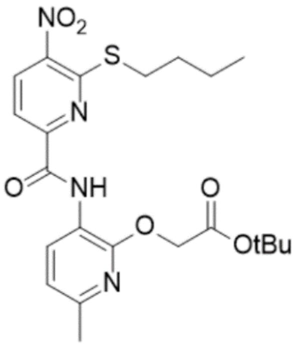

**Supplementary Fig. 39. $^{1}$H-NMR of NS60 Dep**

$^{1}$H NMR (500 MHz, DMSO) δ 0.86 – 0.96 (t, $J$ = 7.4 Hz, 3H), 1.41 – 1.50 (q, $J$ = 7.4 Hz, 2H), 1.54 – 1.72 (p, $J$ = 7.4 Hz, 2H), 2.26 – 2.39 (s, 3H), 3.03 – 3.16 (t, $J$ = 7.3 Hz, 2H), 4.91 – 5.03 (s, 2H), 6.85 – 6.96 (d, $J$ = 7.9 Hz, 1H), 7.95 – 8.03 (d, $J$ = 8.3 Hz, 1H), 8.03 – 8.11 (d, $J$ = 8.1 Hz, 1H), 8.44 – 8.55 (d, $J$ = 7.9 Hz, 1H), 9.78 – 9.83 (s, 1H); HRMS (m/z): [M]+ calcd. for $C_{18}H_{20}N_4O_6S$, 421.1176; found, 421.1176.

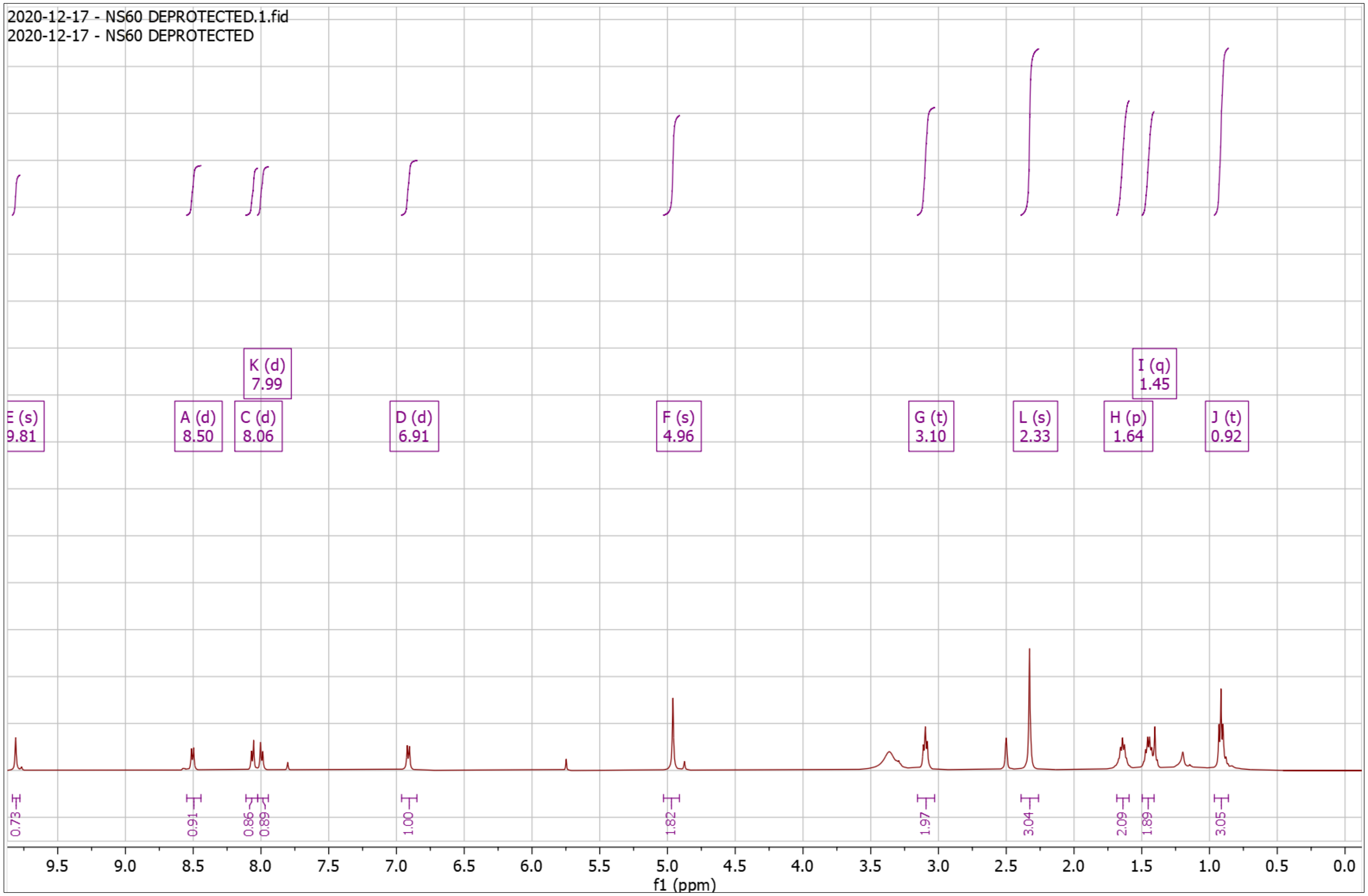


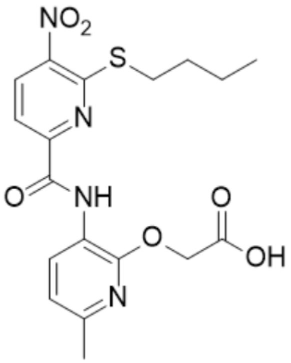

**Supplementary Fig. 40. $^{1}$H-NMR of NS61 Pro**

$^{1}$H NMR (500 MHz, $CDCl_3$) δ 1.46 – 1.50 (s, 9H), 2.36 – 2.40 (s, 3H), 4.22 – 4.26 (s, 2H), 4.85 – 4.89 (s, 2H), 6.25 – 6.29 (d, $J$ = 3.3 Hz, 1H), 6.29 – 6.35 (dd, $J$ = 3.3, 1.9 Hz, 1H), 6.77 – 6.82 (d, $J$ = 7.9 Hz, 1H), 7.35 – 7.41 (d, $J$ = 1.9 Hz, 1H), 7.73 – 7.78 (d, $J$ = 8.1 Hz, 1H), 8.07 – 8.12 (d, $J$ = 8.0 Hz, 1H), 8.61 – 8.66 (d, $J$ = 8.0 Hz, 1H), 9.98 – 10.01 (s, 1H); HRMS (m/z): [M]+ calcd. for $C_{23}H_{24}N_4O_7S$, 501.1439; found, 501.1434.

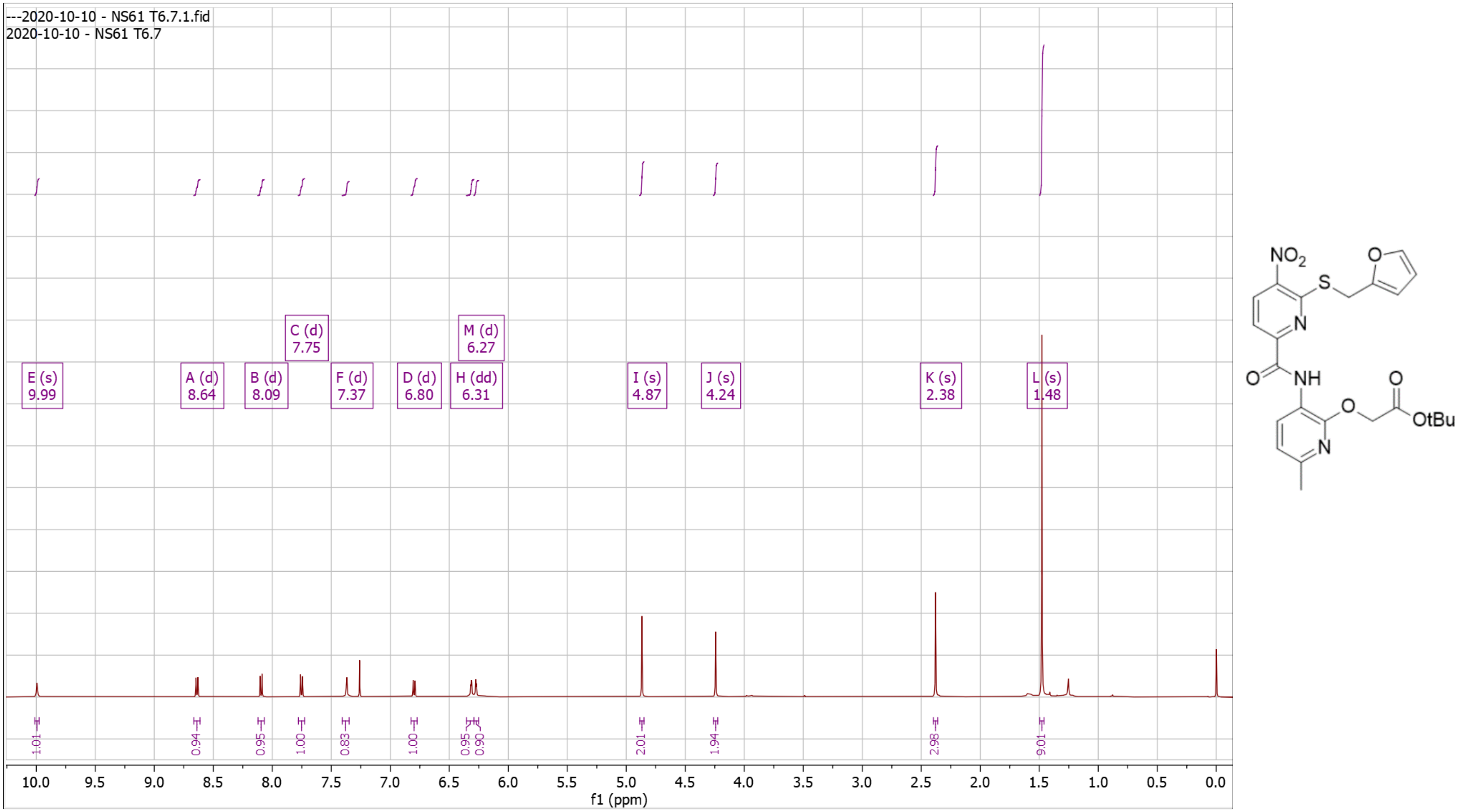

**Supplementary Fig. 41. $^{1}$H-NMR of NS61 Dep**

$^{1}$H NMR (500 MHz, DMSO) δ 2.31 – 2.37 (s, 3H), 4.54 – 4.58 (s, 2H), 4.93 – 4.98 (s, 2H), 6.38 – 6.44 (dd, *J* = 3.2, 1.9 Hz, 1H), 6.44 – 6.50 (d, *J* = 3.3 Hz, 1H), 6.88 – 7.00 (d, *J* = 8.0 Hz, 1H), 7.58 – 7.65 (d, *J* = 1.9 Hz, 1H), 8.08 – 8.15 (d, *J* = 8.1 Hz, 1H), 8.17 – 8.27 (d, *J* = 8.2 Hz, 1H), 8.46 – 8.54 (d, *J* = 7.9 Hz, 1H), 9.80 – 9.89 (s, 1H), 12.77 – 12.97 (s, 1H); HRMS (m/z): [M]+ calcd. for $C_{19}H_{16}N_4O_7S$, 445.0813; found, 445.0821.

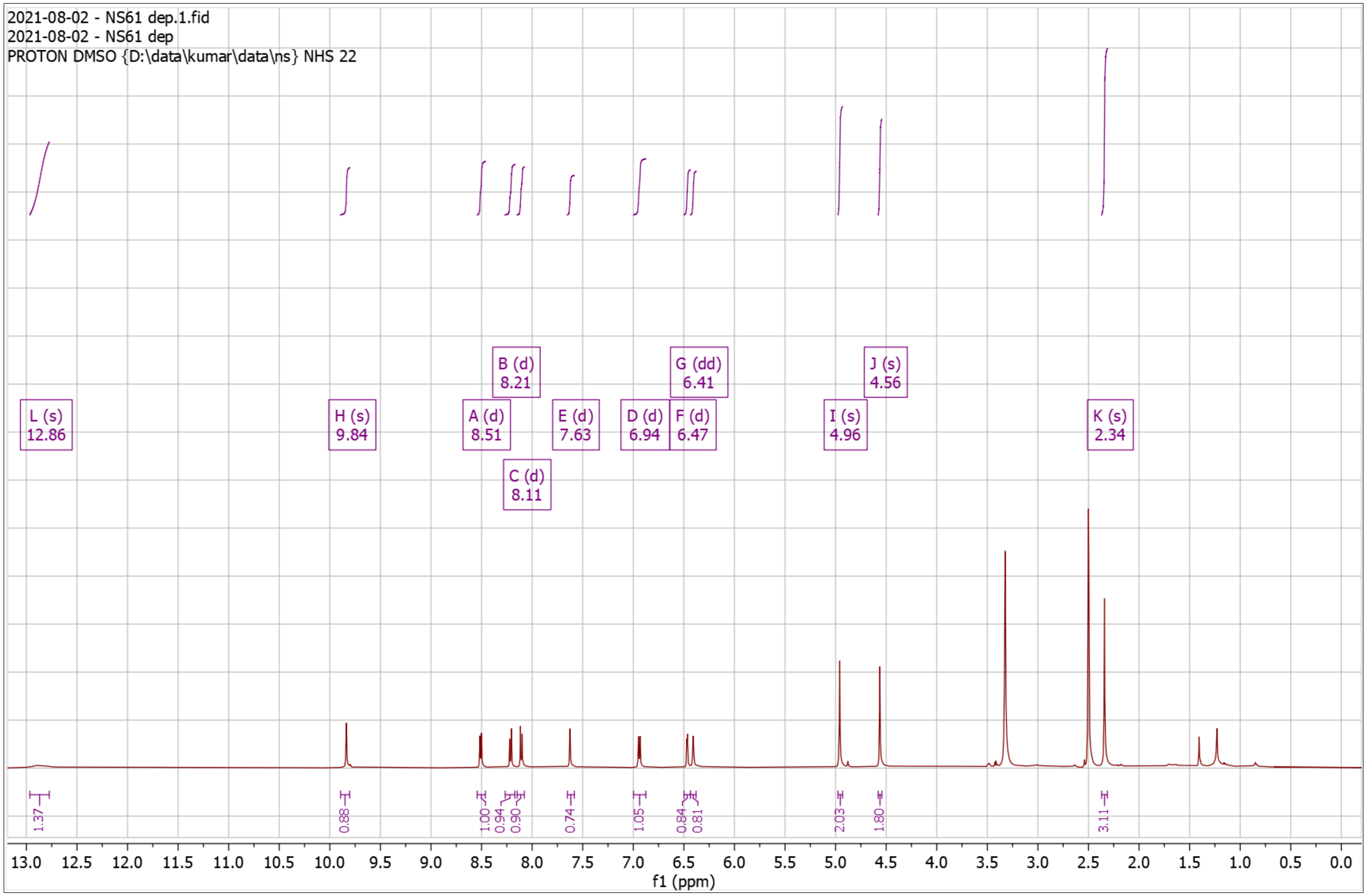


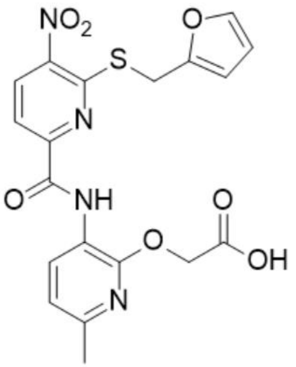

**Supplementary Fig. 42. $^{1}$H-NMR of NS62 Pro**

$^{1}$H NMR (500 MHz, $CDCl_3$) δ 1.46 – 1.49 (s, 18H), 2.36 – 2.41 (s, 3H), 2.67 – 2.74 (t, $J$ = 6.1 Hz, 2H), 4.00 – 4.07 (q, $J$ = 6.0 Hz, 2H), 4.86 – 4.93 (s, 2H), 6.79 – 6.85 (d, $J$ = 7.9 Hz, 1H), 7.55 – 7.61 (d, $J$ = 8.4 Hz, 1H), 8.50 – 8.56 (t, $J$ = 5.9 Hz, 1H), 8.57 – 8.64 (d, $J$ = 8.4 Hz, 1H), 8.64 – 8.71 (d, $J$ = 7.9 Hz, 1H), 10.21 – 10.32 (s, 1H); HRMS (m/z): [M]+ calcd. for $C_{25}H_{33}N_5O_8$, 532.2402; found, 532.2402.

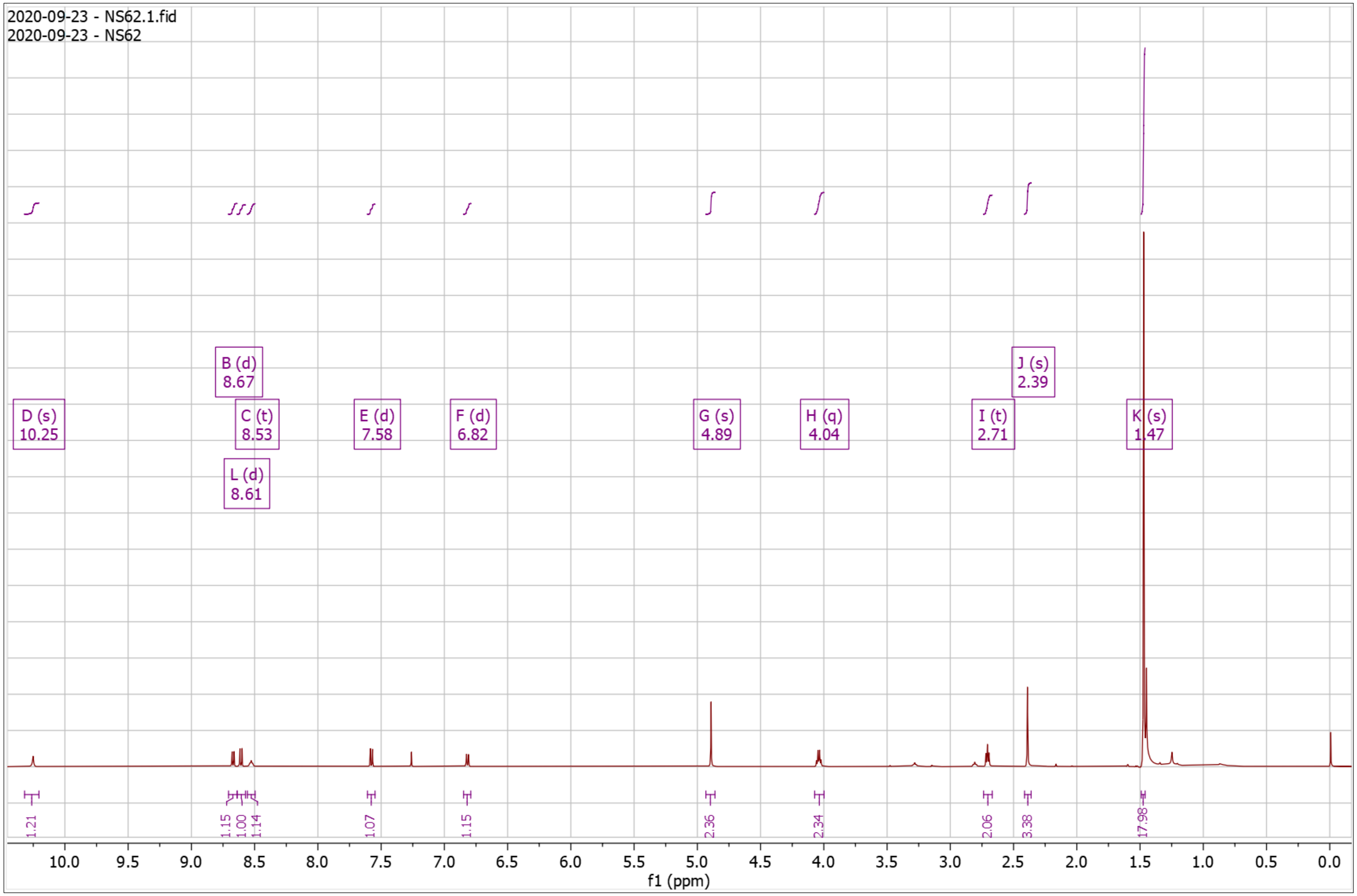


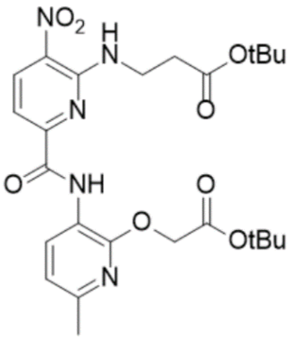

**Supplementary Fig. 43. $^{1}$H-NMR of NS62 Dep**

$^{1}$H NMR (500 MHz, DMSO) δ 2.31 – 2.38 (s, 3H), 2.64 – 2.74 (t, $J$ = 6.8 Hz, 2H), 3.83 – 3.94 (q, $J$ = 6.5 Hz, 2H), 4.86 – 5.00 (s, 2H), 6.91 – 6.96 (d, $J$ = 7.9 Hz, 1H), 7.41 – 7.48 (d, $J$ = 8.7 Hz, 1H), 8.50 – 8.55 (d, $J$ = 7.9 Hz, 1H), 8.59 – 8.72 (dd, 2H), 10.14 – 10.36 (s, 1H); HRMS (m/z): [M]+ calcd. for $C_{17}H_{17}N_5O_8$, 420.1150; found, 420.1143.

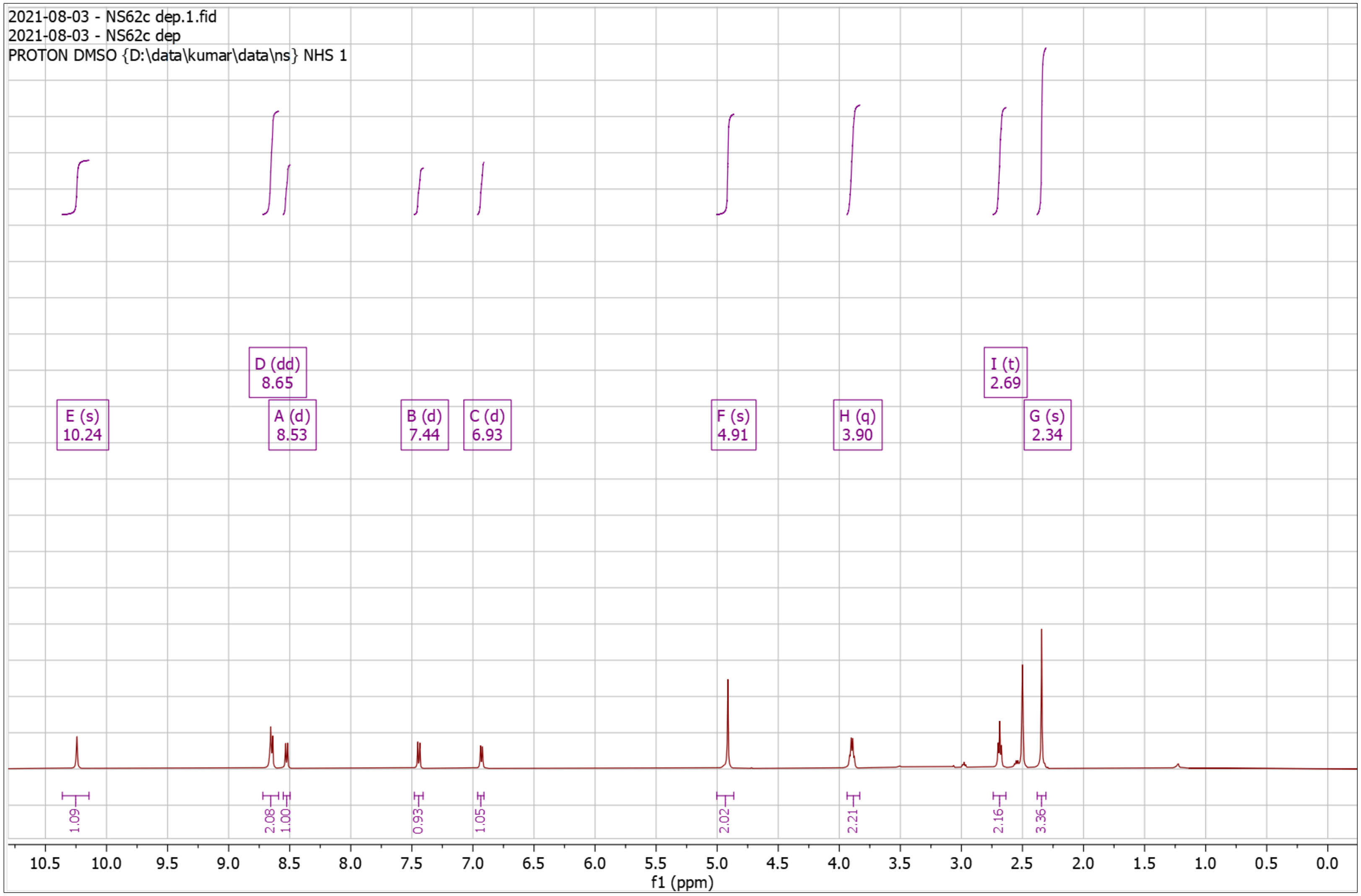


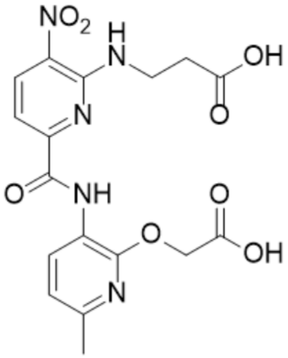

**Supplementary Fig. 44. $^{1}$H-NMR of NS71 Pro**

$^{1}$H NMR (500 MHz, DMSO) δ 1.35 – 1.46 (s, 9H), 2.32 – 2.36 (s, 3H), 2.81 – 2.92 (t, *J* = 6.4 Hz, 2H), 3.88 – 3.97 (q, *J* = 6.0 Hz, 2H), 4.94 – 4.98 (s, 2H), 6.86 – 7.02 (d, *J* = 7.9 Hz, 1H), 7.37 – 7.50 (d, *J* = 8.3 Hz, 1H), 8.47 – 8.53 (d, *J* = 8.0 Hz, 1H), 8.61 – 8.71 (d, *J* = 8.3 Hz, 1H), 9.06 – 9.19 (t, *J* = 4.9 Hz, 1H), 10.19 – 10.30 (s, 1H); HRMS (m/z): [M]+ calcd. for $C_{20}H_{25}N_5O_9S$, 512.1446; found, 512.1439.

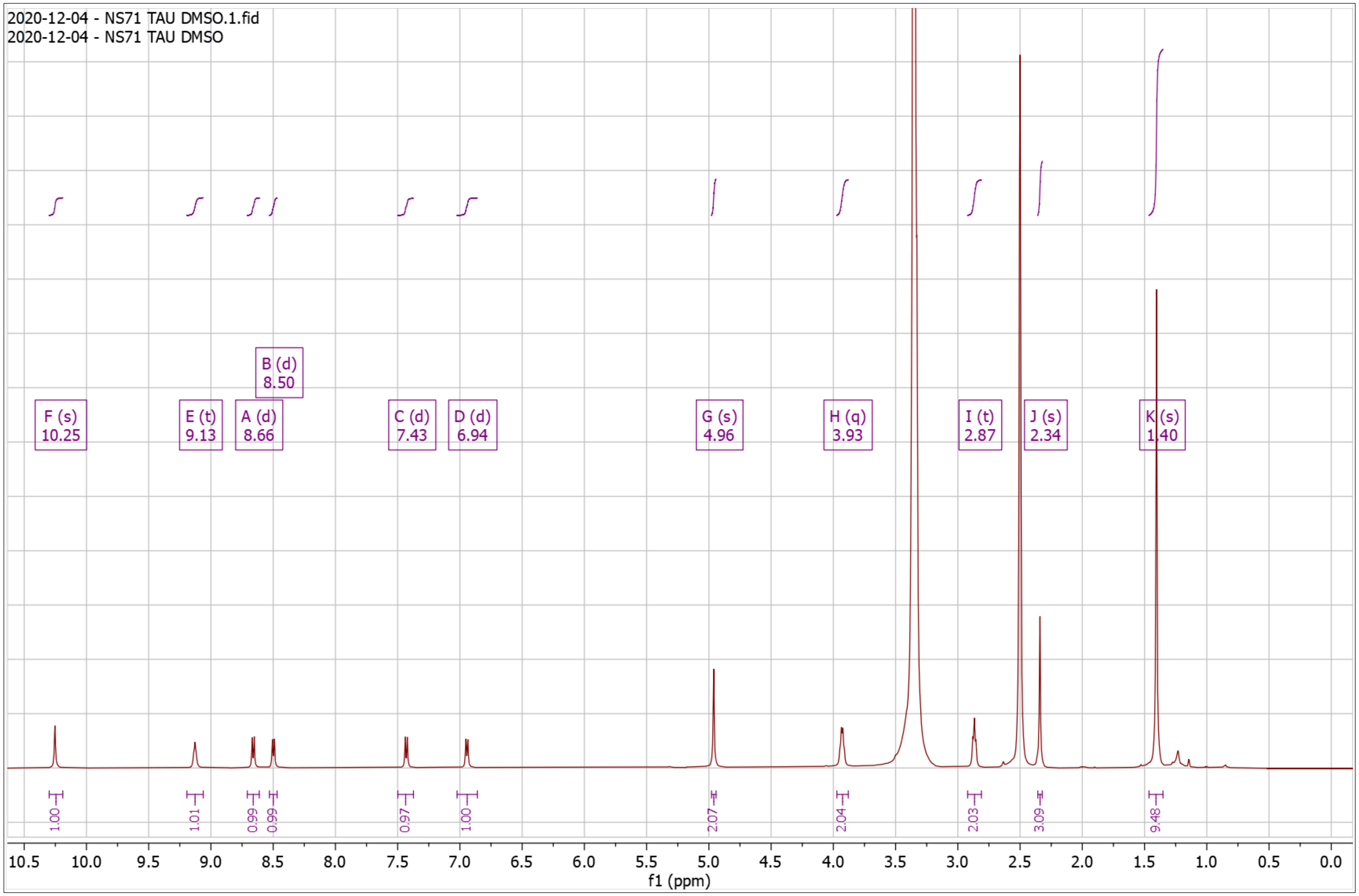


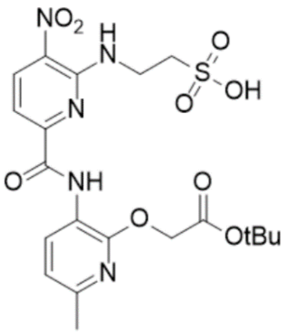

**Supplementary Fig. 45. $^{1}$H-NMR of NS71 Dep**

$^{1}$H NMR (500 MHz, DMSO) δ 2.26 – 2.41 (s, 3H), 2.82 – 2.91 (t, *J* = 6.2 Hz, 2H), 3.90 – 4.02 (q, *J* = 5.8 Hz, 2H), 4.88 – 5.02 (s, 2H), 6.87 – 6.97 (d, *J* = 7.9 Hz, 1H), 7.37 – 7.48 (d, *J* = 8.4 Hz, 1H), 8.41 – 8.54 (d, *J* = 7.9 Hz, 1H), 8.59 – 8.70 (d, *J* = 8.4 Hz, 1H), 9.07 – 9.20 (t, *J* = 5.1 Hz, 1H), 10.23 – 10.42 (s, 1H); HRMS (m/z): [M]+ calcd. for $C_{16}H_{17}N_5O_9S$, 456.0820; found, 456.0820.

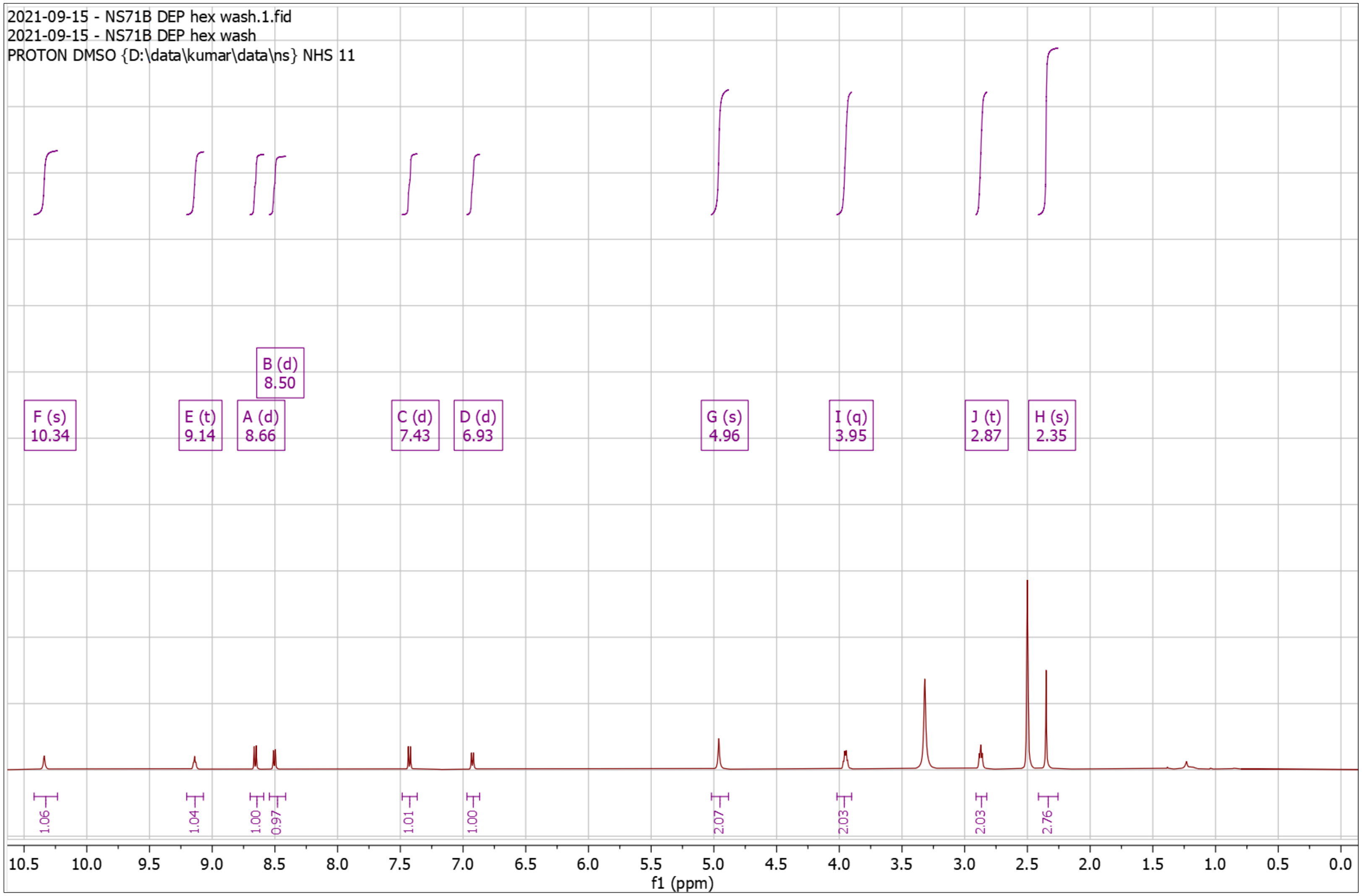


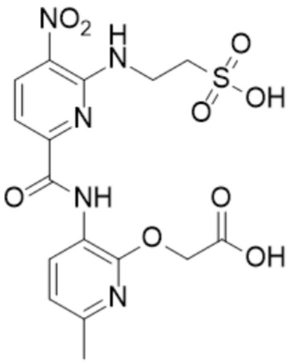

**Supplementary Fig. 46. $^{1}$H-NMR of NS72 Pro**

$^{1}$H NMR (500 MHz, $CDCl_3$) δ 1.45 – 1.50 (s, 9H), 2.36 – 2.43 (s, 3H), 3.18 – 3.29 (t, *J* = 6.8 Hz, 2H), 4.07 – 4.17 (q, *J* = 6.5 Hz, 2H), 4.63 – 4.67 (s, 2H), 6.80 – 6.86 (d, *J* = 7.9 Hz, 1H), 7.04 – 7.10 (t, *J* = 7.5 Hz, 1H), 7.14 – 7.22 (m, 2H), 7.34 – 7.39 (d, *J* = 8.2 Hz, 1H), 7.49 – 7.55 (d, *J* = 8.4 Hz, 1H), 7.61 – 7.66 (d, *J* = 8.0 Hz, 1H), 8.05 – 8.08 (s, 1H), 8.29 – 8.33 (t, 1H), 8.53 – 8.58 (d, *J* = 8.4 Hz, 1H), 8.65 – 8.70 (d, *J* = 7.9 Hz, 1H), 10.17 – 10.28 (s, 1H); HRMS (m/z): [M]+ calcd. for $C_{28}H_{30}N_6O_6$, 547.2300; found, 547.2299.

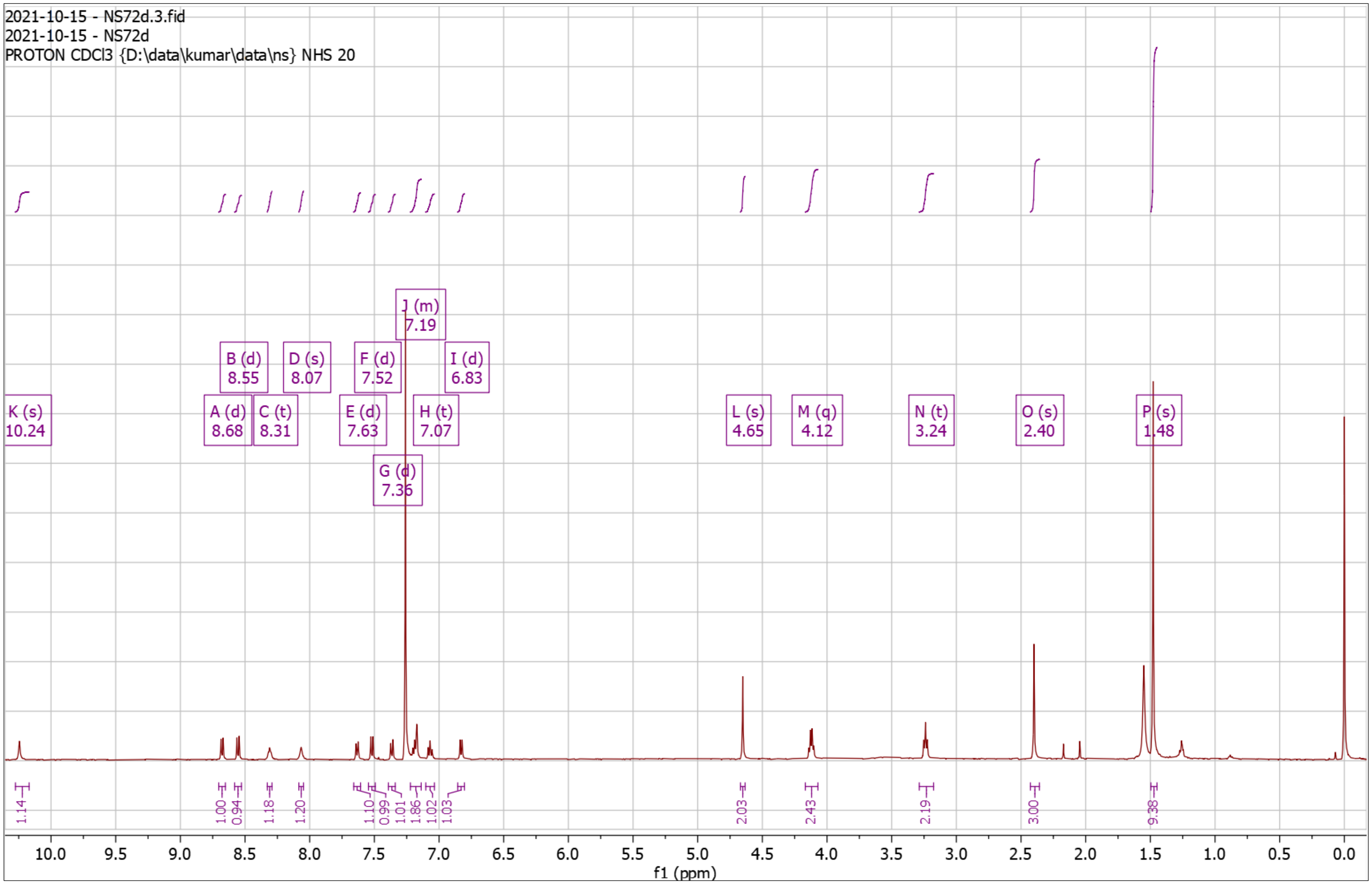


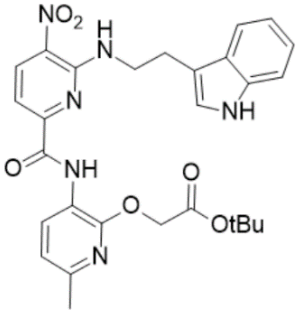

**Supplementary Fig. 47. $^{1}$H-NMR of NS72 Dep**

$^{1}$H NMR (500 MHz, DMSO) δ 2.32 – 2.35 (s, 3H), 3.05 – 3.12 (t, *J* = 6.8 Hz, 2H), 3.93 – 4.04 (q, *J* = 6.5 Hz, 2H), 4.61 – 4.65 (s, 2H), 6.81 – 6.86 (t, *J* = 7.4 Hz, 1H), 6.86 – 6.92 (d, *J* = 7.9 Hz, 1H), 6.94 – 7.02 (t, *J* = 7.5 Hz, 1H), 7.22 – 7.25 (d, *J* = 2.3 Hz, 1H), 7.24 – 7.30 (d, *J* = 8.1 Hz, 1H), 7.31 – 7.38 (d, *J* = 8.4 Hz, 1H), 7.50 – 7.56 (d, *J* = 7.9 Hz, 1H), 8.48 – 8.53 (d, *J* = 7.8 Hz, 1H), 8.53 – 8.65 (dd, *J* = 9.6, 6.9 Hz, 2H), 10.33 – 10.41 (s, 1H), 10.79 – 10.87 (m, 1H). HRMS (m/z): [M]+ calcd. for $C_{24}H_{22}N_6O_6$, 491.1674; found, 491.1678.

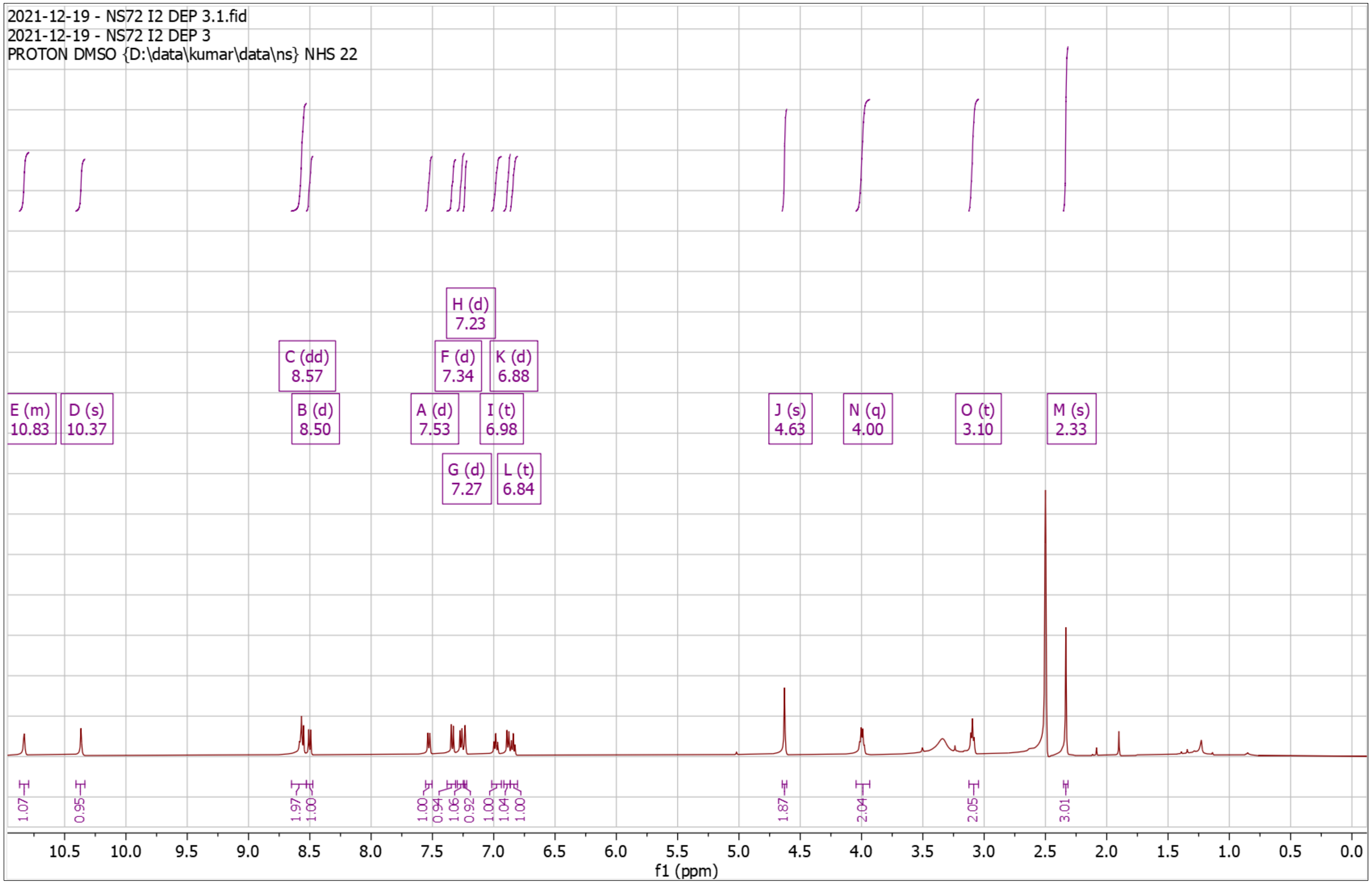


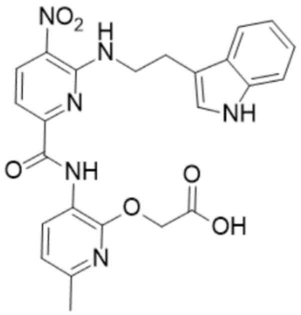

**Supplementary Fig. 48. $^1$H-NMR of NS119 Pro**

$^1$H NMR (500 MHz, $CDCl_3$) δ 1.39 – 1.41 (s, 9H), 1.48 – 1.51 (s, 9H), 2.36 – 2.44 (s, 3H), 3.47 – 3.60 (q, $J$ = 6.0 Hz, 2H), 3.84 – 3.99 (q, $J$ = 5.9 Hz, 2H), 4.84 – 4.95 (s, 2H), 4.99 – 5.09 (s, 1H), 6.80 – 6.86 (d, $J$ = 7.9 Hz, 1H), 7.56 – 7.62 (d, $J$ = 8.4 Hz, 1H), 8.35 – 8.47 (s, 1H), 8.59 – 8.64 (d, $J$ = 8.4 Hz, 1H), 8.65 – 8.70 (d, $J$ = 7.9 Hz, 1H), 10.13 – 10.25 (s, 1H); HRMS (m/z): [M]+ calcd. for $C_{25}H_{34}N_6O_8$, 547.2511; found, 547.2510.

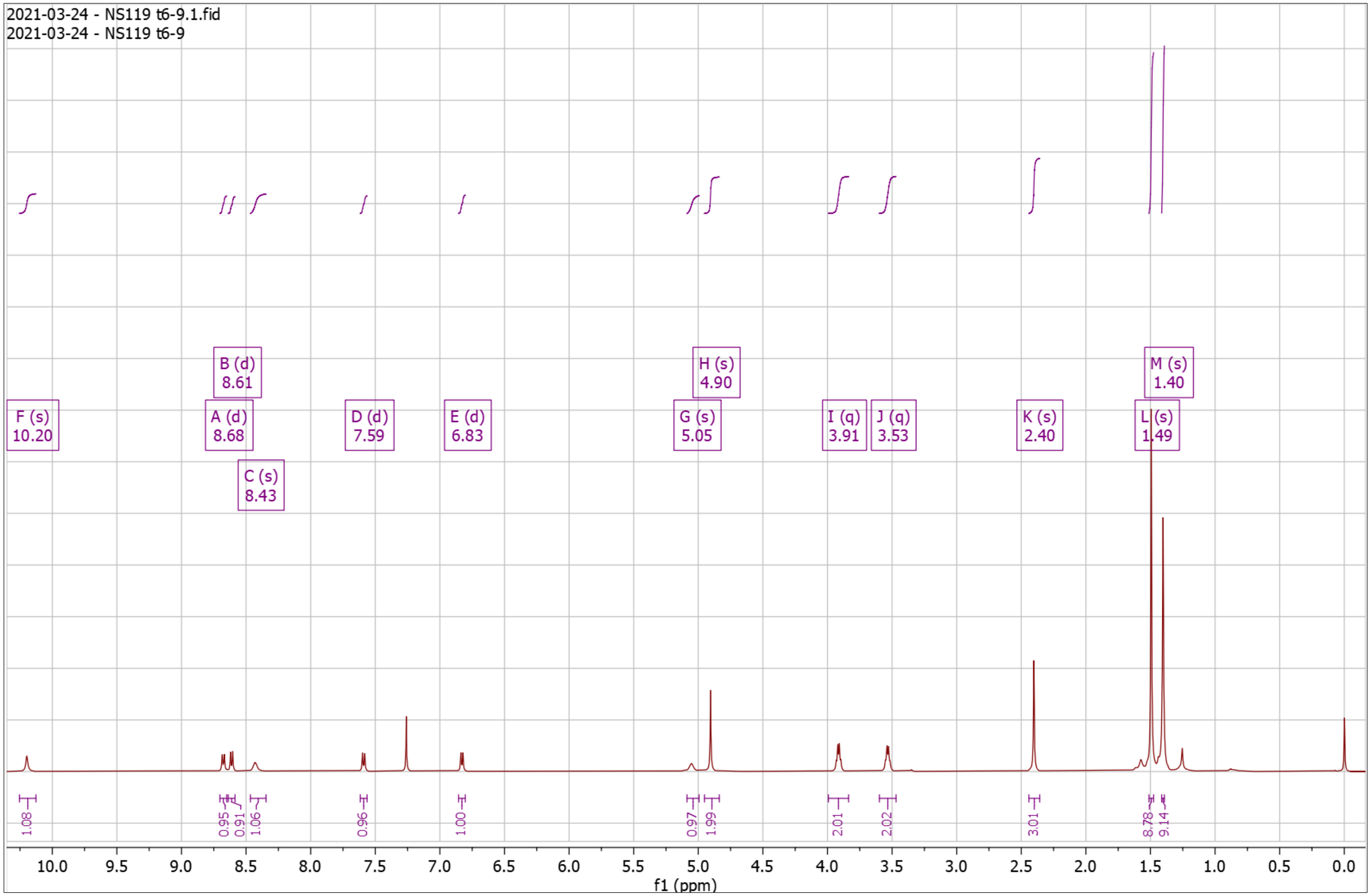


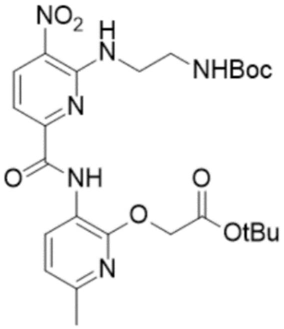

**Supplementary Fig. 49. $^{1}$H-NMR of NS119 Dep**

$^{1}$H NMR (500 MHz, DMSO) δ 2.32 – 2.38 (s, 3H), 3.13 – 3.21 (t, $J$ = 5.5 Hz, 2H), 3.89 – 3.98 (q, $J$ = 5.6 Hz, 2H), 4.96 – 4.99 (s, 2H), 6.94 – 6.99 (d, $J$ = 7.9 Hz, 1H), 7.48 – 7.53 (d, $J$ = 8.4 Hz, 1H), 7.72 – 7.89 (s, 4H), 8.44 – 8.49 (d, $J$ = 7.8 Hz, 1H), 8.58 – 8.66 (t, $J$ = 5.9 Hz, 1H), 8.66 – 8.74 (d, $J$ = 8.4 Hz, 1H), 10.07 – 10.21 (s, 1H); HRMS (m/z): [M]+ calcd. for $C_{16}H_{18}N_6O_6$, 391.1361; found, 391.1365.

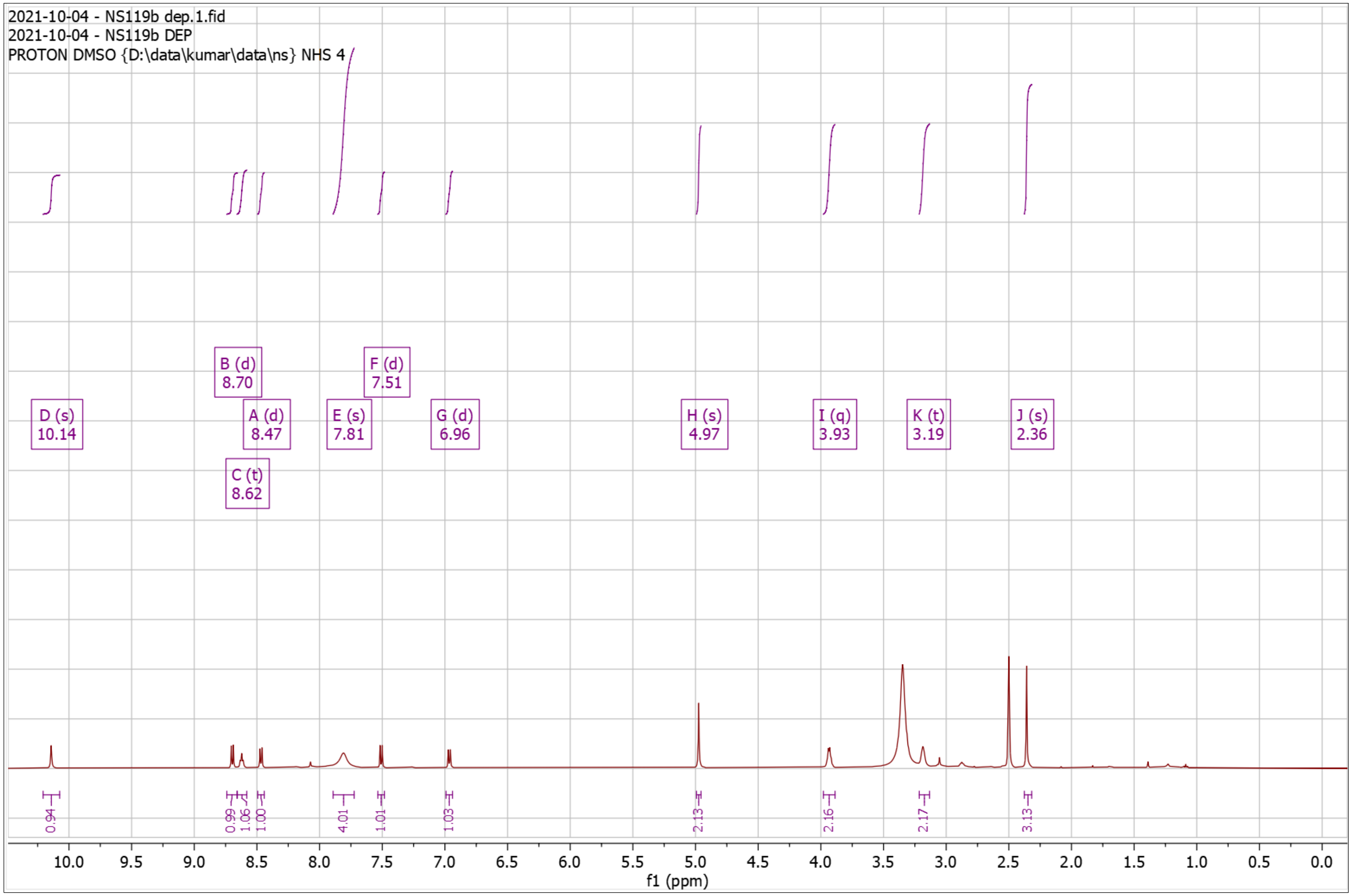


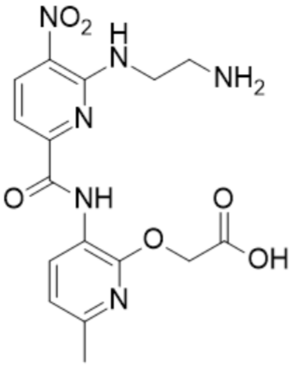

**Supplementary Fig. 50. ¹H-NMR of NS137 Pro**

¹H NMR (500 MHz, $CDCl_3$) δ 1.02 – 1.10 (d, *J* = 6.7 Hz, 6H), 1.41 – 1.49 (s, 9H), 1.97 – 2.14 (dp, *J* = 13.6, 6.9 Hz, 1H), 2.35 – 2.41 (s, 3H), 3.52 – 3.67 (t, *J* = 6.4 Hz, 2H), 4.79 – 4.94 (s, 2H), 6.72 – 6.88 (d, *J* = 7.8 Hz, 1H), 7.48 – 7.57 (d, *J* = 8.4 Hz, 1H), 8.25 – 8.42 (t, *J* = 6.0 Hz, 1H), 8.53 – 8.62 (d, *J* = 8.4 Hz, 1H), 8.62 – 8.69 (d, *J* = 7.9 Hz, 1H), 10.09 – 10.31 (s, 1H); HRMS (m/z): [M]+ calcd. for $C_{22}H_{29}N_5O_6$, 460.2191; found, 460.2188.

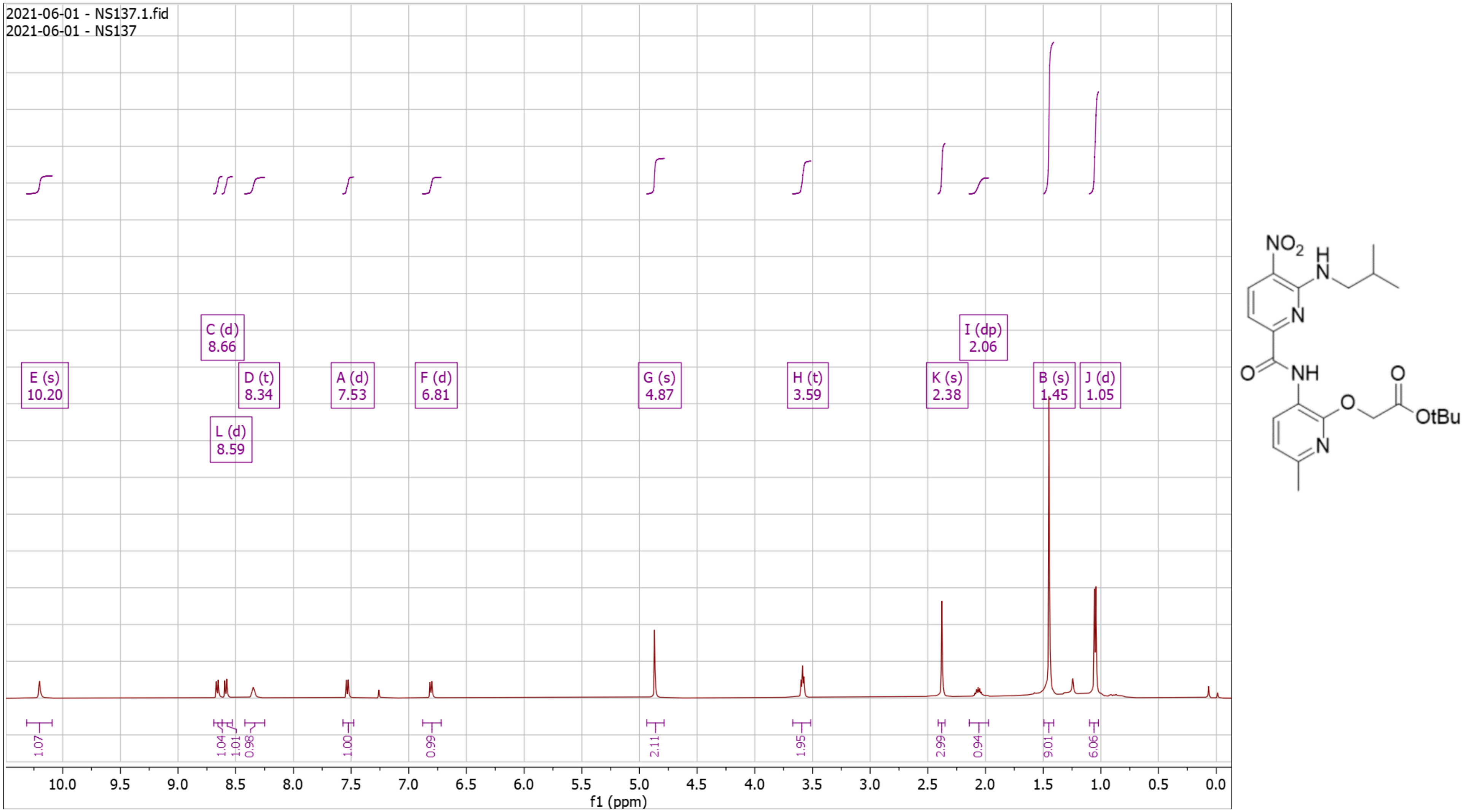

**Supplementary Fig. 51. $^{1}$H-NMR of NS137 Dep**

$^{1}$H NMR (500 MHz, DMSO) δ 0.94 – 0.96 (d, *J* = 6.7 Hz, 6H), 1.99 – 2.10 (tt, *J* = 13.3, 6.4 Hz, 1H), 2.33 – 2.37 (s, 3H), 3.48 – 3.54 (t, *J* = 6.4 Hz, 2H), 4.96 – 4.99 (s, 2H), 6.93 – 6.99 (d, *J* = 7.9 Hz, 1H), 7.40 – 7.45 (d, *J* = 8.3 Hz, 1H), 8.51 – 8.58 (d, *J* = 8.0 Hz, 1H), 8.58 – 8.63 (t, *J* = 6.0 Hz, 1H), 8.62 – 8.70 (d, *J* = 8.4 Hz, 1H), 10.10 – 10.17 (s, 1H); HRMS (m/z): [M]+ calcd. for $C_{18}H_{21}N_5O_6$, 404.1565; found, 404.1565.

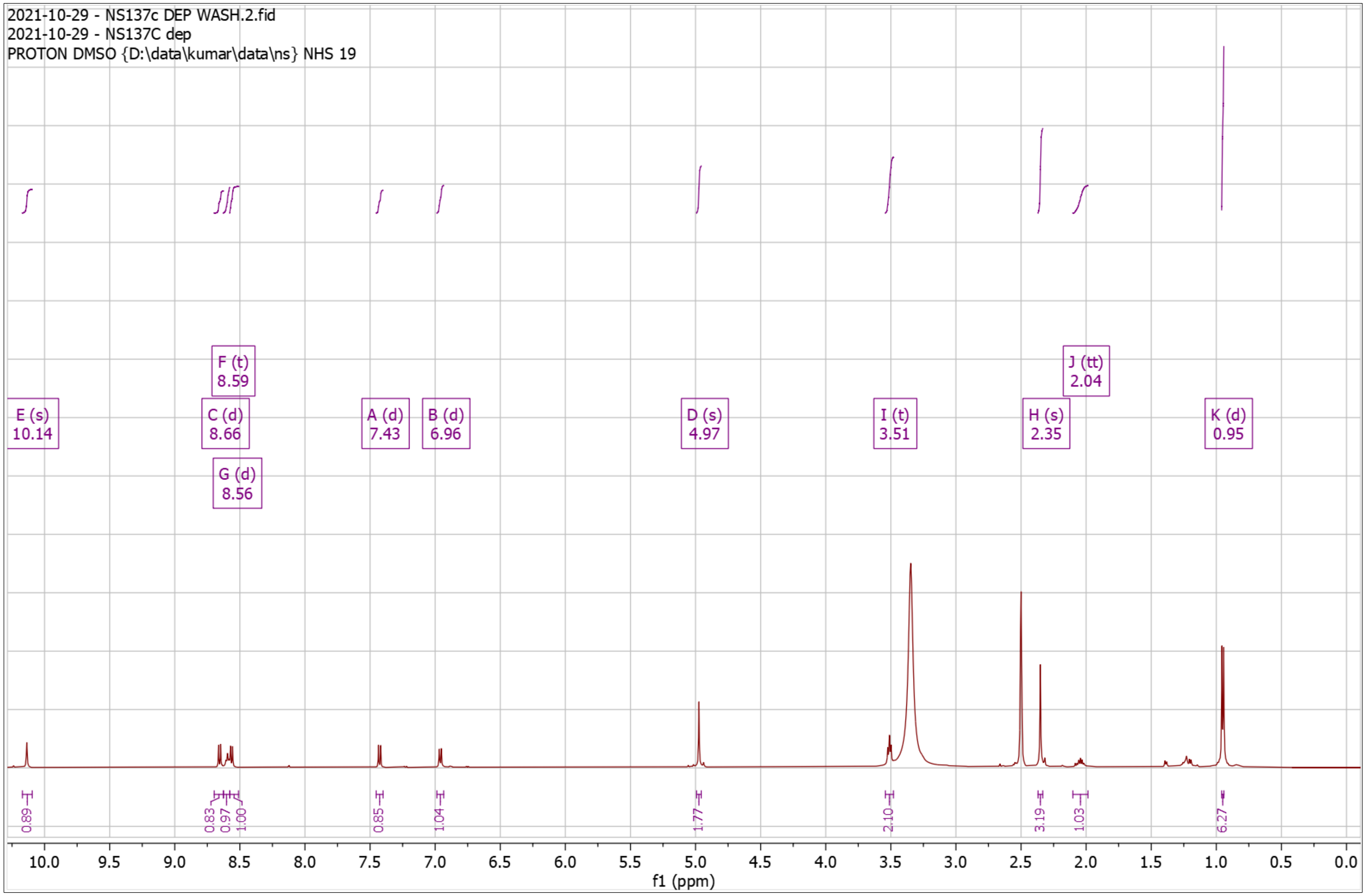


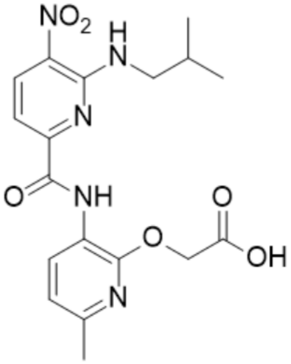

**Supplementary Fig. 52. $^1$H-NMR of NS157 Pro**

$^1$H NMR (500 MHz, $CDCl_3$) δ 0.80 – 0.95 (m, 5H), 1.30 – 1.39 (m, 4H), 1.46 – 1.47 (s, 9H), 1.71 – 1.82 (p, *J* = 7.1 Hz, 2H), 2.36 – 2.41 (s, 3H), 3.71 – 3.81 (q, *J* = 6.5 Hz, 2H), 4.84 – 4.88 (s, 2H), 6.80 – 6.85 (d, *J* = 7.9 Hz, 1H), 7.53 – 7.58 (d, *J* = 8.3 Hz, 1H), 8.22 – 8.28 (t, *J* = 5.8 Hz, 1H), 8.58 – 8.63 (d, 1H), 8.65 – 8.71 (d, *J* = 7.9 Hz, 1H), 10.25 – 10.29 (s, 1H); HRMS (m/z): [M]+ calcd. for $C_{24}H_{33}N_5O_6$, 488.2504; found, 488.2503.

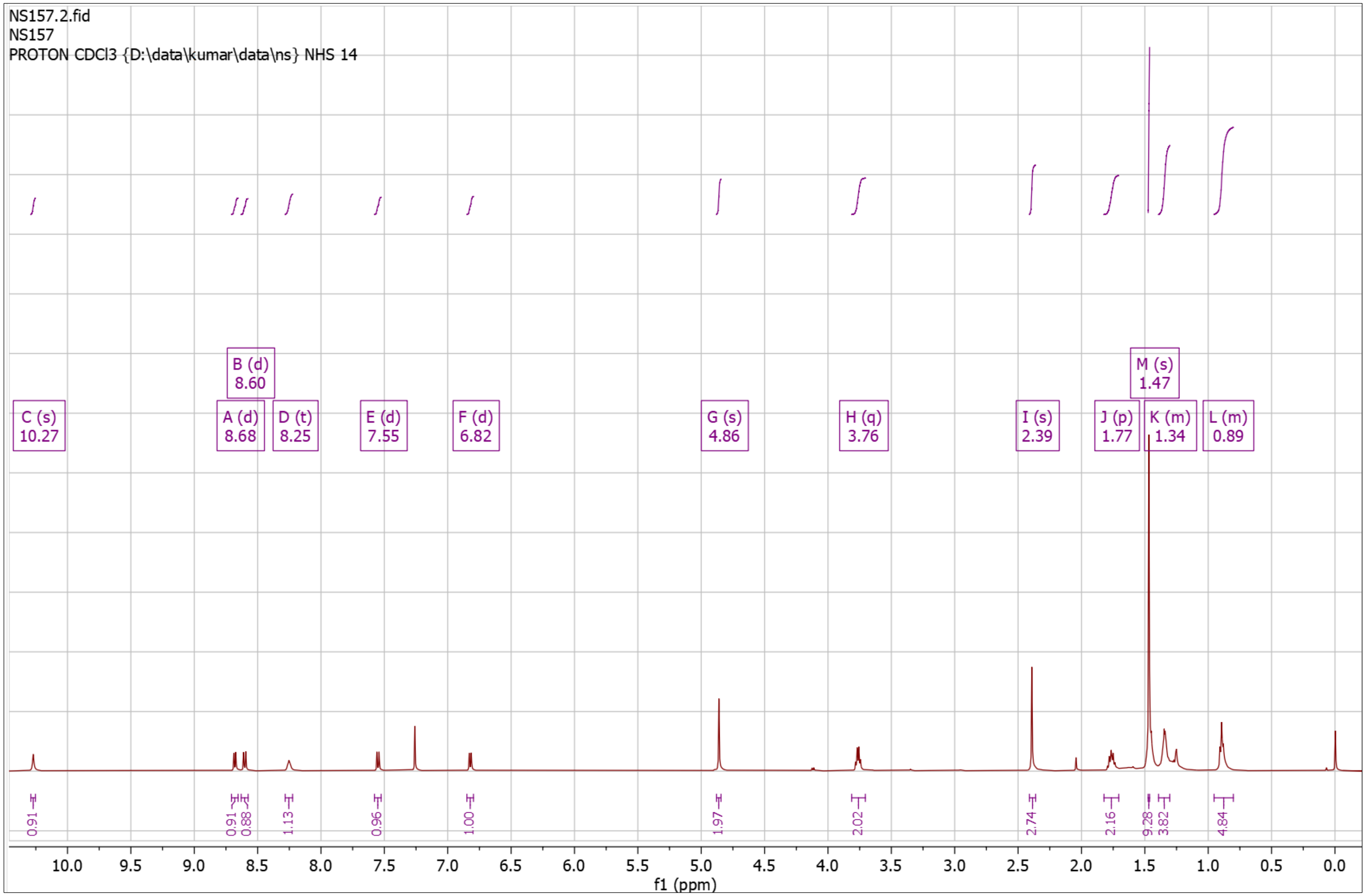


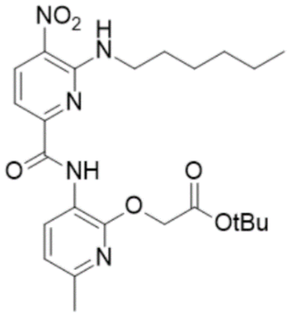

**Supplementary Fig. 53. $^{1}$H-NMR of NS157 Dep**

$^{1}$H NMR (500 MHz, DMSO) δ 0.74 – 0.89 (m, 3H), 1.24 – 1.33 (m, 4H), 1.32 – 1.42 (p, $J$ = 6.9 Hz, 2H), 1.60 – 1.72 (p, $J$ = 7.2 Hz, 2H), 2.29 – 2.34 (s, 3H), 3.61 – 3.76 (q, $J$ = 6.7 Hz, 2H), 4.64 – 4.77 (s, 2H), 6.80 – 6.91 (d, $J$ = 7.9 Hz, 1H), 7.39 – 7.44 (d, $J$ = 8.4 Hz, 1H), 8.45 – 8.53 (d, $J$ = 7.9 Hz, 1H), 8.53 – 8.59 (t, $J$ = 5.8 Hz, 1H), 8.59 – 8.67 (d, $J$ = 8.4 Hz, 1H), 10.35 – 10.50 (s, 1H); HRMS (m/z): [M]+ calcd. for $C_{20}H_{25}N_5O_6$, 432.1878; found, 432.1882.

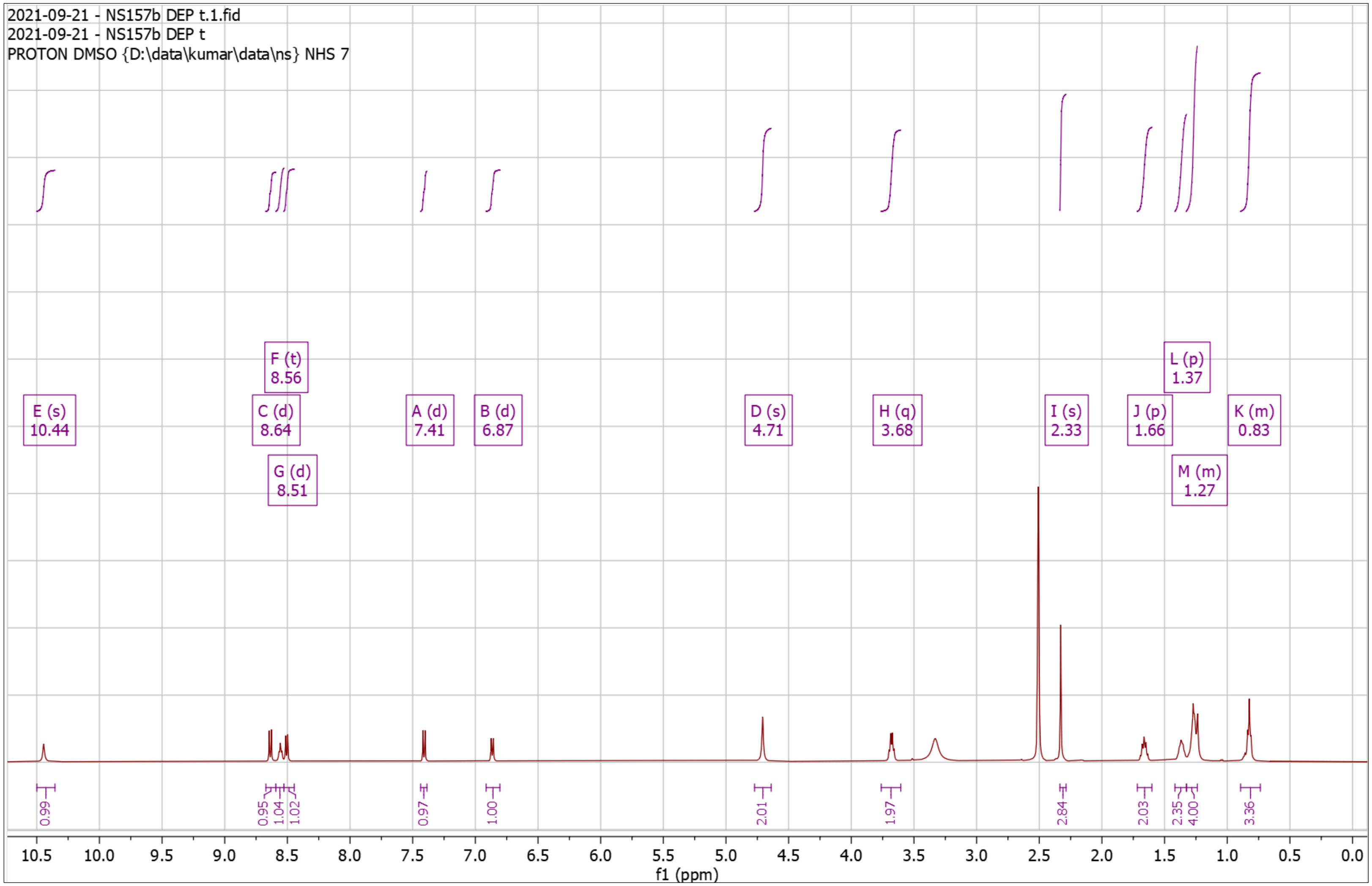


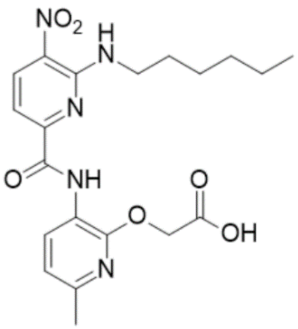

**Supplementary Fig. 54. $^{1}$H-NMR of NS158 Pro**

$^{1}$H NMR (500 MHz, $CDCl_3$) δ 1.49 – 1.51 (s, 9H), 2.38 – 2.41 (s, 3H), 3.93 – 4.02 (d, $J$ = 3.3 Hz, 4H), 4.88 – 4.92 (s, 2H), 6.79 – 6.84 (d, $J$ = 7.9 Hz, 1H), 7.54 – 7.59 (d, $J$ = 8.4 Hz, 1H), 8.41 – 8.45 (t, 1H), 8.57 – 8.62 (d, $J$ = 8.4 Hz, 1H), 8.63 – 8.68 (d, $J$ = 7.9 Hz, 1H), 10.19 – 10.23 (s, 1H); HRMS (m/z): [M]+ calcd. for $C_{20}H_{25}N_5O_7$, 448.1827; found, 448.1827.

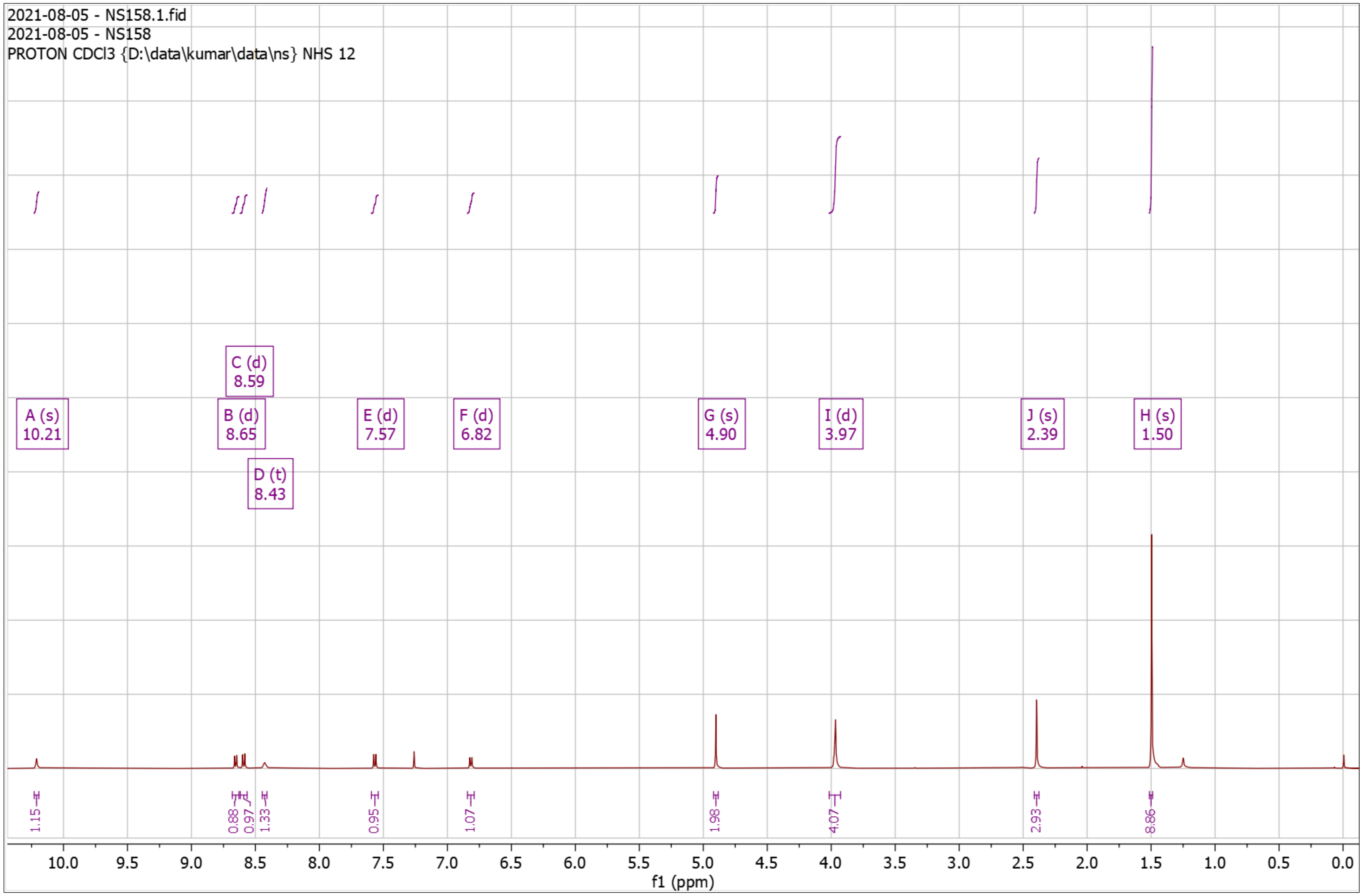


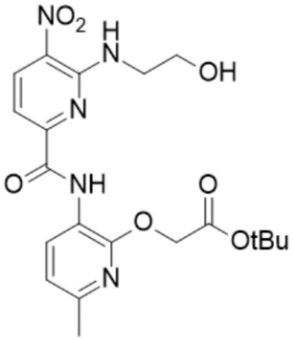

**Supplementary Fig. 55. $^{1}$H-NMR of NS158 Dep**

$^{1}$H NMR (500 MHz, DMSO) δ 2.34 – 2.36 (s, 3H), 3.68 – 3.72 (t, *J* = 5.7 Hz, 2H), 3.72 – 3.78 (t, *J* = 5.5 Hz, 2H), 4.93 – 4.98 (s, 2H), 6.92 – 6.97 (d, *J* = 7.9 Hz, 1H), 7.38 – 7.49 (d, *J* = 8.4 Hz, 1H), 8.47 – 8.53 (d, *J* = 7.9 Hz, 1H), 8.53 – 8.59 (t, *J* = 5.6 Hz, 1H), 8.63 – 8.69 (d, *J* = 8.3 Hz, 1H), 10.31 – 10.33 (s, 1H); HRMS (m/z): [M]+ calcd. for $C_{16}H_{17}N_5O_7$, 392.1201; found, 392.1205.

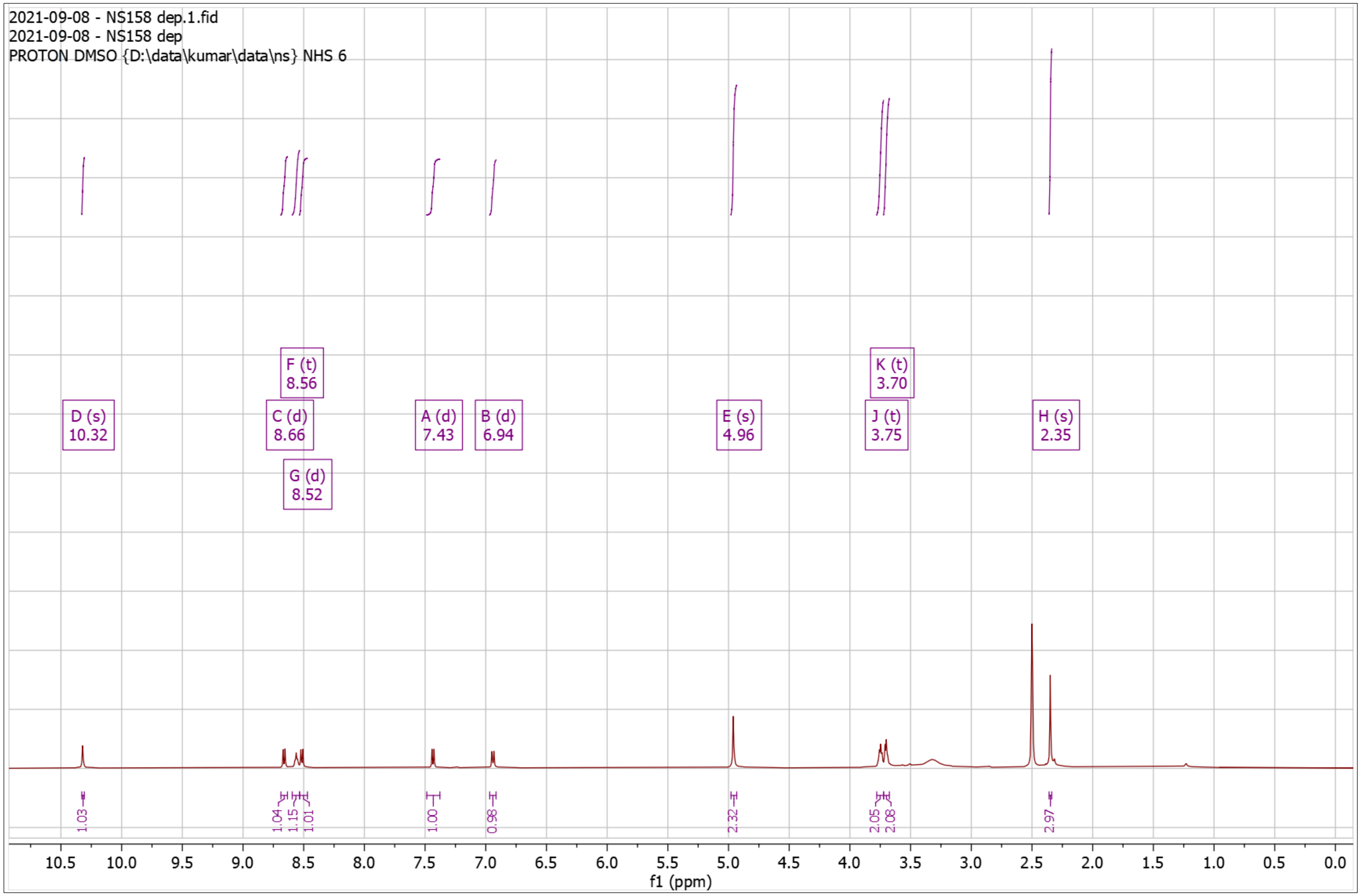


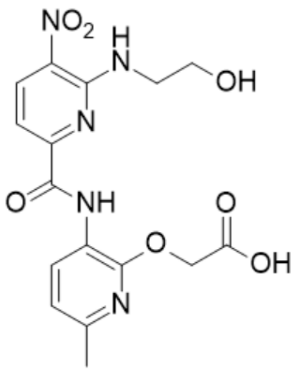

**Supplementary Fig. 56. $^{1}$H-NMR of NS160 Pro**

$^{1}$H NMR (500 MHz, $CDCl_3$) δ 1.43 – 1.48 (s, 9H), 2.38 – 2.41 (s, 3H), 2.96 – 3.03 (t, $J$ = 7.1 Hz, 2H), 3.97 – 4.04 (td, $J$ = 7.1, 5.4 Hz, 2H), 4.67 – 4.71 (s, 2H), 4.80 – 4.94 (s, 1H), 6.74 – 6.80 (d, 2H), 6.80 – 6.85 (d, $J$ = 8.0 Hz, 1H), 7.11 – 7.17 (d, 2H), 7.52 – 7.58 (d, $J$ = 8.5 Hz, 1H), 8.18 – 8.29 (t, 1H), 8.52 – 8.63 (d, $J$ = 8.5 Hz, 1H), 8.65 – 8.70 (d, $J$ = 7.8 Hz, 1H), 10.24 – 10.28 (s, 1H); HRMS (m/z): [M]+ calcd. for $C_{26}H_{29}N_5O_7$, 524.2140; found, 524.2120.

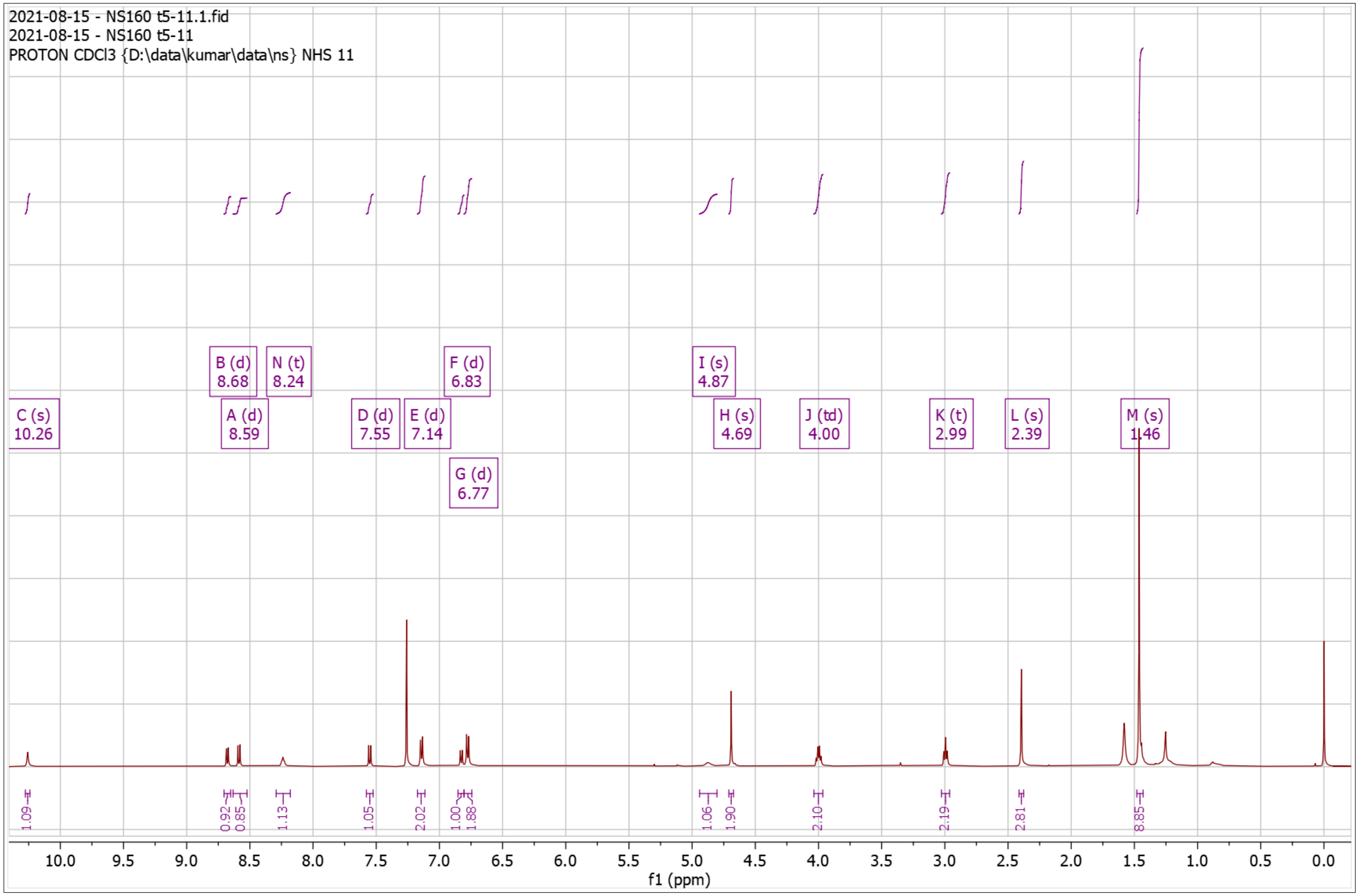


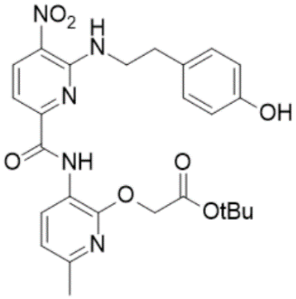

**Supplementary Fig. 57. $^{1}$H-NMR of NS160 Dep**

$^{1}$H NMR (500 MHz, DMSO) δ 2.32 – 2.38 (s, 3H), 2.83 – 2.92 (t, *J* = 7.0 Hz, 2H), 3.83 – 3.95 (q, *J* = 6.7 Hz, 2H), 4.72 – 4.81 (s, 2H), 6.60 – 6.65 (d, *J* = 8.1 Hz, 2H), 6.93 – 6.99 (d, *J* = 7.9 Hz, 1H), 7.02 – 7.08 (d, *J* = 8.0 Hz, 2H), 7.35 – 7.46 (d, *J* = 8.4 Hz, 1H), 8.47 – 8.55 (t, *J* = 5.6 Hz, 1H), 8.55 – 8.59 (d, *J* = 7.9 Hz, 1H), 8.61 – 8.67 (d, *J* = 8.4 Hz, 1H), 9.08 – 9.21 (m, 1H), 10.15 – 10.27 (s, 1H); HRMS (m/z): [M]+ calcd. for $C_{22}H_{21}N_5O_7$, 468.1514; found, 468.1512.

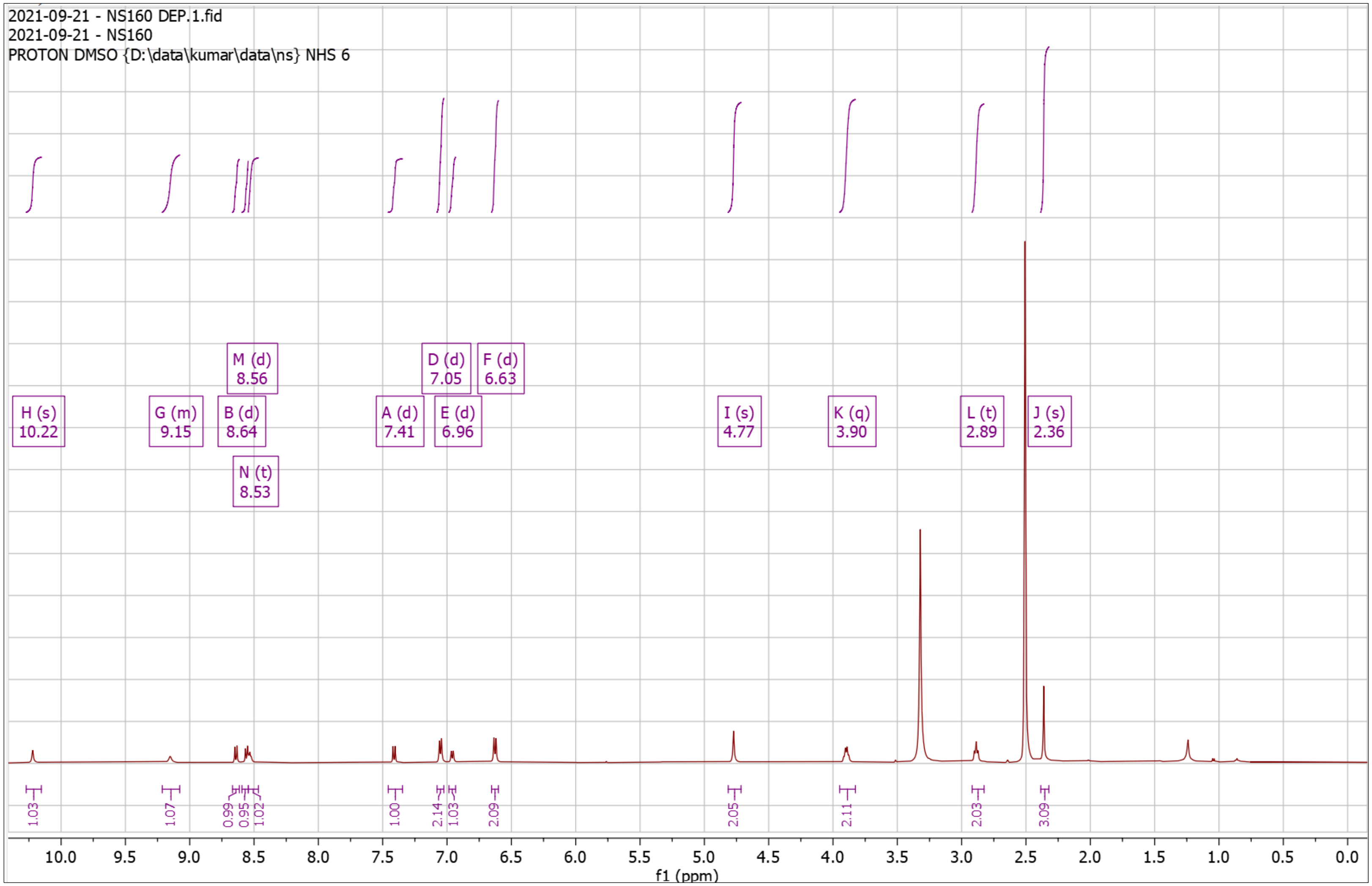


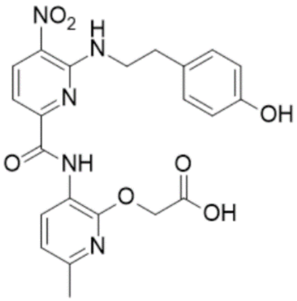

**Supplementary Fig. 58. $^{1}$H-NMR of NS161Pro**

$^{1}$H NMR (500 MHz, $CDCl_3$) δ 1.44 – 1.44 (s, 9H), 2.39 – 2.42 (s, 3H), 2.54 – 2.58 (s, 3H), 4.88 – 4.92 (s, 2H), 5.05 – 5.18 (d, $J$ = 5.3 Hz, 2H), 6.80 – 6.85 (d, $J$ = 7.9 Hz, 1H), 7.61 – 7.66 (d, $J$ = 8.4 Hz, 1H), 8.44 – 8.48 (s, 1H), 8.61 – 8.71 (m, 3H), 9.03 – 9.08 (t, $J$ = 5.4 Hz, 1H), 10.20 – 10.24 (s, 1H); HRMS (m/z): [M]+ calcd. for $C_{24}H_{27}N_7O_6$, 510.2096; found, 510.2095.

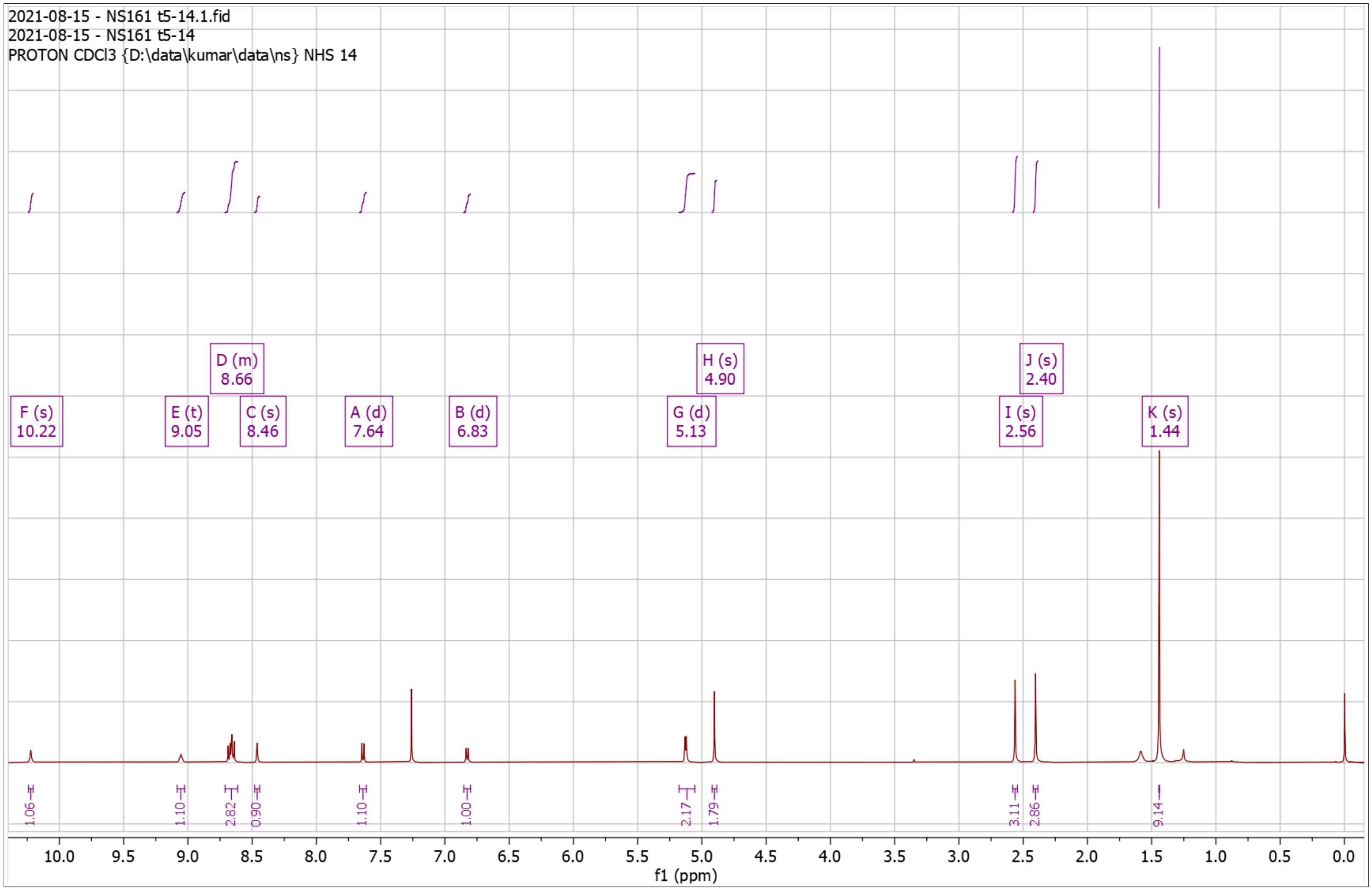


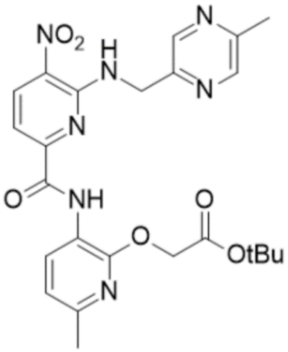

**Supplementary Fig. 59. $^{1}$H-NMR of NS161 Dep**

$^{1}$H NMR (500 MHz, DMSO) δ 2.32 – 2.36 (s, 3H), 2.42 – 2.45 (s, 3H), 4.91 – 4.94 (s, 2H), 4.99 – 5.03 (d, $J$ = 5.7 Hz, 2H), 6.90 – 6.95 (d, $J$ = 7.9 Hz, 1H), 7.46 – 7.51 (d, $J$ = 8.4 Hz, 1H), 8.44 – 8.49 (d, $J$ = 7.8 Hz, 2H), 8.57 – 8.60 (s, 1H), 8.68 – 8.73 (d, $J$ = 8.3 Hz, 1H), 9.21 – 9.27 (t, $J$ = 5.7 Hz, 1H), 10.05 – 10.09 (s, 1H); HRMS (m/z): [M]+ calcd. for $C_{20}H_{19}N_7O_6$, 454.1470; found, 454.1470.

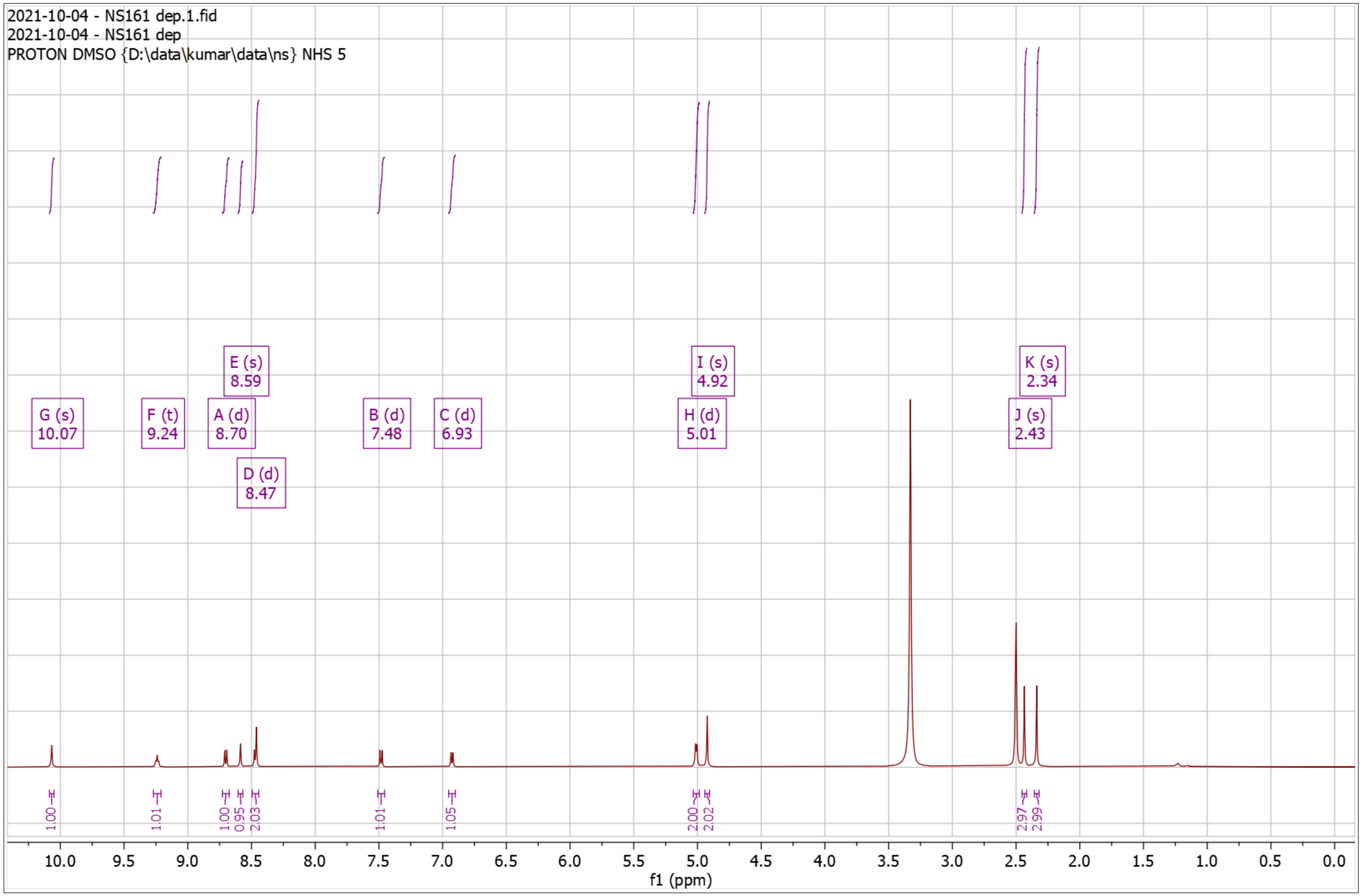


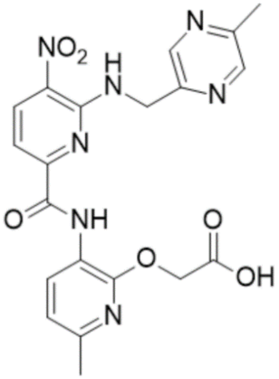

**Supplementary Fig. 60. $^{1}$H-NMR of NS122 Pro**

$^{1}$H NMR (500 MHz, $CDCl_3$) δ 0.99 – 1.33 (m, 5H), 1.40 – 1.49 (s, 9H), 1.63 – 1.91 (m, 6H), 2.34 – 2.39 (s, 3H), 3.41 – 3.49 (t, $J$ = 6.2 Hz, 2H), 4.85 – 4.87 (s, 2H), 4.95 – 5.01 (t, $J$ = 5.7 Hz, 1H), 6.76 – 6.79 (d, $J$ = 7.9 Hz, 1H), 7.57 – 7.62 (d, $J$ = 7.8 Hz, 1H), 7.84 – 7.89 (d, $J$ = 7.9 Hz, 1H), 8.32 – 8.39 (d, $J$ = 8.1 Hz, 1H), 8.39 – 8.44 (d, $J$ = 8.2 Hz, 1H), 8.65 – 8.71 (d, $J$ = 7.9 Hz, 1H), 9.32 – 9.35 (s, 1H), 10.33 – 10.37 (s, 1H). HRMS (m/z): [M]+ calcd. for $C_{31}H_{36}ClN_7O_7$, 654.2438; found, 654.2424.

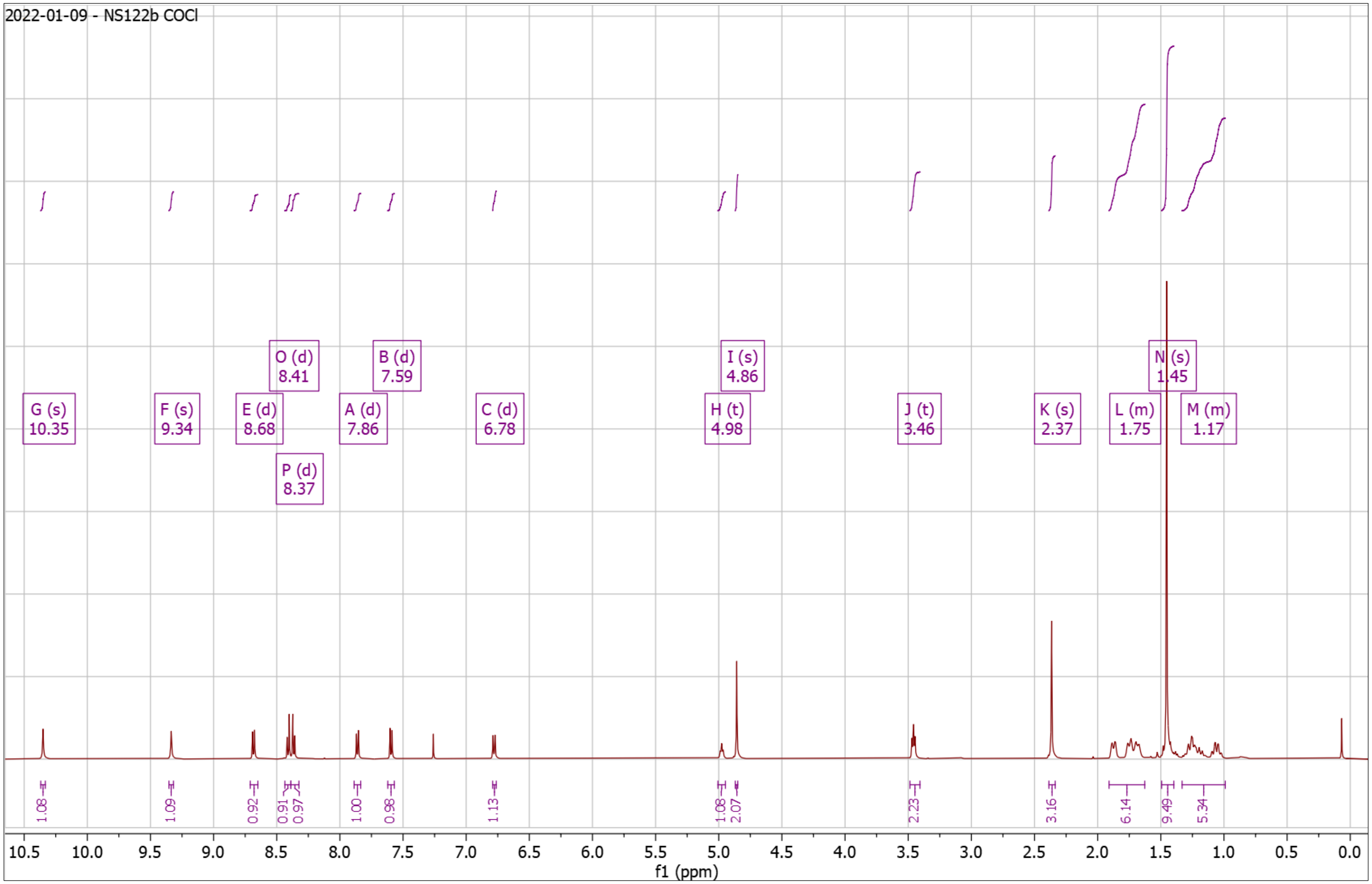


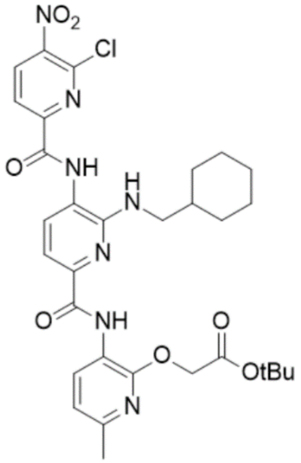

**Supplementary Fig. 61. High Resolution Mass Spectrum for NS122 Pro**

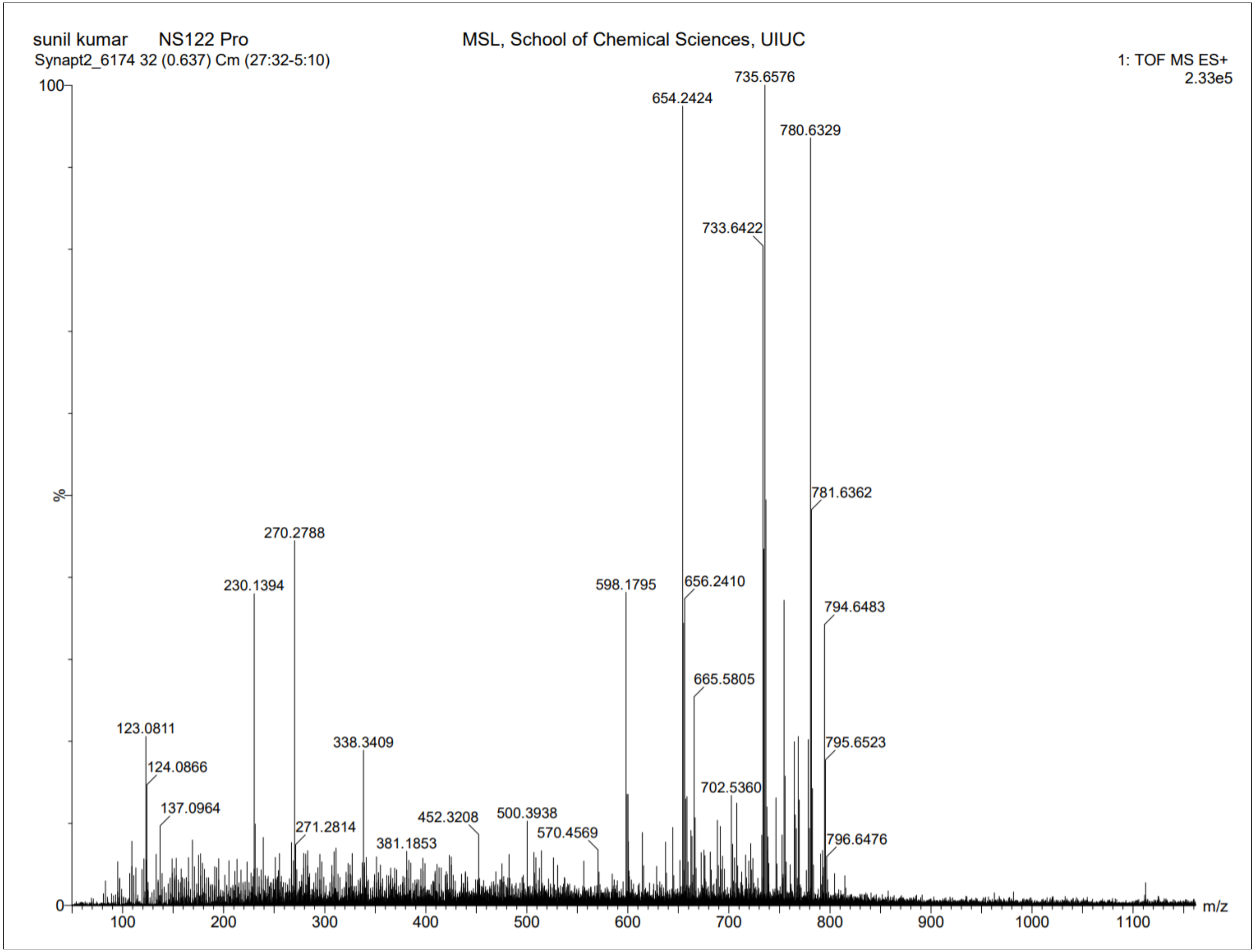

**Supplementary Fig. 62. ¹H-NMR of NS122 Dep**

¹H NMR (500 MHz, DMSO) δ 0.88 – 1.28 (m, 5H), 1.55 – 1.85 (dd, *J* = 85.2, 19.6 Hz, 6H), 2.33 – 2.36 (s, 3H), 4.94 – 4.98 (s, 2H), 6.67 – 6.78 (s, 1H), 6.92 – 6.97 (d, *J* = 7.9 Hz, 1H), 7.34 – 7.39 (d, *J* = 7.6 Hz, 1H), 7.66 – 7.72 (d, *J* = 7.6 Hz, 1H), 8.28 – 8.33 (d, *J* = 8.2 Hz, 1H), 8.60 – 8.68 (d, *J* = 7.8 Hz, 1H), 8.75 – 8.81 (d, *J* = 8.2 Hz, 1H), 10.24 – 10.33 (s, 1H), 10.33 – 10.43 (s, 1H). HRMS (m/z): [M]+ calcd. for $C_{27}H_{28}ClN_7O_7$, 598.1812; found, 598.1802.

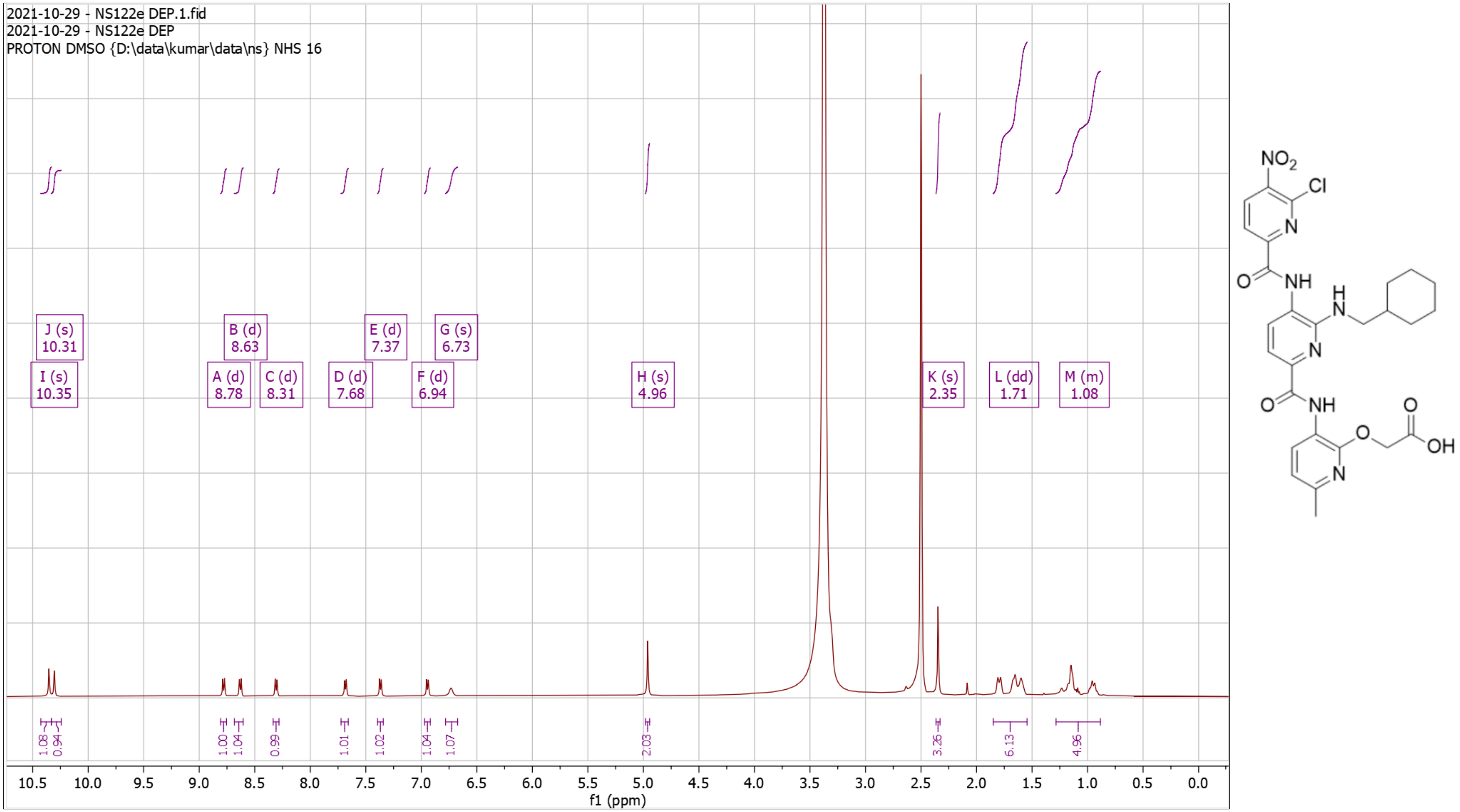

**Supplementary Fig. 63. High Resolution Mass Spectrum for NS122 Dep**

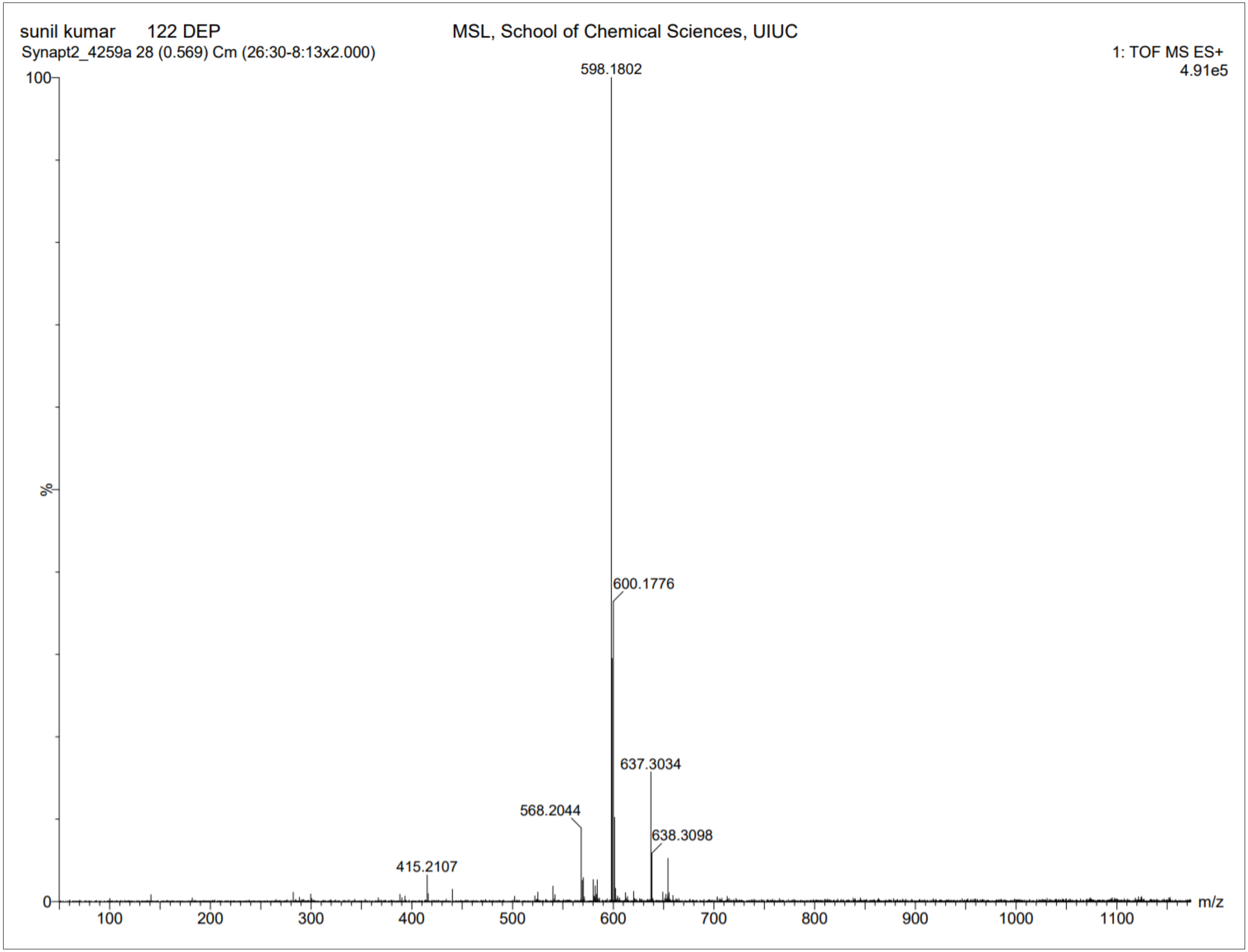

**Supplementary Fig. 64. $^1$H-NMR of NS123 Pro**

$^1$H NMR (500 MHz, CDCl3) δ 1.08 – 1.12 (t, J = 7.5 Hz, 3H), 1.16 – 1.31 (m, 8H), 1.43 – 1.47 (s, 9H), 1.64 – 1.93 (m, 5H), 2.36 – 2.40 (s, 3H), 3.44 – 3.50 (t, J = 6.2 Hz, 2H), 3.62 – 3.69 (m, 2H), 4.81 – 4.85 (t, 1H), 4.85 – 4.88 (s, 2H), 6.78 – 6.83 (d, J = 8.0 Hz, 1H), 7.59 – 7.64 (d, J = 8.4 Hz, 1H), 7.65 – 7.70 (d, J = 7.8 Hz, 1H), 7.92 – 7.98 (d, J = 7.8 Hz, 1H), 8.28 – 8.32 (t, 1H), 8.63 – 8.68 (d, J = 8.3 Hz, 1H), 8.70 – 8.76 (d, J = 7.9 Hz, 1H), 9.38 – 9.41 (s, 1H), 10.36 – 10.40 (s, 1H). HRMS (m/z): [M]+ calcd. for $C_{34}H_{44}N_8O_7$, 677.3406; found, 677.3385.

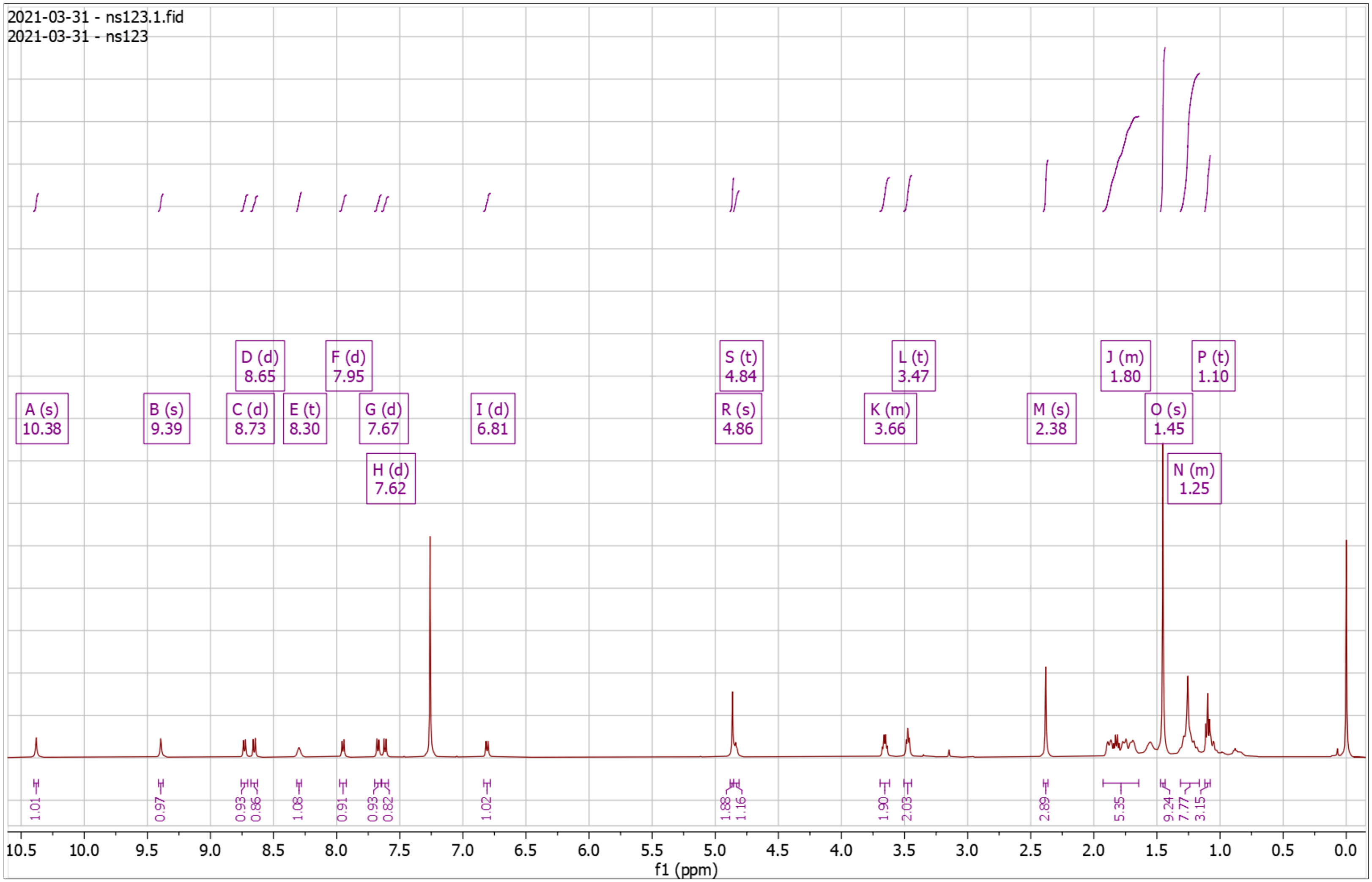


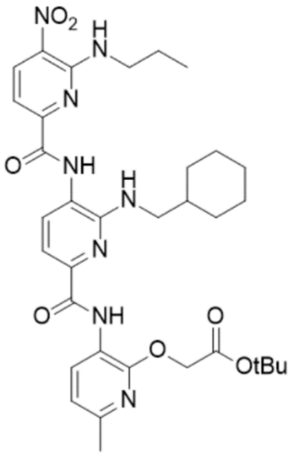

**Supplementary Fig. 65. High Resolution Mass Spectrum for NS123 Pro**

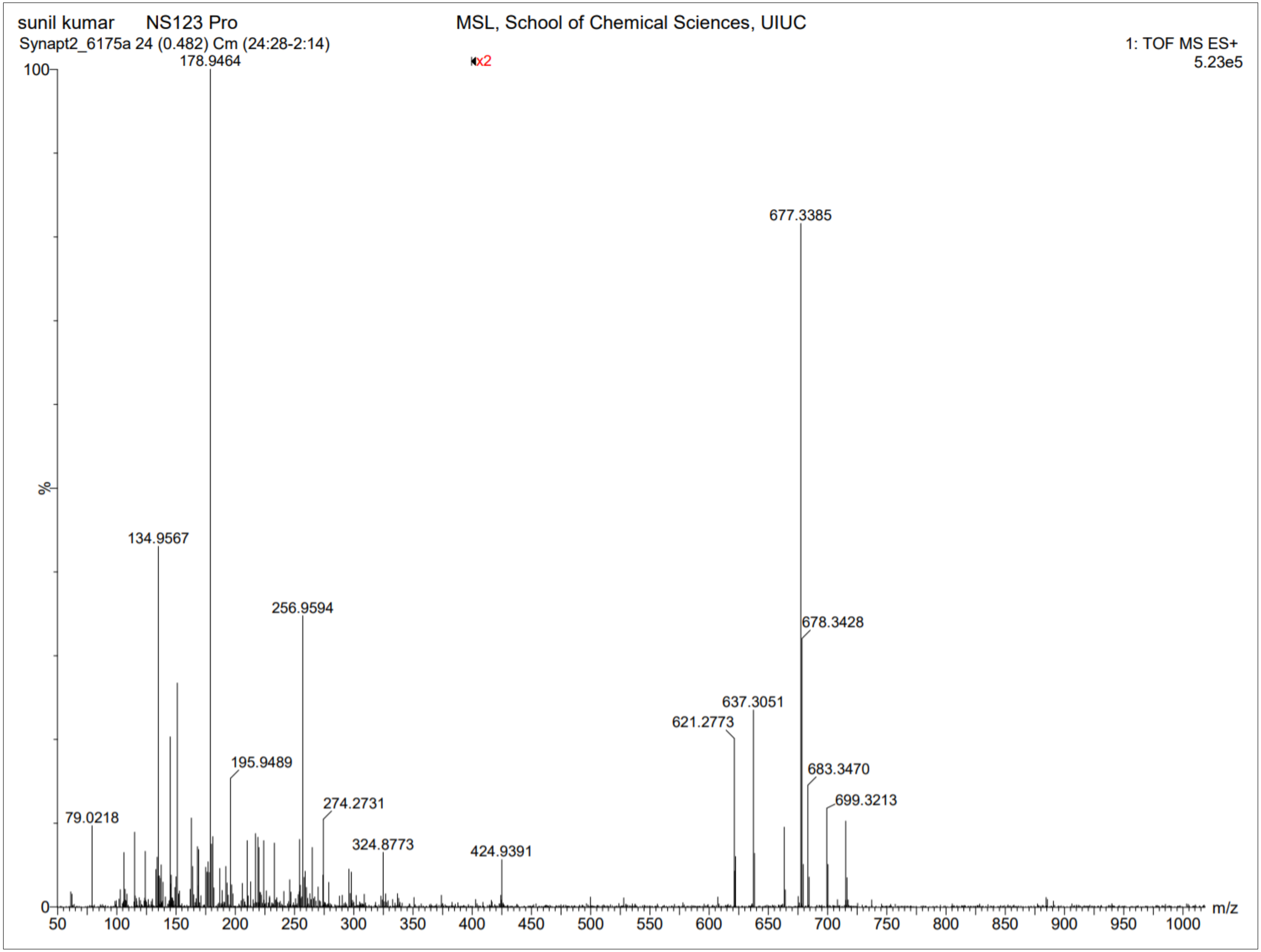

**Supplementary Fig. 66. $^{1}$H-NMR of NS123 Dep**

$^{1}$H NMR (500 MHz, DMSO) δ 0.93 – 0.98 (t, J = 7.4 Hz, 3H), 1.10 – 1.31 (m, 7H), 1.64 – 1.88 (m, 7H), 2.33 – 2.34 (s, 3H), 3.68 – 3.80 (q, J = 6.6 Hz, 2H), 4.88 – 4.95 (s, 2H), 6.67 – 6.77 (t, J = 5.5 Hz, 1H), 6.87 – 6.94 (d, J = 8.0 Hz, 1H), 7.34 – 7.40 (d, J = 8.5 Hz, 1H), 7.40 – 7.46 (d, J = 7.7 Hz, 1H), 7.90 – 7.99 (t, J = 8.1 Hz, 1H), 8.49 – 8.56 (t, J = 5.8 Hz, 1H), 8.58 – 8.68 (dd, J = 9.8, 8.1 Hz, 2H), 9.76 – 9.86 (s, 1H), 10.35 – 10.41 (s, 1H). HRMS (m/z): [M]+ calcd. for $C_{30}H_{36}N_8O_7$, 621.2780; found, 621.2772.

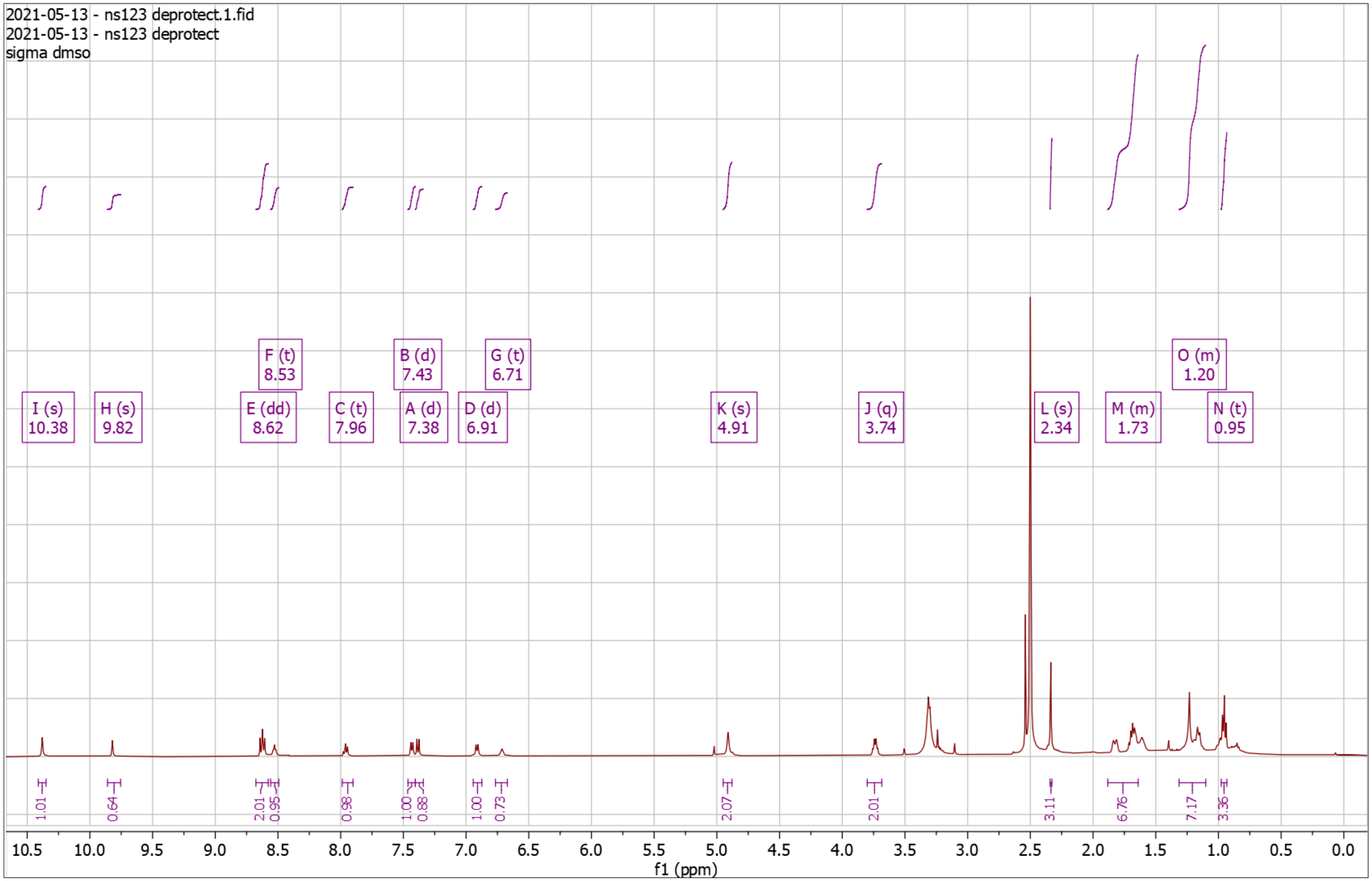


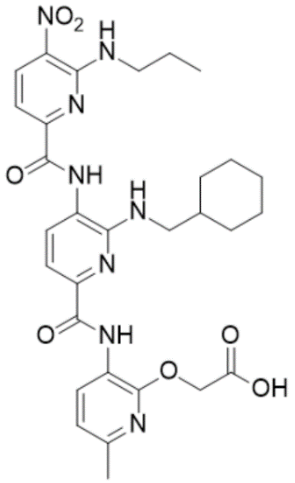

**Supplementary Fig. 70. High Resolution Mass Spectrum for NS123 Dep**

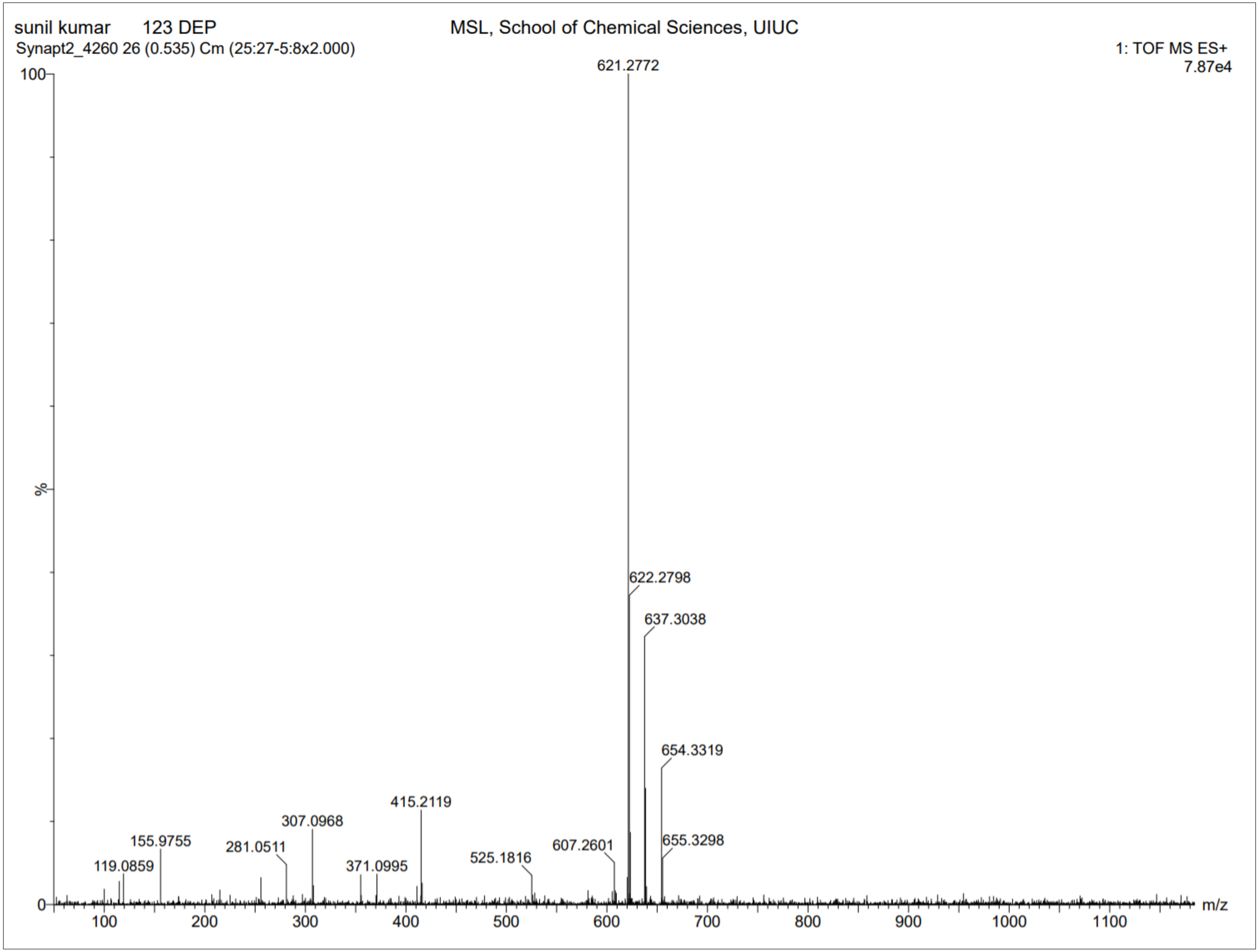

**Supplementary Fig. 71. $^{1}$H-NMR of NS125 Pro**

$^{1}$H NMR (500 MHz, $CDCl_3$) δ 0.84 – 1.20 (m, 5H), 1.20 – 1.22 (s, 9H), 1.43 – 1.46 (s, 9H), 1.64 – 1.92 (m, 6H), 2.37 – 2.40 (s, 3H), 3.38 – 3.43 (q, *J* = 7.6 Hz, 2H), 3.43 – 3.45 (d, *J* = 6.8 Hz, 2H), 3.77 – 3.88 (q, *J* = 6.8 Hz, 2H), 4.81 – 4.86 (t, 1H), 4.86 – 4.90 (s, 2H), 6.78 – 6.83 (d, *J* = 7.9 Hz, 1H), 7.56 – 7.61 (d, *J* = 7.8 Hz, 1H), 7.65 – 7.70 (d, *J* = 8.4 Hz, 1H), 7.80 – 7.86 (d, *J* = 7.8 Hz, 1H), 8.34 – 8.38 (t, *J* = 7.8 Hz, 1H), 8.61 – 8.66 (d, *J* = 8.5 Hz, 1H), 8.72 – 8.77 (d, *J* = 7.9 Hz, 1H), 10.25 – 10.29 (s, 1H), 10.44 – 10.48 (s, 1H). HRMS (m/z): [M]+ calcd. for $C_{38}H_{51}N_9O_9$, 778.3883; found, 778.3856.

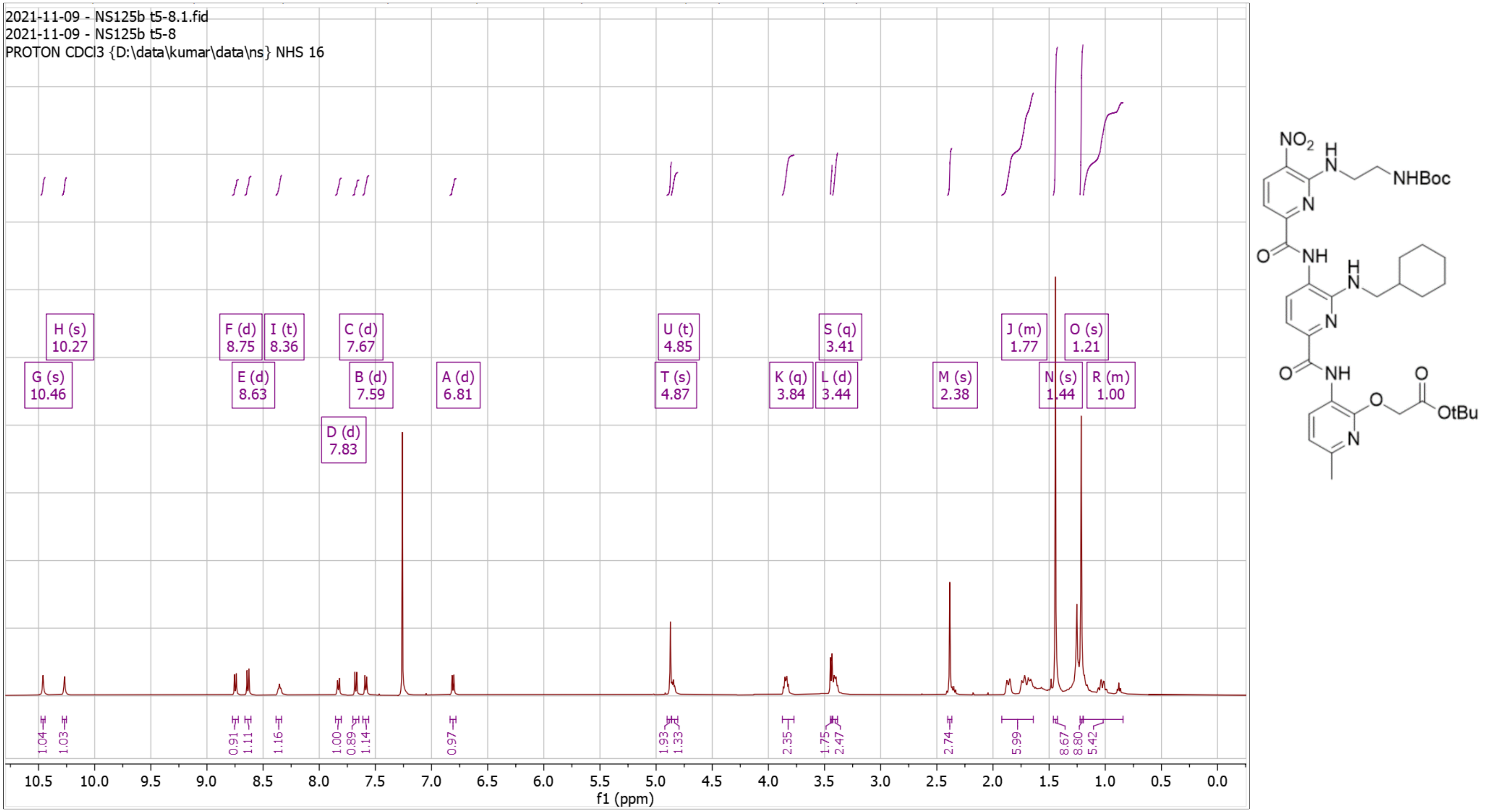

**Supplementary Fig. 72. High Resolution Mass Spectrum for NS125 Pro**

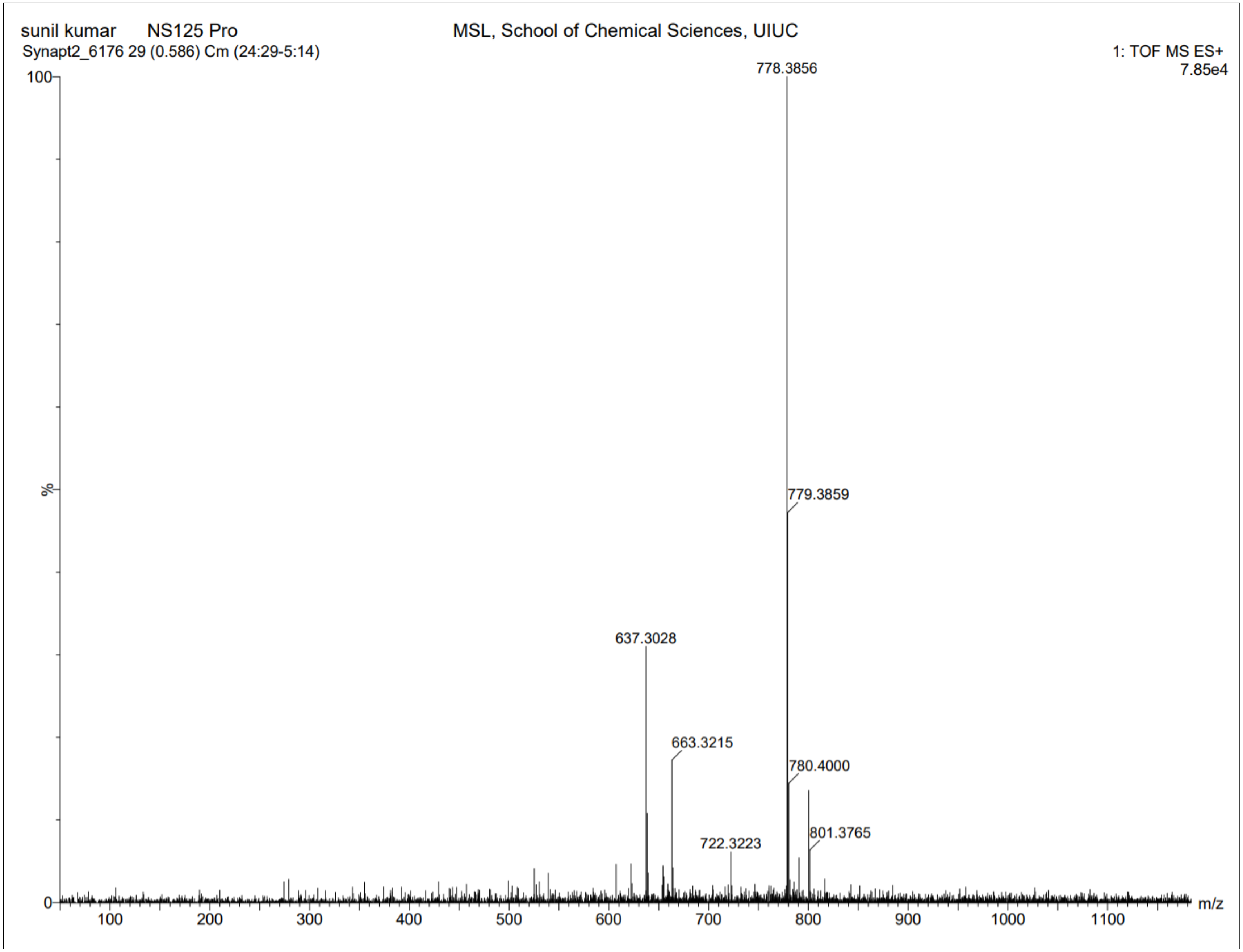

**Supplementary Fig. 73. [1]H-NMR of NS125 Dep**

[1]H NMR (500 MHz, DMSO) δ 0.89 – 1.27 (dq, *J* = 98.6, 11.9 Hz, 6H), 1.56 – 1.87 (m, 7H), 2.32 – 2.35 (s, 3H), 3.07 – 3.13 (t, *J* = 5.7 Hz, 2H), 3.98 – 4.09 (q, *J* = 5.9 Hz, 2H), 4.92 – 4.96 (s, 2H), 6.79 – 6.83 (t, 1H), 6.91 – 6.96 (d, *J* = 7.9 Hz, 1H), 7.38 – 7.43 (d, *J* = 7.7 Hz, 1H), 7.44 – 7.49 (d, *J* = 8.4 Hz, 1H), 7.74 – 7.79 (d, *J* = 7.7 Hz, 1H), 8.59 – 8.73 (m, 3H), 10.33 – 10.36 (s, 1H). HRMS (m/z): [M]+ calcd. for $C_{29}H_{35}N_9O_7$, 622.2732; found, 622.2714.

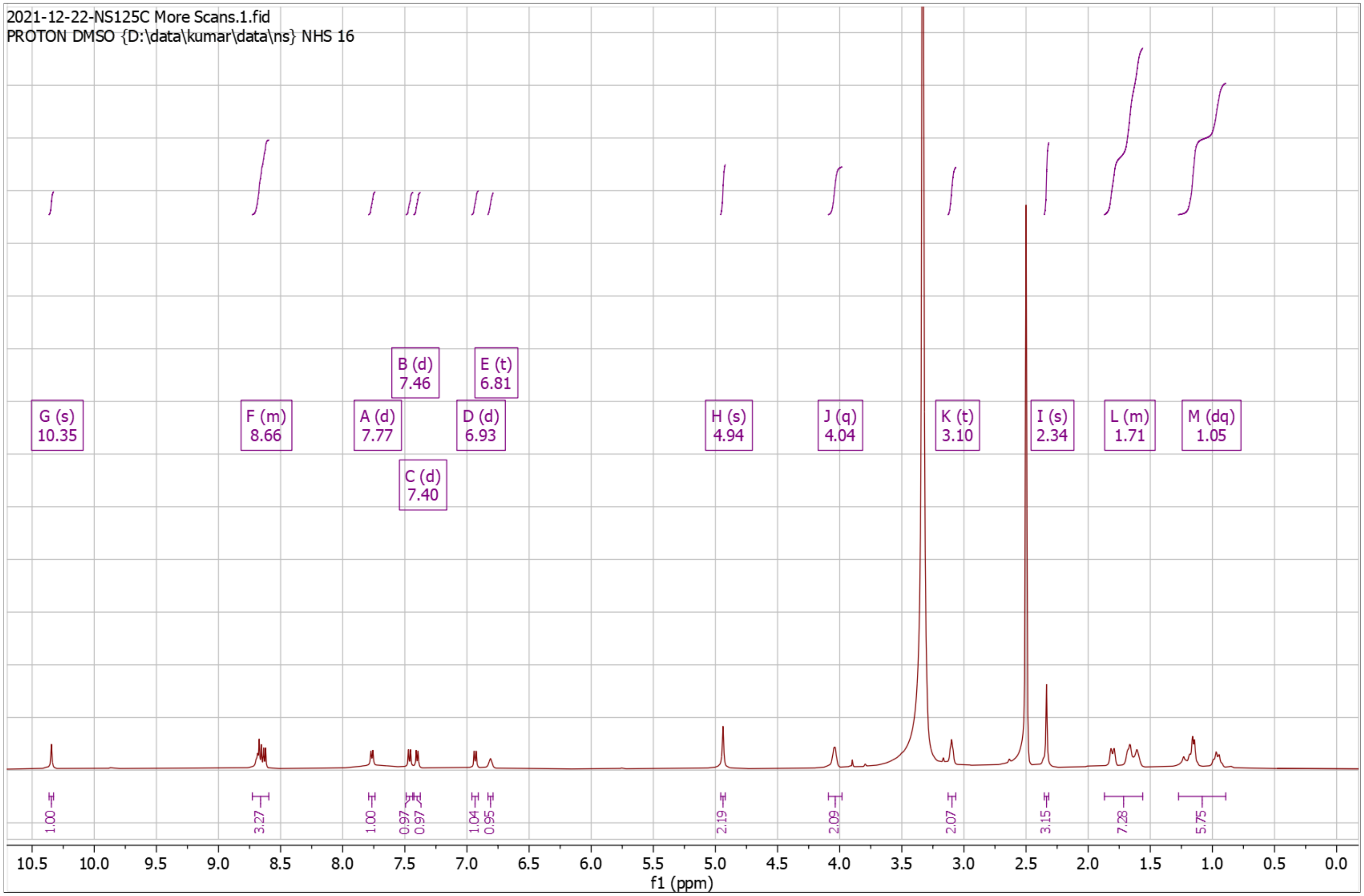


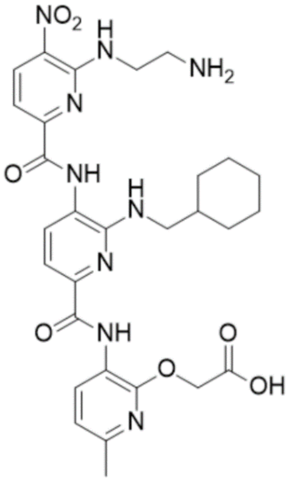

**Supplementary Fig. 74. High Resolution Mass Spectrum for NS125 Dep**

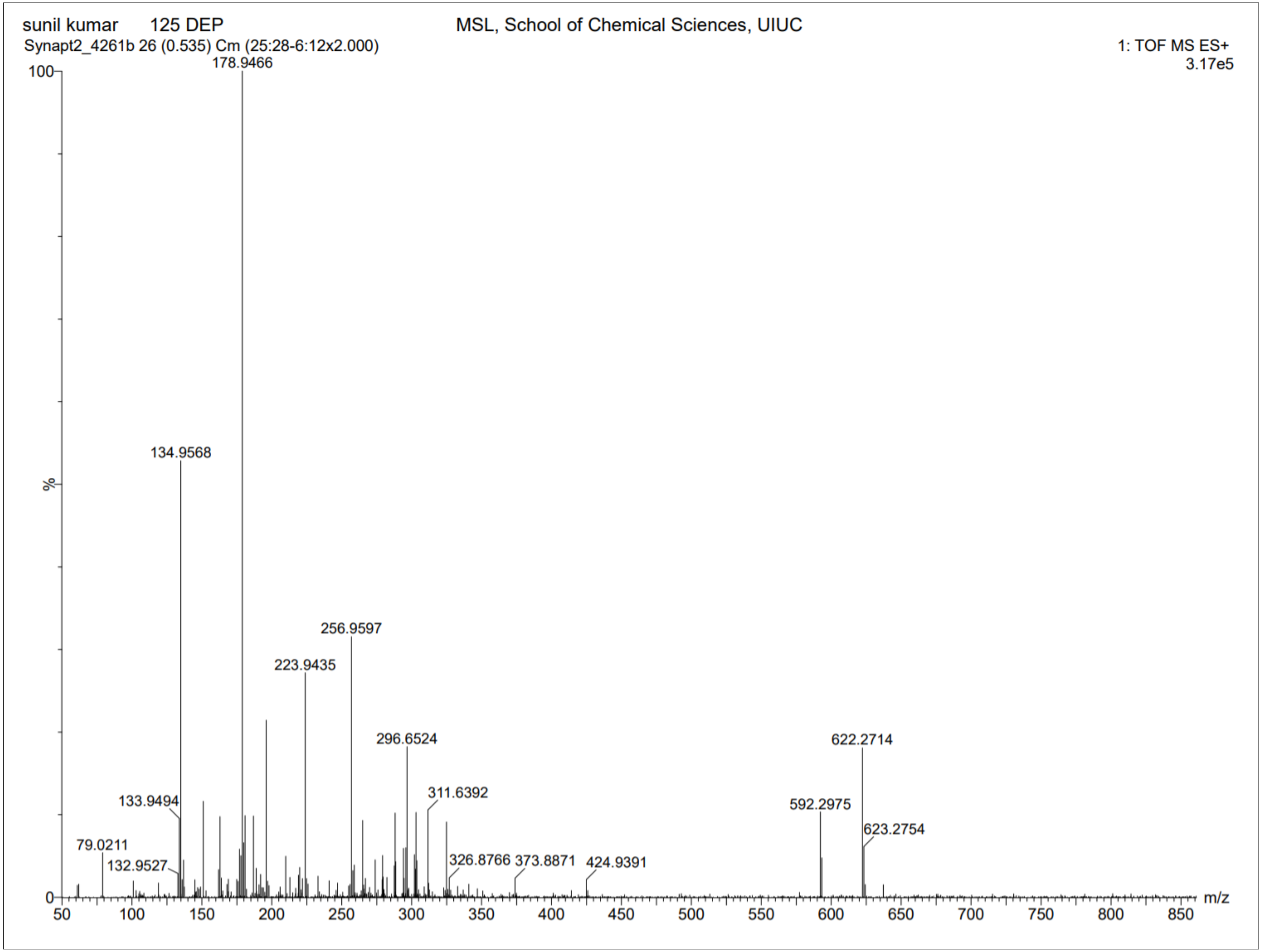

**Supplementary Fig. 75.** $^{1}$**H-NMR of NS126 Pro**

$^{1}$H NMR (500 MHz, $CDCl_3$) δ 1.00 – 1.39 (m, 10H), 1.43 – 1.47 (s, 9H), 1.65 – 1.94 (m, 12H), 2.36 – 2.42 (s, 3H), 3.42 – 3.50 (t, *J* = 6.2 Hz, 2H), 3.50 – 3.59 (t, *J* = 6.1 Hz, 2H), 4.81 – 4.88 (s, 2H), 4.88 – 4.94 (t, *J* = 6.0 Hz, 1H), 6.76 – 6.85 (d, *J* = 8.0 Hz, 1H), 7.54 – 7.63 (d, *J* = 8.4 Hz, 1H), 7.63 – 7.71 (d, *J* = 7.8 Hz, 1H), 7.85 – 7.96 (d, *J* = 7.9 Hz, 1H), 8.30 – 8.41 (t, *J* = 5.7 Hz, 1H), 8.58 – 8.69 (d, *J* = 8.4 Hz, 1H), 8.69 – 8.78 (d, *J* = 7.9 Hz, 1H), 9.28 – 9.43 (s, 1H), 10.29 – 10.47 (s, 1H). HRMS (m/z): [M]+ calcd. for $C_{38}H_{50}N_8O_7$, 731.3875; found, 731.3856.

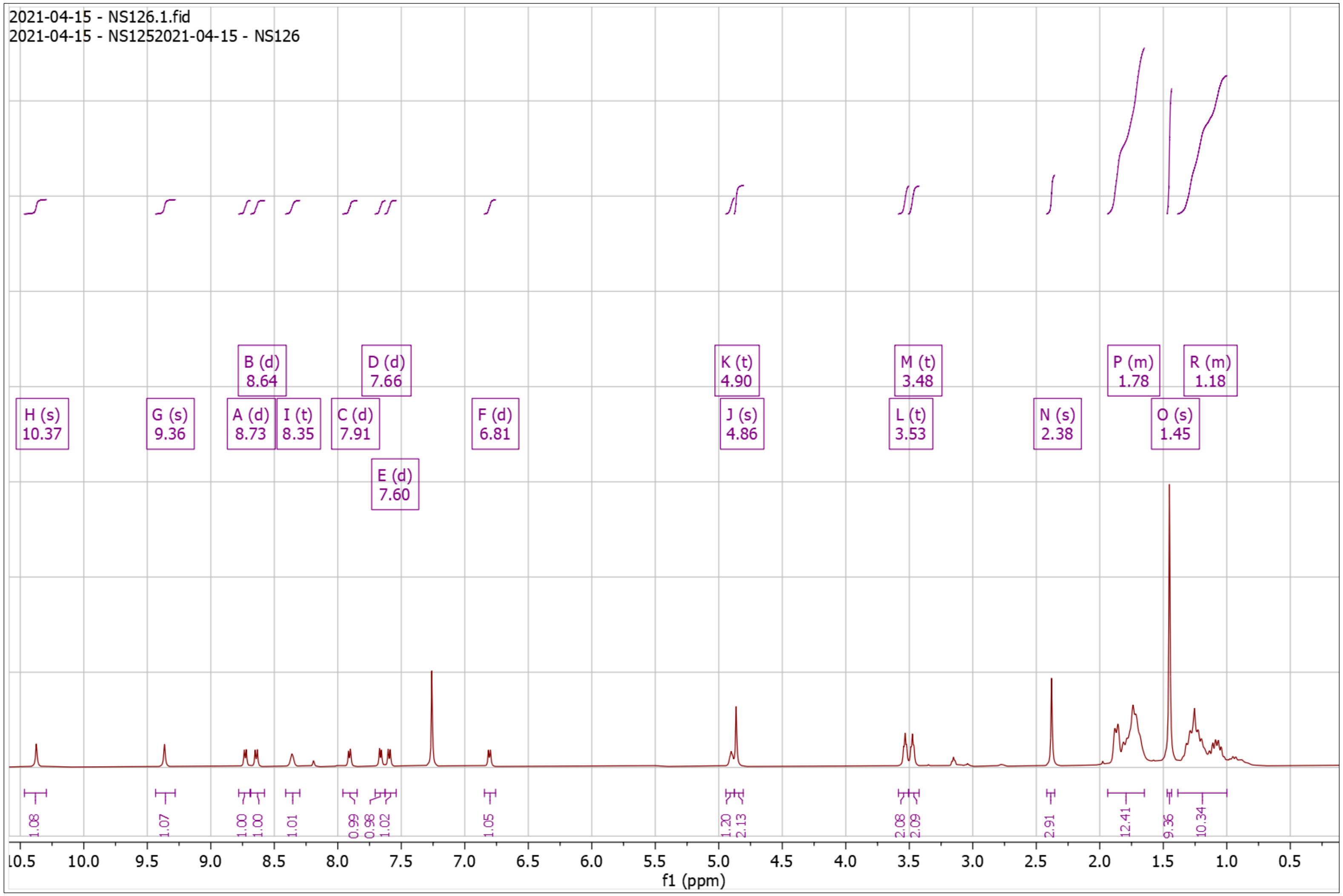


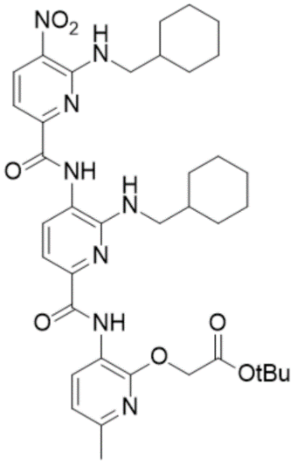

**Supplementary Fig. 76. High Resolution Mass Spectrum for NS126 Pro**

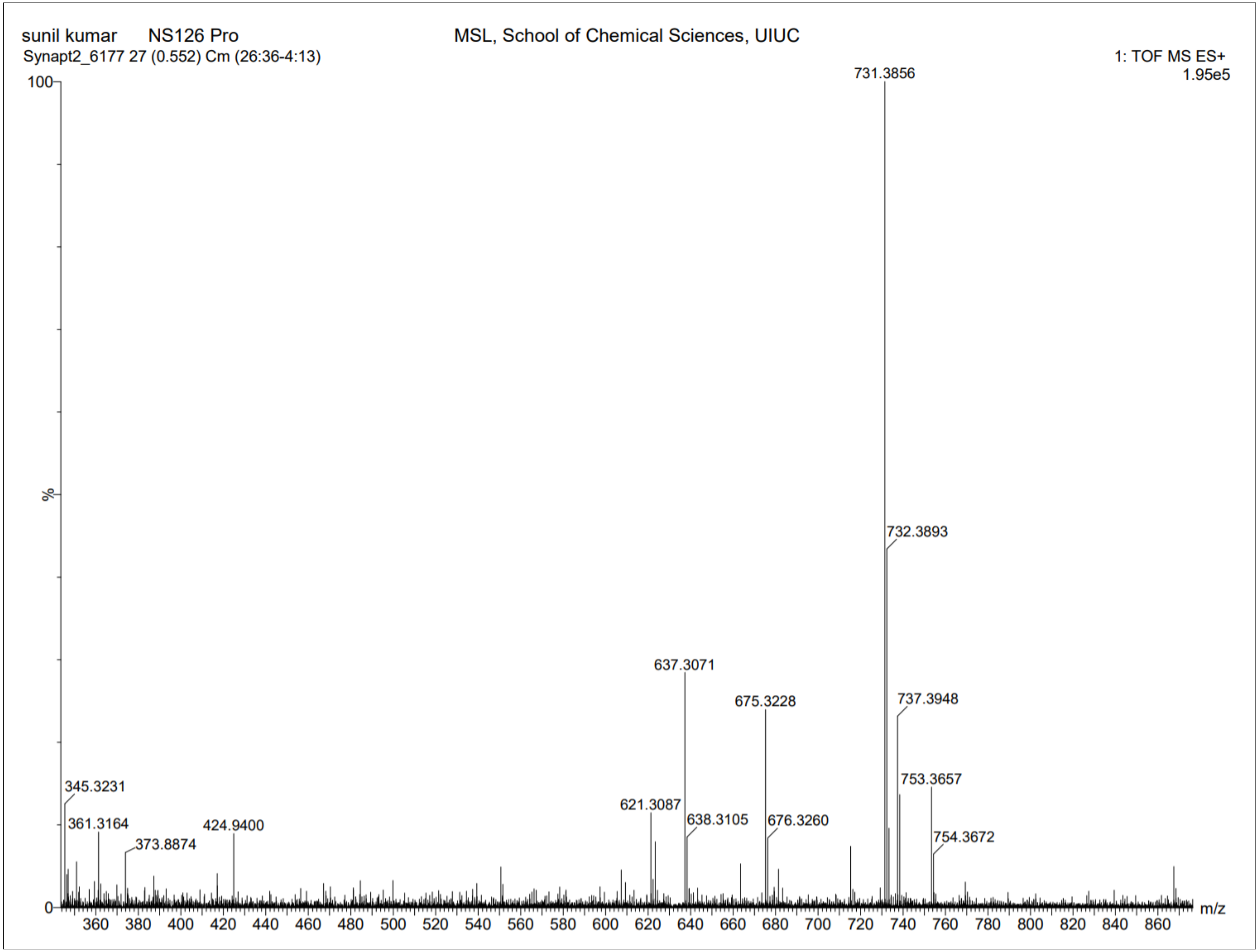

**Supplementary Fig. 77. $^{1}$H-NMR of NS126 Dep**

$^{1}$H NMR (500 MHz, DMSO) δ 0.86 – 1.27 (m, 10H), 1.56 – 1.90 (m, 12H), 2.31 – 2.37 (s, 3H), 3.63 – 3.69 (t, *J* = 6.4 Hz, 2H), 4.92 – 4.96 (s, 2H), 6.72 – 6.80 (t, *J* = 5.8 Hz, 1H), 6.91 – 6.96 (d, *J* = 8.0 Hz, 1H), 7.36 – 7.48 (dd, *J* = 26.2, 8.1 Hz, 2H), 7.92 – 7.97 (d, *J* = 7.8 Hz, 1H), 8.46 – 8.53 (d, *J* = 6.2 Hz, 1H), 8.60 – 8.66 (dd, *J* = 8.2, 4.7 Hz, 2H), 9.78 – 9.81 (s, 1H), 10.35 – 10.38 (s, 1H). HRMS (m/z): [M]+ calcd. for $C_{34}H_{42}N_8O_7$, 675.3249; found, 675.3245.

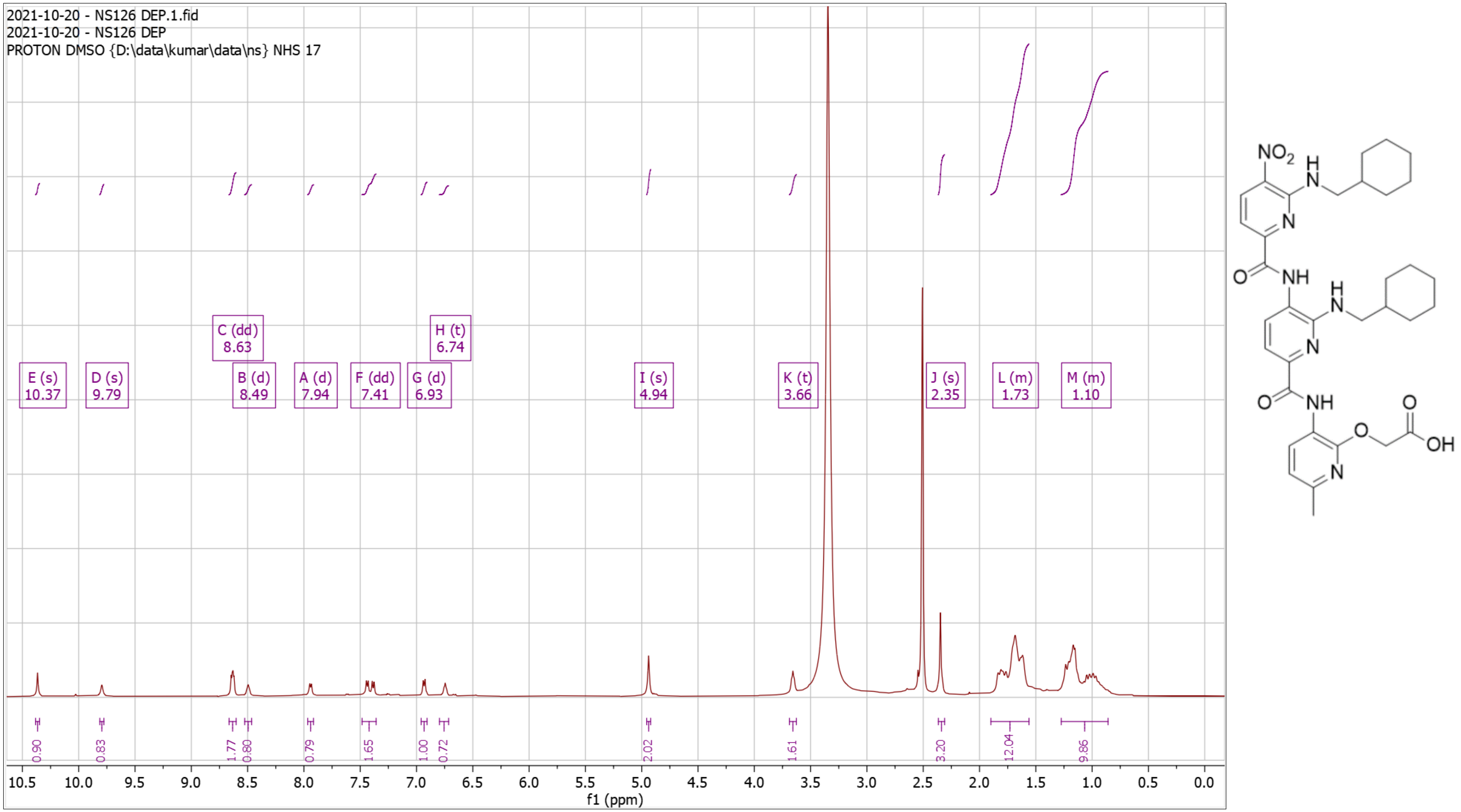

**Supplementary Fig. 78. High Resolution Mass Spectrum for NS126 Dep**

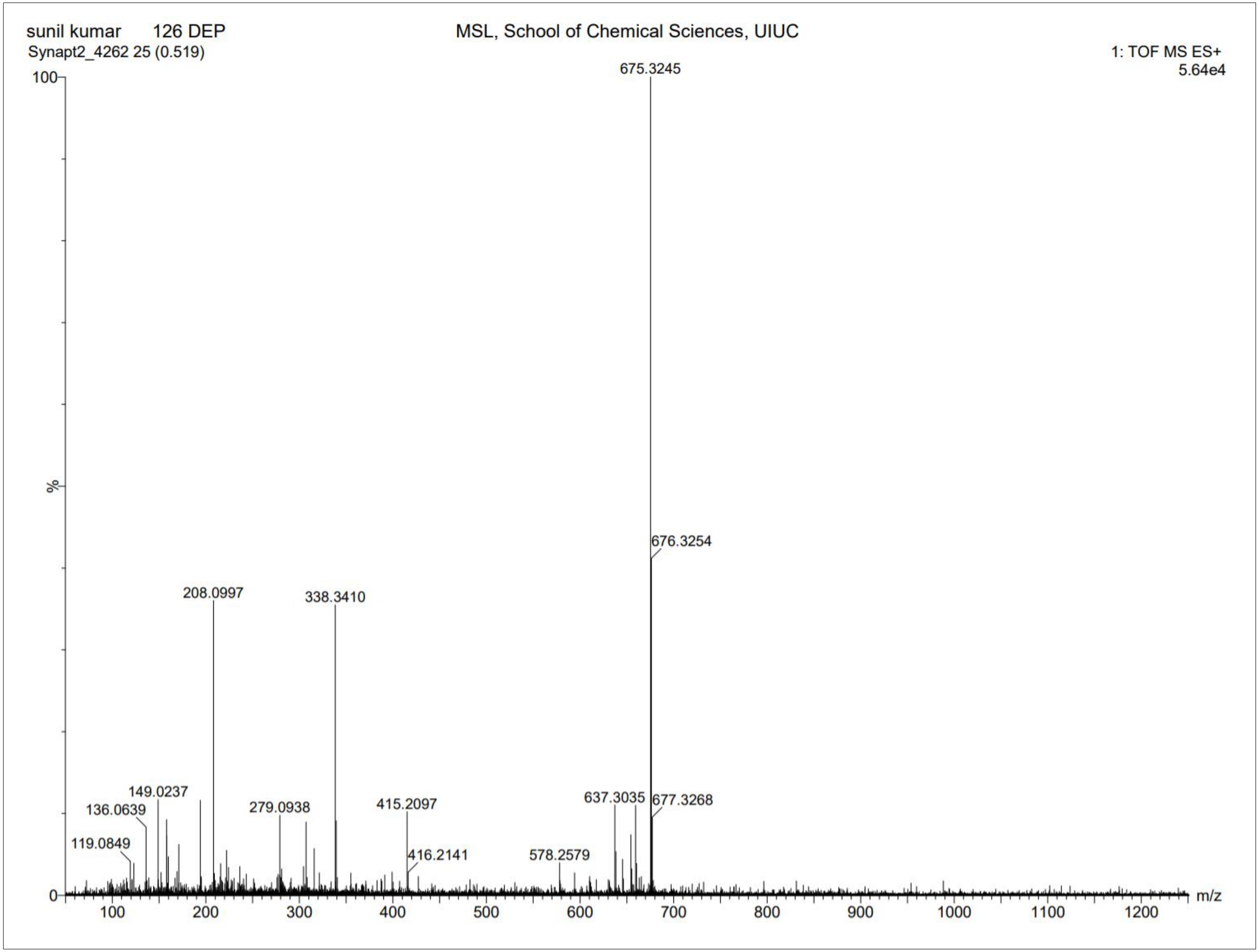

**Supplementary Fig. 79. $^{1}$H-NMR of NS127 Pro**

$^{1}$H NMR (500 MHz, DMSO) δ 0.80 – 1.17 (m, 5H), 1.38 – 1.42 (s, 9H), 1.55 – 1.82 (m, 6H), 2.31 – 2.36 (s, 3H), 2.94 – 3.04 (t, *J* = 7.2 Hz, 2H), 3.30 – 3.31 (t, 3H), 3.99 – 4.07 (q, *J* = 6.8 Hz, 2H), 4.86 – 4.89 (s, 2H), 6.73 – 6.78 (t, *J* = 6.2 Hz, 1H), 6.92 – 6.97 (d, *J* = 7.9 Hz, 1H), 7.13 – 7.33 (m, 5H), 7.35 – 7.42 (d, *J* = 8.4 Hz, 1H), 7.42 – 7.47 (d, *J* = 7.7 Hz, 1H), 7.90 – 7.97 (d, *J* = 7.8 Hz, 1H), 8.52 – 8.58 (t, *J* = 5.8 Hz, 1H), 8.60 – 8.66 (dd, *J* = 8.1, 3.4 Hz, 2H), 9.82 – 9.87 (s, 1H), 10.33 – 10.37 (s, 1H). HRMS (m/z): [M]+ calcd. for $C_{39}H_{46}N_8O_7$, 739.3562; found, 739.3551.

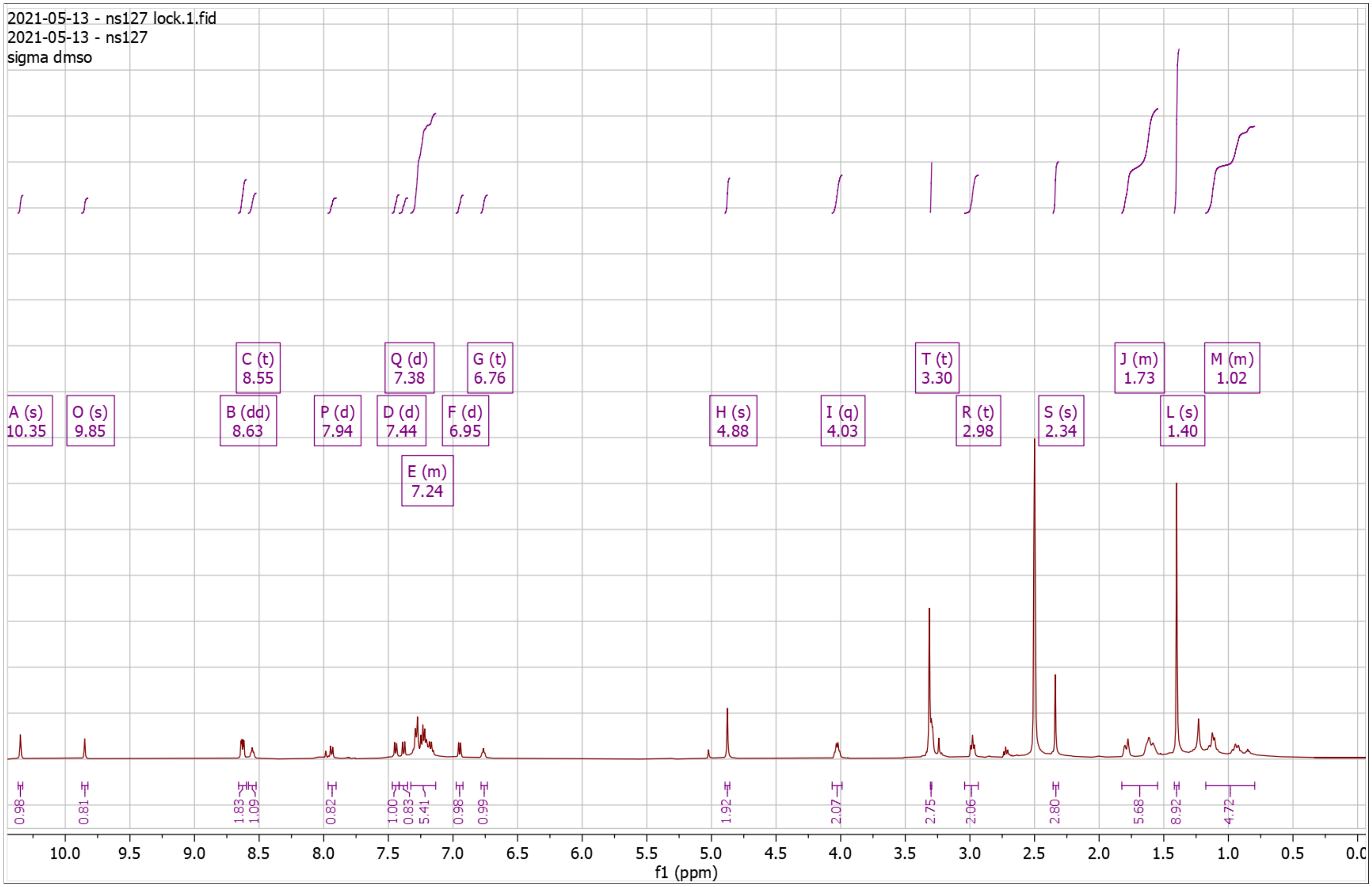


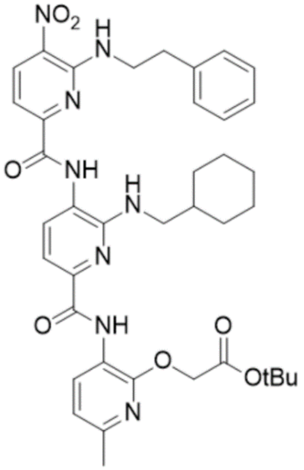

**Supplementary Fig. 80. High Resolution Mass Spectrum for NS127 Pro**

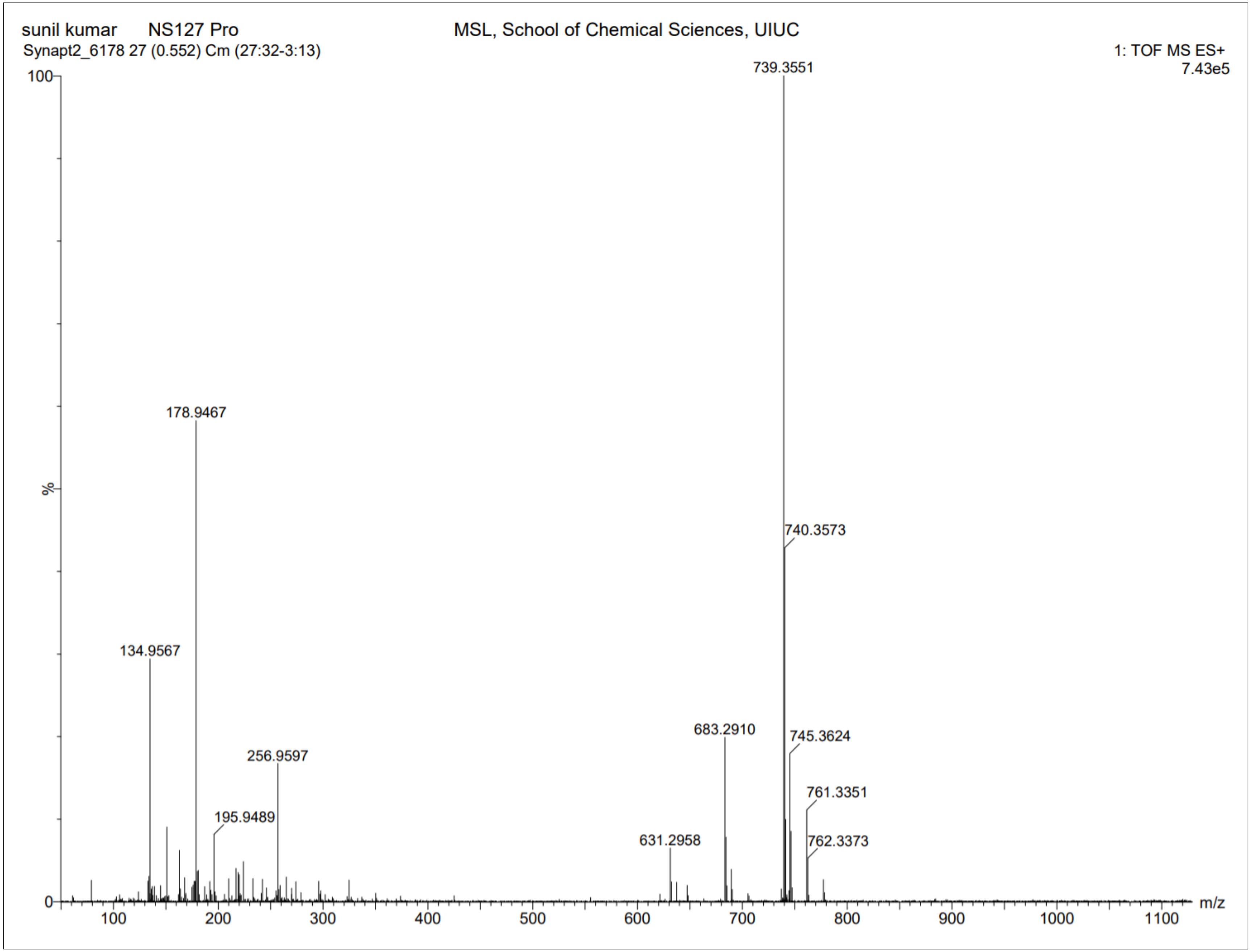

**Supplementary Fig. 81. $^{1}$H-NMR of NS127 Dep**

$^{1}$H NMR (500 MHz, DMSO) δ 0.89 – 1.16 (m, 5H), 1.52 – 1.86 (m, 6H), 2.31 – 2.40 (s, 3H), 2.92 – 3.04 (t, *J* = 7.2 Hz, 2H), 3.23 – 3.33 (t, 2H), 3.99 – 4.07 (q, *J* = 6.7 Hz, 2H), 4.91 – 5.00 (s, 2H), 6.69 – 6.79 (t, *J* = 5.7 Hz, 1H), 6.88 – 6.98 (d, *J* = 7.9 Hz, 1H), 7.12 – 7.31 (m, 5H), 7.35 – 7.41 (d, *J* = 8.4 Hz, 1H), 7.41 – 7.48 (d, *J* = 7.7 Hz, 1H), 7.86 – 7.99 (d, *J* = 7.7 Hz, 1H), 8.48 – 8.58 (t, *J* = 5.8 Hz, 1H), 8.59 – 8.66 (dd, *J* = 8.2, 4.5 Hz, 2H), 9.80 – 9.88 (s, 1H), 10.33 – 10.39 (s, 1H). HRMS (m/z): [M]+ calcd. for $C_{35}H_{38}N_8O_7$, 683.2936; found, 683.2925.

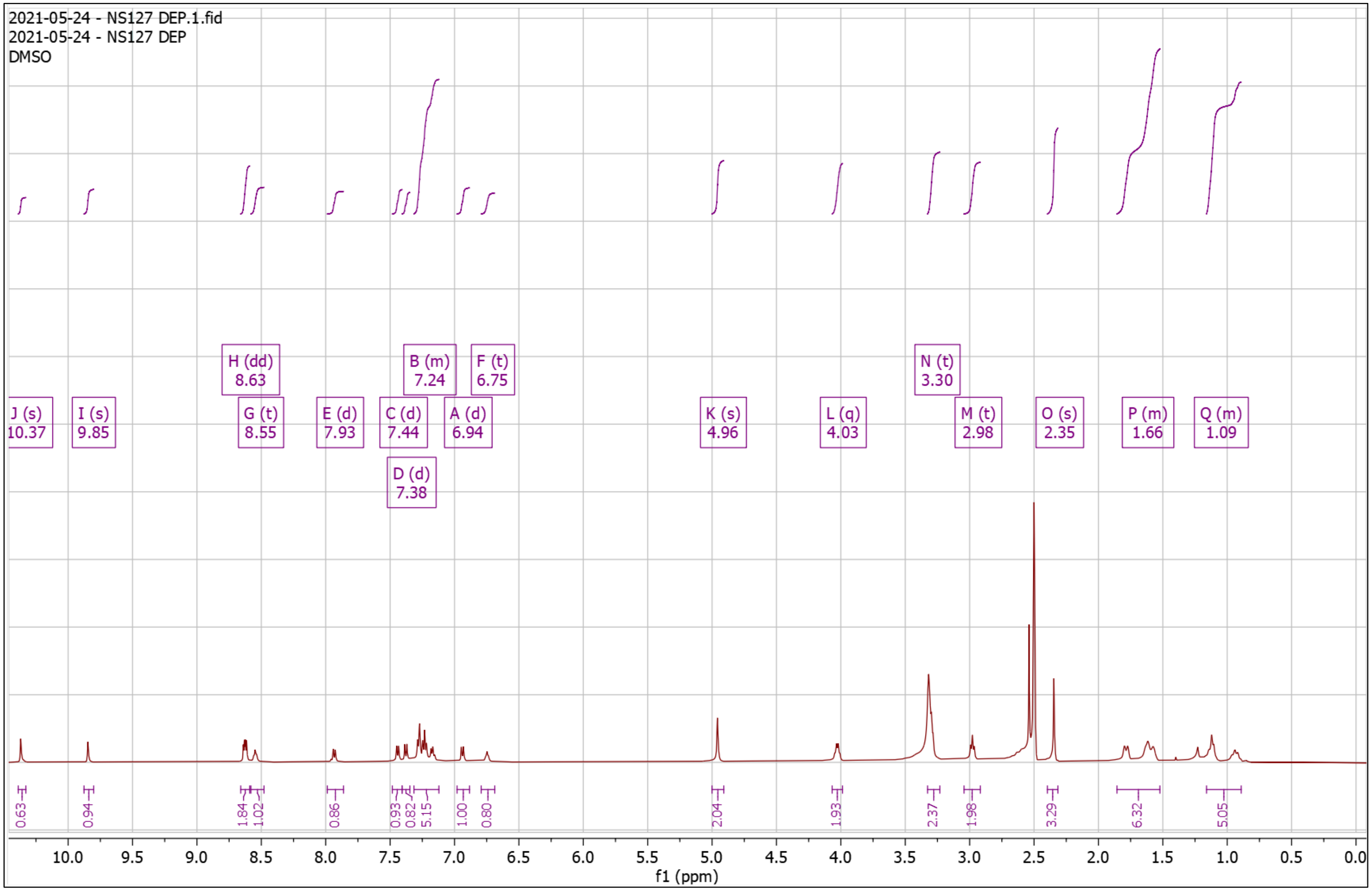


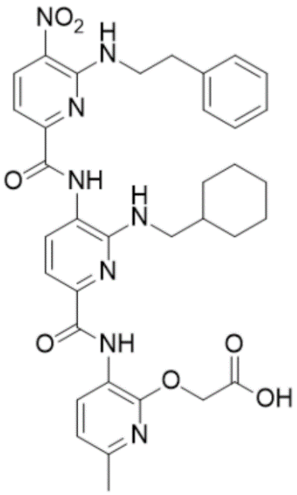

**Supplementary Fig. 82. High Resolution Mass Spectrum for NS127 Dep**

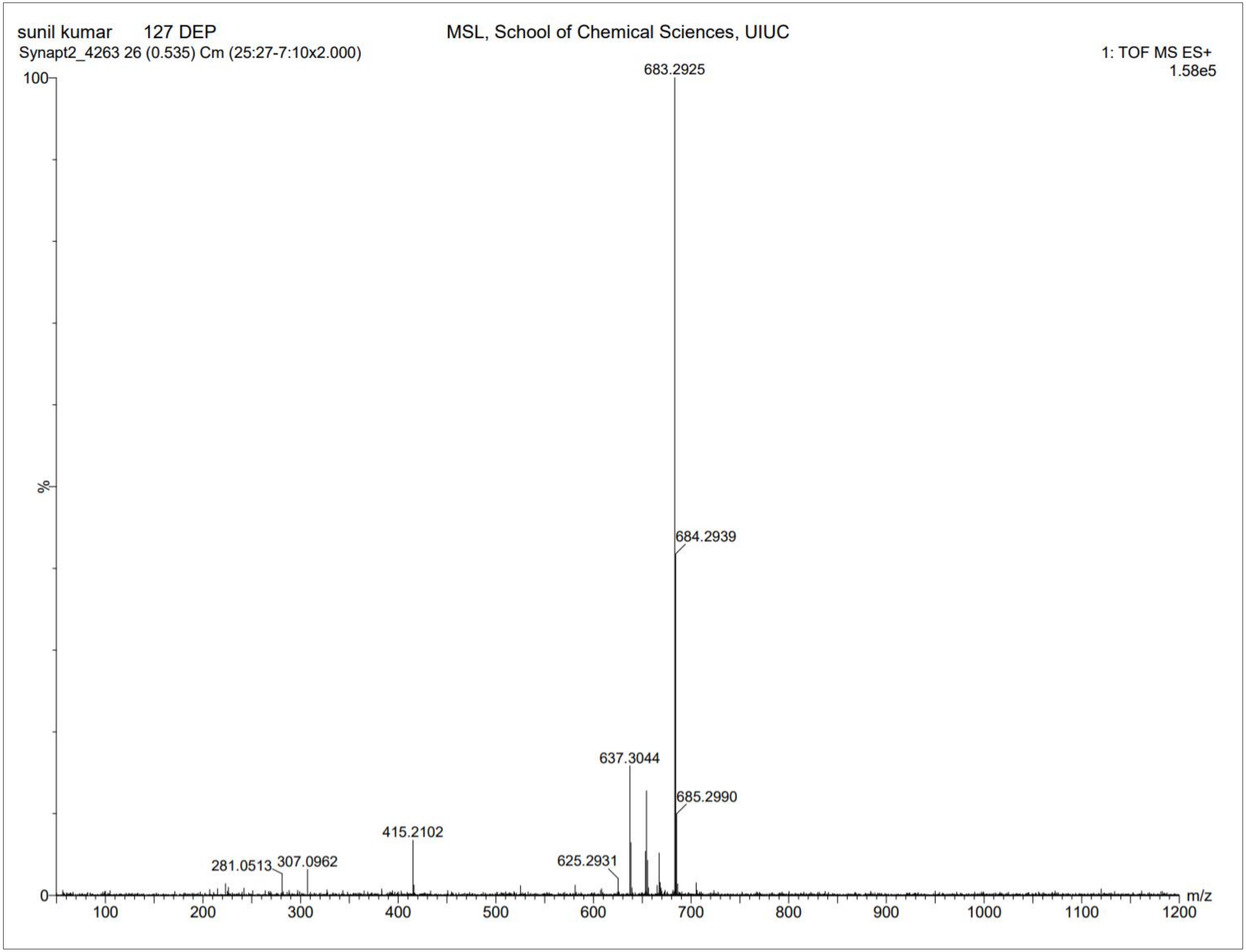

**Supplementary Fig. 83. $^{1}$H-NMR of NS129 Pro**

$^{1}$H NMR (500 MHz, DMSO) δ 0.90 – 1.17 (dd, $J$ = 81.8, 10.1 Hz, 5H), 1.29 – 1.49 (s, 9H), 1.50 – 1.87 (m, 6H), 2.30 – 2.37 (s, 3H), 3.27 – 3.40 (d, $J$ = 6.7 Hz, 2H), 4.83 – 4.88 (s, 2H), 4.88 – 4.99 (s, 2H), 6.28 – 6.36 (s, 1H), 6.91 – 6.97 (d, $J$ = 7.9 Hz, 1H), 7.38 – 7.49 (t, $J$ = 7.5 Hz, 2H), 7.66 – 7.73 (s, 1H), 8.04 – 8.17 (d, $J$ = 7.7 Hz, 1H), 8.59 – 8.68 (m, 2H), 8.96 – 9.04 (s, 1H), 10.17 – 10.29 (s, 1H), 10.31 – 10.45 (s, 1H). HRMS (m/z): [M]+ calcd. for $C_{35}H_{42}N_{10}O_{7}$, 715.3311; found, 715.3275.

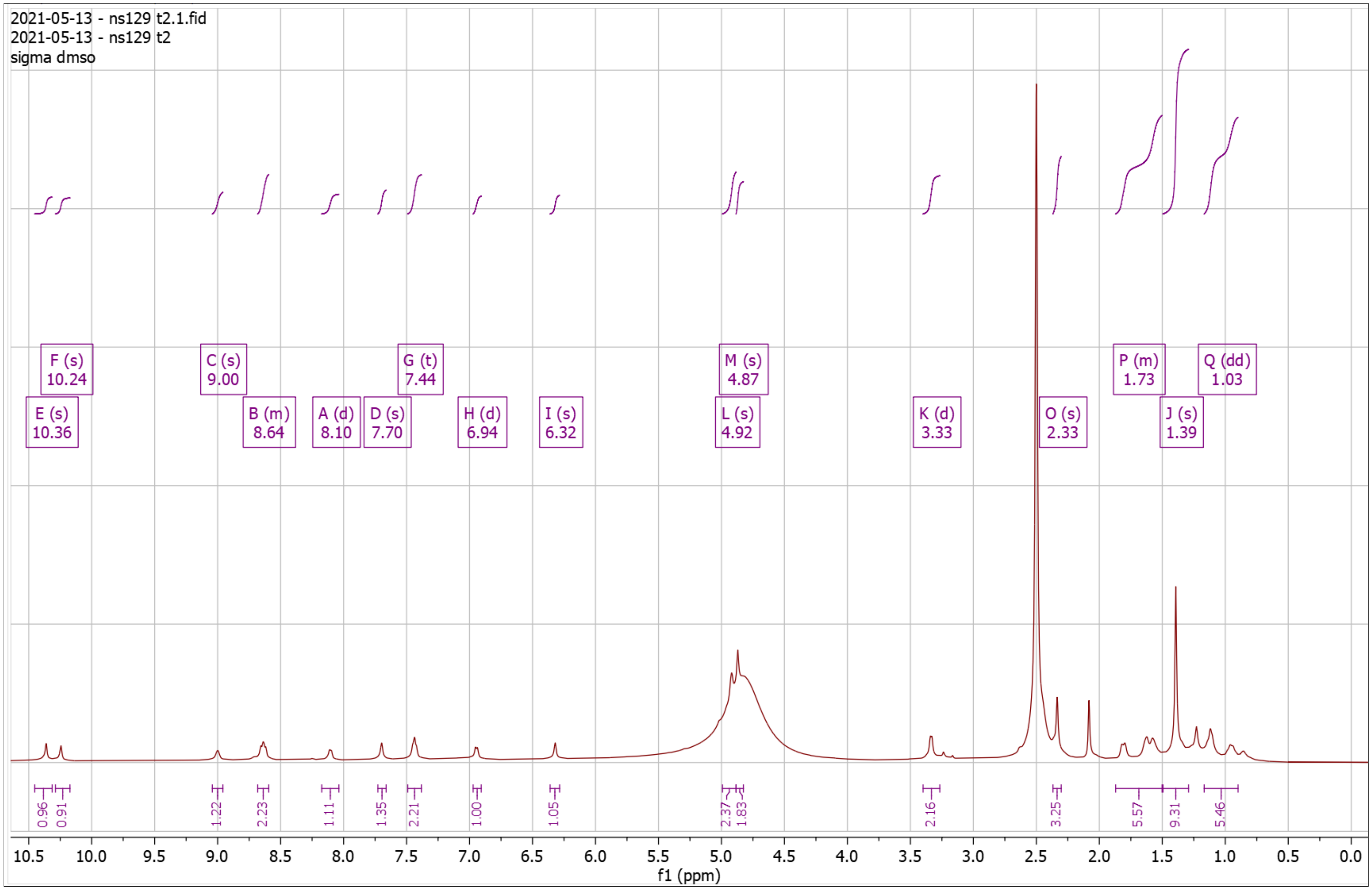


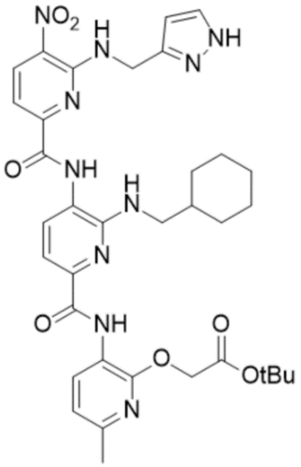

**Supplementary Fig. 84. High Resolution Mass Spectrum for NS129 Pro**

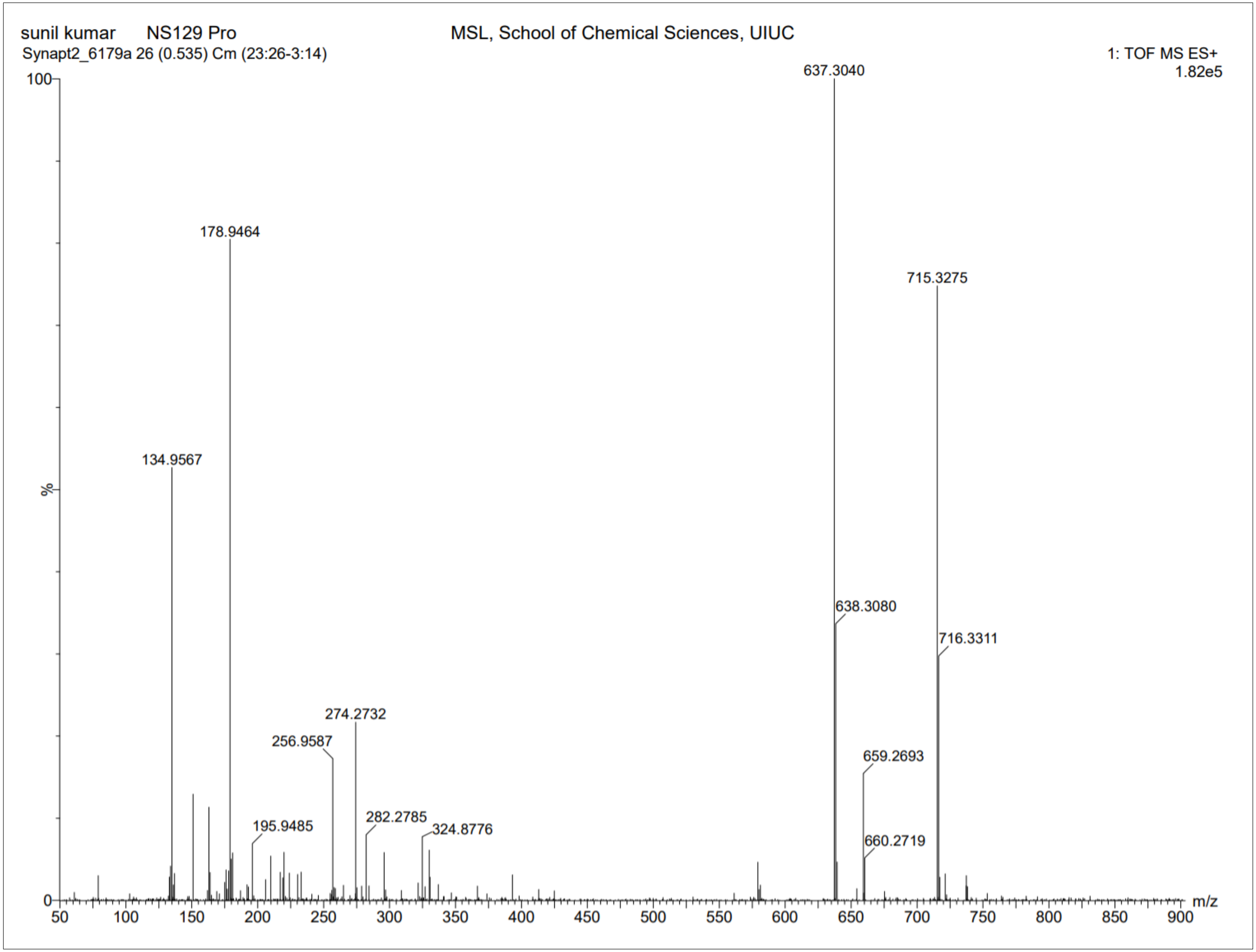

**Supplementary Fig. 85. $^{1}$H-NMR of NS129 Dep**

$^{1}$H NMR (500 MHz, DMSO) δ 0.78 – 1.32 (m, 5H), 1.50 – 1.95 (m, 6H), 2.33 – 2.37 (s, 3H), 3.31 – 3.36 (d, $J$ = 6.1 Hz, 2H), 4.87 – 4.92 (d, $J$ = 5.1 Hz, 2H), 4.95 – 4.98 (s, 2H), 6.26 – 6.30 (s, 1H), 6.92 – 6.97 (d, $J$ = 8.0 Hz, 1H), 7.10 – 7.14 (s, 1H), 7.16 – 7.22 (d, $J$ = 10.3 Hz, 1H), 7.40 – 7.48 (dd, $J$ = 13.0, 8.0 Hz, 2H), 7.62 – 7.66 (s, 1H), 8.13 – 8.18 (d, $J$ = 8.0 Hz, 1H), 8.61 – 8.68 (t, $J$ = 8.2 Hz, 2H), 8.97 – 9.03 (d, $J$ = 5.8 Hz, 1H), 10.27 – 10.31 (s, 1H), 10.38 – 10.41 (s, 1H). HRMS (m/z): [M]+ calcd. for $C_{31}H_{34}N_{10}O_7$, 659.2685; found, 659.2673.

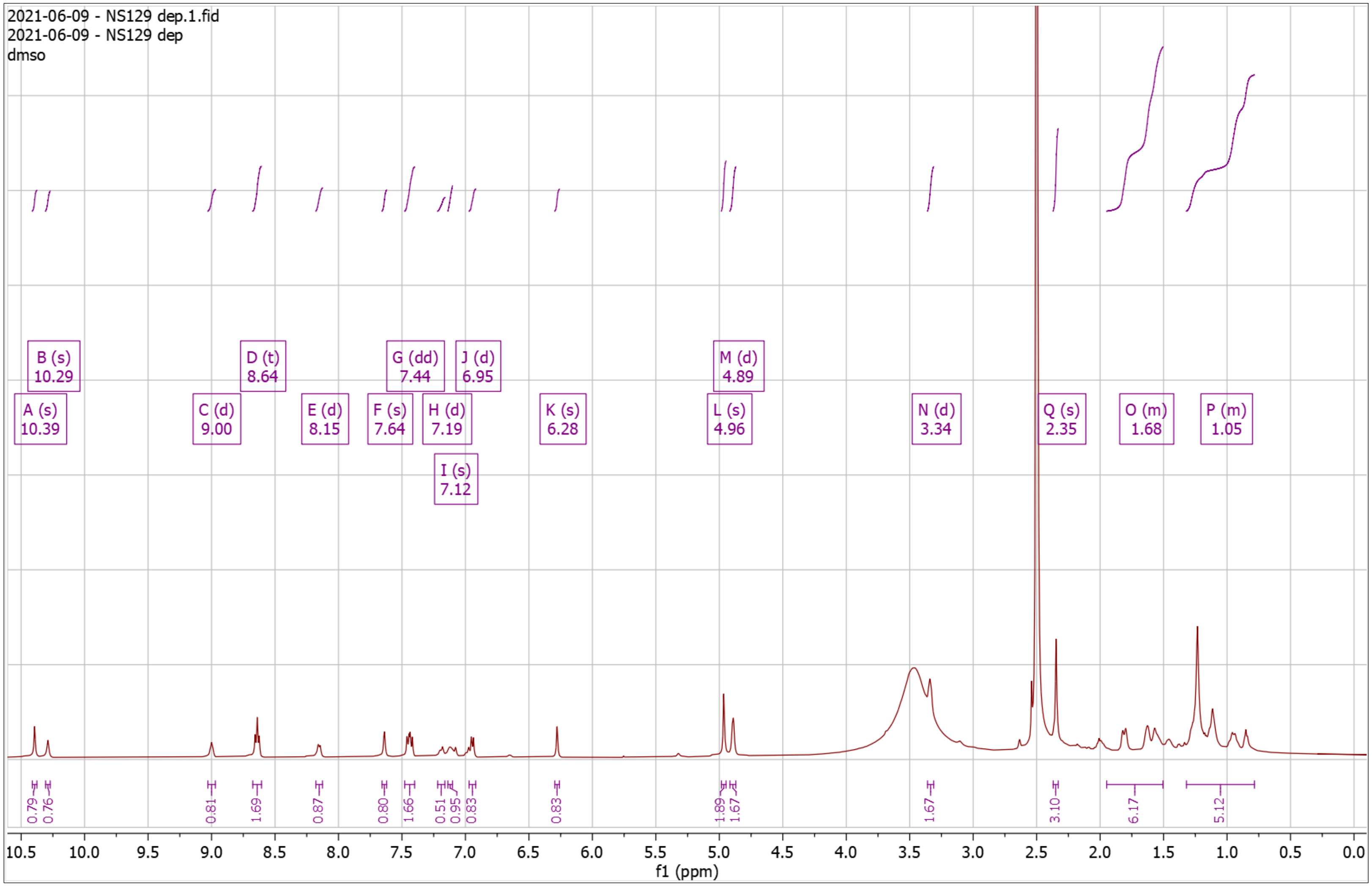


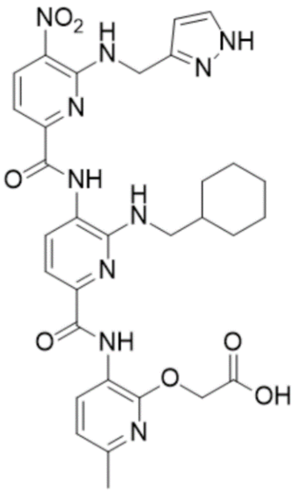

**Supplementary Fig. 86. High Resolution Mass Spectrum for NS129 Dep**

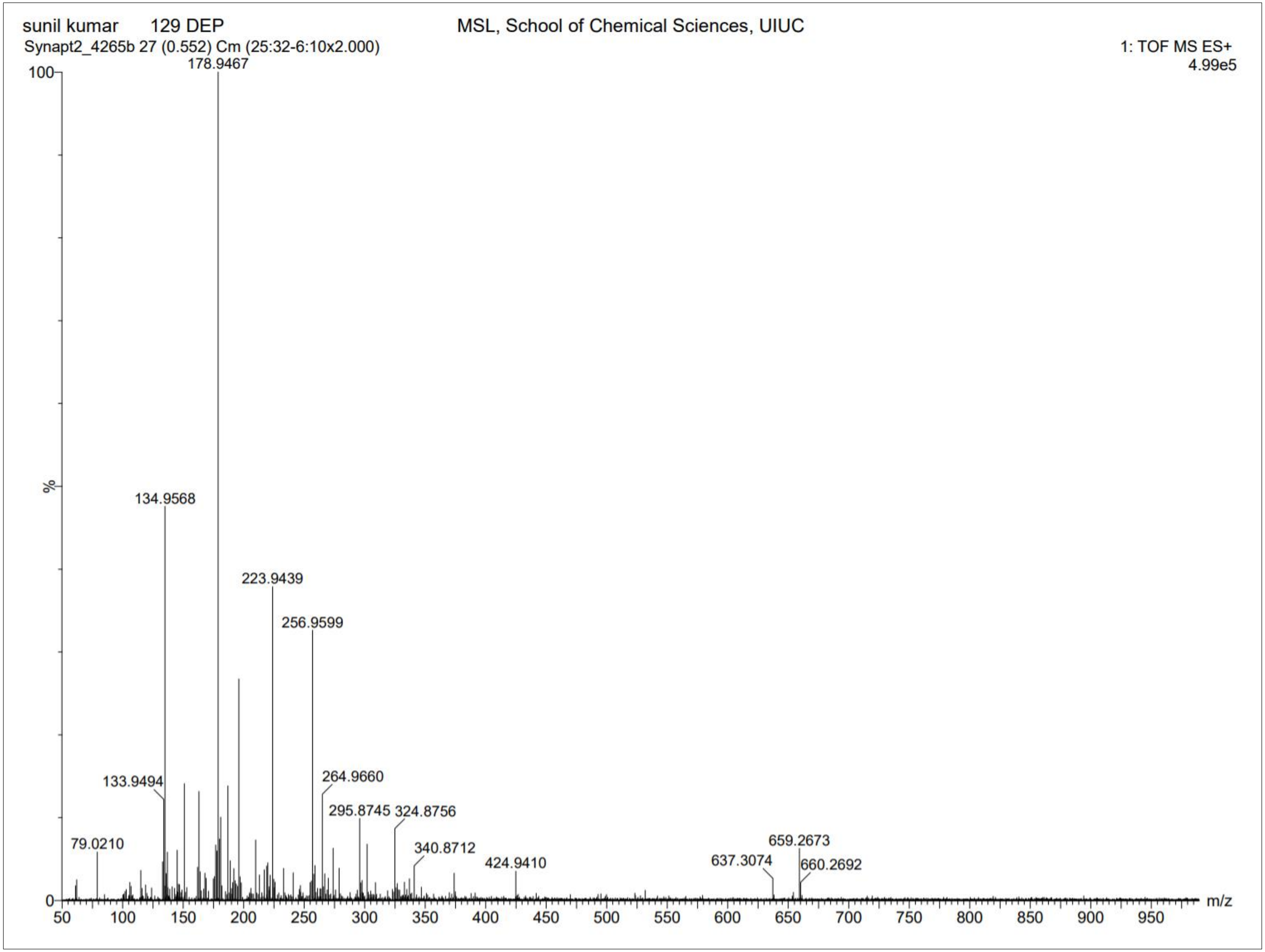

**Supplementary Fig. 87. [1]H-NMR of NS130 Pro**

[1]H NMR (500 MHz, $CDCl_3$) δ 0.76 – 1.21 (m, 5H), 1.24 – 1.33 (s, 9H), 1.33 – 1.39 (s, 9H), 1.56 – 1.91 (m, 6H), 2.30 – 2.33 (s, 3H), 2.54 – 2.60 (t, *J* = 6.1 Hz, 2H), 3.38 – 3.43 (d, *J* = 6.8 Hz, 2H), 3.97 – 4.04 (q, *J* = 6.2 Hz, 2H), 4.77 – 4.82 (s, 2H), 6.72 – 6.77 (d, *J* = 8.0 Hz, 1H), 7.54 – 7.60 (dd, *J* = 8.1, 3.4 Hz, 2H), 7.98 – 8.04 (d, *J* = 7.9 Hz, 1H), 8.55 – 8.60 (d, *J* = 8.4 Hz, 1H), 8.65 – 8.70 (d, *J* = 7.9 Hz, 1H), 9.52 – 9.56 (s, 1H), 10.31 – 10.34 (s, 1H). HRMS (m/z): [M]+ calcd. for $C_{38}H_{50}N_8O_9$, 763.3774; found, 763.3752.

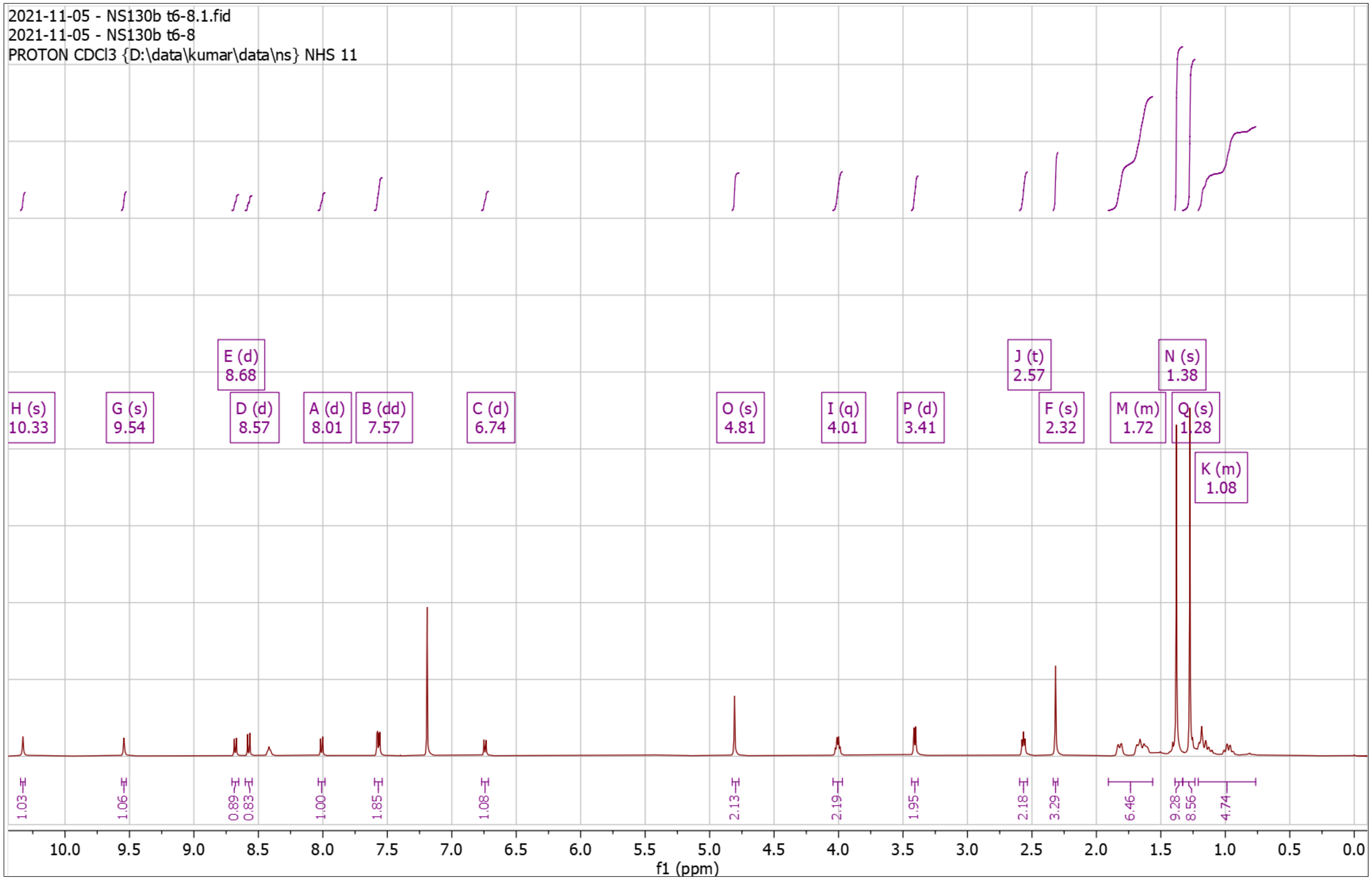


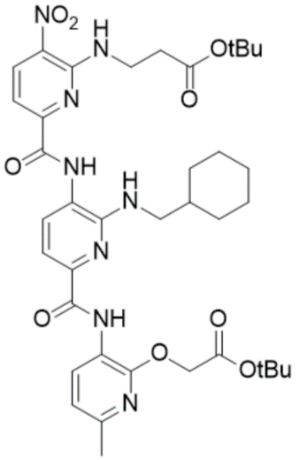

**Supplementary Fig. 88. High Resolution Mass Spectrum for NS130 Pro**

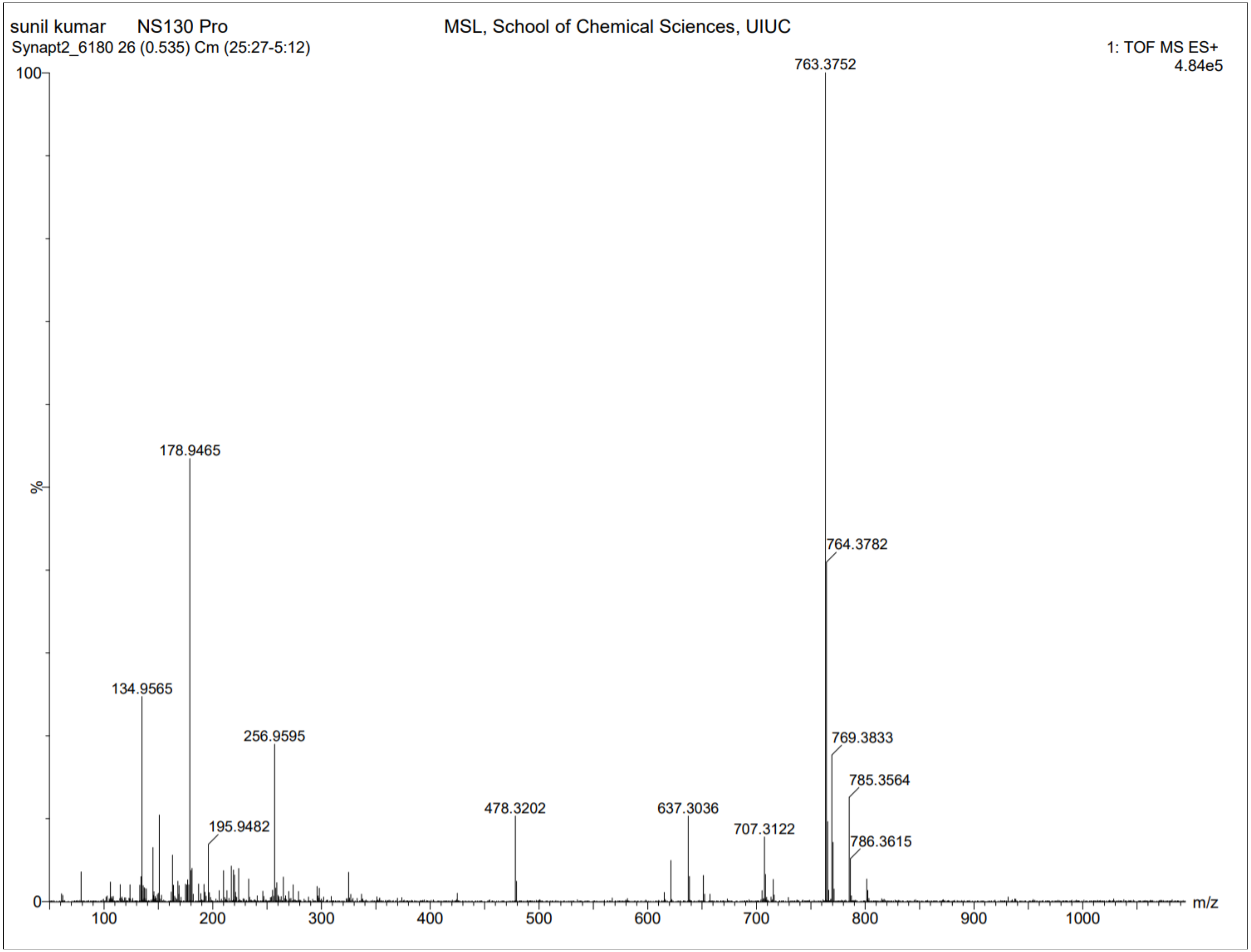

**Supplementary Fig. 89. $^1$H-NMR of NS130 Dep**

$^1$H NMR (500 MHz, DMSO) δ 0.90 – 1.27 (dt, $J$ = 96.8, 13.0 Hz, 5H), 1.56 – 1.86 (dd, $J$ = 92.9, 19.4 Hz, 6H), 2.34 – 2.35 (s, 3H), 2.61 – 2.67 (t, $J$ = 6.4 Hz, 2H), 3.98 – 4.06 (q, $J$ = 6.3 Hz, 2H), 4.94 – 4.98 (s, 2H), 6.69 – 6.75 (t, $J$ = 5.8 Hz, 1H), 6.92 – 6.97 (d, $J$ = 8.0 Hz, 1H), 7.38 – 7.44 (d, $J$ = 8.1 Hz, 2H), 7.84 – 7.89 (d, $J$ = 7.7 Hz, 1H), 8.61 – 8.67 (dd, $J$ = 8.2, 4.9 Hz, 2H), 8.68 – 8.74 (t, $J$ = 5.9 Hz, 1H), 9.91 – 9.95 (s, 1H), 10.35 – 10.38 (s, 1H). HRMS (m/z): [M]+ calcd. for $C_{30}H_{34}N_8O_9$, 651.2522; found, 651.2520.

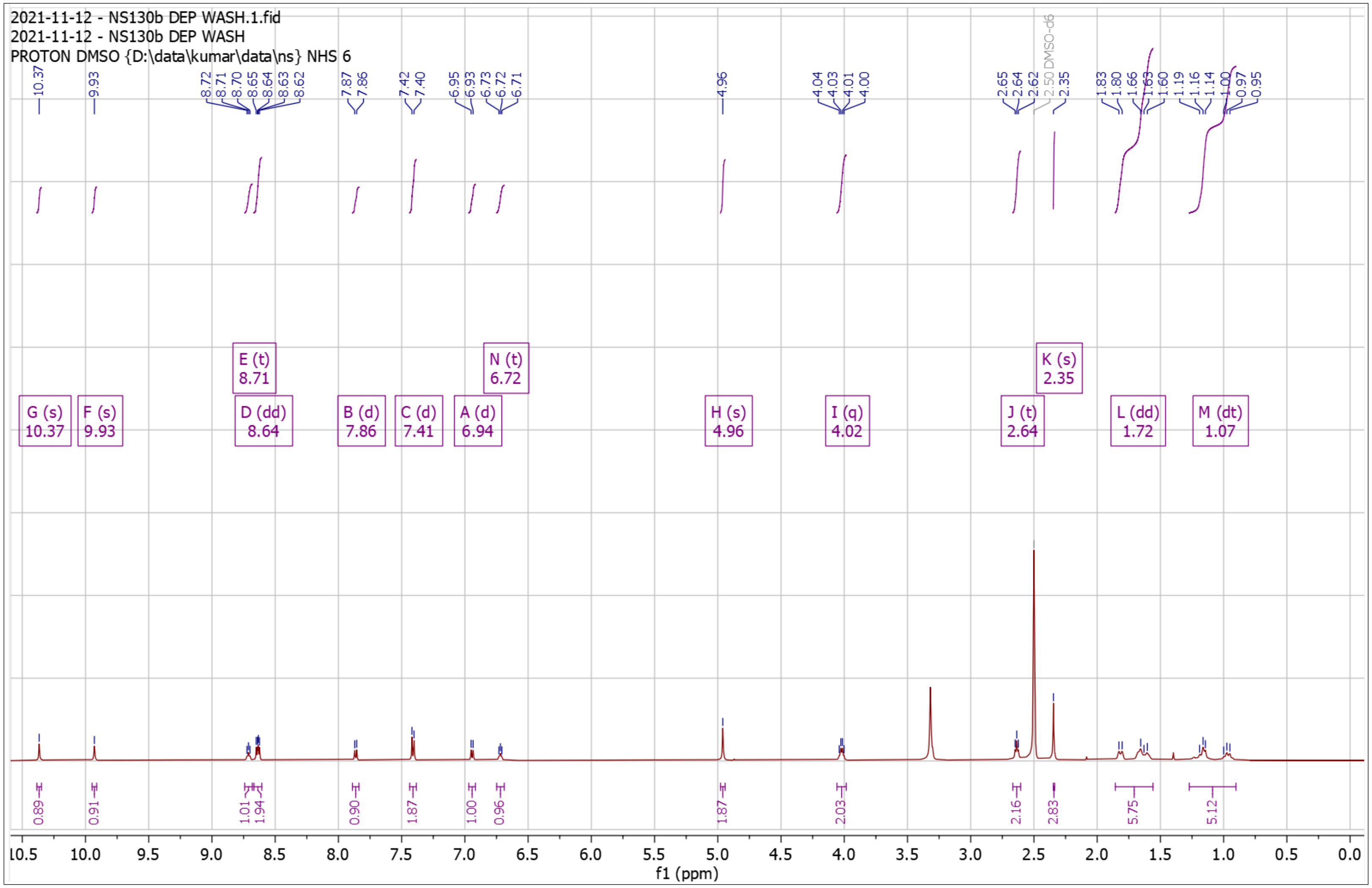


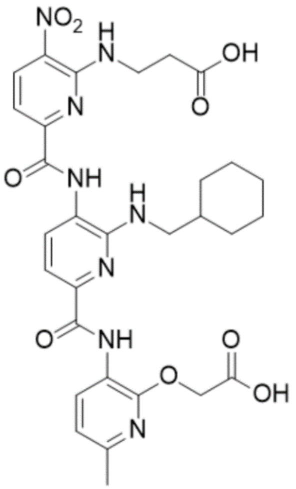

**Supplementary Fig. 90. High Resolution Mass Spectrum for NS130 Dep**

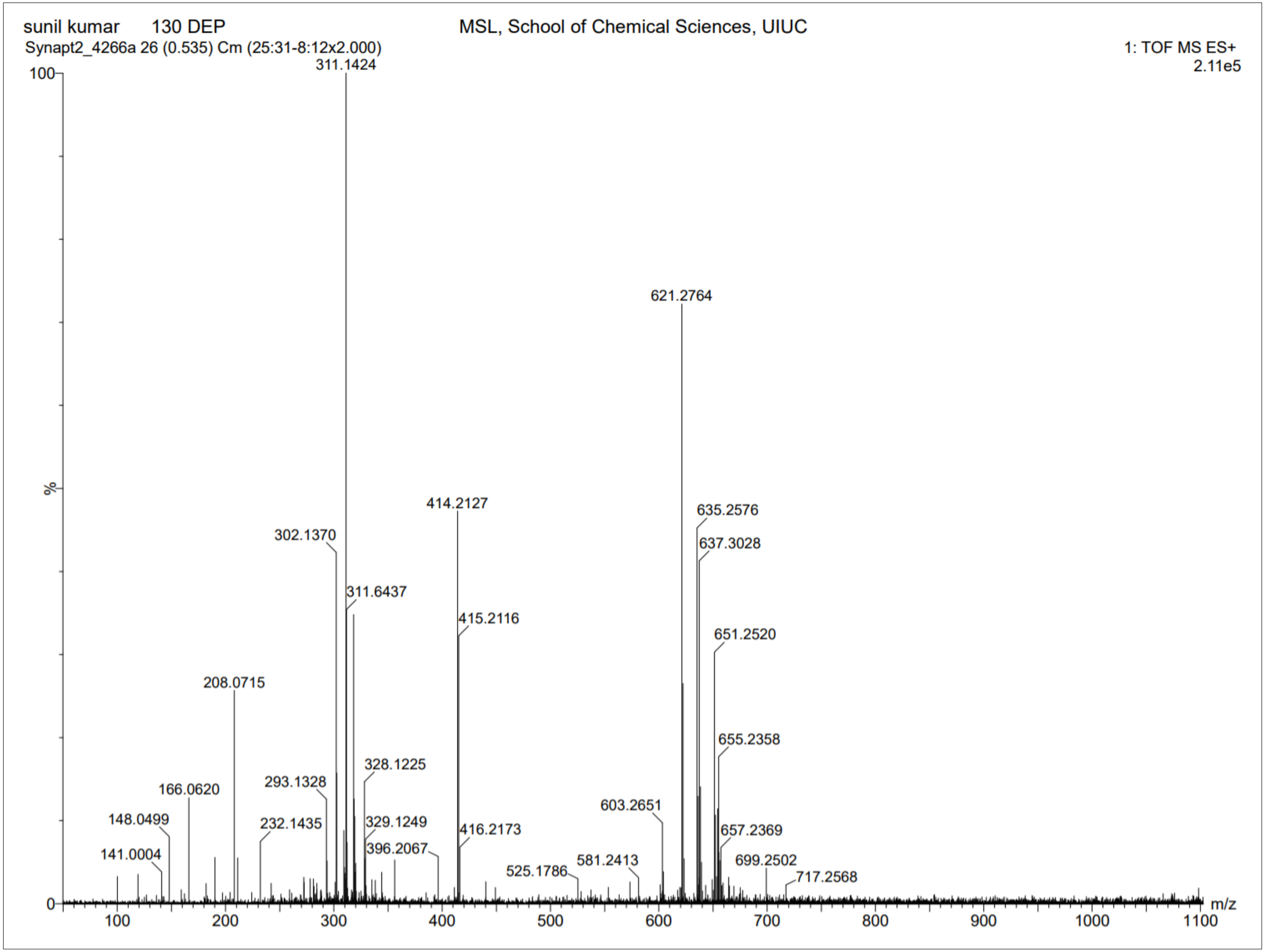

**Supplementary Fig. 91. $^{1}$H-NMR of NS131 Pro**

$^{1}$H NMR (500 MHz, DMSO) δ 0.91 – 1.30 (m, 5H), 1.39 – 1.40 (s, 9H), 1.60 – 1.87 (m, 6H), 2.32 – 2.35 (s, 3H), 2.75 – 2.84 (t, *J* = 6.2 Hz, 2H), 3.97 – 4.13 (q, *J* = 6.1 Hz, 2H), 4.80 – 4.97 (s, 2H), 6.72 – 6.84 (t, *J* = 5.4 Hz, 1H), 6.91 – 6.98 (d, *J* = 7.9 Hz, 1H), 7.35 – 7.45 (dd, *J* = 8.1, 4.4 Hz, 2H), 7.85 – 7.94 (d, *J* = 7.6 Hz, 1H), 8.57 – 8.67 (dd, *J* = 8.2, 4.0 Hz, 2H), 9.05 – 9.14 (t, *J* = 5.5 Hz, 1H), 10.28 – 10.34 (s, 1H), 10.34 – 10.42 (s, 1H). HRMS (m/z): [M]+ calcd. for $C_{33}H_{42}N_8O_{10}S$, 743.2817; found, 743.2793.

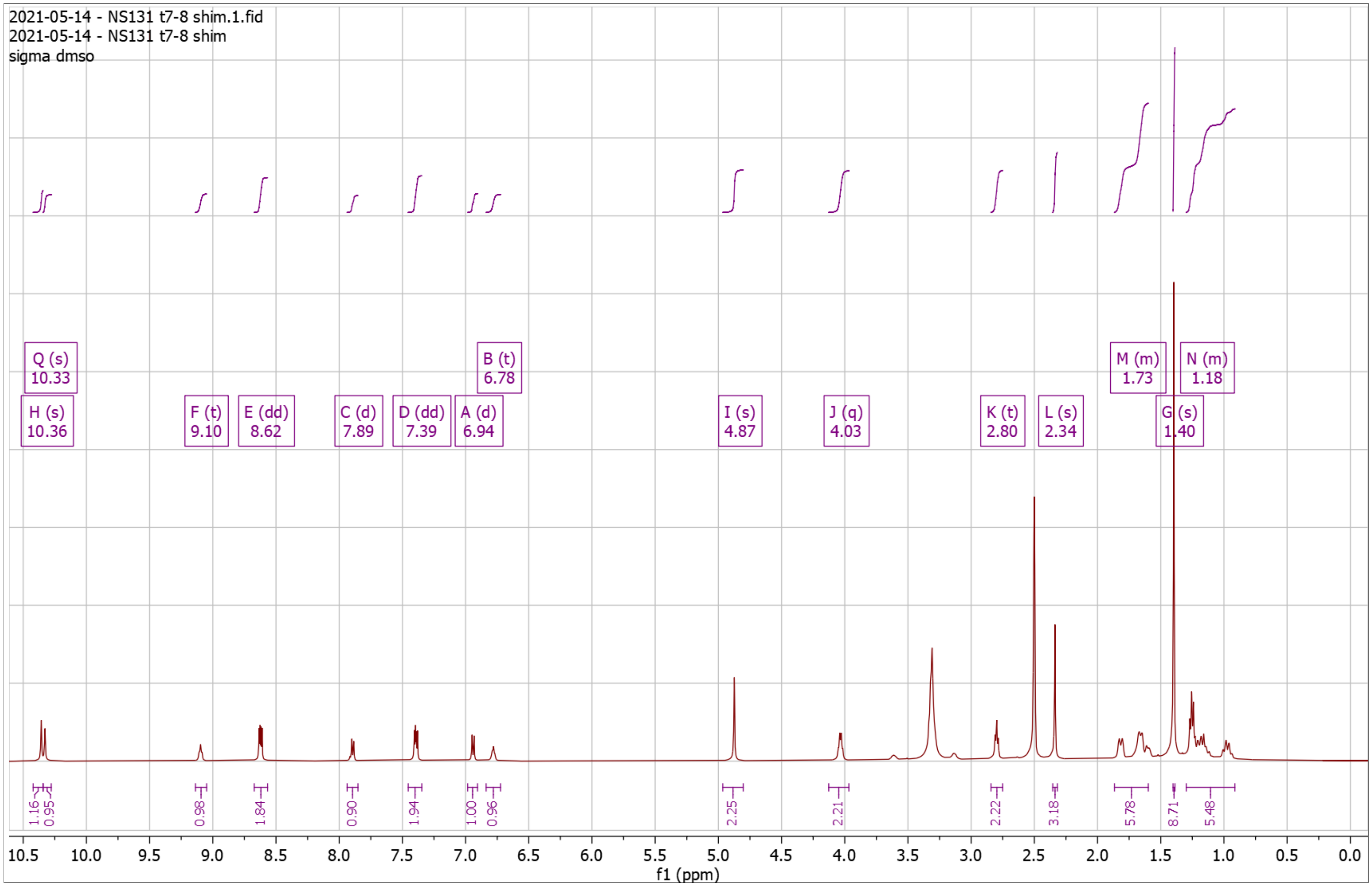


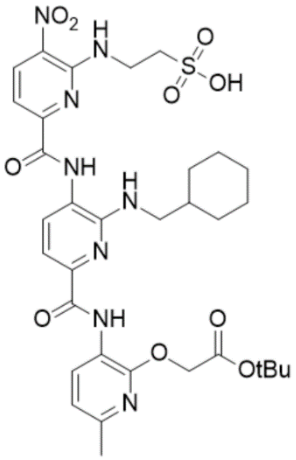

**Supplementary Fig. 92. High Resolution Mass Spectrum for NS131 Pro**

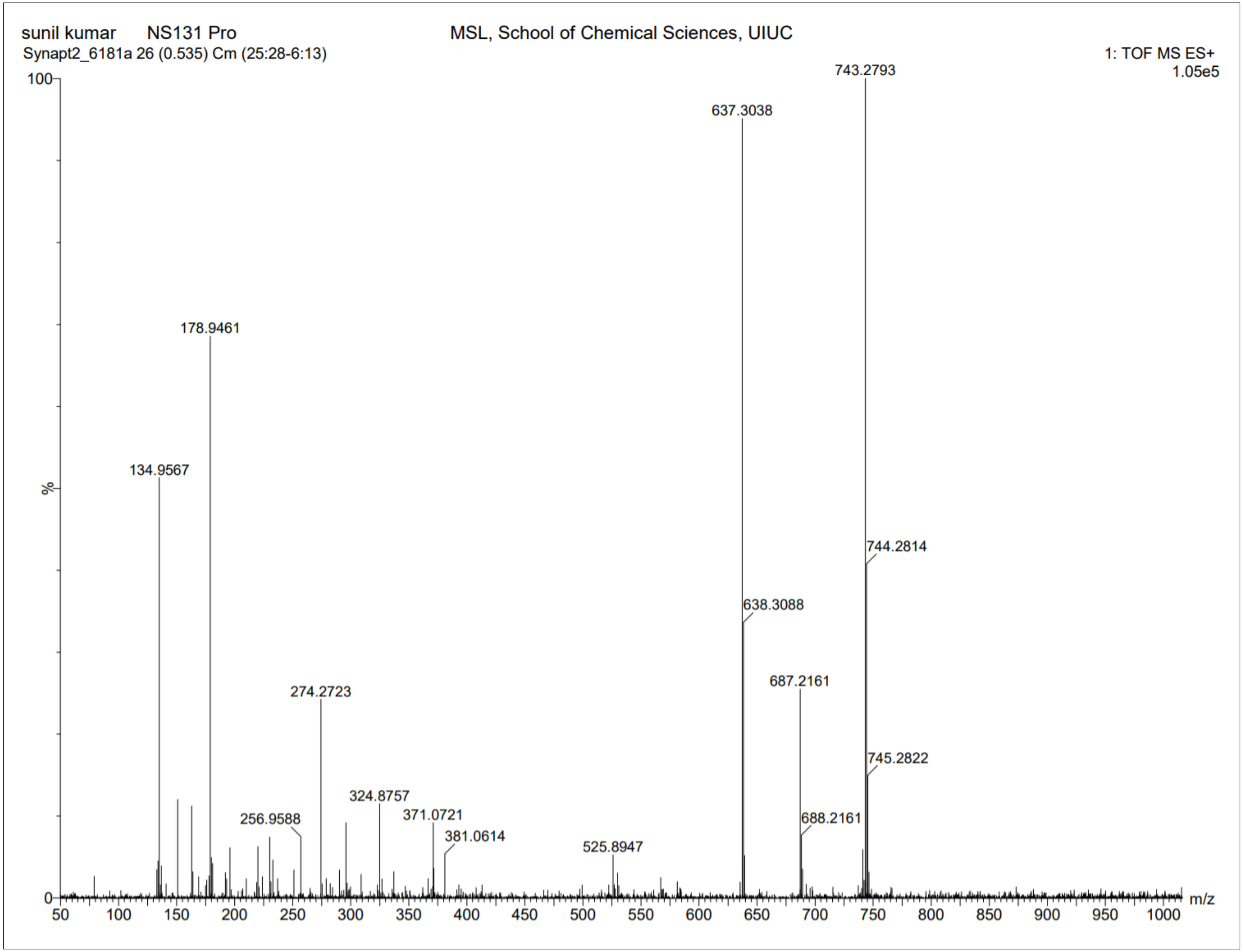

**Supplementary Fig. 93. $^{1}$H-NMR of NS131 Dep**

$^{1}$H NMR (500 MHz, DMSO) δ 0.84 – 1.19 (m, 5H), 1.53 – 1.78 (m, 6H), 2.25 – 2.29 (s, 3H), 2.69 – 2.75 (t, *J* = 6.3 Hz, 2H), 3.92 – 4.00 (q, *J* = 6.1 Hz, 2H), 4.86 – 4.90 (s, 2H), 6.67 – 6.74 (t, *J* = 5.6 Hz, 1H), 6.84 – 6.90 (d, *J* = 7.9 Hz, 1H), 7.29 – 7.35 (dd, *J* = 8.1, 4.3 Hz, 2H), 7.80 – 7.85 (d, *J* = 7.7 Hz, 1H), 8.53 – 8.61 (dd, *J* = 8.1, 3.2 Hz, 2H), 9.00 – 9.06 (t, *J* = 5.6 Hz, 1H), 10.25 – 10.28 (s, 1H), 10.31 – 10.35 (s, 1H). HRMS (m/z): [M]+ calcd. for $C_{29}H_{34}N_{8}O_{10}S$, 687.2191; found, 687.2180.

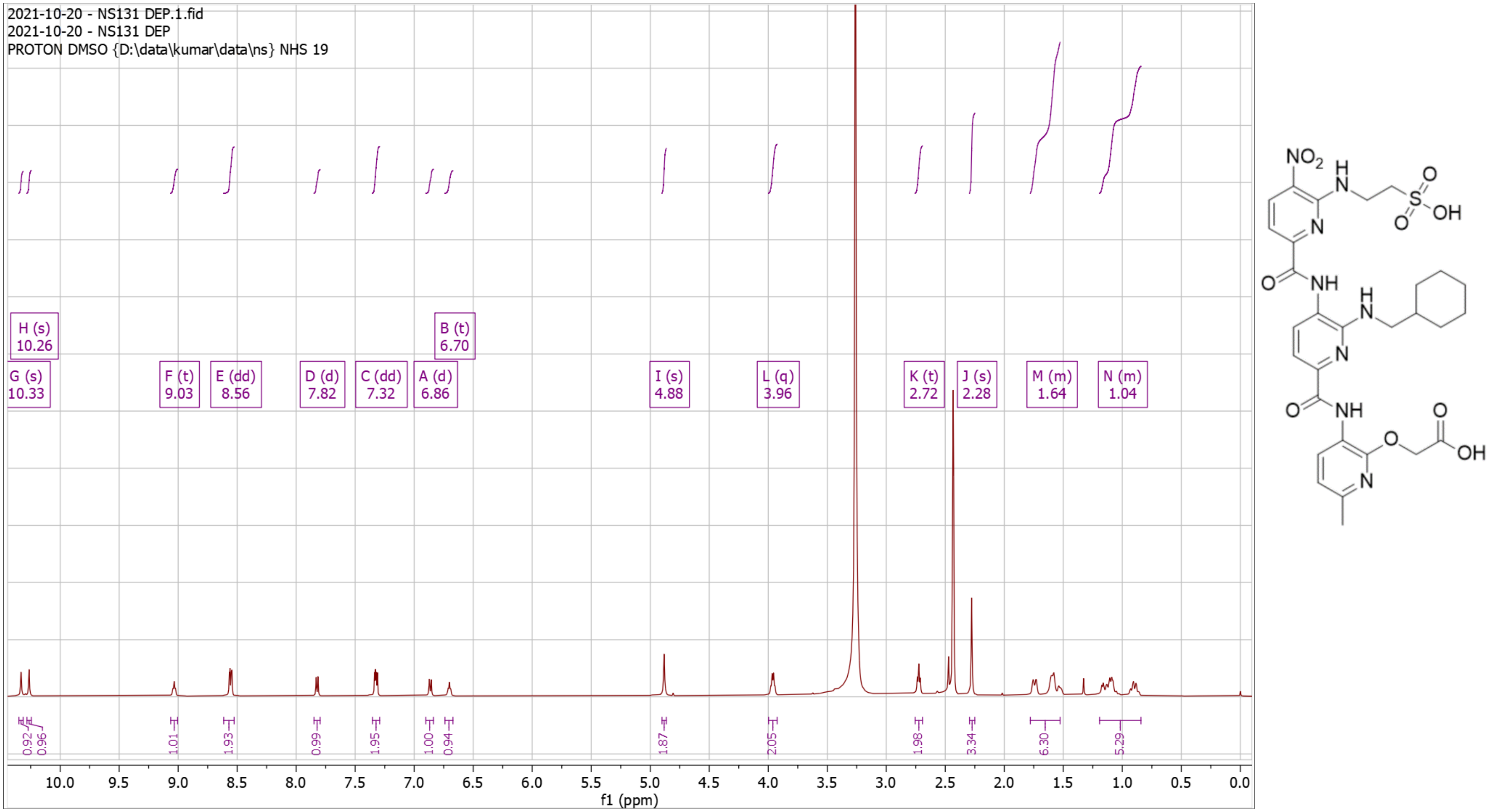

**Supplementary Fig. 94. High Resolution Mass Spectrum for NS131 Dep**

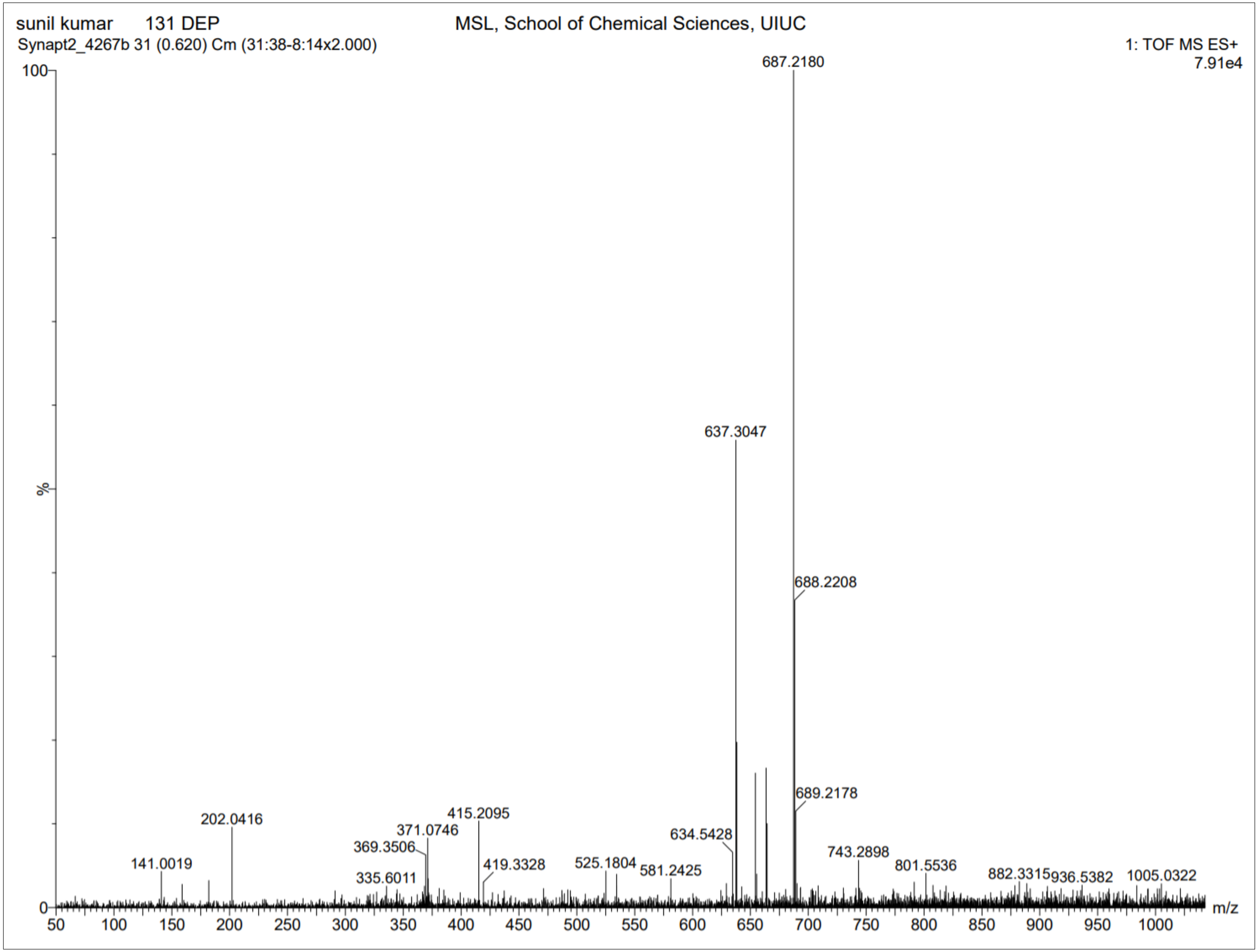

**Supplementary Fig. 95. $^1$H-NMR of NS132 Pro**

$^1$H NMR (500 MHz, DMSO) δ 0.81 – 1.18 (dt, $J$ = 94.1, 11.7 Hz, 5H), 1.36 – 1.42 (s, 9H), 1.50 – 1.80 (m, 6H), 2.33 – 2.35 (s, 3H), 3.06 – 3.13 (t, $J$ = 7.1 Hz, 2H), 3.23 – 3.29 (t, $J$ = 6.1 Hz, 2H), 3.98 – 4.13 (q, $J$ = 6.8 Hz, 2H), 4.83 – 4.90 (s, 2H), 6.71 – 6.76 (t, $J$ = 5.6 Hz, 1H), 6.76 – 6.82 (t, $J$ = 7.5 Hz, 1H), 6.91 – 6.97 (d, $J$ = 7.9 Hz, 1H), 6.97 – 7.03 (t, $J$ = 7.5 Hz, 1H), 7.19 – 7.23 (d, $J$ = 2.3 Hz, 1H), 7.27 – 7.31 (d, $J$ = 8.1 Hz, 1H), 7.34 – 7.39 (d, $J$ = 8.4 Hz, 1H), 7.41 – 7.47 (d, $J$ = 7.7 Hz, 1H), 7.51 – 7.58 (d, $J$ = 7.9 Hz, 1H), 7.92 – 7.97 (d, $J$ = 7.9 Hz, 1H), 8.55 – 8.68 (m, 3H), 9.81 – 9.87 (s, 1H), 10.31 – 10.38 (s, 1H), 10.79 – 10.85 (m, 1H). HRMS (m/z): [M]+ calcd. for $C_{41}H_{47}N_9O_7$, 778.3671; found, 778.3665.

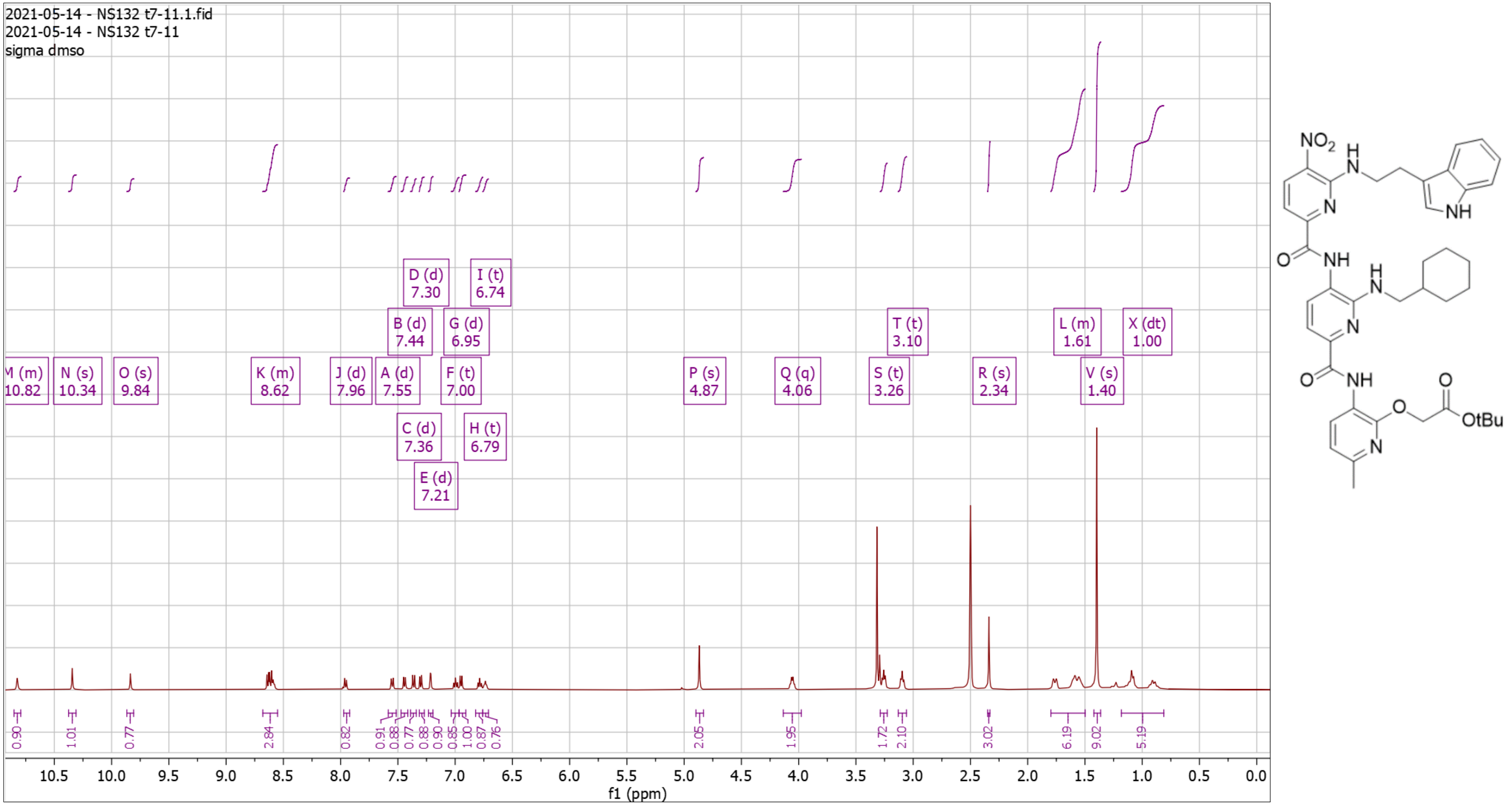

**Supplementary Fig. 96. High Resolution Mass Spectrum for NS132 Pro**

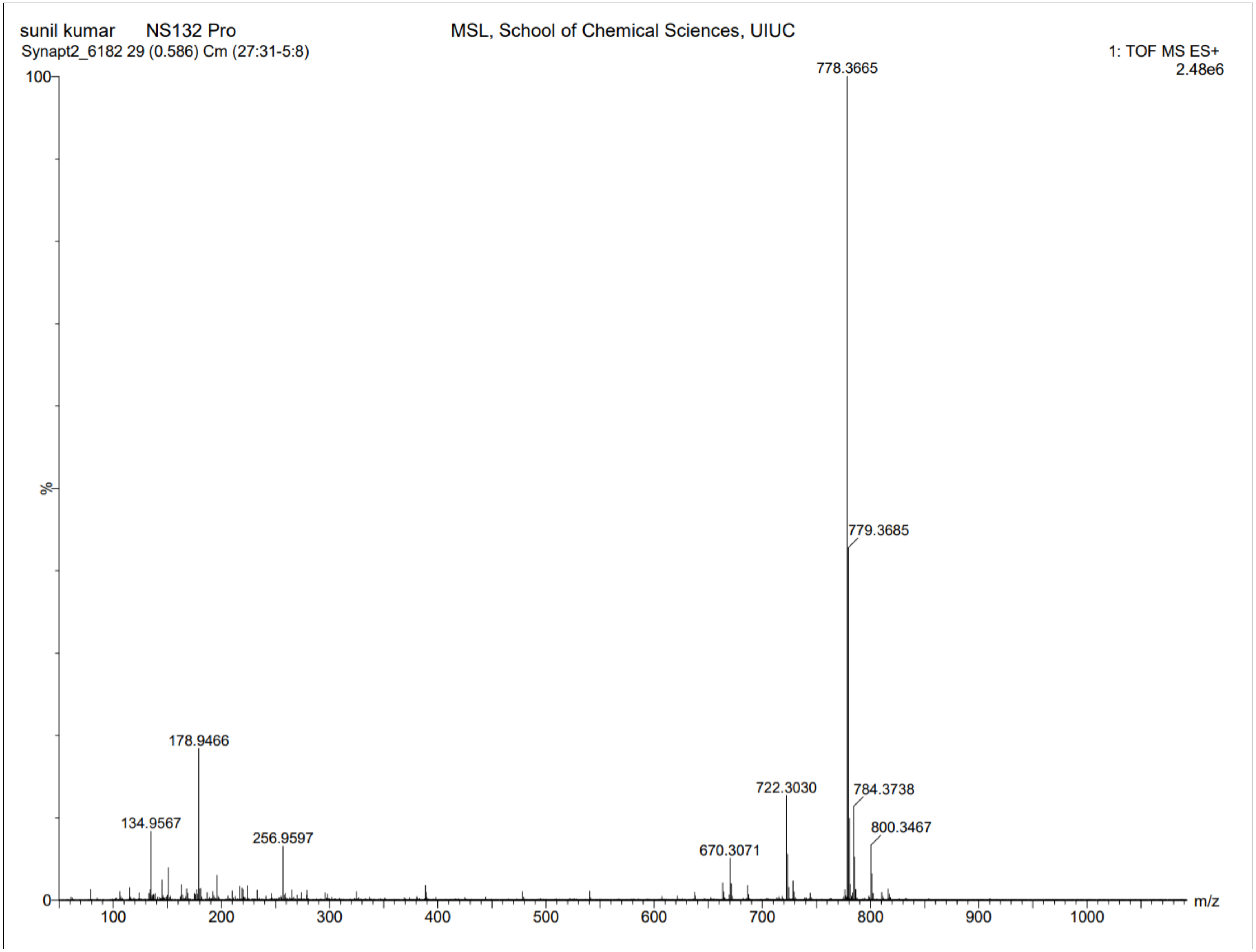

**Supplementary Fig. 97. $^{1}$H-NMR of NS132 Dep**

$^{1}$H NMR (500 MHz, DMSO) δ 0.84 – 1.13 (m, 5H), 1.49 – 1.83 (m, 6H), 2.33 – 2.36 (s, 3H), 3.06 – 3.16 (t, $J$ = 7.1 Hz, 2H), 3.22 – 3.30 (t, $J$ = 6.2 Hz, 2H), 3.99 – 4.14 (q, $J$ = 6.7 Hz, 2H), 4.94 – 4.98 (s, 2H), 6.69 – 6.77 (t, $J$ = 5.7 Hz, 1H), 6.77 – 6.83 (t, $J$ = 7.4 Hz, 1H), 6.91 – 6.96 (d, $J$ = 7.9 Hz, 1H), 6.97 – 7.03 (t, $J$ = 7.6 Hz, 1H), 7.20 – 7.24 (d, $J$ = 2.3 Hz, 1H), 7.28 – 7.33 (d, $J$ = 8.1 Hz, 1H), 7.33 – 7.41 (d, $J$ = 8.5 Hz, 1H), 7.41 – 7.47 (d, $J$ = 7.7 Hz, 1H), 7.53 – 7.58 (d, $J$ = 7.9 Hz, 1H), 7.93 – 7.98 (d, $J$ = 7.7 Hz, 1H), 8.56 – 8.66 (td, $J$ = 13.7, 12.0, 6.8 Hz, 3H), 9.83 – 9.87 (s, 1H), 10.33 – 10.38 (s, 1H), 10.83 – 10.87 (s, 1H), 12.88 – 13.07 (s, 1H). HRMS (m/z): [M]+ calcd. for $C_{37}H_{39}N_9O_7$, 722.3045; found, 722.3034.

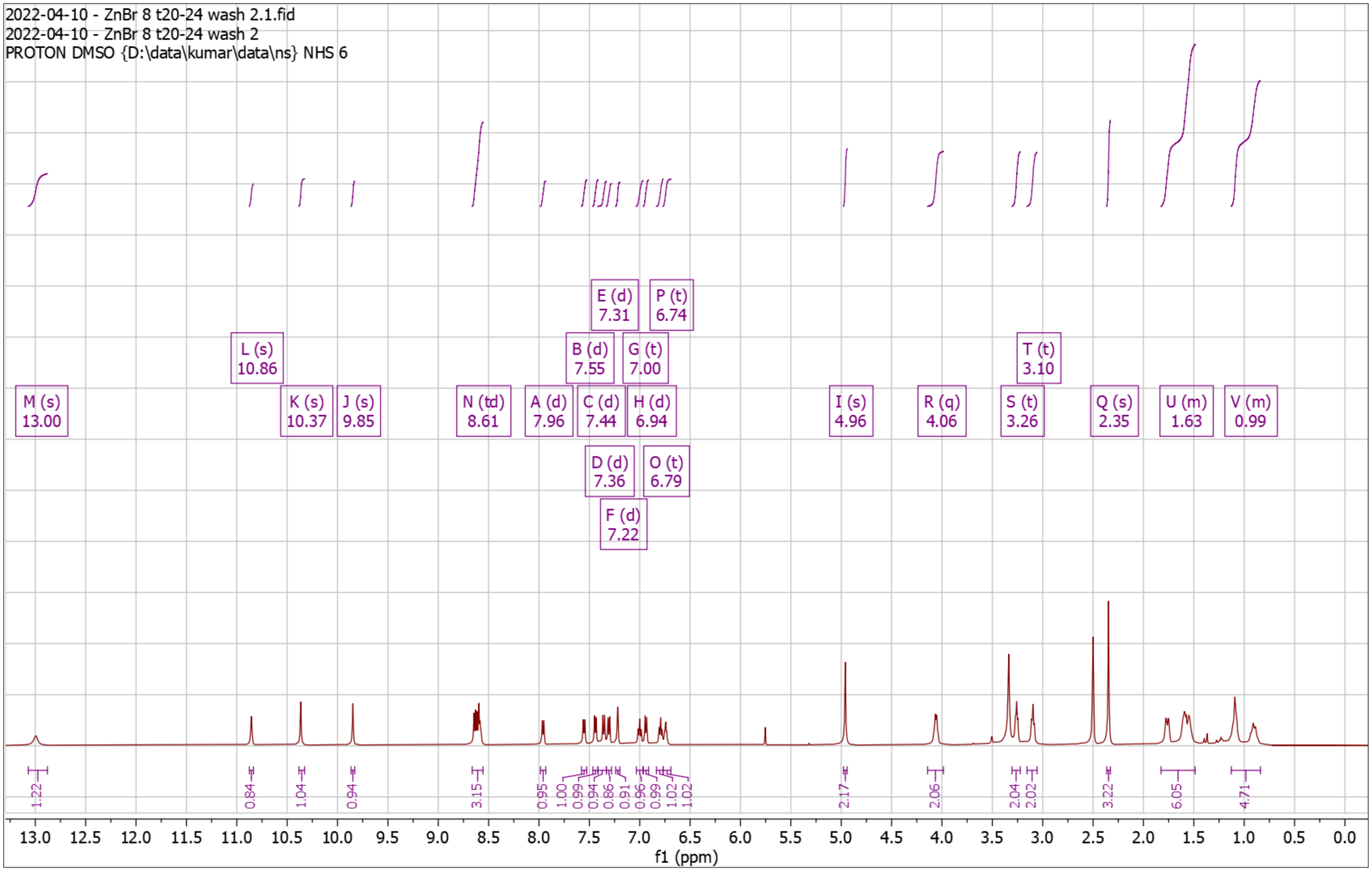


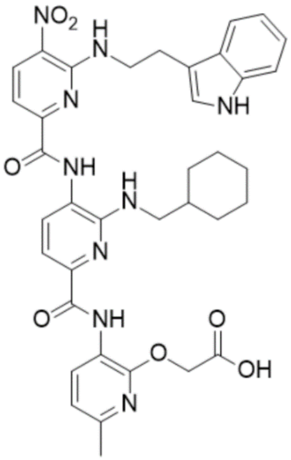

**Supplementary Fig. 98. $^{13}$C-NMR of NS132 Dep**

13C NMR (126 MHz, DMSO) δ 23.25, 24.73, 25.52, 26.06, 30.69, 37.21, 41.66, 47.07, 54.90, 69.78, 109.64, 110.08, 111.38, 111.43, 116.49, 118.10, 118.16, 119.69, 120.91, 122.40, 123.03, 126.35, 127.18, 129.20, 132.17, 136.32, 137.37, 143.33, 148.53, 150.91, 151.67, 153.73, 162.21.

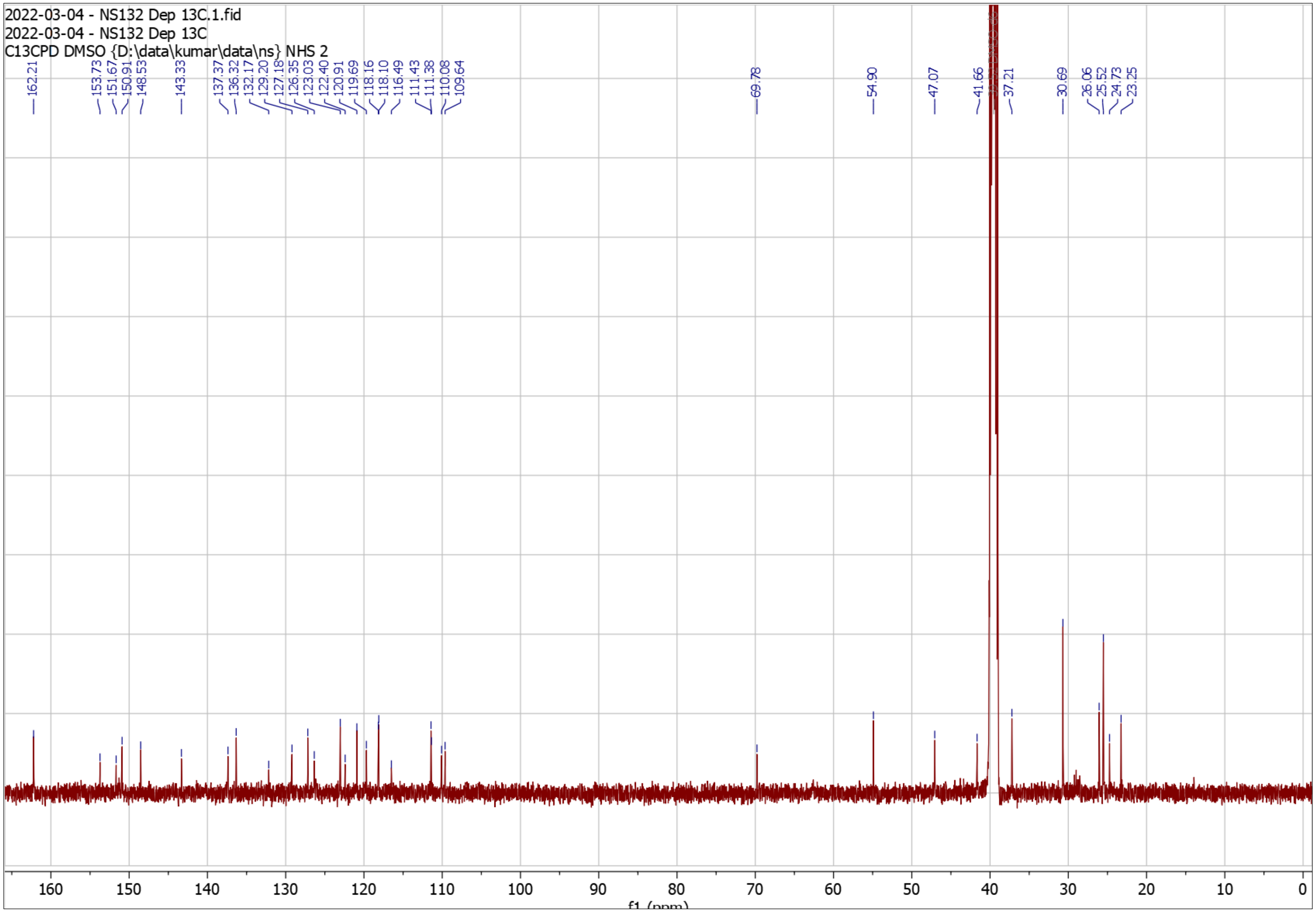


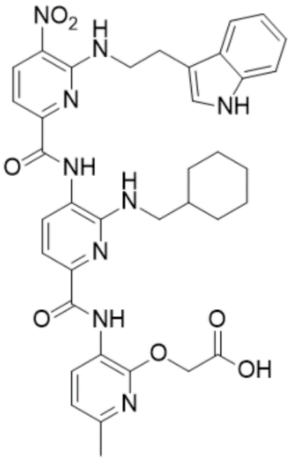

**Supplementary Fig. 99. High Resolution Mass Spectrum for NS132 Dep**

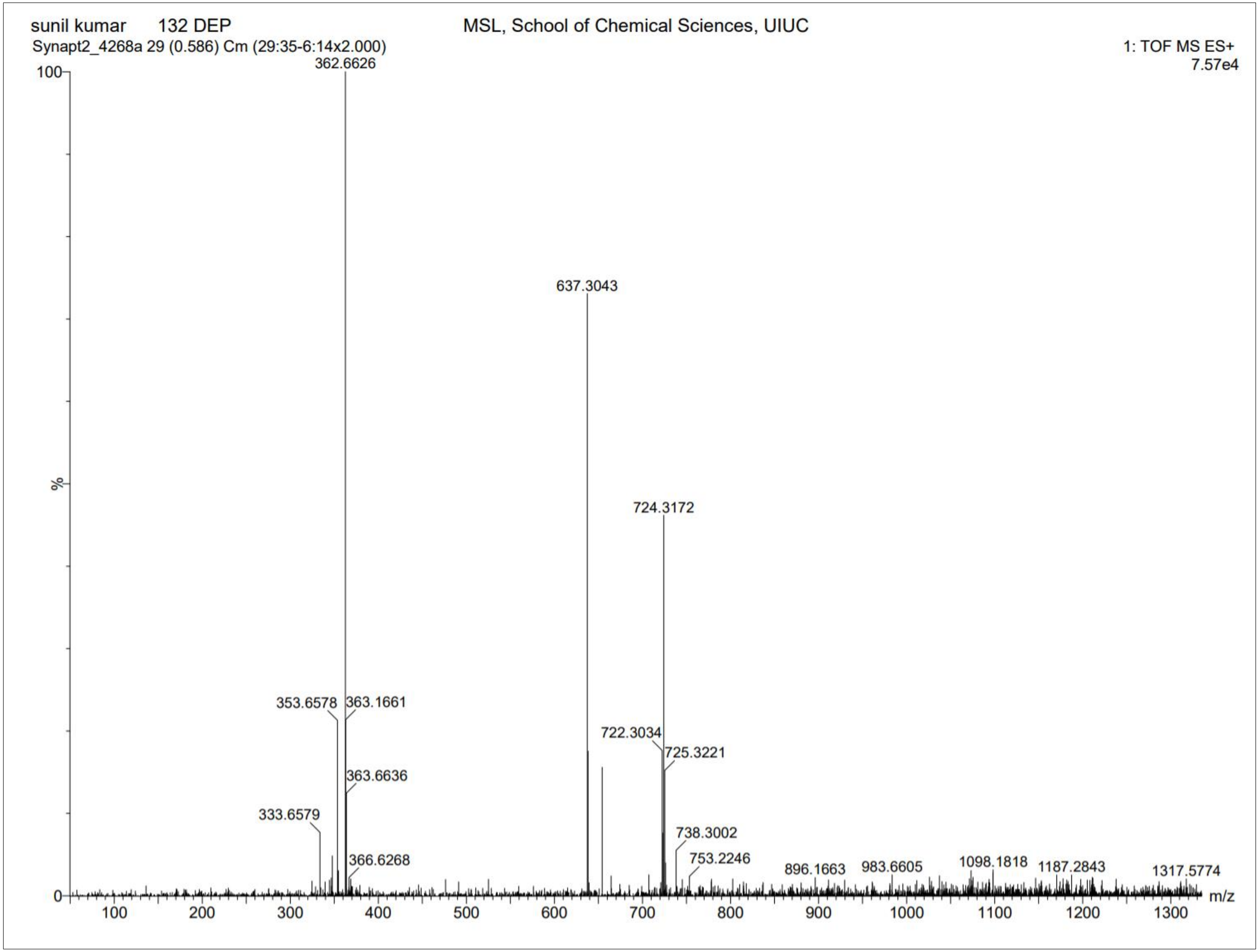

**Supplementary Fig. 100. $^{1}$H-NMR of NS138 Pro**

$^{1}$H NMR (500 MHz, $CDCl_3$) δ 0.72 – 1.08 (m, 3H), 1.08 – 1.10 (d, $J$ = 6.7 Hz, 6H), 1.10 – 1.33 (m, 3H), 1.45 – 1.46 (s, 9H), 1.64 – 1.91 (m, 5H), 2.02 – 2.13 (dt, $J$ = 12.9, 6.5 Hz, 1H), 2.36 – 2.39 (s, 3H), 3.44 – 3.50 (t, $J$ = 6.2 Hz, 2H), 3.50 – 3.55 (dd, $J$ = 6.7, 5.4 Hz, 2H), 4.83 – 4.88 (s, 2H), 6.78 – 6.85 (d, $J$ = 7.9 Hz, 1H), 7.58 – 7.65 (d, $J$ = 8.4 Hz, 1H), 7.65 – 7.69 (d, $J$ = 7.9 Hz, 1H), 7.90 – 7.94 (d, $J$ = 7.9 Hz, 1H), 8.34 – 8.41 (s, 1H), 8.63 – 8.68 (d, $J$ = 8.4 Hz, 1H), 8.71 – 8.75 (d, $J$ = 7.9 Hz, 1H), 9.33 – 9.38 (s, 1H), 10.35 – 10.42 (s, 1H). HRMS (m/z): [M]+ calcd. for $C_{35}H_{46}N_8O_7$, 691.3562; found, 691.3537.

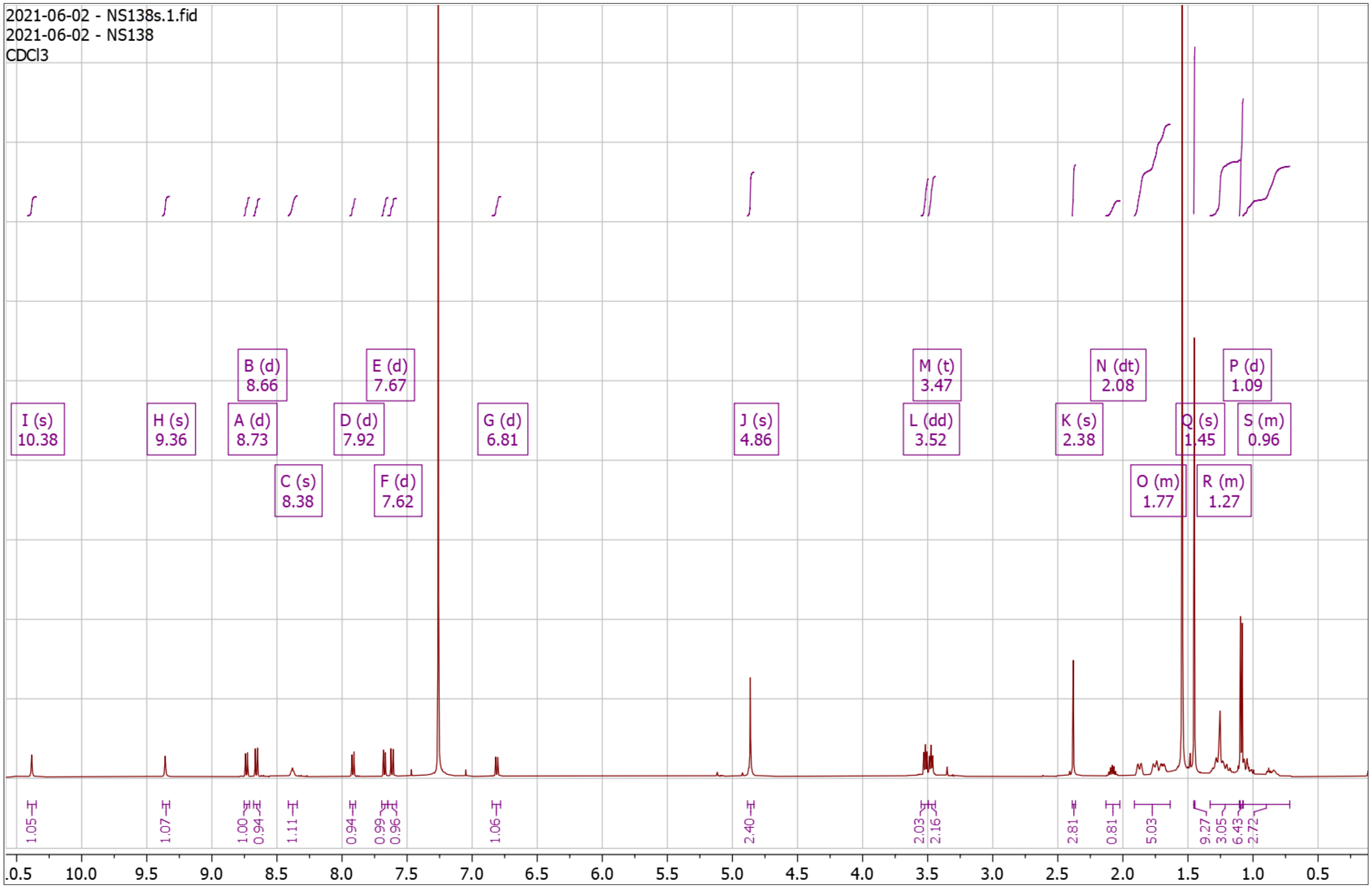


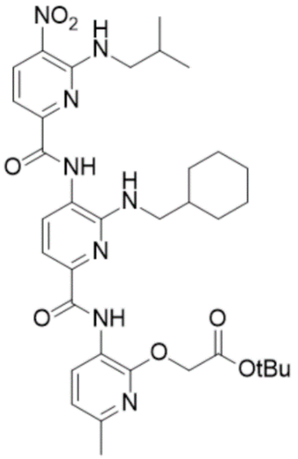

**Supplementary Fig. 101. High Resolution Mass Spectrum for NS138 Pro**

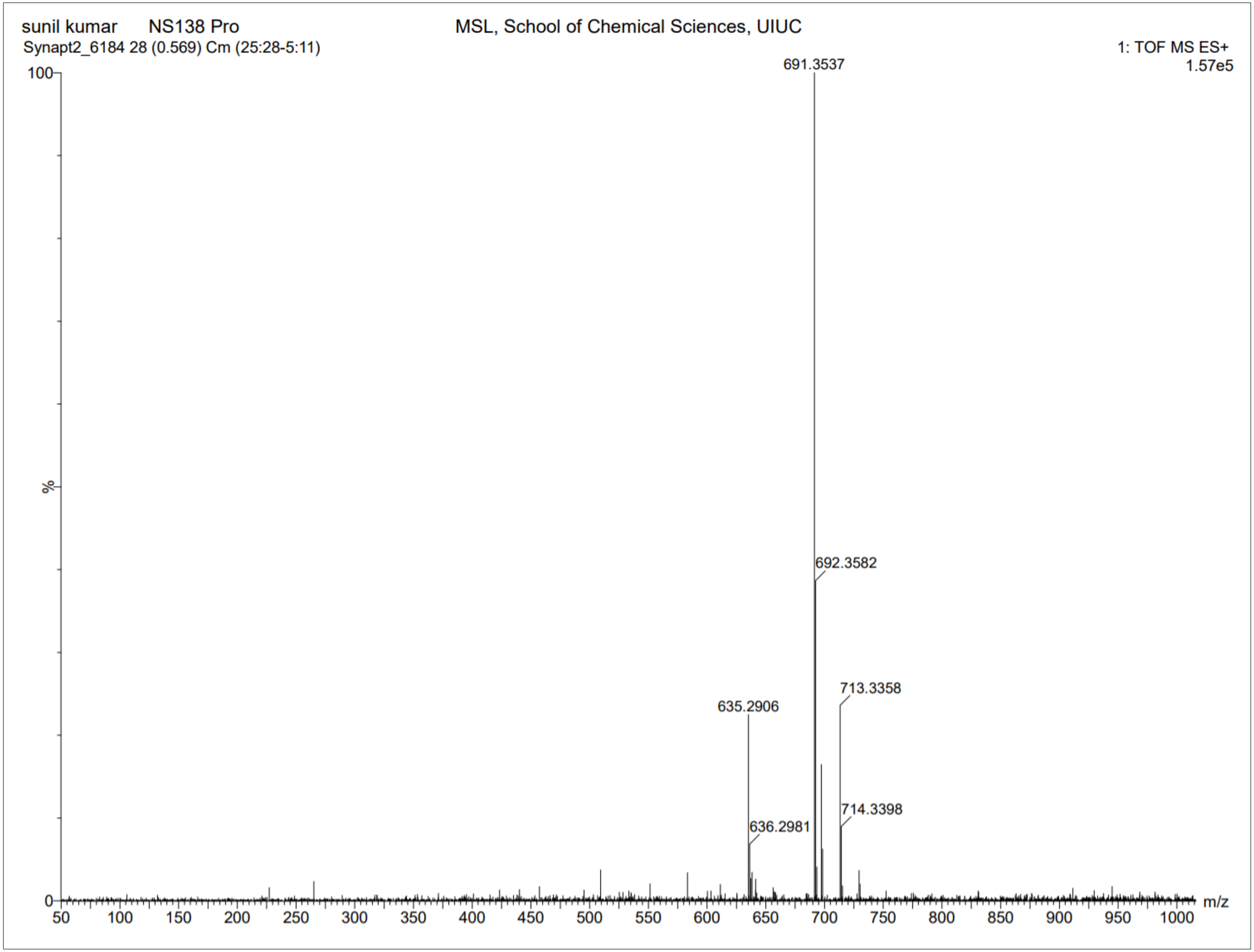

**Supplementary Fig. 102. ¹H-NMR of NS138 Dep**

$^1$H NMR (500 MHz, DMSO) δ 0.94 – 1.00 (d, *J* = 6.6 Hz, 5H), 1.58 – 1.86 (m, 6H), 1.10 – 1.29 (m, 5H), 1.99 – 2.07 (dq, *J* = 14.2, 7.7, 7.2 Hz, 1H), 2.25 – 2.40 (s, 3H), 3.59 – 3.65 (t, *J* = 6.3 Hz, 2H), 4.94 – 4.98 (s, 2H), 6.69 – 6.75 (t, *J* = 5.7 Hz, 1H), 6.91 – 6.96 (d, *J* = 7.9 Hz, 1H), 7.36 – 7.46 (dd, *J* = 22.4, 8.1 Hz, 2H), 7.90 – 7.94 (d, *J* = 7.8 Hz, 1H), 8.49 – 8.56 (t, *J* = 5.9 Hz, 1H), 8.60 – 8.63 (d, *J* = 5.0 Hz, 1H), 8.63 – 8.66 (d, *J* = 5.5 Hz, 1H), 9.77 – 9.80 (s, 1H), 10.32 – 10.37 (s, 1H). HRMS (m/z): [M]+ calcd. for $C_{31}H_{38}N_8O_7$, 635.2936; found, 635.2932.

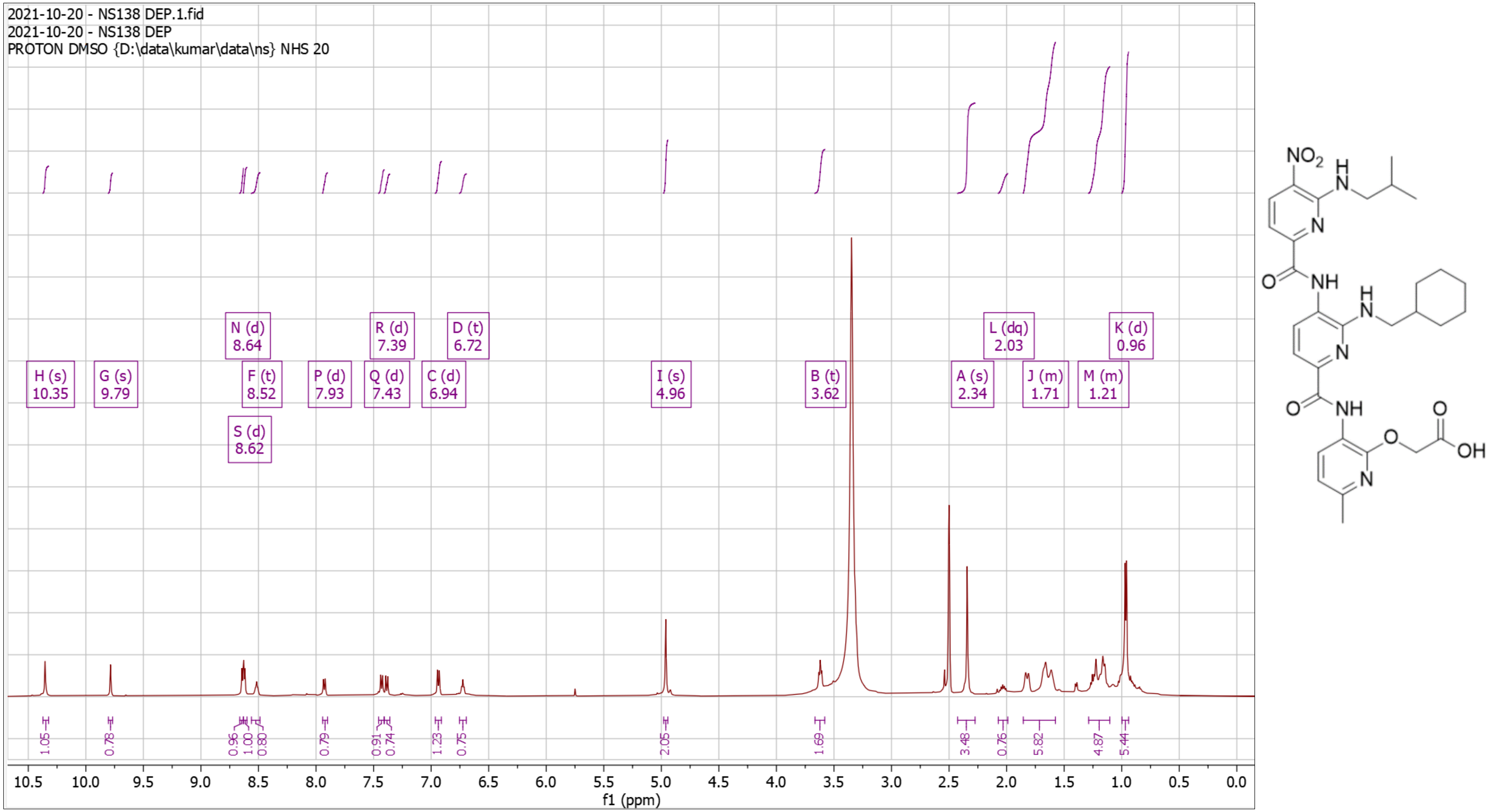

**Supplementary Fig. 103. High Resolution Mass Spectrum for NS138 Dep**

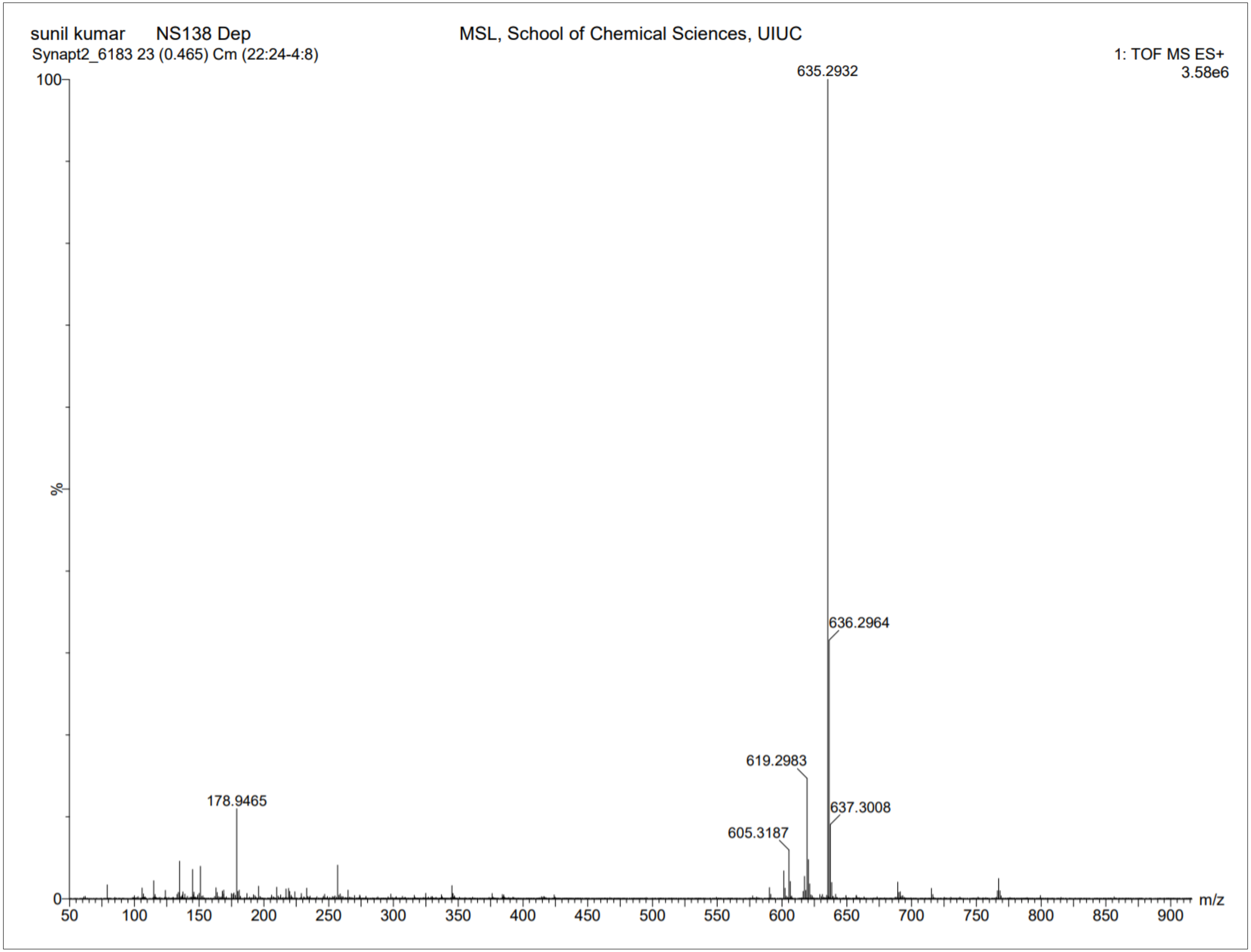

**Supplementary Fig. 104. $^{1}$H-NMR of NS165 Pro**

$^{1}$H NMR (500 MHz, CDCl3) δ 0.85 – 0.92 (m, 3H), 1.00 – 1.38 (m, 8H), 1.44 – 1.46 (s, 9H), 1.64 – 1.92 (m, 8H), 2.35 – 2.40 (s, 3H), 2.93 – 3.01 (t, J = 7.7 Hz, 3H), 3.38 – 3.51 (t, J = 6.2 Hz, 2H), 3.59 – 3.72 (td, J = 7.0, 5.2 Hz, 2H), 4.81 – 4.87 (s, 2H), 4.87 – 4.95 (t, J = 5.7 Hz, 1H), 6.72 – 6.83 (d, J = 7.9 Hz, 1H), 7.52 – 7.62 (d, J = 8.4 Hz, 1H), 7.62 – 7.69 (d, J = 7.8 Hz, 1H), 7.86 – 7.97 (d, J = 7.8 Hz, 1H), 8.20 – 8.32 (t, J = 5.3 Hz, 1H), 8.58 – 8.67 (d, J = 8.4 Hz, 1H), 8.67 – 8.74 (d, J = 7.9 Hz, 1H), 9.32 – 9.42 (s, 1H), 10.30 – 10.40 (s, 1H). HRMS (m/z): [M]+ calcd. for $C_{37}H_{50}N_8O_7$, 719.3875; found, 719.3874.

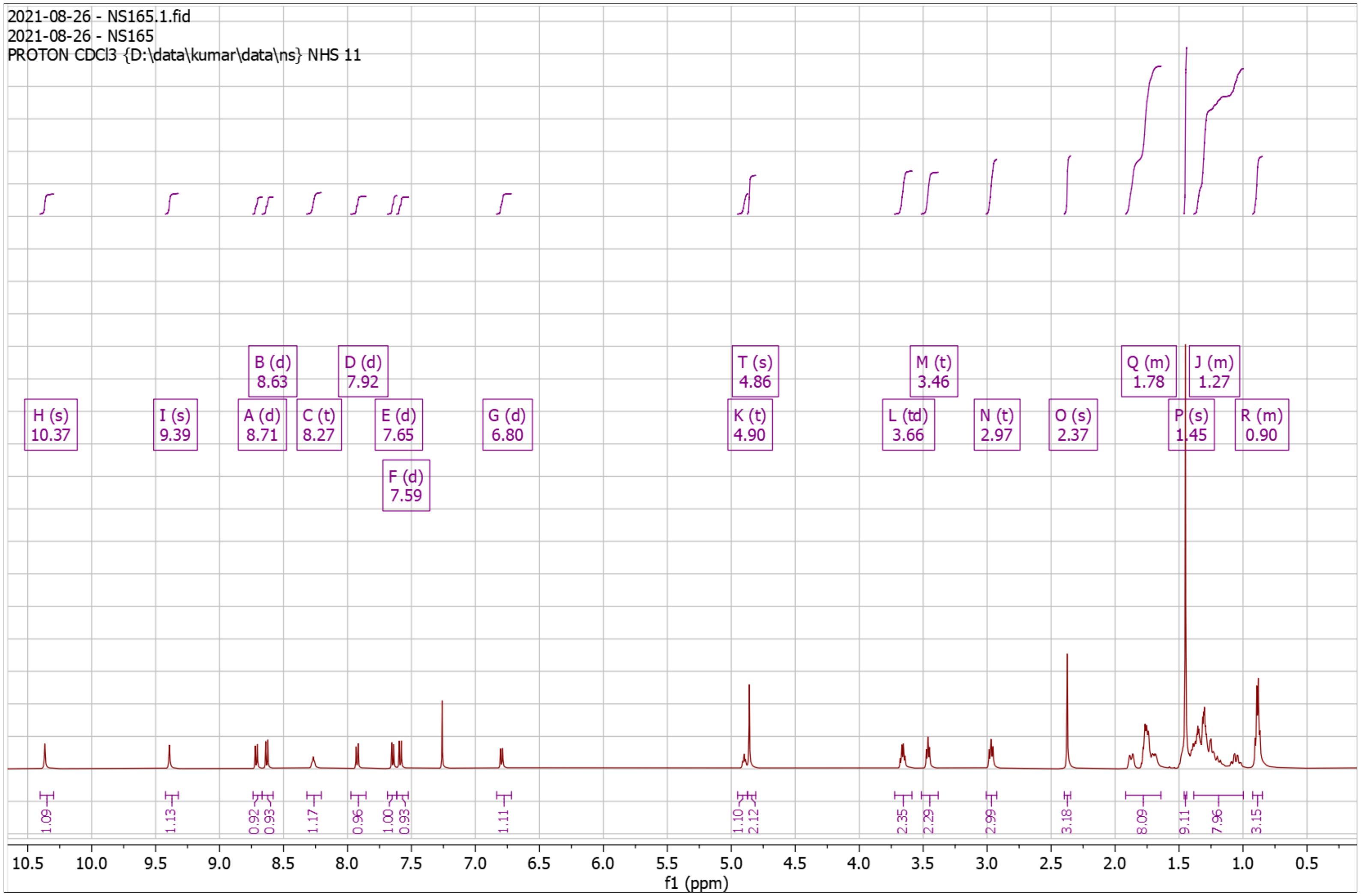


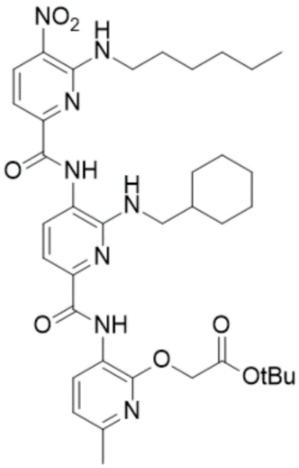

**Supplementary Fig. 105. High Resolution Mass Spectrum for NS165 Pro**

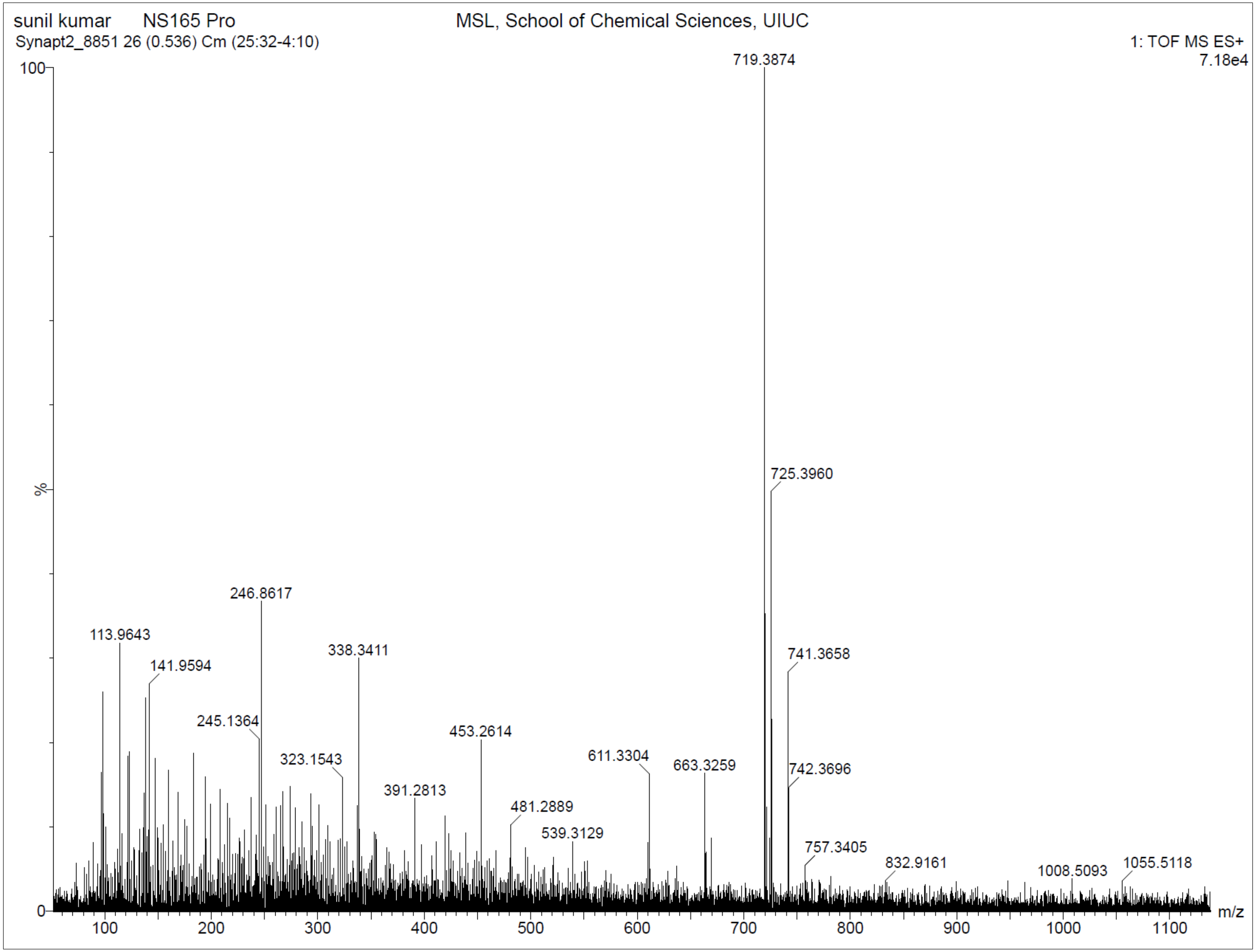

**Supplementary Fig. 106. $^{1}$H-NMR of NS165 Dep**

$^{1}$H NMR (500 MHz, DMSO) δ 0.76 – 0.84 (t, J = 6.9 Hz, 3H), 0.92 – 1.41 (m, 11H), 1.57 – 1.87 (m, 8H), 2.29 – 2.39 (s, 3H), 3.29 – 3.33 (t, 2H), 3.70 – 3.81 (q, J = 6.6 Hz, 2H), 4.93 – 5.02 (s, 2H), 6.73 – 6.80 (t, J = 5.7 Hz, 1H), 6.91 – 6.96 (d, J = 7.9 Hz, 1H), 7.34 – 7.41 (d, J = 8.4 Hz, 1H), 7.41 – 7.46 (d, J = 7.8 Hz, 1H), 7.94 – 7.99 (d, J = 7.8 Hz, 1H), 8.45 – 8.57 (t, J = 5.7 Hz, 1H), 8.57 – 8.69 (dd, J = 8.2, 3.0 Hz, 2H), 9.75 – 9.82 (s, 1H), 10.29 – 10.43 (s, 1H). HRMS (m/z): [M]+ calcd. for $C_{33}H_{42}N_8O_7$, 663.3249; found, 663.3253.

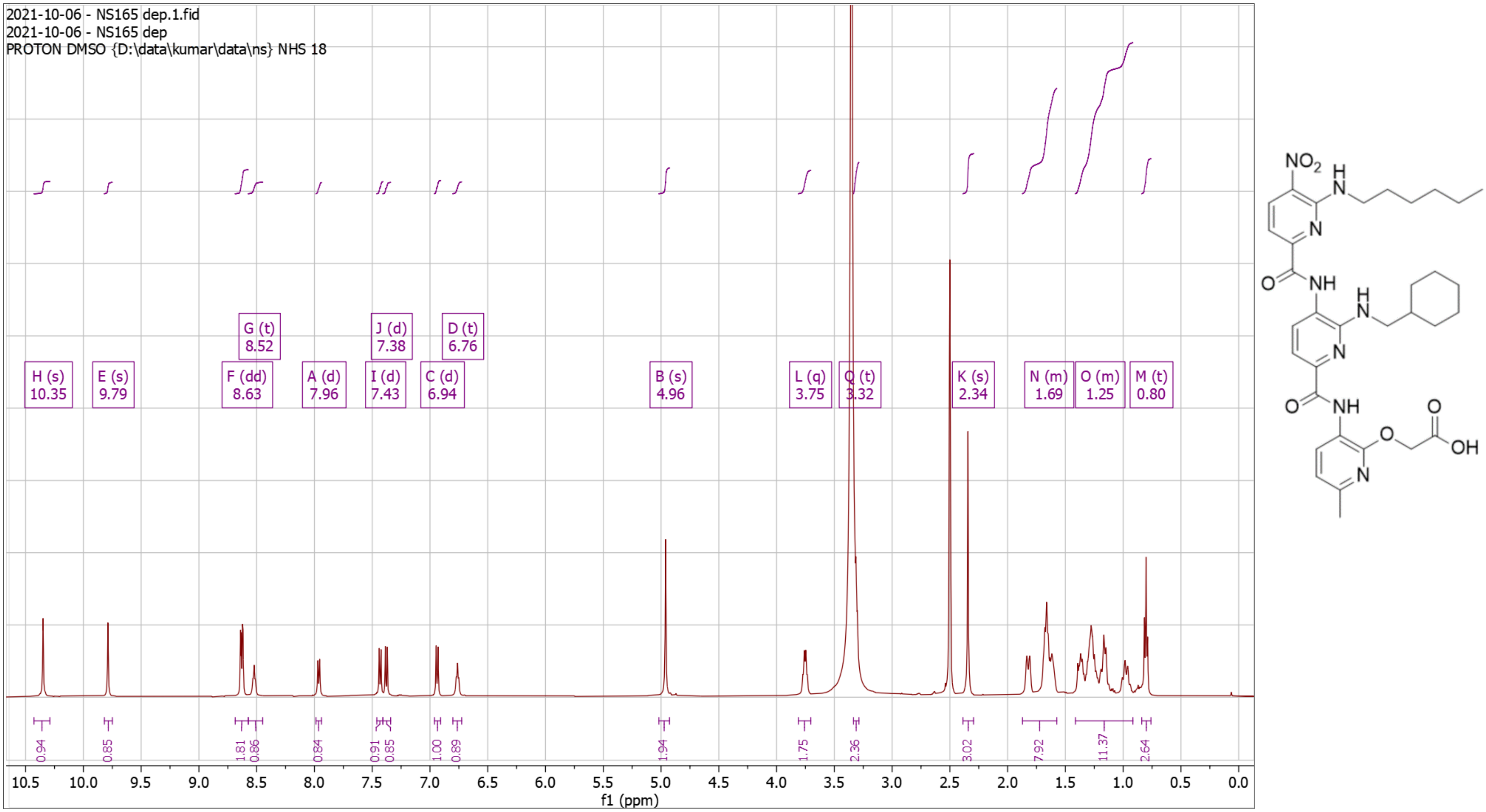

**Supplementary Fig. 107. High Resolution Mass Spectrum for NS165 Dep**

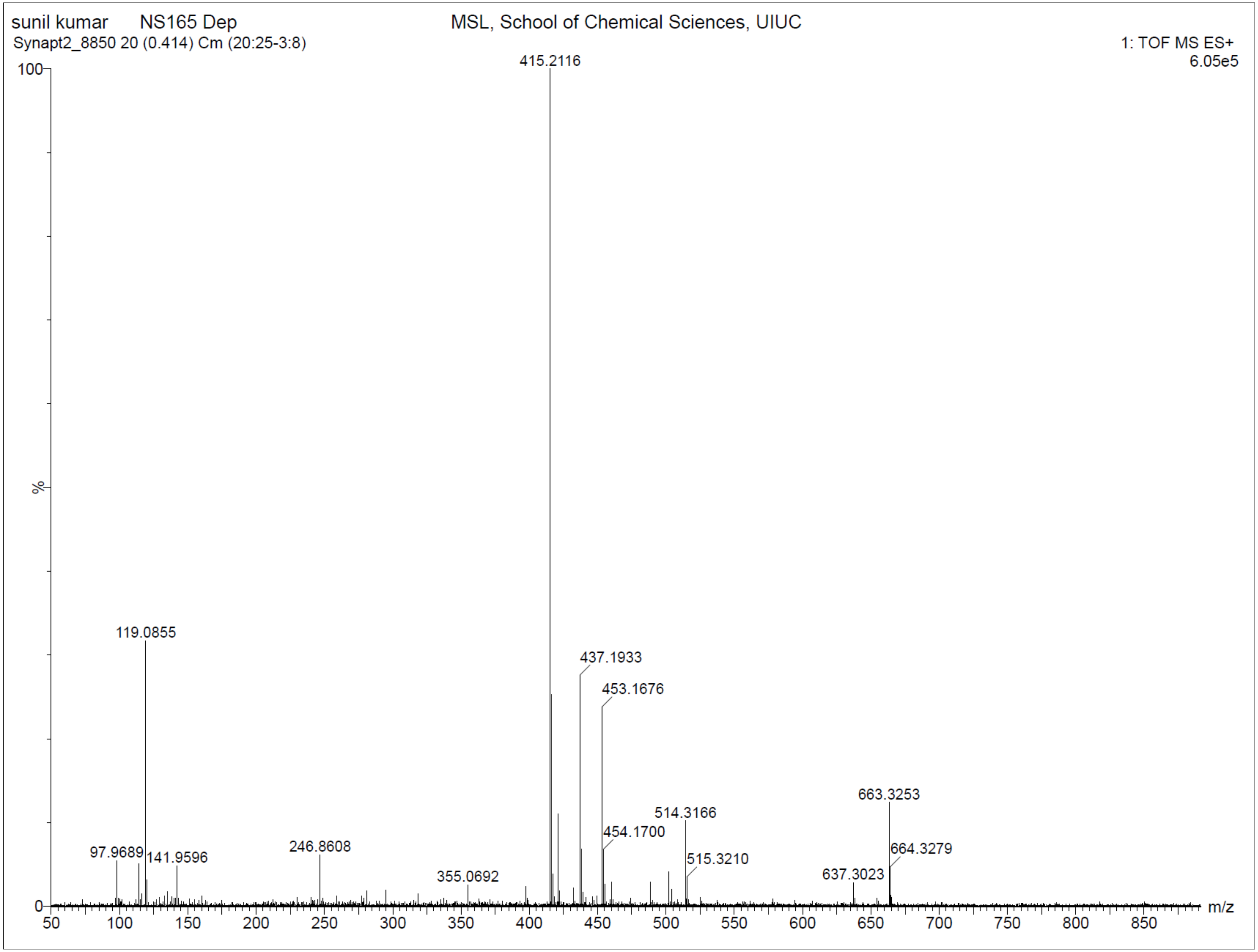

**Supplementary Fig. 108. $^{1}$H-NMR of NS166 Pro**

$^{1}$H NMR (500 MHz, $CDCl_3$) δ 0.87 – 1.35 (m, 5H), 1.44 – 1.48 (s, 9H), 1.65 – 1.92 (m, 6H), 2.33 – 2.41 (s, 3H), 3.38 – 3.47 (t, *J* = 6.2 Hz, 2H), 3.86 – 3.94 (q, *J* = 5.4 Hz, 2H), 3.97 – 4.04 (t, *J* = 4.9 Hz, 2H), 4.82 – 4.89 (s, 2H), 5.23 – 5.29 (t, *J* = 5.5 Hz, 1H), 6.72 – 6.80 (d, *J* = 7.9 Hz, 1H), 7.56 – 7.64 (d, *J* = 8.2 Hz, 2H), 8.06 – 8.16 (d, *J* = 7.8 Hz, 1H), 8.47 – 8.51 (t, *J* = 5.6 Hz, 1H), 8.54 – 8.62 (d, *J* = 8.4 Hz, 1H), 8.63 – 8.71 (d, *J* = 7.8 Hz, 1H), 9.45 – 9.53 (s, 1H), 10.26 – 10.33 (s, 1H). HRMS (m/z): [M]+ calcd. for $C_{33}H_{42}N_8O_8$, 679.3198; found, 679.3203.

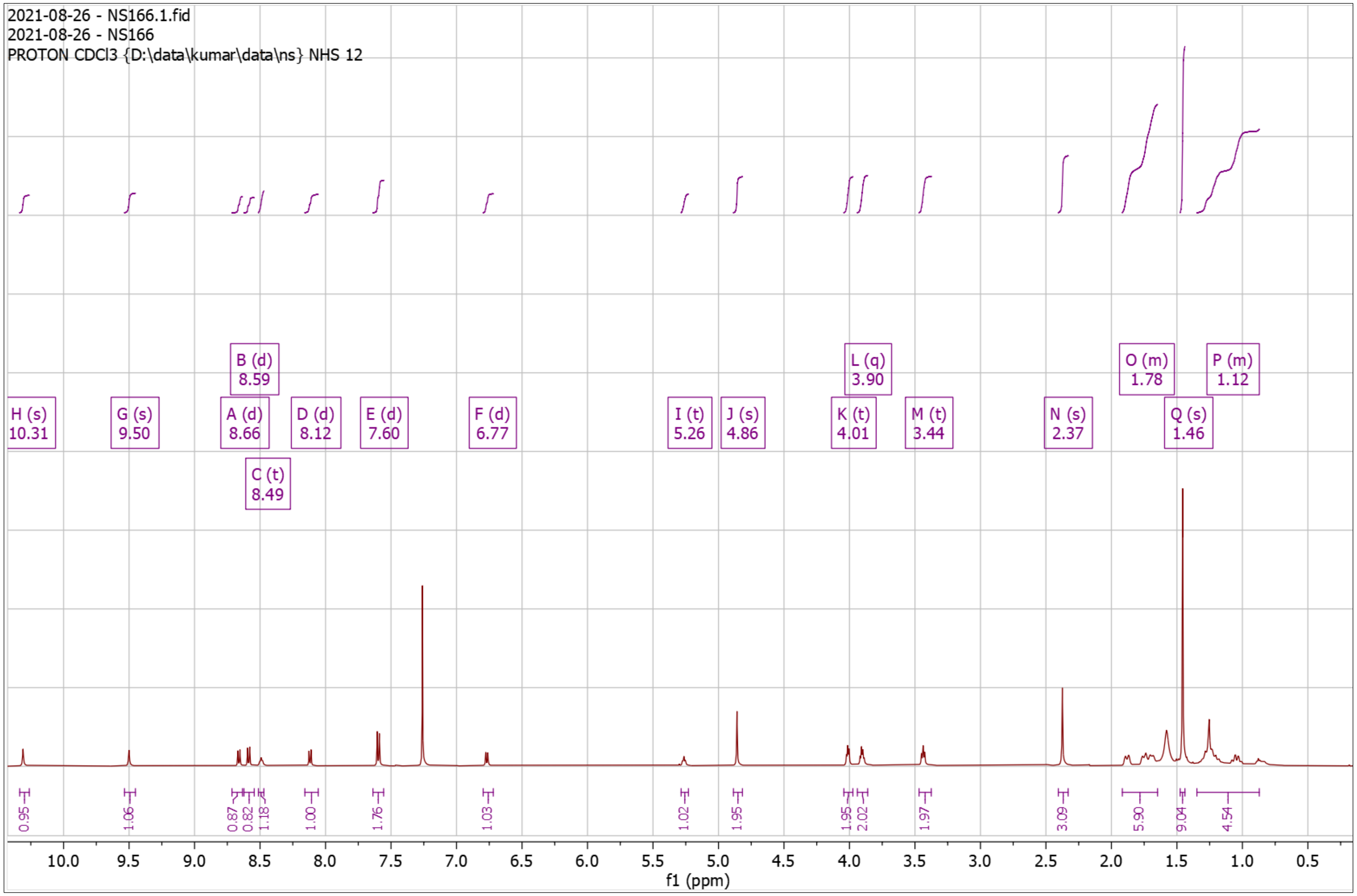


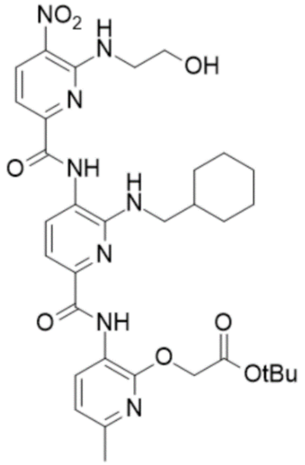

**Supplementary Fig. 109. High Resolution Mass Spectrum for NS166 Pro**

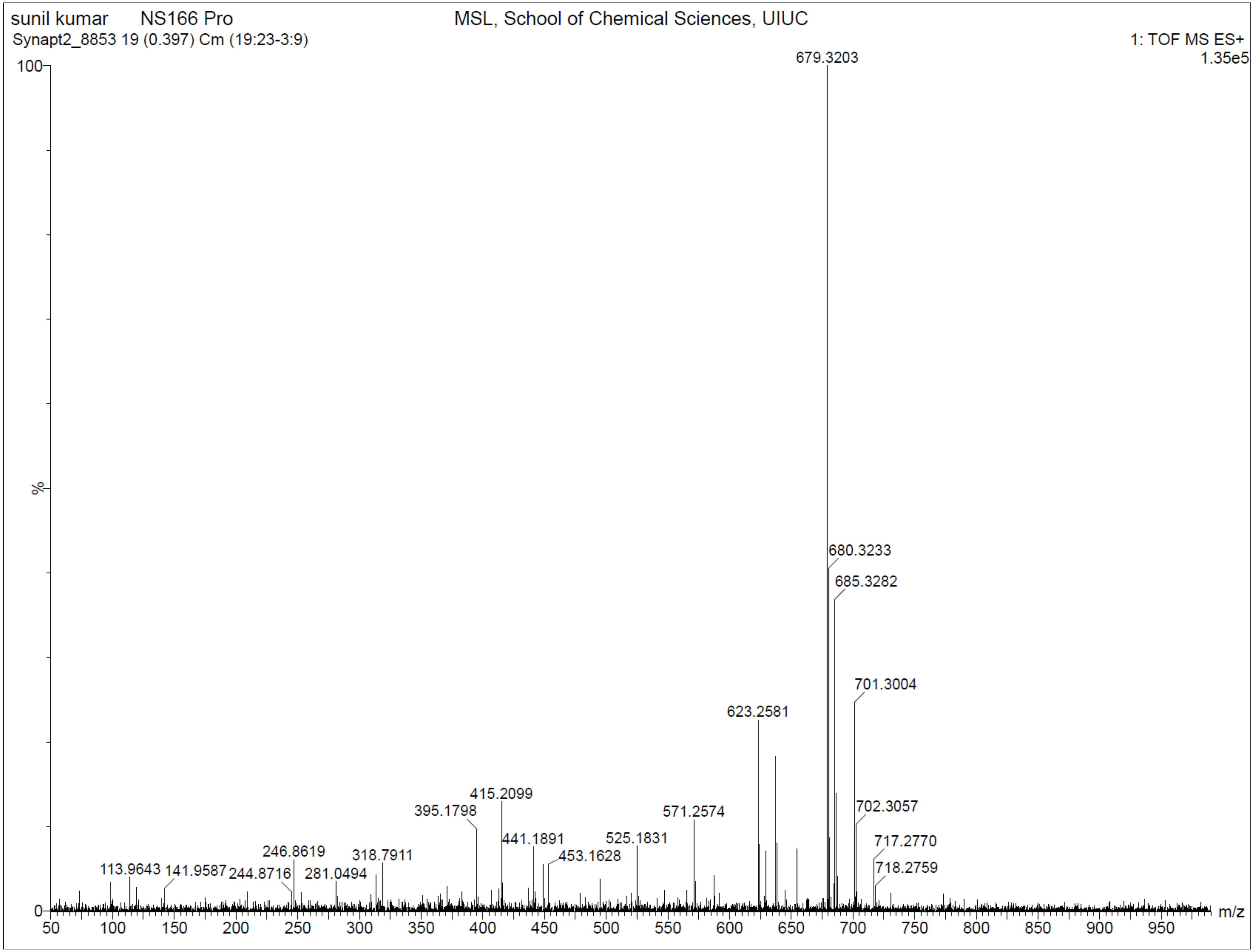

**Supplementary Fig. 110. $^{1}$H-NMR of NS166 Dep**

$^{1}$H NMR (500 MHz, DMSO) δ 0.90 – 1.25 (dt, $J$ = 96.3, 12.3 Hz, 5H), 1.54 – 1.85 (m, 6H), 2.29 – 2.32 (s, 3H), 3.27 – 3.32 (t, $J$ = 6.1 Hz, 2H), 3.64 – 3.70 (t, $J$ = 5.5 Hz, 2H), 3.83 – 3.90 (q, $J$ = 5.5 Hz, 2H), 4.75 – 4.79 (s, 2H), 6.65 – 6.75 (t, $J$ = 5.6 Hz, 1H), 6.78 – 6.88 (d, $J$ = 7.9 Hz, 1H), 7.33 – 7.47 (dd, $J$ = 8.1, 5.5 Hz, 2H), 7.88 – 7.94 (d, $J$ = 7.8 Hz, 1H), 8.52 – 8.60 (m, 2H), 8.60 – 8.70 (d, $J$ = 8.4 Hz, 1H), 9.89 – 10.03 (s, 1H), 10.34 – 10.48 (s, 1H). HRMS (m/z): [M]+ calcd. for $C_{29}H_{34}N_{8}O_{8}$, 623.2572; found, 623.2581.

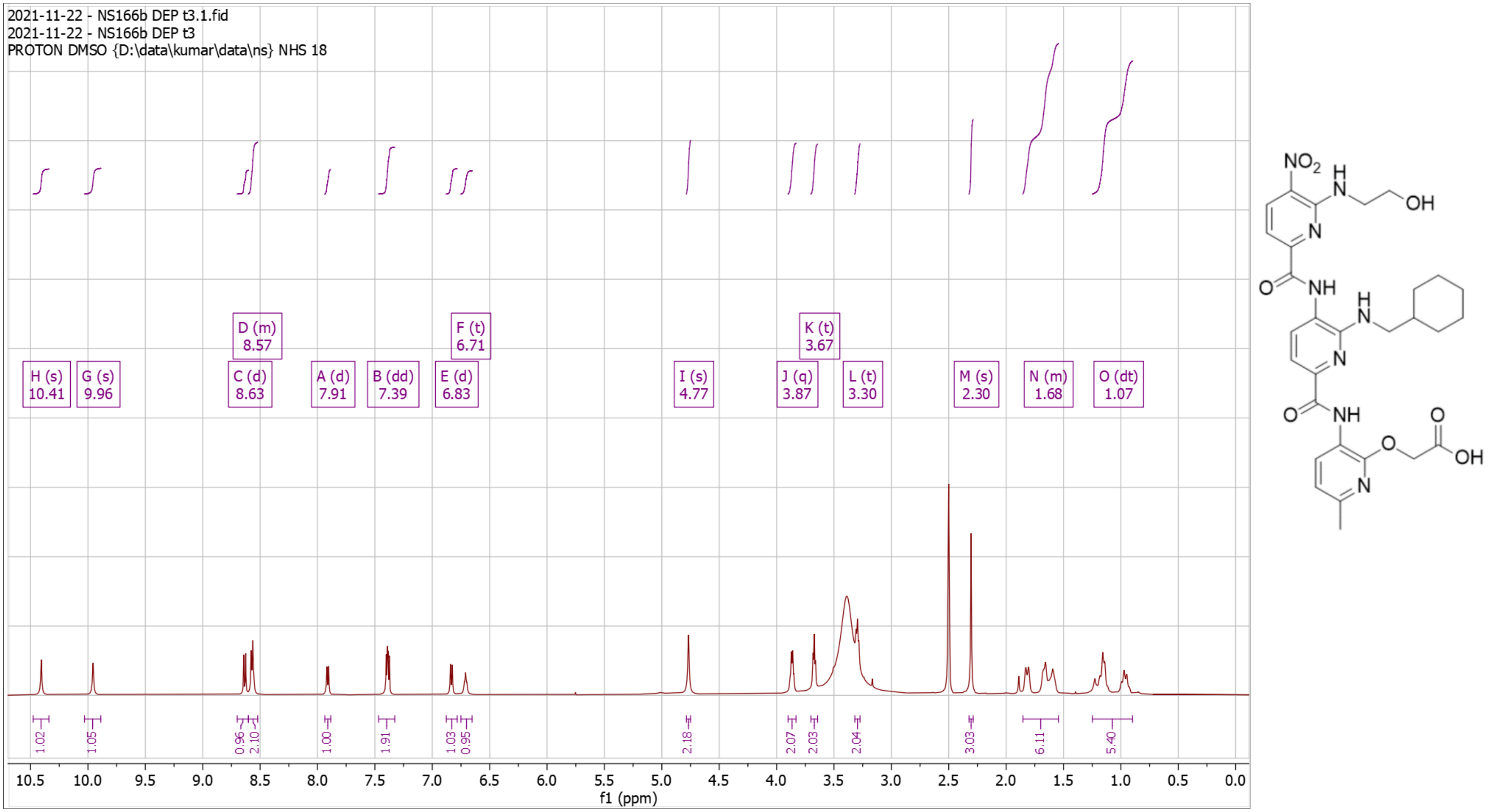

**Supplementary Fig. 111. High Resolution Mass Spectrum for NS166 Dep**

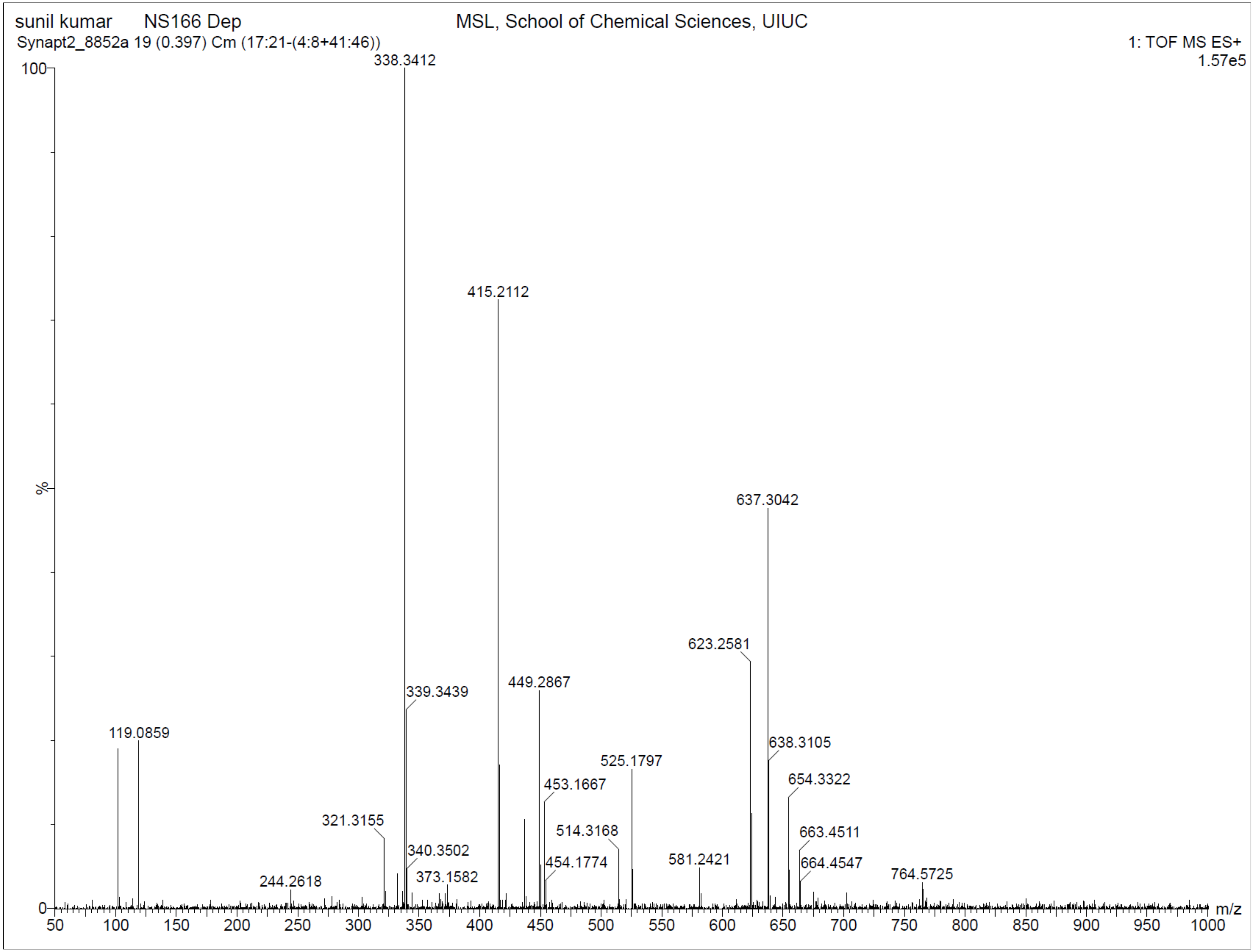

**Supplementary Fig. 112. $^{1}$H-NMR of NS168 Pro**

$^{1}$H NMR (500 MHz, $CDCl_3$) δ 0.85 – 1.30 (m, 5H), 1.45 – 1.49 (s, 9H), 1.53 – 1.90 (m, 6H), 2.31 – 2.44 (s, 3H), 2.89 – 3.05 (t, *J* = 7.0 Hz, 2H), 3.36 – 3.47 (t, *J* = 6.2 Hz, 2H), 3.83 – 3.95 (q, *J* = 6.6 Hz, 2H), 4.80 – 4.94 (m, 3H), 5.32 – 5.45 (s, 1H), 6.66 – 6.76 (d, *J* = 8.3 Hz, 2H), 6.76 – 6.83 (d, *J* = 7.9 Hz, 1H), 7.04 – 7.10 (d, *J* = 8.1 Hz, 2H), 7.54 – 7.61 (d, *J* = 8.3 Hz, 1H), 7.60 – 7.68 (d, *J* = 7.8 Hz, 1H), 7.84 – 7.93 (d, *J* = 7.8 Hz, 1H), 8.25 – 8.36 (t, *J* = 5.4 Hz, 1H), 8.56 – 8.65 (d, *J* = 8.3 Hz, 1H), 8.68 – 8.74 (d, *J* = 7.9 Hz, 1H), 9.27 – 9.41 (s, 1H), 10.35 – 10.42 (s, 1H). HRMS (m/z): [M]+ calcd. for $C_{39}H_{46}N_8O_8$, 755.3511; found, 755.3497.

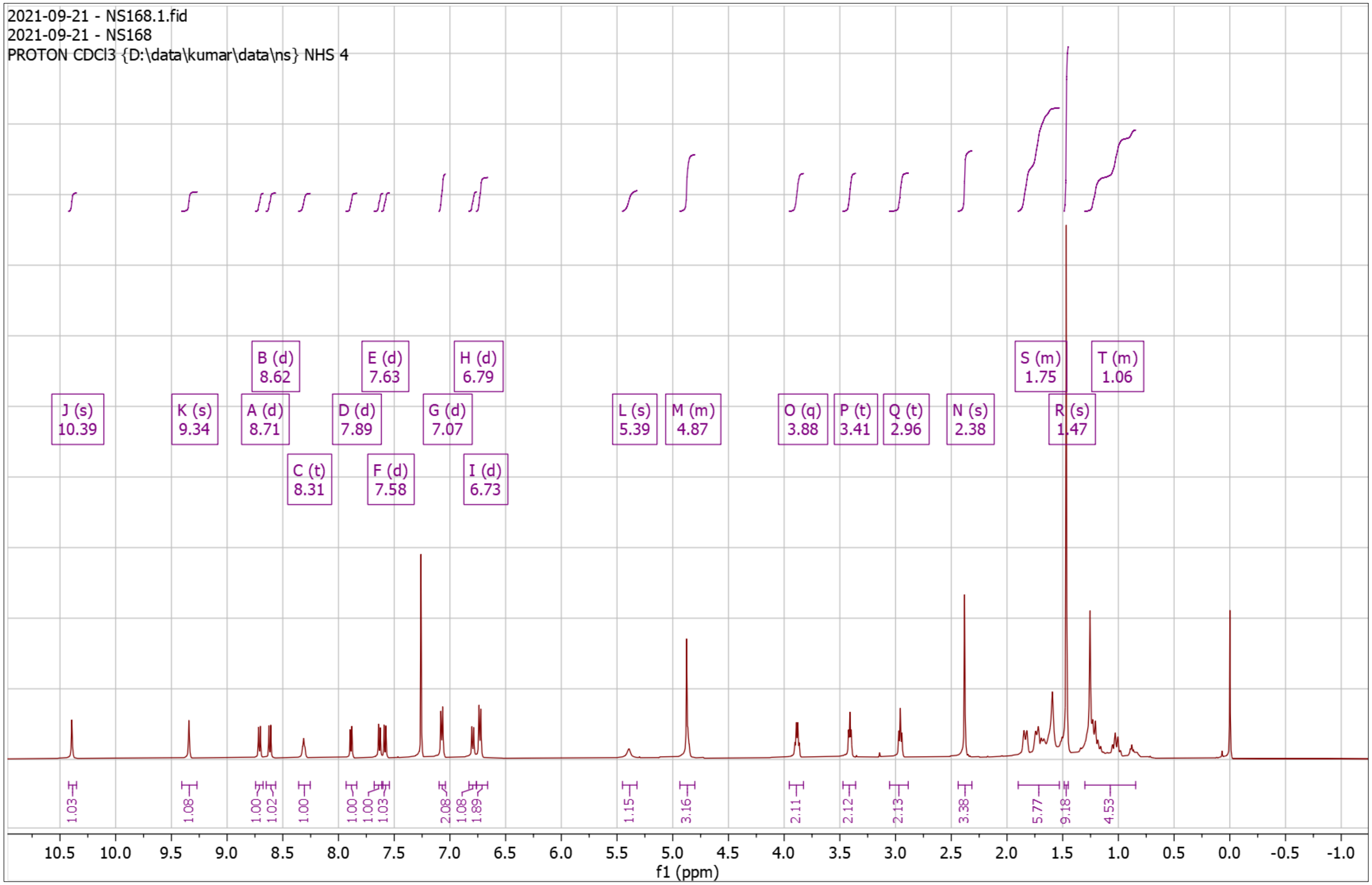


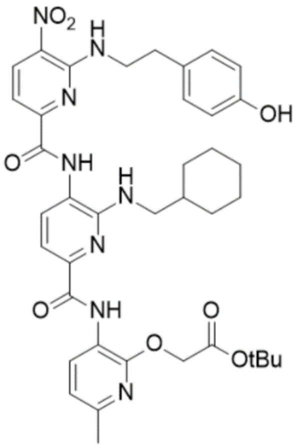

**Supplementary Fig. 113. High Resolution Mass Spectrum for NS168 Pro**

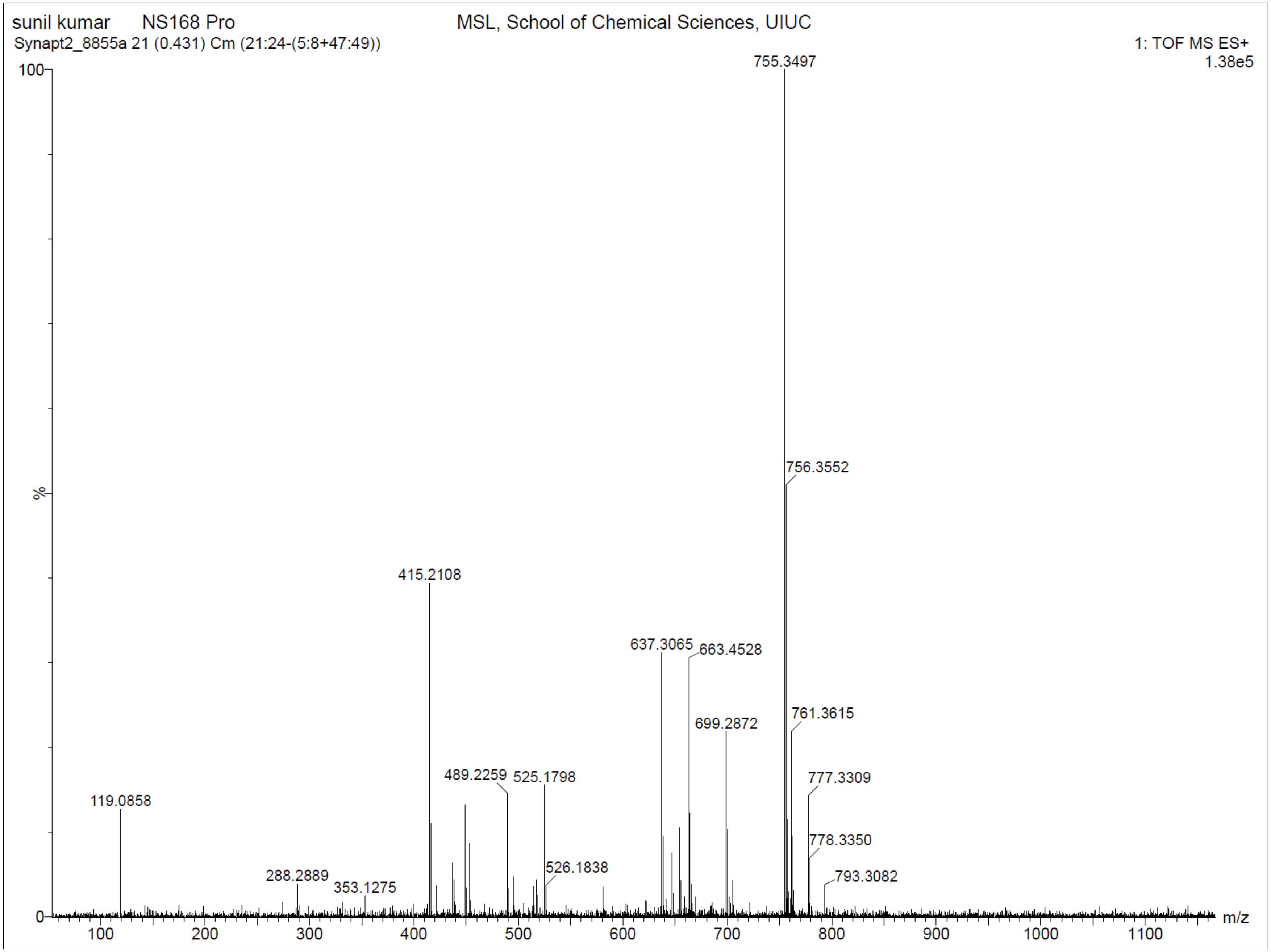

**Supplementary Fig. 114. $^{1}$H-NMR of NS168 Dep**

$^{1}$H NMR (500 MHz, DMSO) δ 0.77 – 1.21 (m, 5H), 1.49 – 1.90 (m, 6H), 2.33 – 2.37 (s, 3H), 2.82 – 2.89 (t, *J* = 7.1 Hz, 2H), 3.27 – 3.31 (t, *J* = 6.4 Hz, 2H), 3.92 – 3.99 (q, *J* = 6.7 Hz, 2H), 4.96 – 4.98 (s, 2H), 6.60 – 6.65 (d, *J* = 8.3 Hz, 2H), 6.73 – 6.79 (t, *J* = 5.7 Hz, 1H), 6.92 – 6.97 (d, *J* = 7.9 Hz, 1H), 7.04 – 7.09 (d, *J* = 8.1 Hz, 2H), 7.35 – 7.40 (d, *J* = 8.4 Hz, 1H), 7.41 – 7.47 (d, *J* = 7.7 Hz, 1H), 7.90 – 7.95 (d, *J* = 7.8 Hz, 1H), 8.47 – 8.52 (t, *J* = 5.7 Hz, 1H), 8.60 – 8.67 (dd, *J* = 8.1, 6.3 Hz, 2H), 9.14 – 9.17 (s, 1H), 9.84 – 9.87 (s, 1H), 10.35 – 10.39 (s, 1H). HRMS (m/z): [M]+ calcd. for $C_{35}H_{38}N_{8}O_{8}$, 699.2885; found, 699.2873.

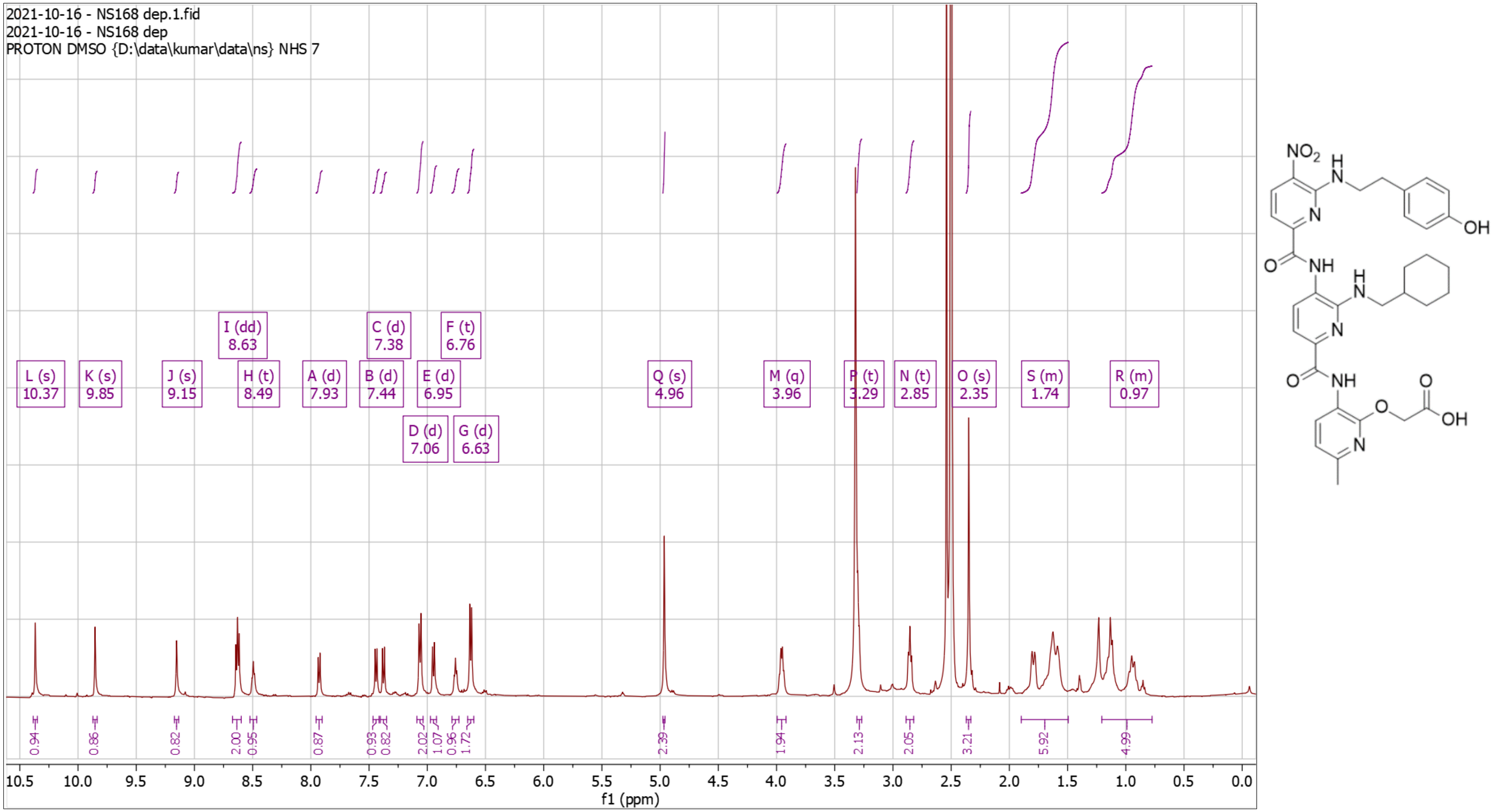

**Supplementary Fig. 115. High Resolution Mass Spectrum for NS168 Dep**

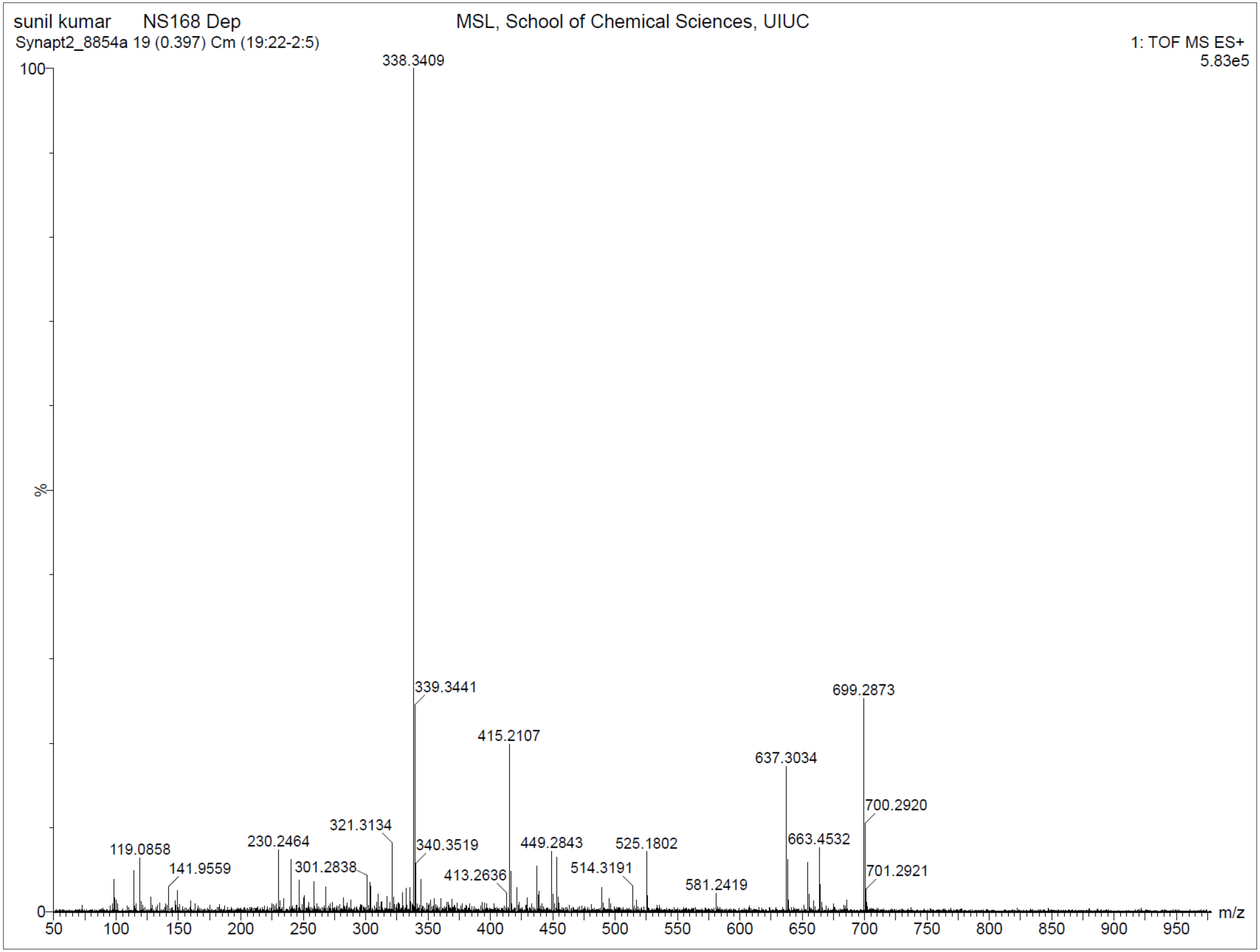

**Supplementary Fig. 116. $^1$H-NMR of NS169 Pro**

$^1$H NMR (500 MHz, CDCl3) δ 0.71 – 1.38 (m, 5H), 1.44 – 1.47 (s, 9H), 1.55 – 1.96 (m, 6H), 2.37 – 2.40 (s, 3H), 2.50 – 2.55 (s, 3H), 3.49 – 3.54 (d, J = 6.8 Hz, 2H), 4.85 – 4.89 (s, 2H), 4.93 – 4.97 (d, J = 5.6 Hz, 2H), 6.79 – 6.84 (d, J = 7.9 Hz, 1H), 7.63 – 7.69 (dd, J = 8.1, 3.4 Hz, 2H), 8.21 – 8.29 (m, 2H), 8.61 – 8.67 (m, 2H), 8.73 – 8.78 (d, J = 7.9 Hz, 1H), 8.86 – 8.92 (t, J = 5.6 Hz, 1H), 9.93 – 9.96 (s, 1H), 10.40 – 10.43 (s, 1H). HRMS (m/z): [M]+ calcd. for $C_{37}H_{44}N_{10}O_7$, 741.3467; found, 741.3457.

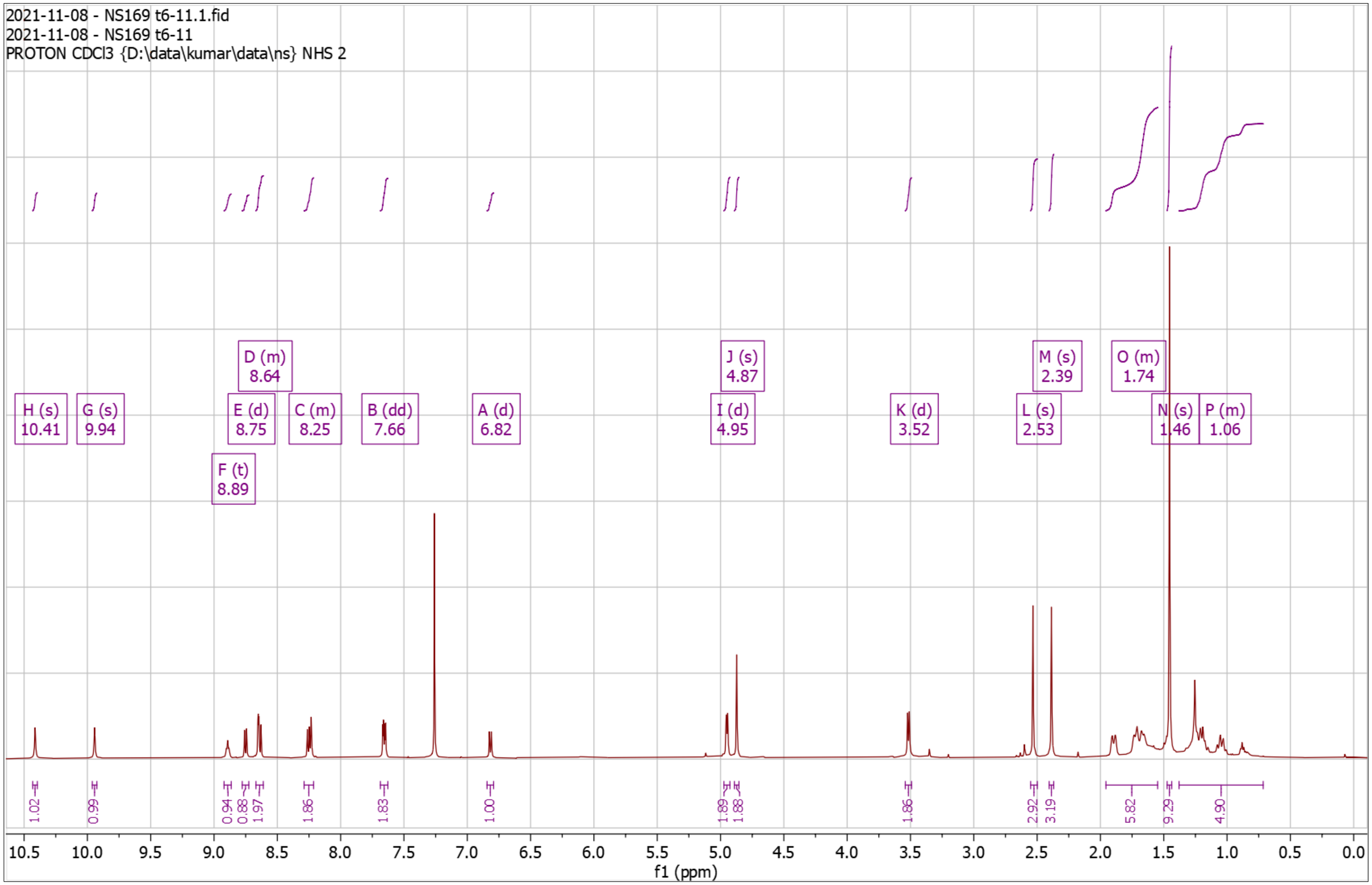


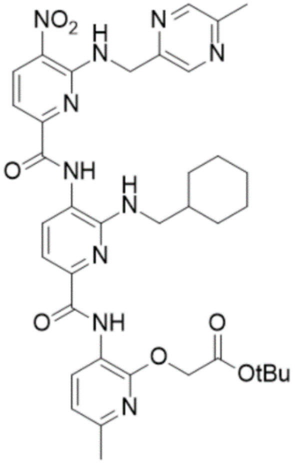

**Supplementary Fig. 117. High Resolution Mass Spectrum for NS169 Pro**

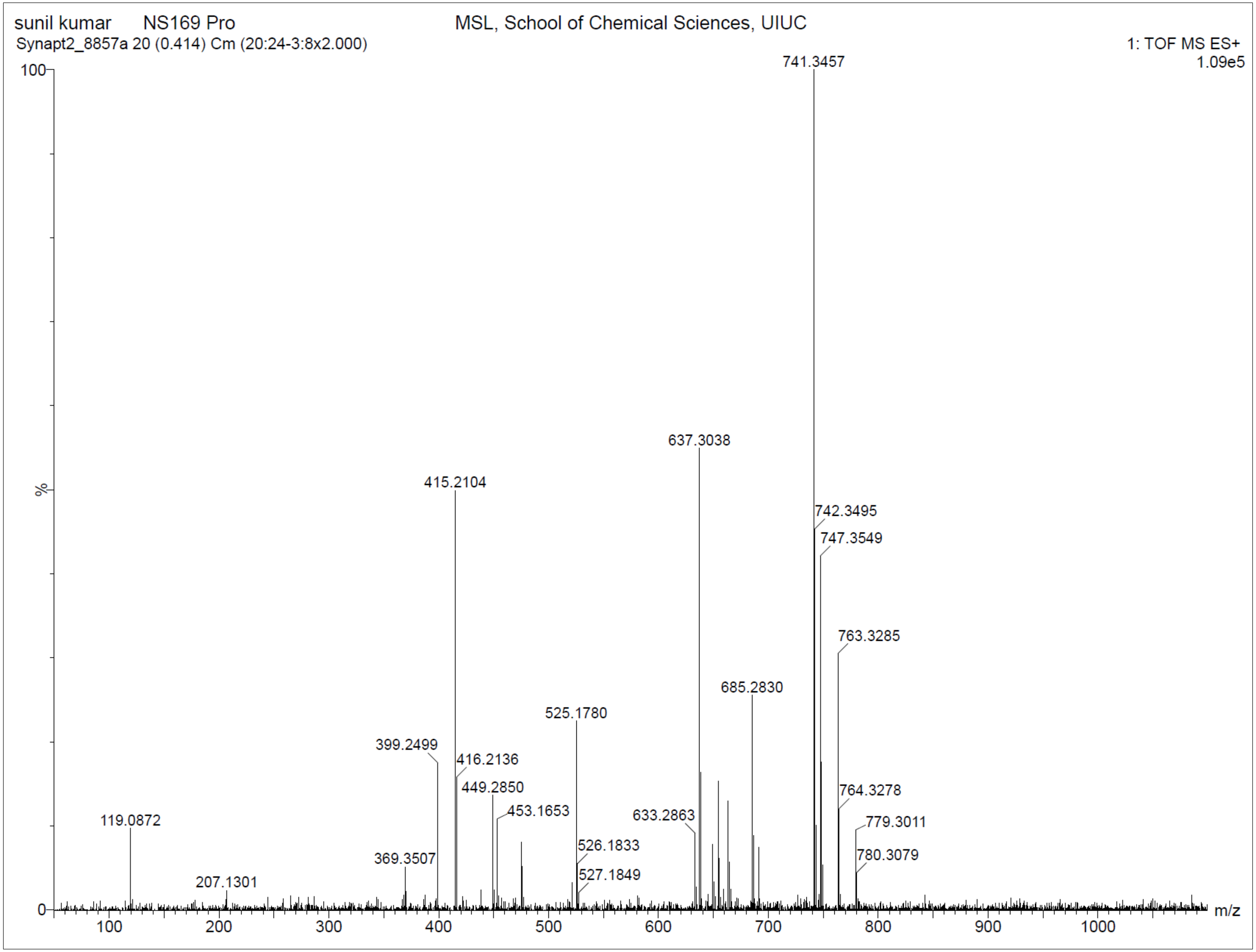

**Supplementary Fig. 118. $^{1}$H-NMR of NS169 Dep**

$^{1}$H NMR (500 MHz, DMSO) δ 0.90 – 1.19 (dd, J = 83.9, 10.2 Hz, 5H), 1.50 – 1.85 (m, 6H), 2.34 – 2.36 (s, 3H), 2.45 – 2.46 (s, 3H), 3.33 – 3.37 (d, J = 6.9 Hz, 2H), 4.92 – 4.99 (s, 2H), 5.05 – 5.16 (d, J = 5.6 Hz, 2H), 6.74 – 6.84 (t, J = 5.9 Hz, 1H), 6.91 – 6.97 (d, J = 8.0 Hz, 1H), 7.36 – 7.48 (dd, J = 8.1, 5.4 Hz, 2H), 7.87 – 7.94 (d, J = 7.8 Hz, 1H), 8.38 – 8.46 (s, 1H), 8.60 – 8.73 (m, 3H), 9.18 – 9.26 (t, J = 5.6 Hz, 1H), 9.96 – 10.05 (s, 1H), 10.31 – 10.42 (s, 1H). HRMS (m/z): [M]+ calcd. for $C_{33}H_{36}N_{10}O_7$, 685.2841; found, 685.2844.

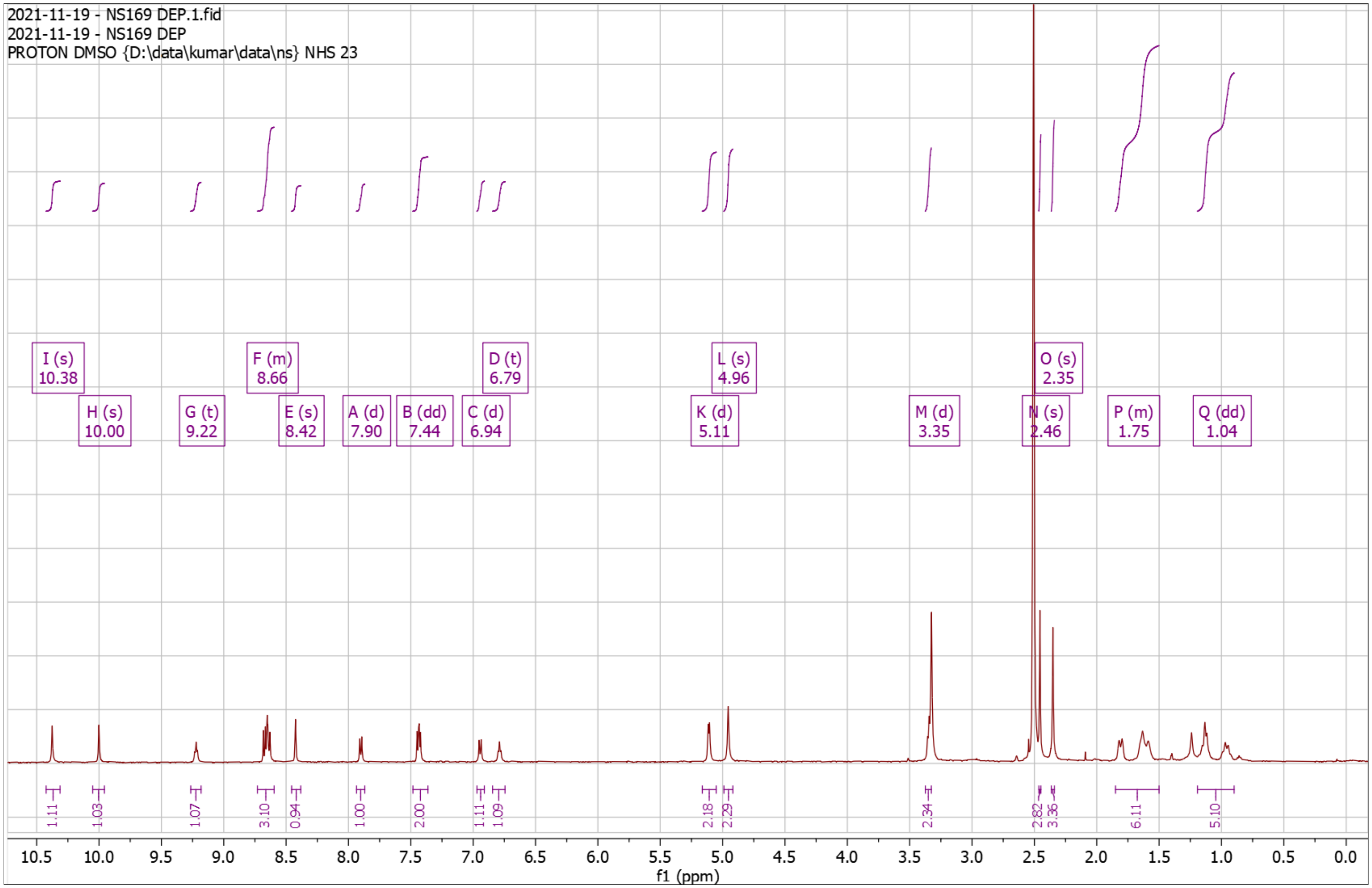


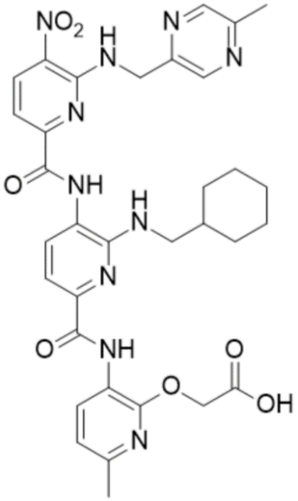

**Supplementary Fig. 119. High Resolution Mass Spectrum for NS169 Dep**

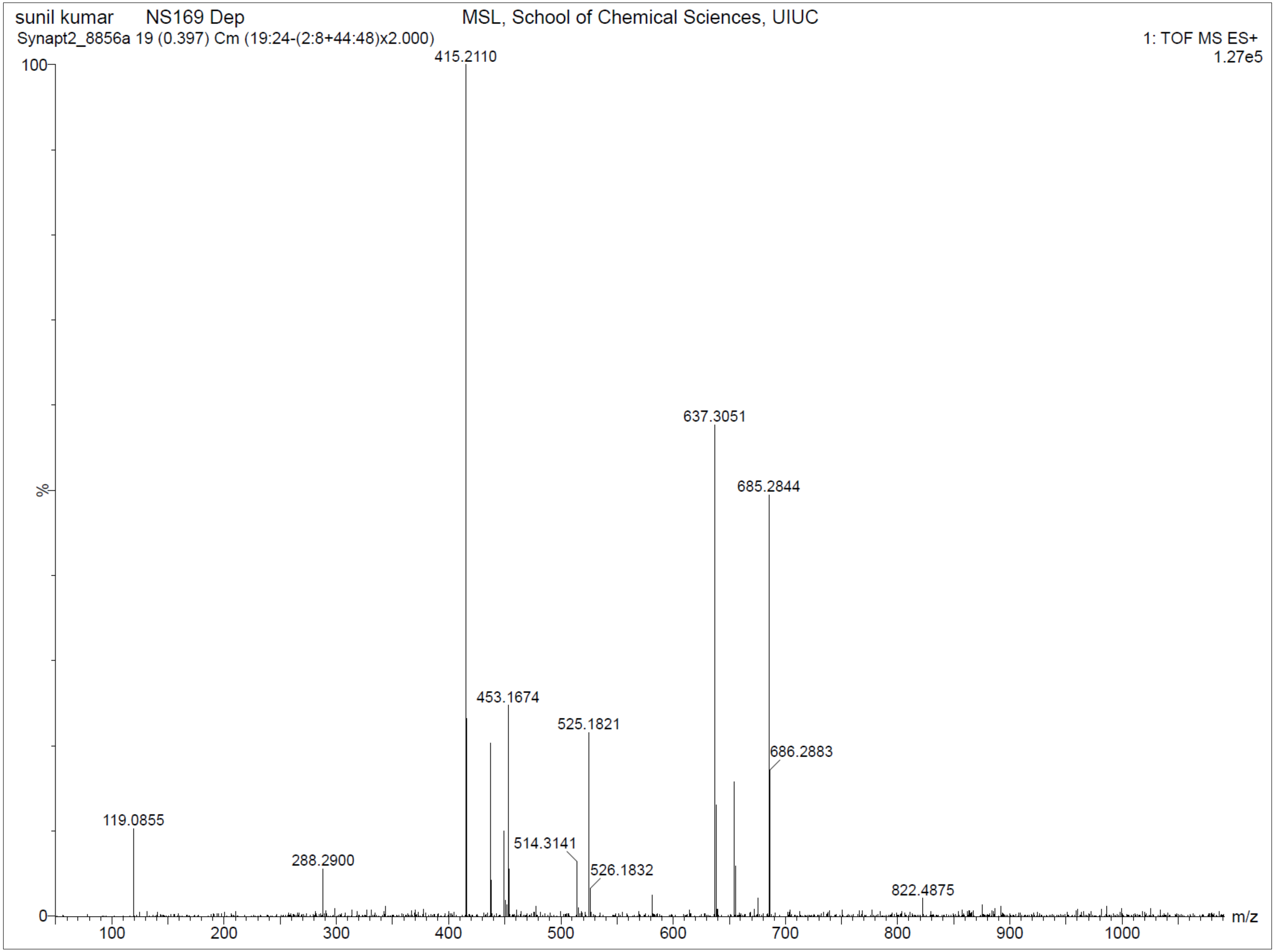

**Supplementary Fig. 120. $^{1}$H-NMR of NS174 Pro**

$^{1}$H NMR (500 MHz, CDCl3) δ 0.82 – 1.34 (m, 5H), 1.42 – 1.50 (s, 9H), 1.64 – 1.93 (m, 6H), 2.09 – 2.11 (t, J = 2.7 Hz, 1H), 2.36 – 2.39 (s, 3H), 2.64 – 2.69 (td, J = 6.6, 2.7 Hz, 2H), 3.43 – 3.52 (d, J = 5.7 Hz, 2H), 3.85 – 3.93 (q, J = 6.3 Hz, 2H), 4.84 – 4.88 (s, 2H), 4.93 – 5.00 (s, 1H), 6.77 – 6.82 (d, J = 7.9 Hz, 1H), 7.62 – 7.67 (dd, J = 8.1, 2.5 Hz, 2H), 7.91 – 7.97 (d, J = 7.9 Hz, 1H), 8.47 – 8.53 (t, J = 5.9 Hz, 1H), 8.63 – 8.68 (d, J = 8.4 Hz, 1H), 8.68 – 8.73 (d, J = 7.9 Hz, 1H), 9.36 – 9.40 (s, 1H), 10.34 – 10.38 (s, 1H). HRMS (m/z): [M]+ calcd. for $C_{35}H_{42}N_8O_7$, 687.3249; found, 687.3240.

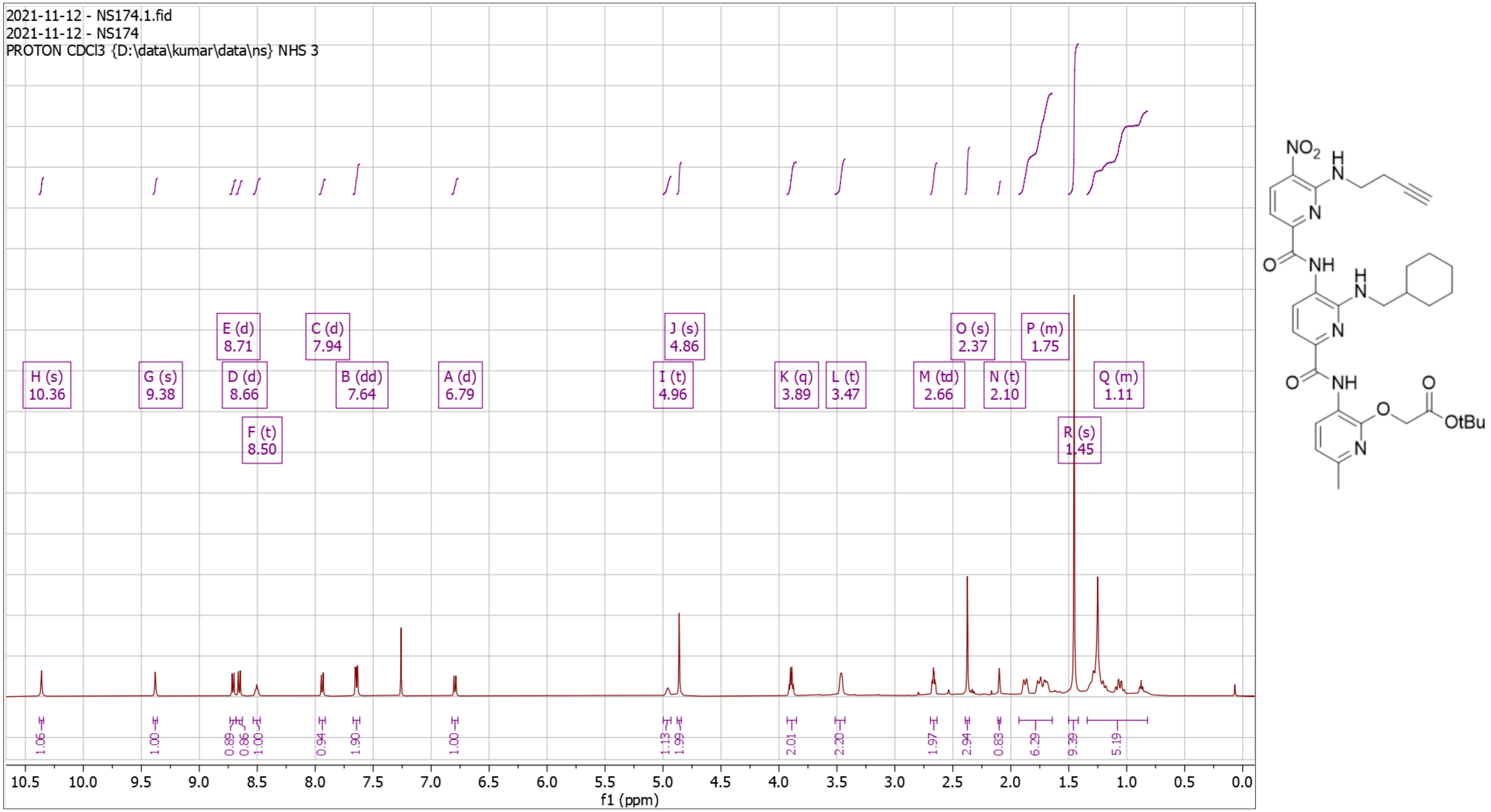

**Supplementary Fig. 121. High Resolution Mass Spectrum for NS174 Pro**

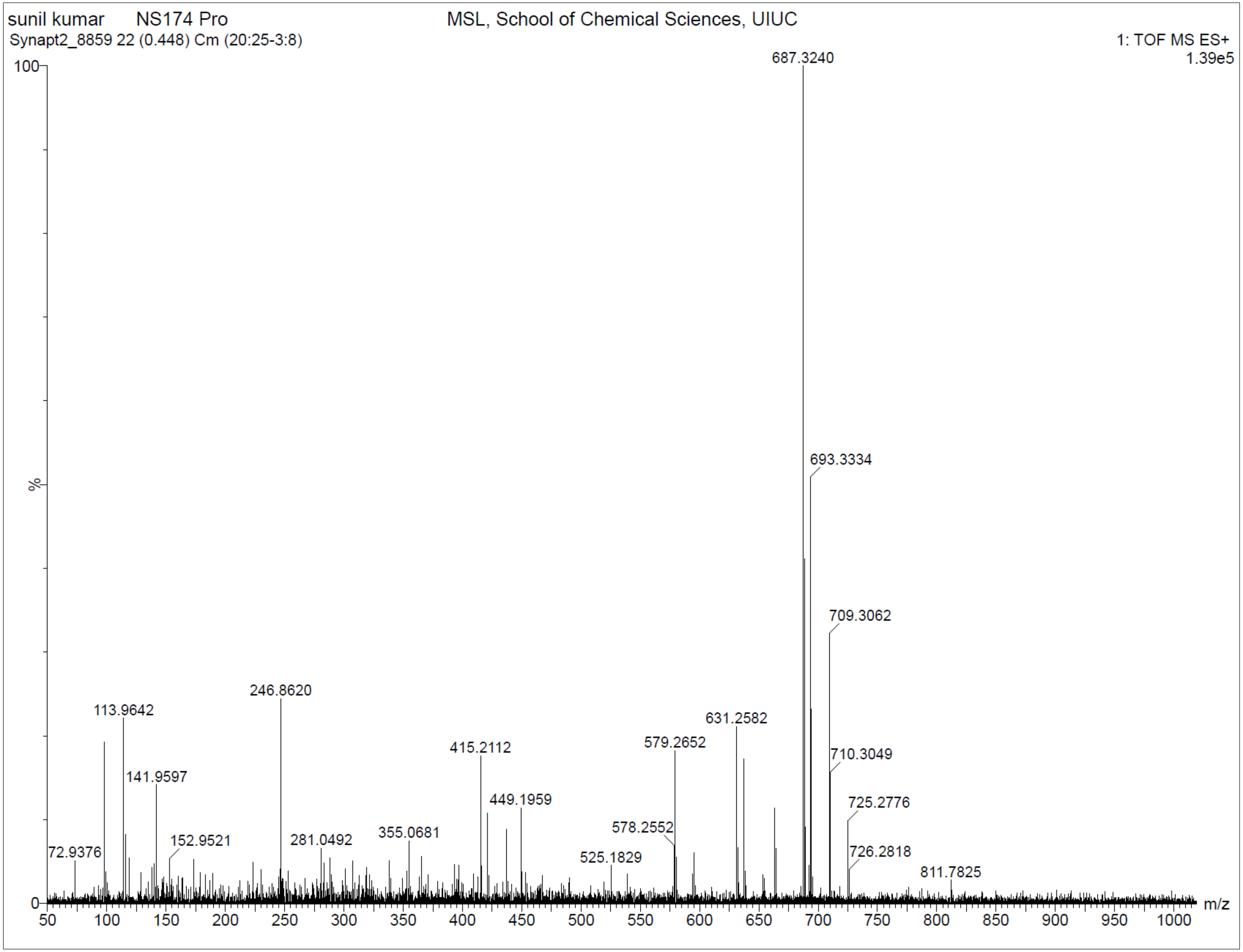

**Supplementary Fig. 122. $^{1}$H-NMR of NS174 Dep**

$^{1}$H NMR (500 MHz, DMSO) δ 0.90 – 1.28 (m, 5H), 1.55 – 1.88 (m, 6H), 2.32 – 2.36 (s, 3H), 2.57 – 2.63 (td, J = 6.8, 2.7 Hz, 2H), 2.84 – 2.91 (t, J = 2.6 Hz, 1H), 3.28 – 3.34 (t, J = 6.3 Hz, 2H), 3.86 – 3.99 (q, J = 6.6 Hz, 2H), 4.94 – 4.97 (s, 2H), 6.65 – 6.77 (t, J = 5.8 Hz, 1H), 6.88 – 6.99 (d, J = 8.0 Hz, 1H), 7.38 – 7.47 (d, J = 8.1 Hz, 2H), 7.83 – 7.91 (d, J = 7.7 Hz, 1H), 8.58 – 8.71 (m, 3H), 9.85 – 9.95 (s, 1H), 10.32 – 10.42 (s, 1H), 12.80 – 13.13 (s, 1H). HRMS (m/z): [M]+ calcd. for $C_{31}H_{34}N_8O_7$, 631.2623; found, 631.2605.

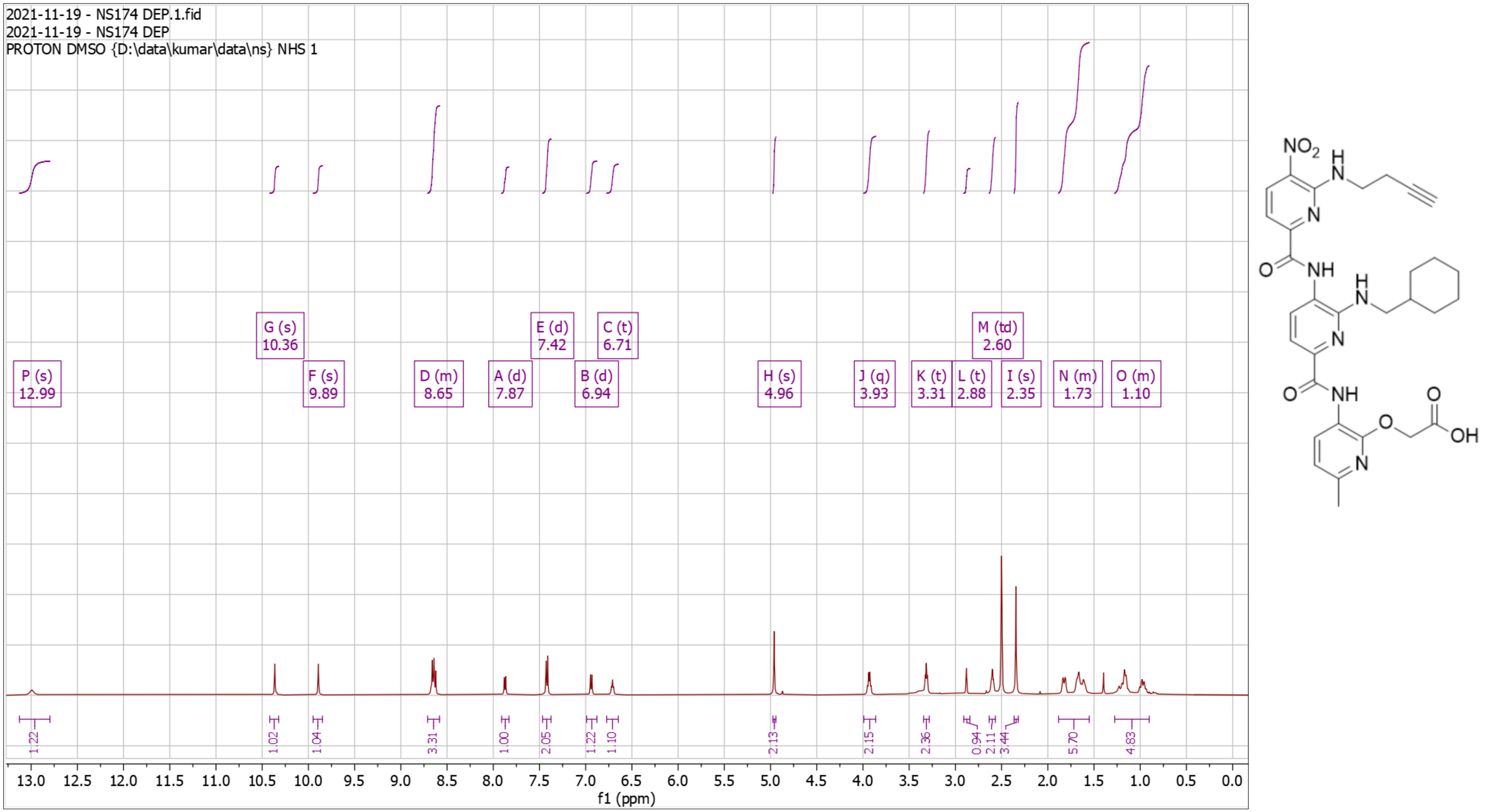

**Supplementary Fig. 123. High Resolution Mass Spectrum for NS174 Dep**

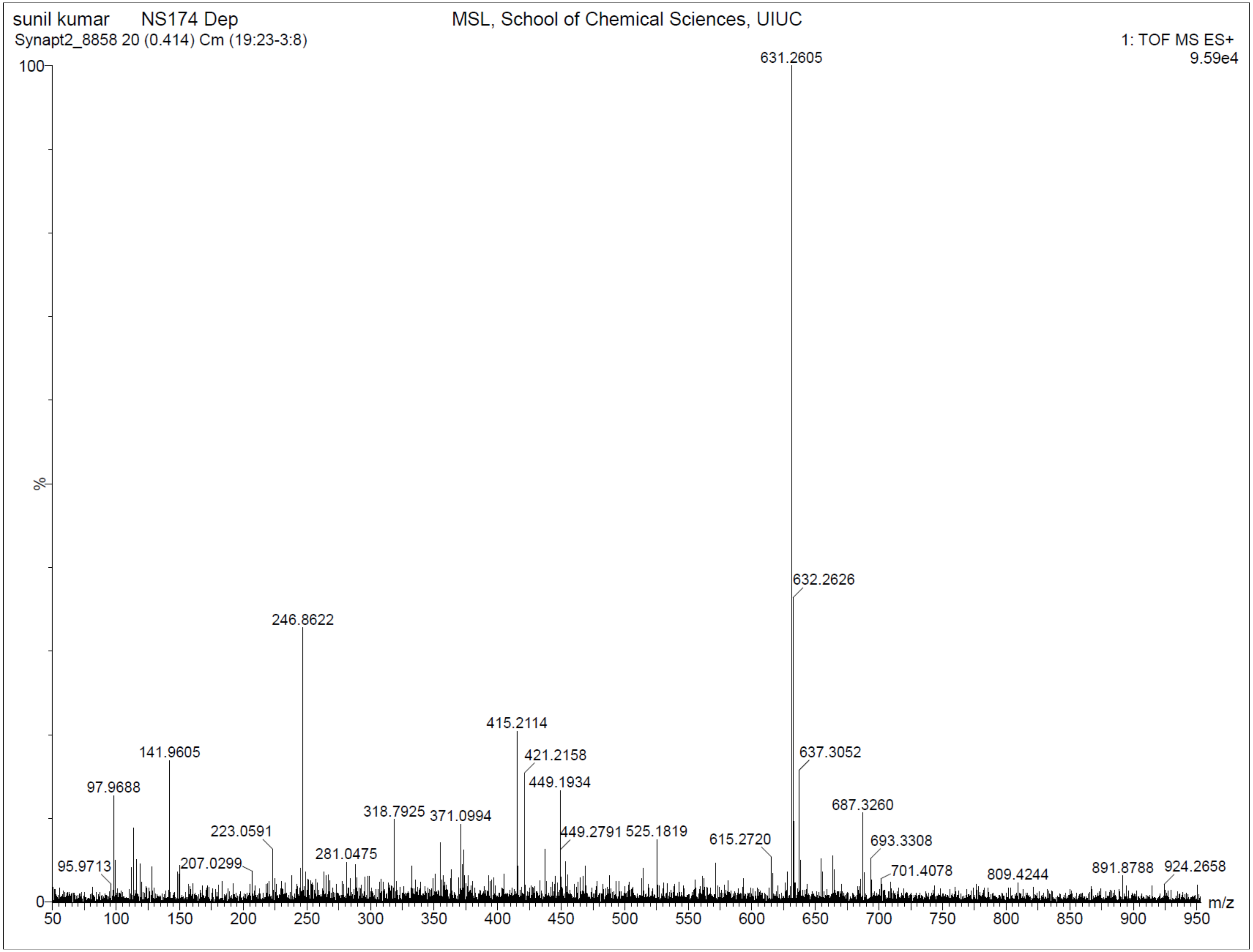

**Supplementary Fig. 124. $^{1}$H-NMR of NS163**

$^{1}$H NMR (500 MHz, DMSO) δ 0.99 – 1.33 (m, 4H), 1.39 – 1.80 (m, 6H), 2.30 – 2.38 (d, J = 4.0 Hz, 3H), 3.05 – 3.14 (d, J = 13.0 Hz, 2H), 3.97 – 4.11 (q, J = 6.7 Hz, 2H), 4.94 – 4.97 (s, 1H), 5.00 – 5.04 (s, 1H), 6.77 – 6.83 (dd, J = 9.7, 4.8 Hz, 1H), 6.92 – 6.97 (dd, J = 8.0, 4.6 Hz, 1H), 6.98 – 7.04 (dt, J = 9.3, 4.6 Hz, 1H), 7.20 – 7.23 (d, J = 2.5 Hz, 1H), 7.29 – 7.32 (d, J = 7.6 Hz, 1H), 7.35 – 7.38 (d, J = 8.4 Hz, 1H), 7.42 – 7.45 (dd, J = 7.7, 3.3 Hz, 1H), 7.54 – 7.58 (dd, J = 8.0, 3.2 Hz, 1H), 7.94 – 7.98 (d, J = 7.7 Hz, 1H), 8.56 – 8.65 (m, 3H), 9.83 – 9.86 (s, 1H), 10.32 – 10.39 (s, 1H), 10.83 – 10.85 (d, J = 2.7 Hz, 1H).HRMS (m/z): [M]+ calcd. for $C_{37}H_{41}N_{10}O_7$, 737.3160; found, 737.3156.

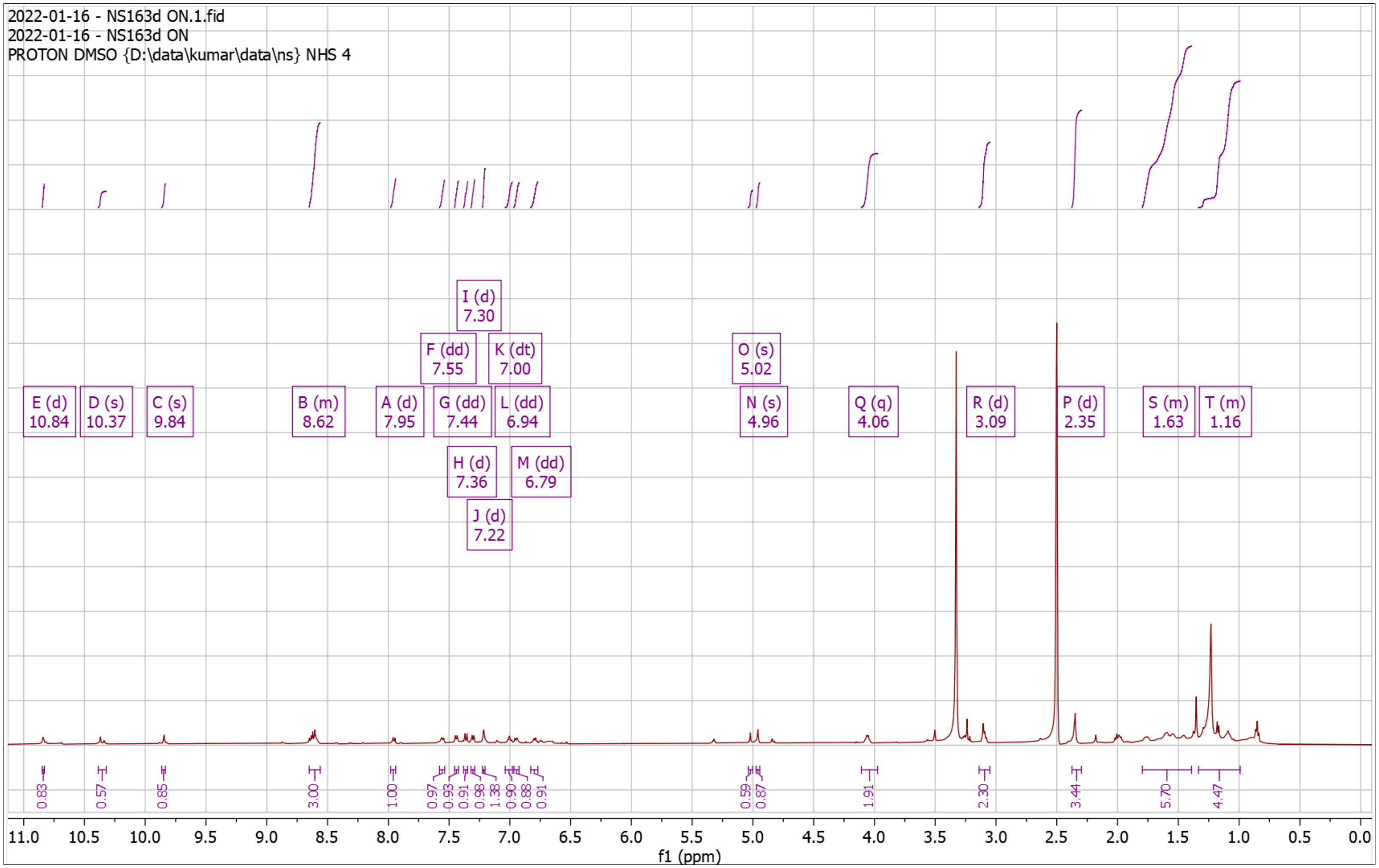


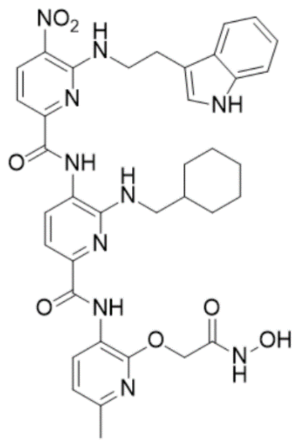

**Supplementary Fig. 125. High Resolution Mass Spectrum for NS163**

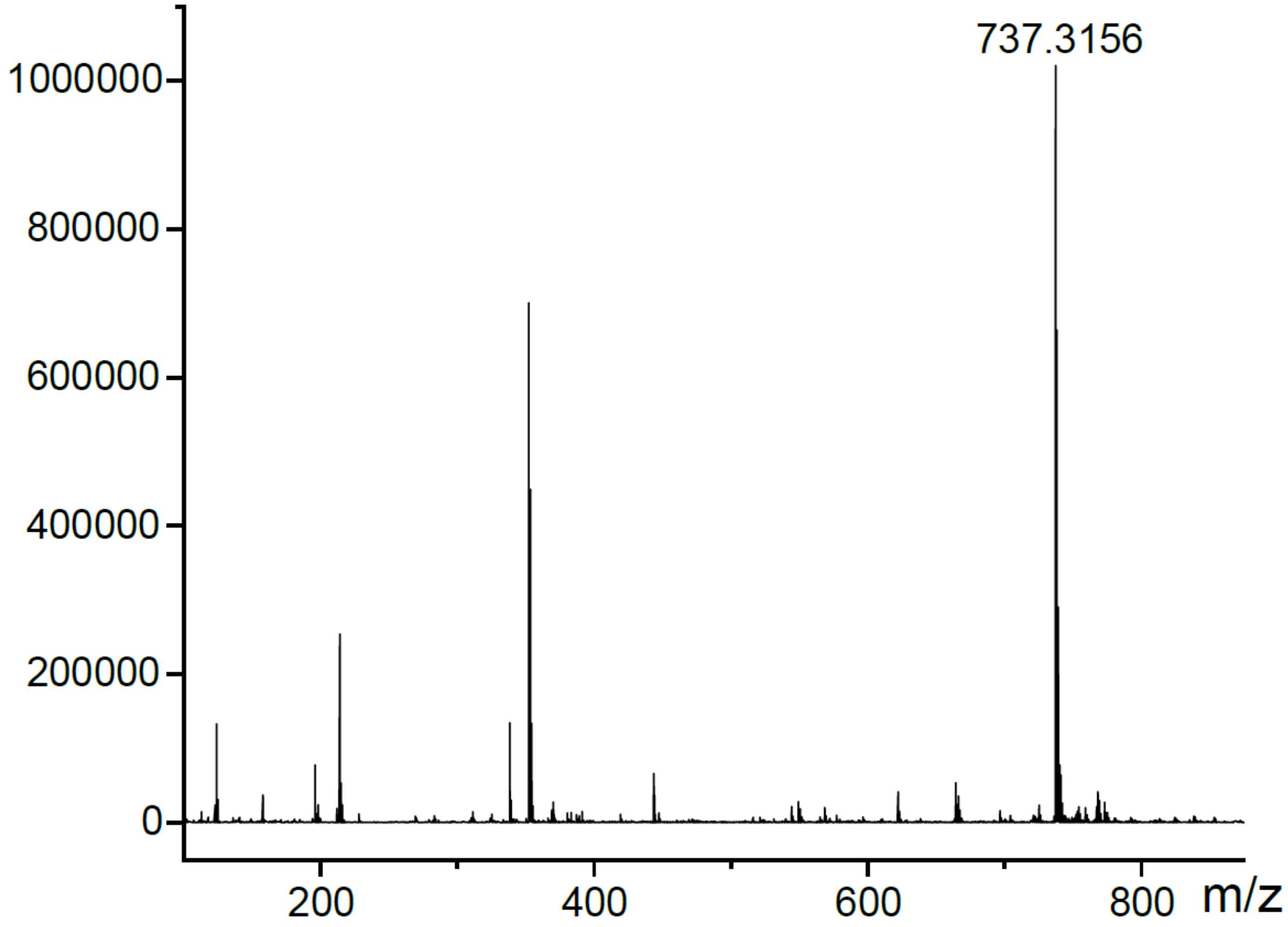

**References:**


1. Yadav, J. S., Balanarsaiah, E., Raghavendra, S. & Satyanarayana, M. Chemoselective hydrolysis of tert-butyl esters in acetonitrile using molecular iodine as a mild and efficient catalyst. *Tetrahedron Lett.* **47**, 4921–4924 (2006).

2. Wu, Y., Limburg, D. C., Wilkinson, D. E., Vaal, M. J. & Hamilton, G. S. A mild deprotection procedure for tert-butyl esters and tert-butyl ethers using $ZnBr_2$ in methylene chloride. *Tetrahedron Lett.* **41**, 2847–2849 (2000).

3. Huang, C., Ren, G., Zhou, H. & Wang, C. A new method for purification of recombinant human α-synuclein in Escherichia coli. *Protein Expr. Purif.* **42**, 173–177 (2005).

4. Ahmed, J. *et al.* Foldamers reveal and validate therapeutic targets associated with toxic α-synuclein self-assembly. *Nat Commun* **13**, 2273 (2022).

5. Eliezer, D., Kutluay, E., Bussell, R. & Browne, G. Conformational properties of α-synuclein in its free and lipid-associated states 1 1Edited by P. E. Wright. *J. Mol. Biol.* **307**, 1061–1073 (2001).

6. Harrington, A. J., Yacoubian, T. A., Slone, S. R., Caldwell, K. A. & Caldwell, G. A. Functional analysis of VPS41-mediated neuroprotection in Caenorhabditis elegans and mammalian models of Parkinson's disease. *J. Neurosci.* **32**, 2142–2153 (2012).

7. Brenner, S. The genetics of Caenorhabditis elegans. Genetics. **77**, 71–94 (1974).

8. Chaudhuri, J., Parihar, M. & Pires-daSilva, A. An introduction to worm lab: from culturing worms to mutagenesis. *J. Vis. Exp.* 2293 (2011).

9. Simonetta, S. Chemotaxis in *C.elegans* (ARENA system). *Phylumtech* https://www.phylumtech.com/home/en/chemotaxis-in-c-elegans-arena/ (2019).

10. Margie, O., Palmer, C. & Chin-Sang, I. C. elegans chemotaxis assay. *J. Vis. Exp.* 50069 (2013).

11. Garcia-Moreno, J. C., Porta de la Riva, M., Martínez-Lara, E., Siles, E. & Cañuelo, A. Tyrosol, a simple phenol from EVOO, targets multiple pathogenic mechanisms of neurodegeneration in a C. elegans model of parkinson's disease. *Neurobiol. Aging* **82**, 60–68 (2019).

12. Currey, H. N., Malinkevich, A., Melquist, P. & Liachko, N. F. Arena-based activity profiling of tau and TDP-43 transgenic C. elegans. *MicroPubl. Biol.* (2020).

13. Habchi, J. *et al.* An anticancer drug suppresses the primary nucleation reaction that initiates the production of the toxic Aβ42 aggregates linked with Alzheimer's disease. *Sci. Adv.* **2**, e1501244 (2016).

14. Yoon, D., Lee, M.-H. & Cha, D. Measurement of intracellular ROS in Caenorhabditis elegans using 2',7'-Dichlorodihydrofluorescein diacetate. *BIO-Protoc.* **8**, (2018).
