## Supplementary material for "A 2D Fragment-Assisted Protein Mimetic Approach to Rescue α-Synuclein Aggregation Mediated Early and Post-Disease Parkinson’s Phenotypes": Movies (S1-S8)

### Slide 1
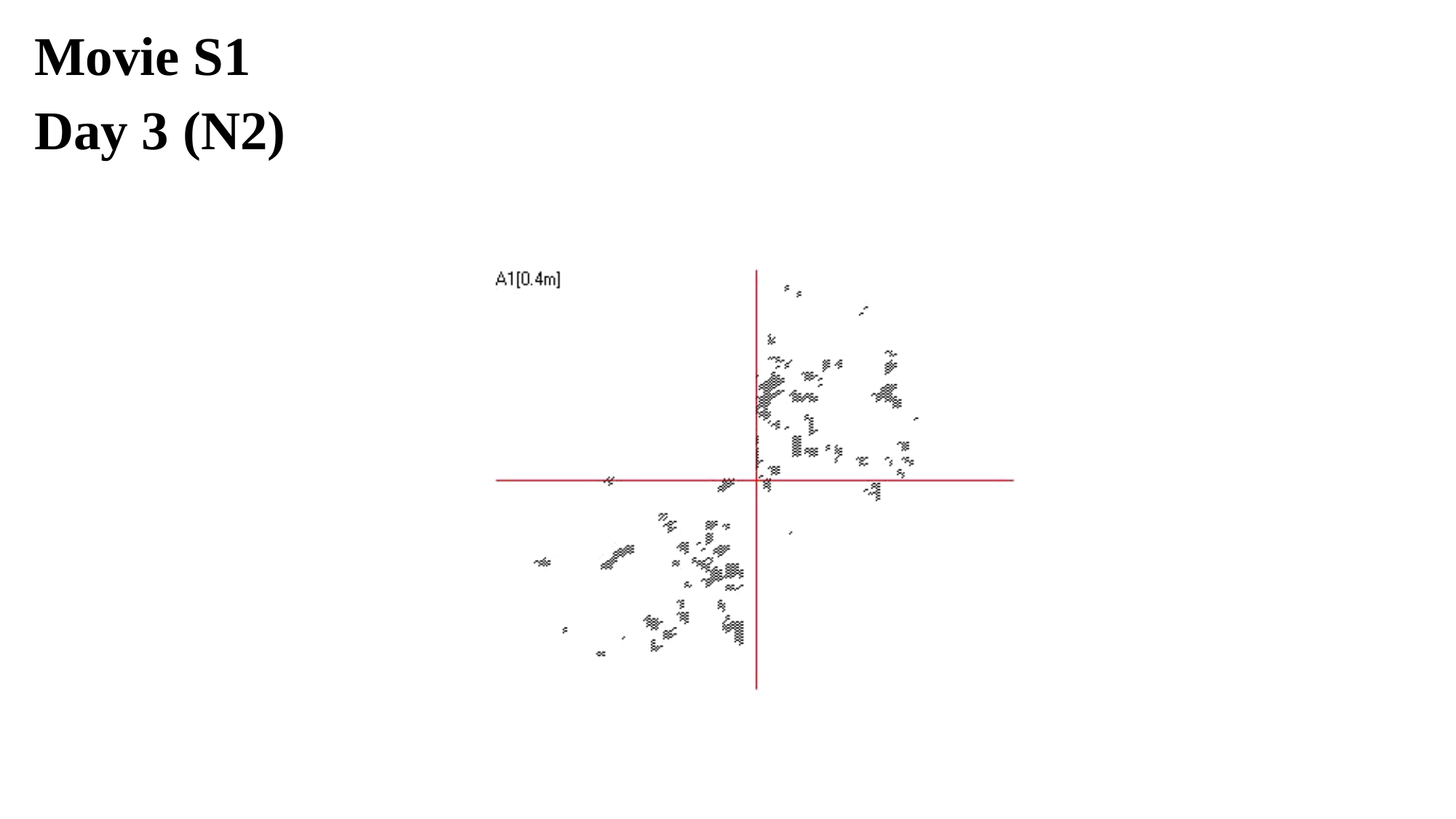

Movie S1
Day 3 (N2)

### Slide 2
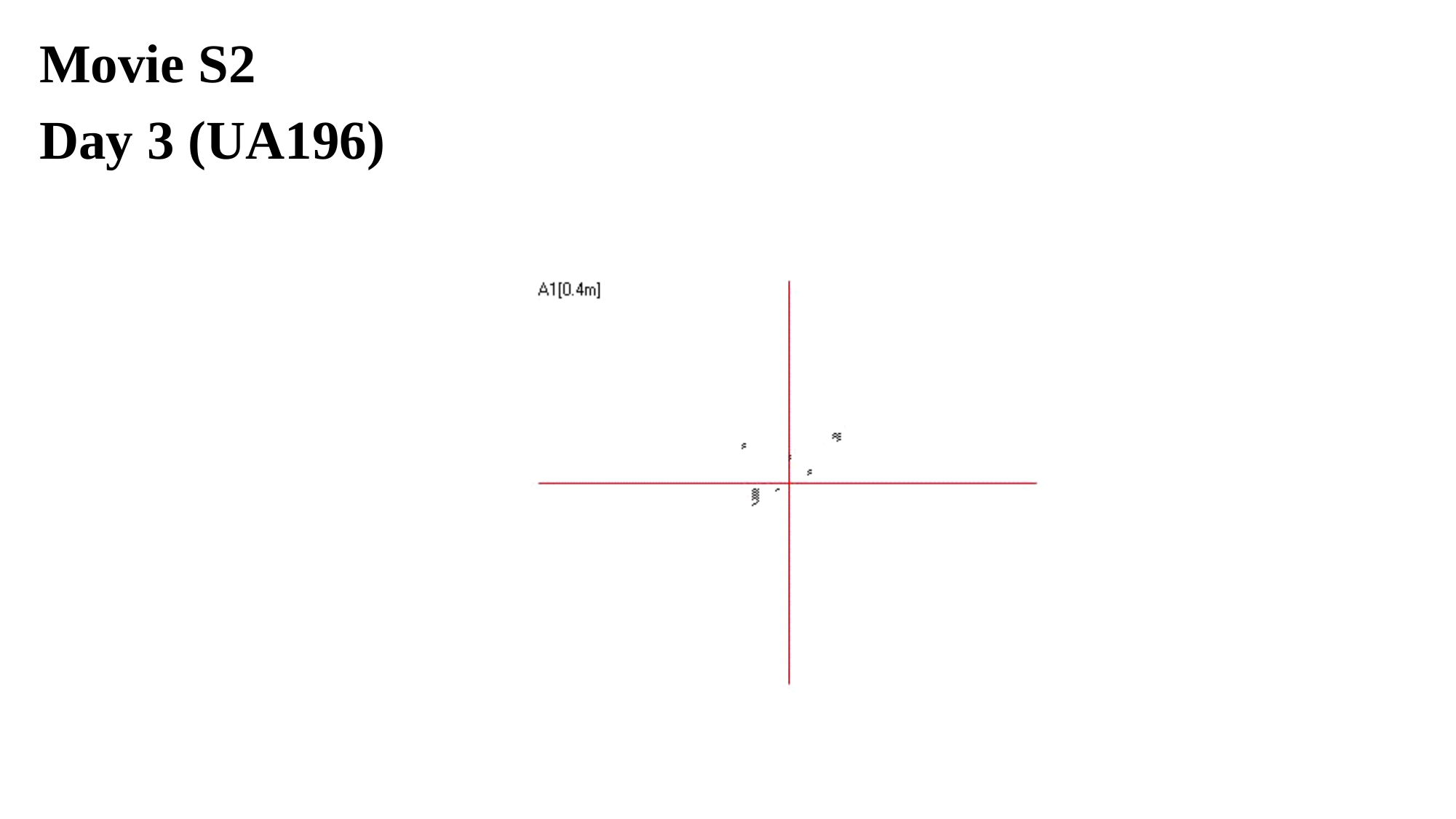

Movie S2
Day 3 (UA196)

### Slide 3
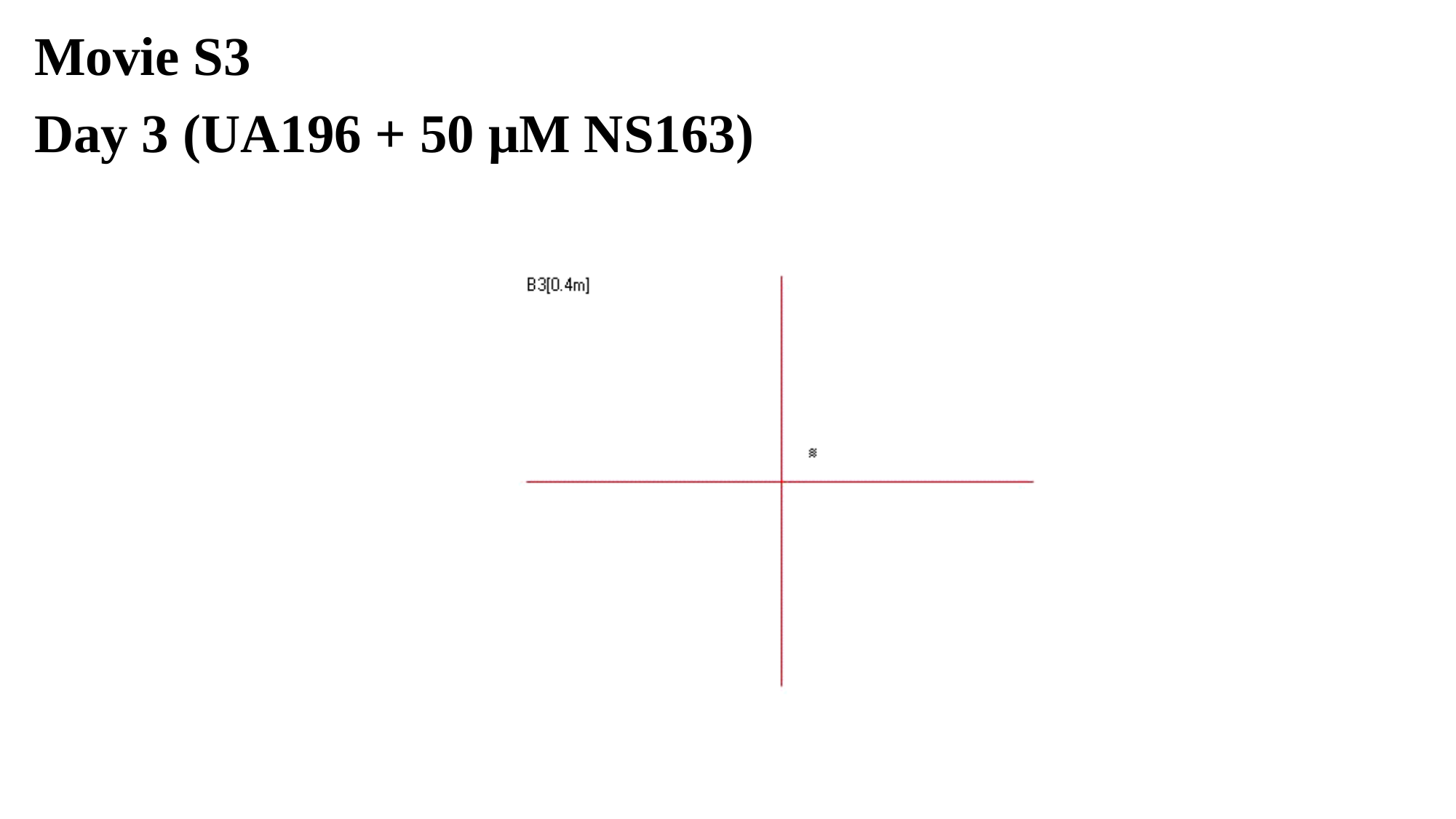

Movie S3
Day 3 (UA196 + 50 μM NS163)

### Slide 4
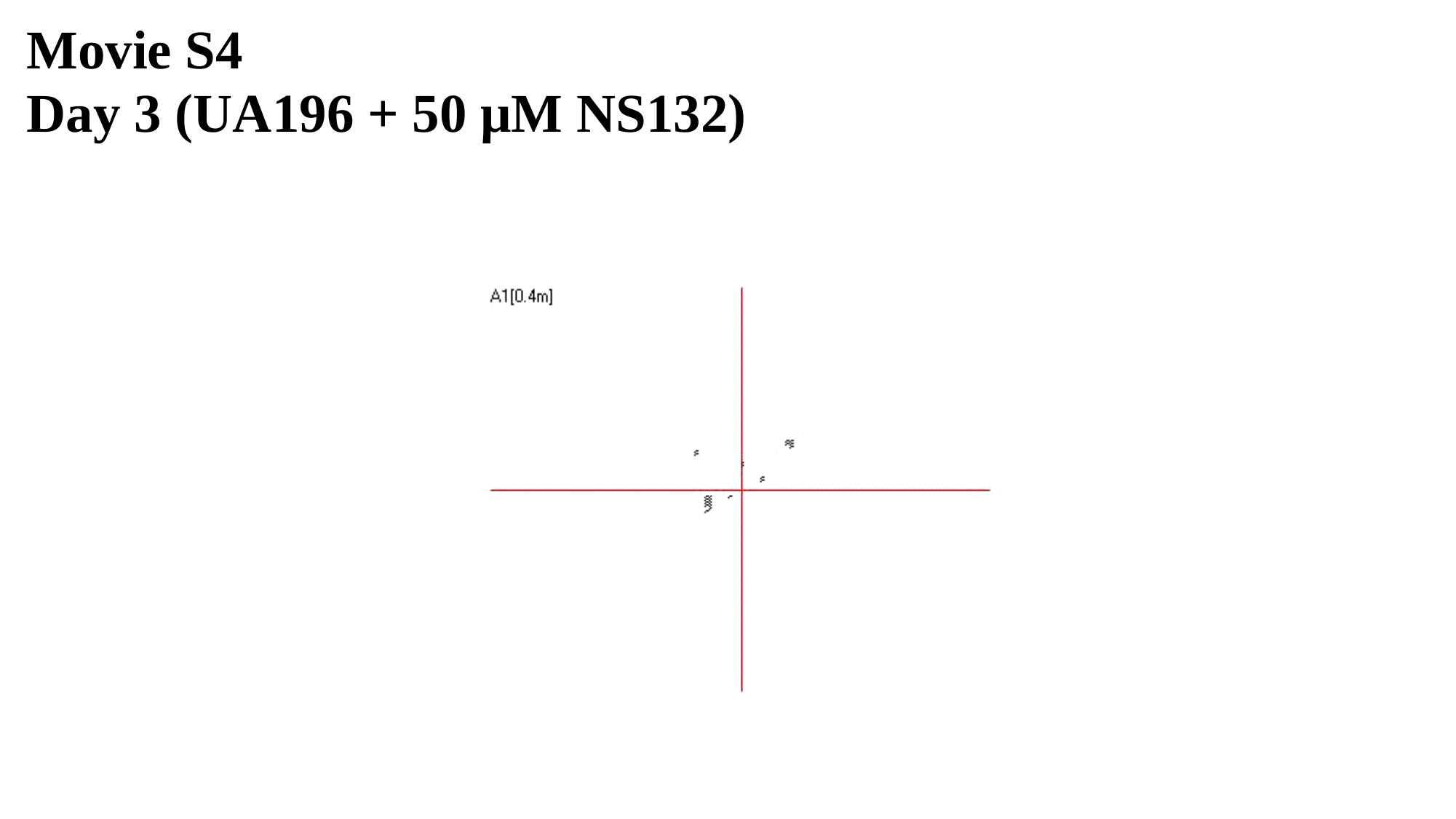

Movie S4
Day 3 (UA196 + 50 μM NS132)

### Slide 5
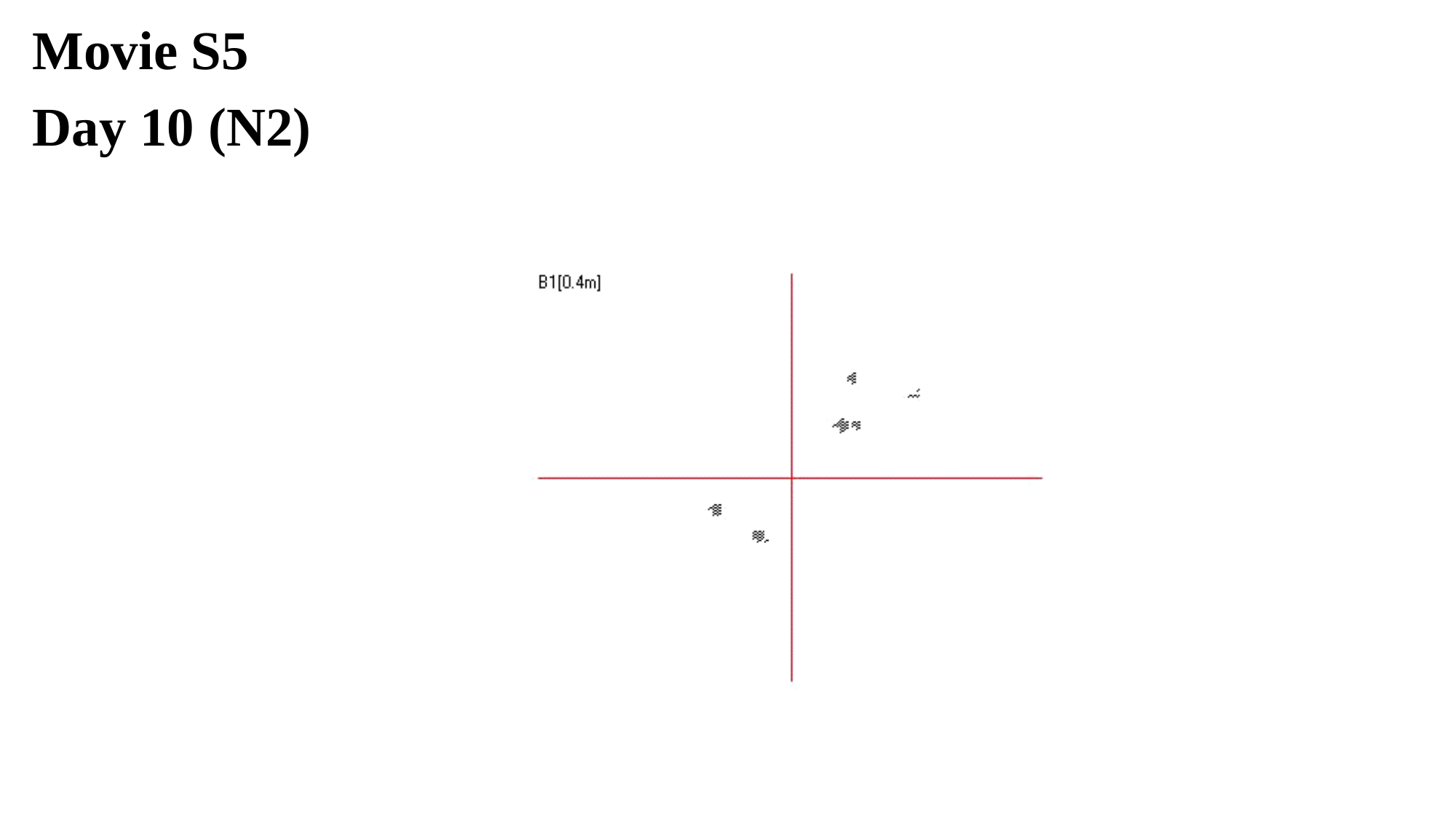

Movie S5
Day 10 (N2)

### Slide 6
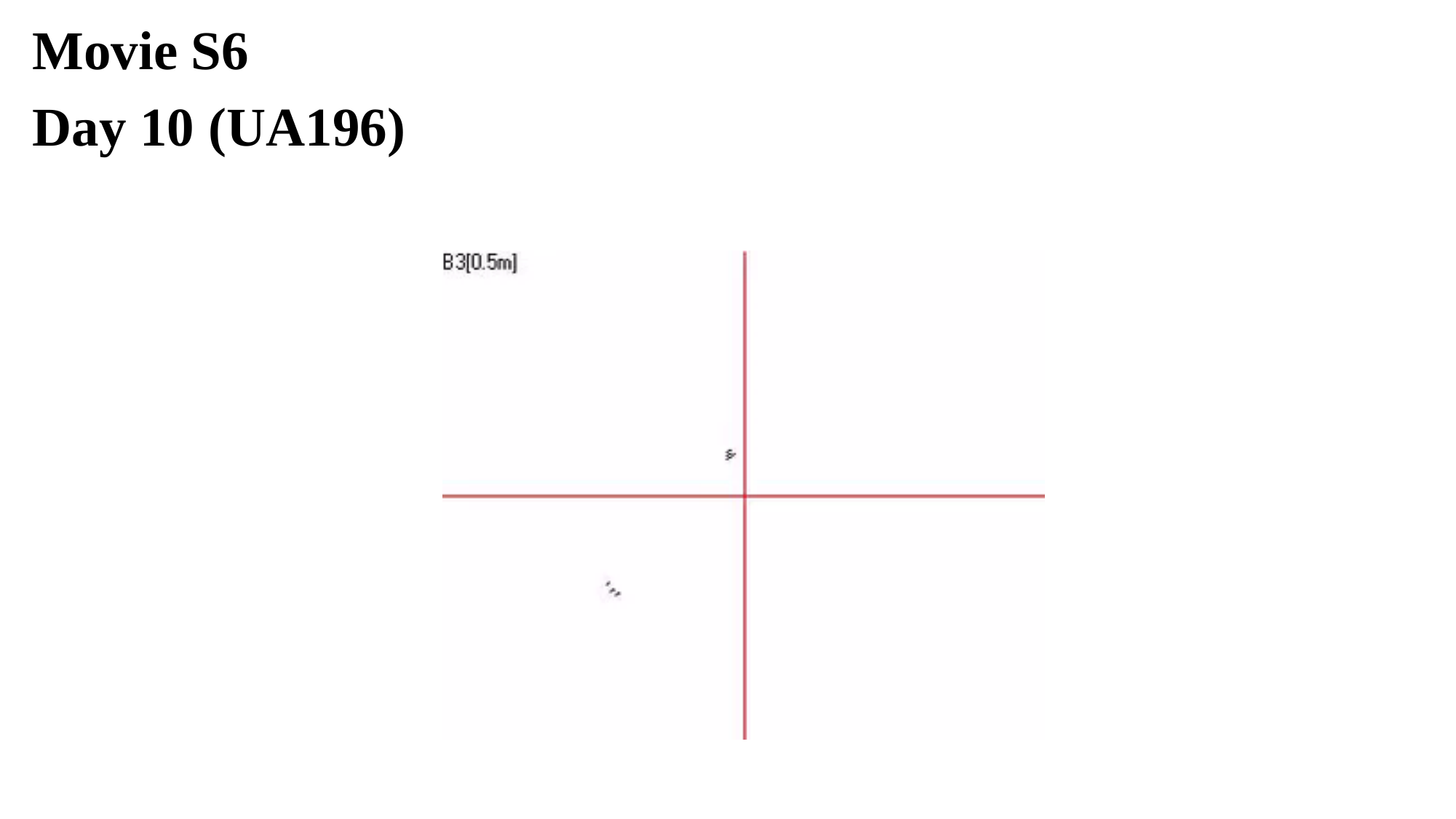

Movie S6
Day 10 (UA196)

### Slide 7
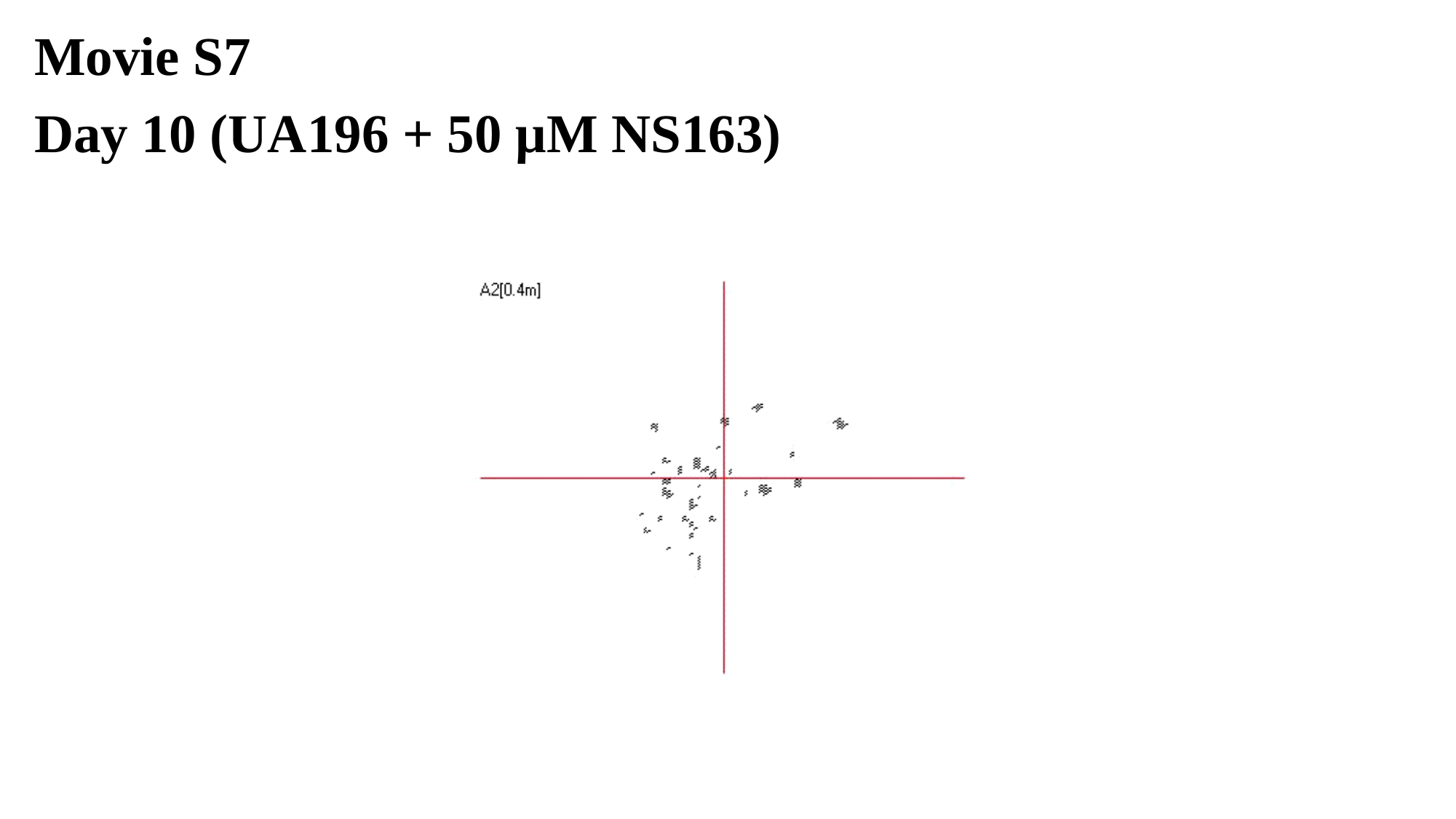

Movie S7
Day 10 (UA196 + 50 μM NS163)

### Slide 8
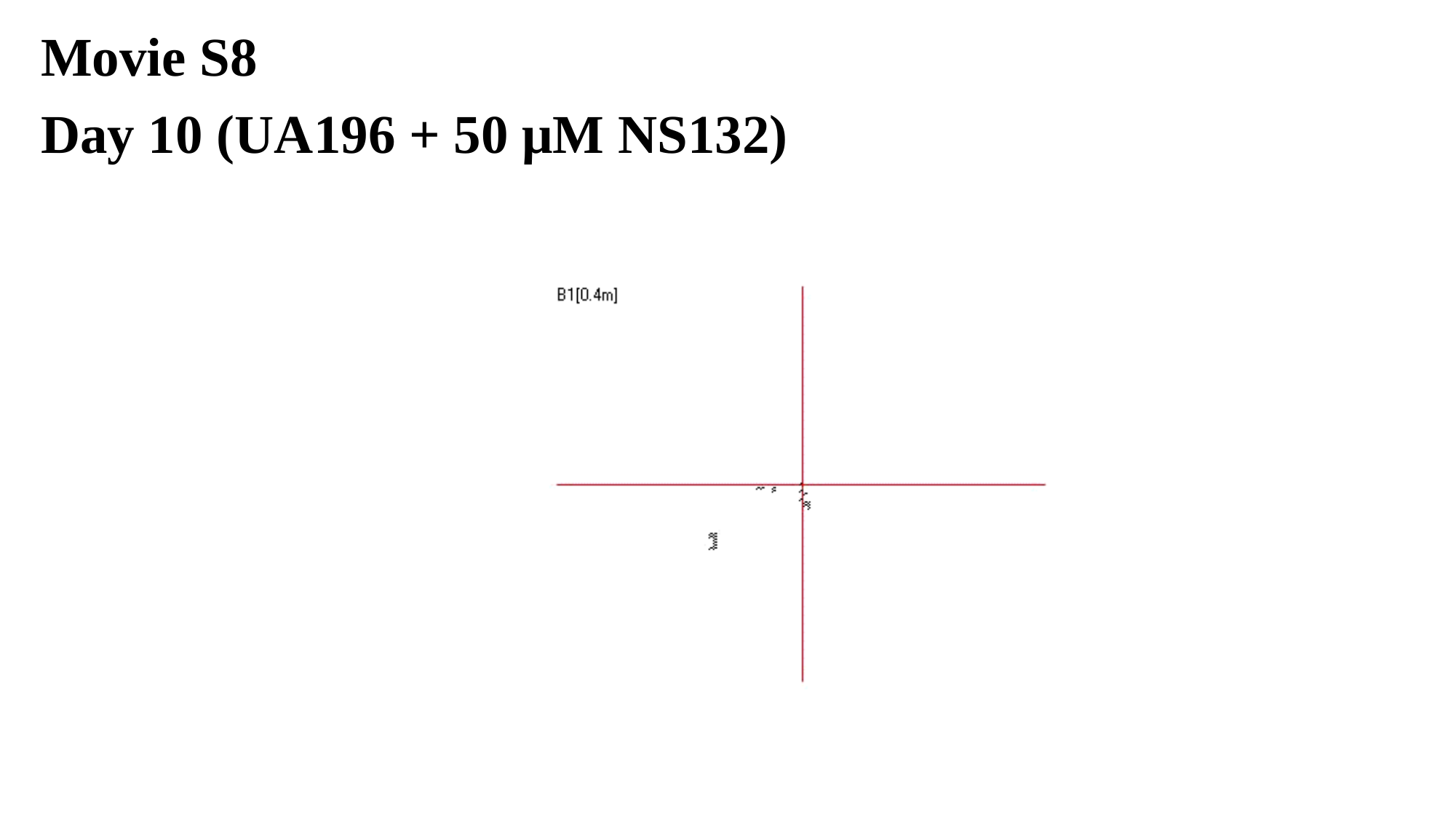

Movie S8
Day 10 (UA196 + 50 μM NS132)
